# Dual Stabilization of S-Adenosylmethionine for Enzymatic DNA Labeling

**DOI:** 10.64898/2026.01.19.700355

**Authors:** Jonas Bucevičius, Rūta Gerasimaitė, Gražvydas Lukinavičius

## Abstract

S-Adenosyl-L-methionine (AdoMet) analogues are powerful tool for site-specific biomolecular labeling via methyltransferase (MTase) catalyzed transfer reactions. However, their utility is often limited by poor chemical stability under enzymatic reaction conditions. Here, we report a new class of stabilized AdoMet analogues featuring a conformationally constrained proline side chain in place of homoalanine. This substitution inhibits intramolecular cyclization, a major decomposition pathway. Combination with selenonium modification, which suppresses depurination, yields analogues with up to a 90-fold increase half-life relative to AdoMet. These cofactors retain activity with DNA MTases, and allow sequence-specific labeling of plasmid DNA using both two-step and single-step approaches with fluorescent dyes.

## INTRODUCTION

S-adenosyl-L-methionine (AdoMet) is one of the most ubiquitous cofactors present in all living organisms, playing a central role in a wide array of biochemical reactions and intracellular regulatory pathways^1^. Methyltransferases (MTases) catalyze S_N_2-like reactions between nucleophiles in biomolecules and the electrophilic carbon next to the sulfonium center of AdoMet^2^. Transfer of larger substituents from AdoMet analogues is usually hampered, but the reaction efficiency can be restored by using synthetic analogues bearing sp- or sp^2^-hybridized carbon at the *β*-position to the sulfonium center^3–8^, which stabilizes the intermediate state and increases the transfer rate through a secondary orbital overlap effect. The transfer of larger groups from synthetic AdoMet-analogs by native or engineered MTases is exploited in late-stage modification of complex organic molecules^9–11^, biomolecule enrichment^12–14^, labeling^15, 16^, sequencing^12, 17, 18^ and optical genome mapping^19–21^. However, the AdoMet and especially its activated analogs are chemically unstable under enzymatic reaction conditions. They decay mainly via two main pathways: intramolecular cyclization leading to formation of homoserine lactone and 5⍰-deoxy-5⍰-(alkylthio)adenosine or its derivatives, and depurination resulting in adenine base and homocysteine sulfonium ribose derivatives^22^ (Figure 1).

**Figure 1.**
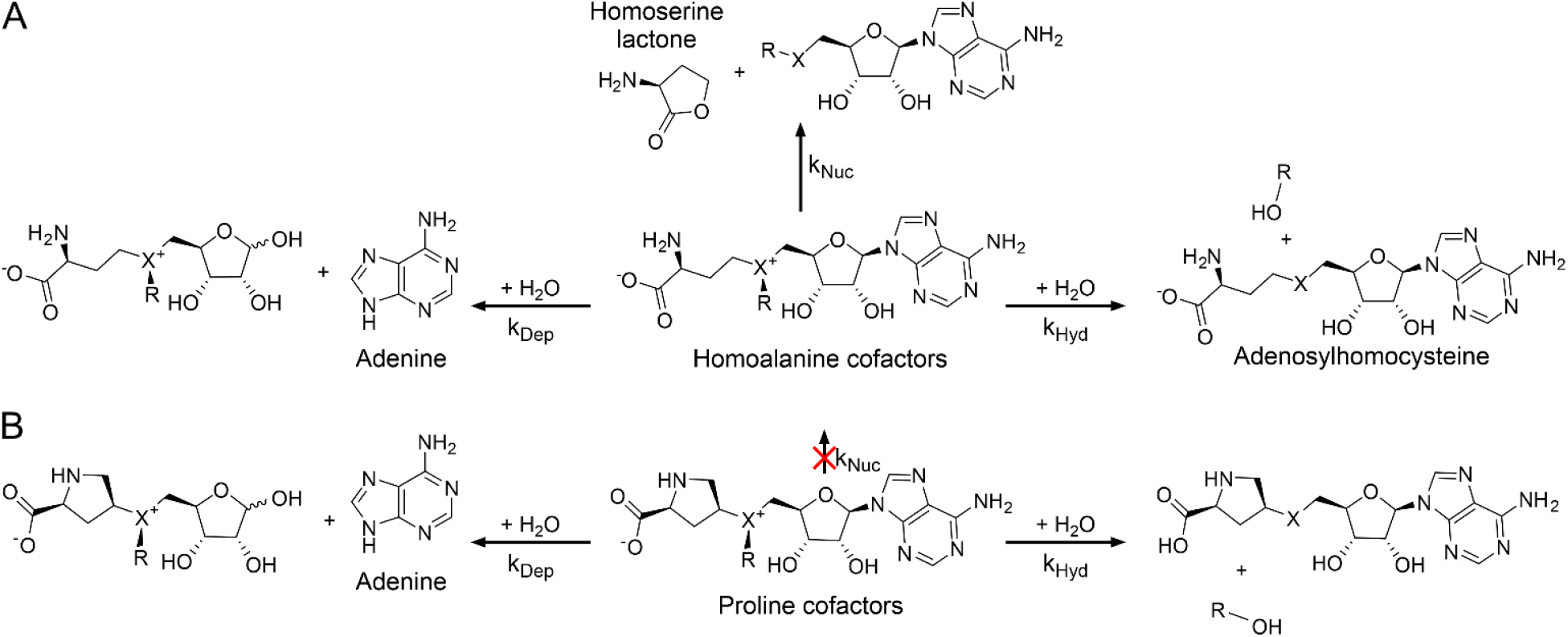
Major decomposition pathways of homoalanine (A) and proline (B) cofactors. Decomposition at physiological pH of AdoMet and its analogs can proceed via multiple pathways depending on structure of transferable side chain (R) and chalcogenonium center (X). Dominating decay products allow estimation of each pathway rate: k_Nuc_ – intramolecular nucleophilic cyclization, k_Dep_ – depurination and k_Hyd_ – hydrolysis. Replacement of homoalanine with proline introduces steric hindrance and effectively eliminates intramolecular cyclization.

Since the first examples of Methyltransferase-Directed Transfer of Activated Groups (mTAG)^3–5^, there have been only a few reports of chemically more stable designs of AdoMet analogs, but the enzymatic activity was either lost^23^ or maintained only with promiscuous smallmolecule transferases such as DnrK, COMT, PNMT, NNMT and none of them maintained activity with DNA MTases^11, 24–27^. Herein we report an AdoMet analog design wherein homoalanine side chain is replaced with a conformationally rigid proline side-chain eliminating intramolecular cyclization decay pathway. When combined with sulfur-to-selenium substitution, that reduces depurination at higher pH, the modified cofactors display up to 90-fold enhanced stability under enzymatic reaction conditions. We demonstrate that these cofactor analogues maintain enzymatic activities with the widely used thermophilic adenine DNA MTase TaqI and an engineered cytosine DNA MTase HhaI.

## RESULTS AND DISCUSSION

### Design, Synthesis, and Initial Enzymatic Activity Testing

We hypothesized that replacing the homoalanine side chain with a proline residue would inhibit the intramolecular cyclization pathway by restricting access to a favorable reactive conformation. To explore this, we synthesized a series of AdoMet analogs incorporating all possible configurations at the chiral centers of the proline-derived side chain (Scheme 1 and Figures S1-S4) and evaluated their activity with the DNA methyltransferases: TaqI and HhaI Q82A/Y254S/N304A variant, engineered to work efficiently with activated AdoMet analogs^28^. The synthesis commenced from enantiomerically pure N-Boc-4-hydroxyproline methyl esters (Scheme 1). The hydroxy group was converted to a mesylate, followed by S_N_2-type nucleophilic substitution with either potassium thioacetate (KSAc) or potassium selenocyanate (KSeCN), affording intermediates **5–9**. Subsequent generation of the corresponding thiolate or selenolate anions enabled nucleophilic substitution of either 5⍰-tosyladenosine or 5⍰-chloro-7-deazaadenosine to yield compounds **10–15**. After ester hydrolysis and Boc deprotection, the resulting compounds **16–21** were alkylated by using mesylates (**SI-1**-**SI-3)** or triflates (**SI-4** and **SI-5)** to furnish diastereomeric mixtures of the final cofactors **22–36**. Obtained diastereomeric mixtures were separated by preparative reverse-phase HPLC (Figure S5). In case of cofactors with methyl transferable group (**28ab**-**30ab**) we were unable to separate diastereomers, therefore these compounds were tested and used as two diastereomer mixtures. In all other cases, separations were successful.

**Scheme 1.**
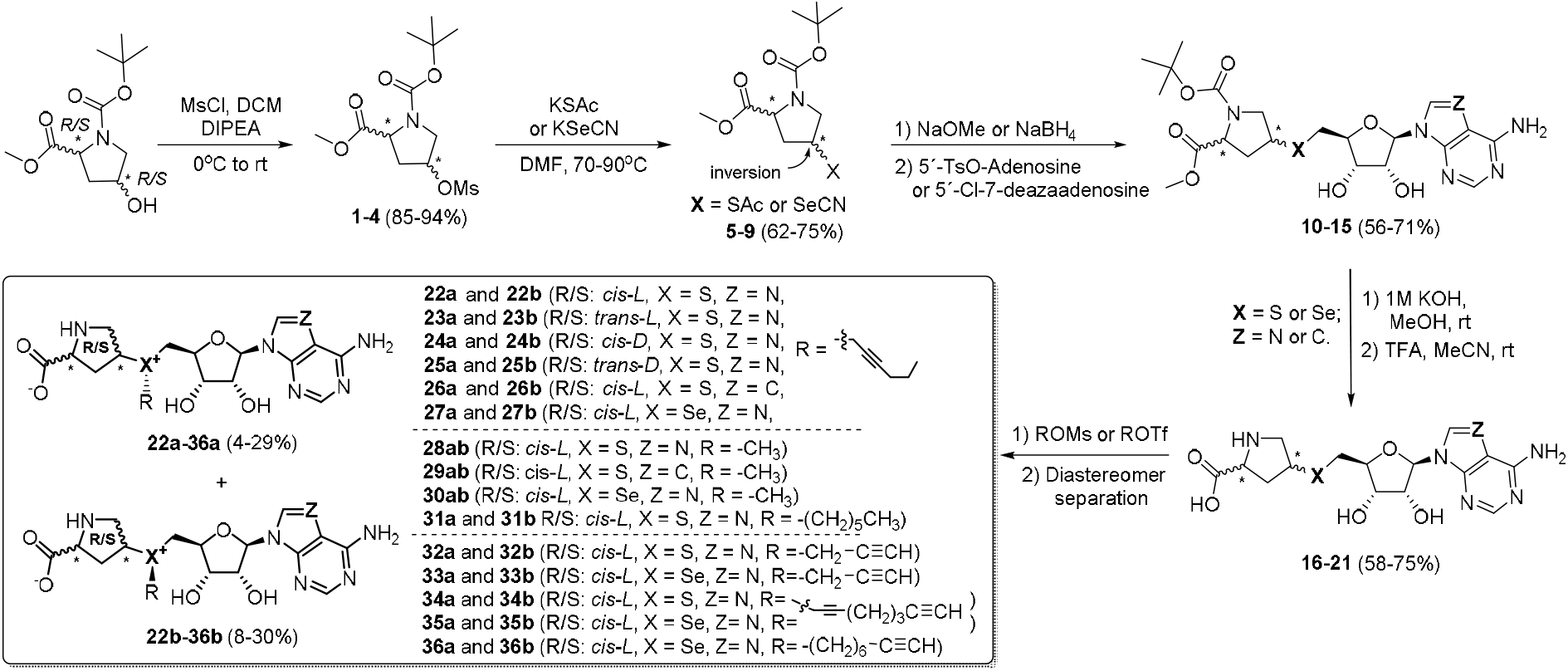
Synthesis scheme of proline cofactors 22-36.

The activity of the new AdoMet analogs in DNA MTase–catalyzed reactions was evaluated using a DNA protection assay, which relies on the inability of restriction enzymes to cleave DNA containing modified bases within their recognition sequences (Figure S6). Cofactors **22a**– **25a** and **22b**–**25b**, bearing a transferable hex-2-ynyl side chain, were tested with the thermophilic DNA adenine-N6 methyltransferase TaqI^3^. Among these, only the *cis*-*L*-proline and *cis*-*D*-proline analogs **22b** and **24b** exhibited enzymatic activity (Figure S7). By analogy with the active stereoisomer of AdoMet, the sulfonium or selenonium center in these cofactors is presumed to adopt the R configuration. We further assume that all other cofactors exhibiting longer retention times (**26b, 27b, 31-36b**) during preparative HPLC purification the same R configuration. The *cis*-*L*-proline analog **22b** was also active with the M.HhaI Q82A/Y254S/N304A variant (Figure S8). On this basis, all subsequent analogs were designed around the *cis*-*L*-proline scaffold as more promising variant.

For comparative analysis, sulfonium- and selenonium-based cofactors (**37b**-**40b**) bearing the canonical homoalanine side chain and a series of transferable groups (methyl, propargyl, and hex-2-ynyl) were prepared (Figure 2A). The synthesis involved alkylation of Sadenosylhomocysteine or Se-adenosylhomocysteine with the corresponding alkyl mesylates or methyl triflate (Figure S9) followed by preparative HPLC purification.

**Figure 2.**
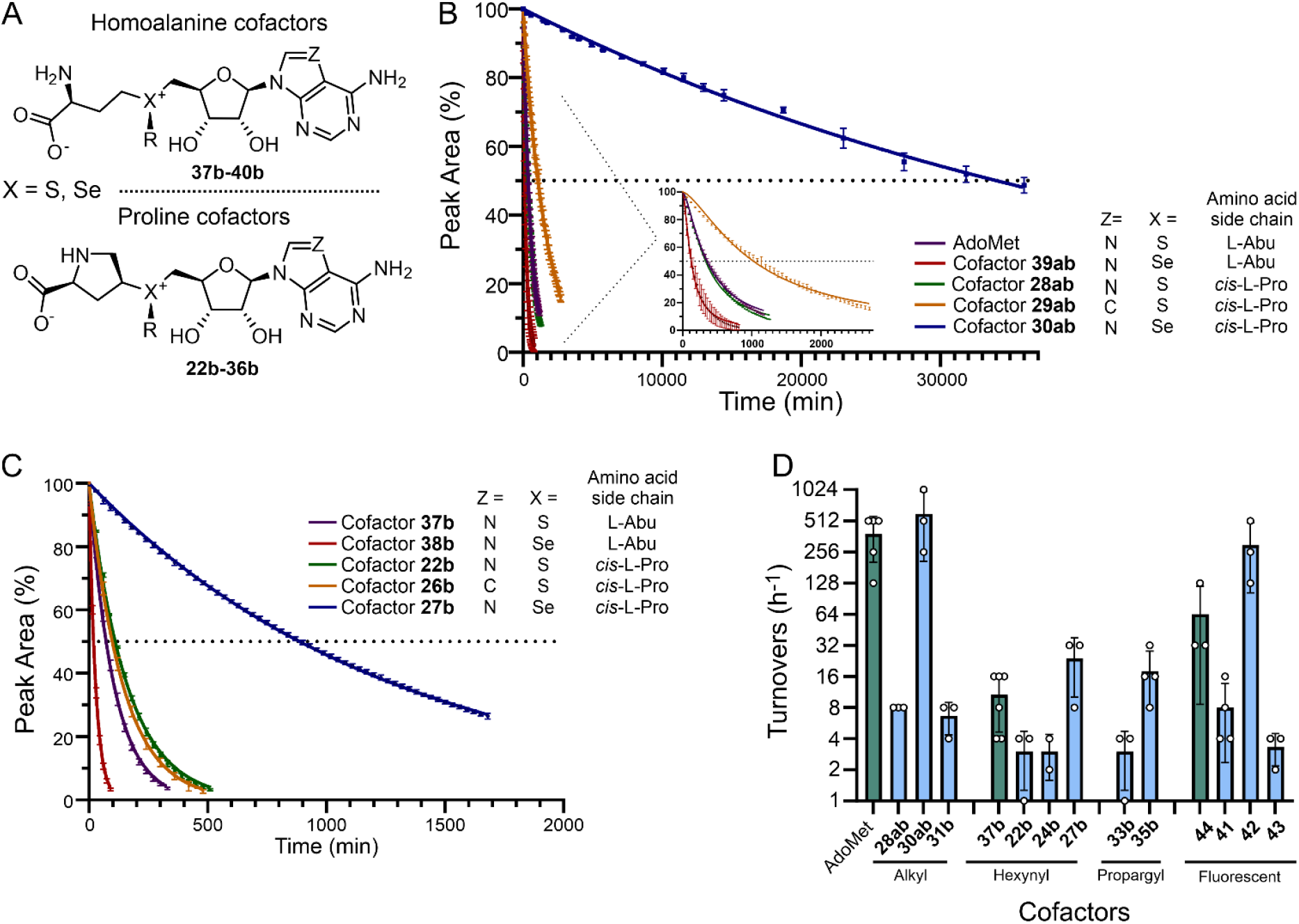
Stability and activity of homoalanine and proline cofactors. (A) General structures of homoalanine and proline cofactors. Degradation kinetics of AdoMet or its analogues containing transferable methyl (B) or hex-2-ynyl groups (C) in 50 mM Tris-HCl (pH 8.0) at 37°C, N = 3. (D) Apparent turnover numbers of M.TaqI with homoalanine (green bars) and proline (blue bars) cofactors measured via DNA protection assay in 50 mM Tris-MOPS (pH 8.0) at 65°C, N ≥ 3.

For single-step labeling of DNA with fluorescent dyes, we synthesized *cis*-*L*-proline cofactors **41**–**43** by conjugating terminal alkyne-bearing intermediates **34b**-**36b** with 4-MeSiR-PEG_4_-N_3_ via CuAAC reaction, introducing a fluorescent reporter directly into the transferable moiety in a 30-67% yields. For comparison, a classical homoalanine-based cofactor analog **44** was prepared similarly from AdoHcyN_3_^5^ and 4-MeSiR-PEG_4_-alkyne (Figure S10).

### Stability, Degradation and Kinetics

The chemical stability and decomposition products of prepared cofactors were evaluated in 50 mM Tris– HCl buffer (pH 8.0) at 37 °C (Figure 2). Time-dependent decomposition analysis revealed behavior consistent with first-order decay kinetics, indicative of (pseudo)unimolecular decomposition processes (Figures 2, S11– S15, and Table 1). AdoMet exhibited a half-life of 366 minutes (Figure 2B), with equal contributions from intramolecular cyclization and depurination pathways (Table 1 and Figure S16). Substitution of the sulfonium center with selenonium (SeAdoMet^22^ **39ab**) suppressed the depurination pathway (Figure S16), but accelerated intramolecular cyclization due to the higher nucleofugality of diselenyl ethers^29^, resulting in a reduced half-life of 133 minutes. We noticed that the decomposition product, methylselenoadenosine, was partially oxidized to methylselenoxideadenosine under the analytical conditions^30^, and this oxidation was taken into account in the half-life and degradation reaction rate calculations. Importantly, the selenium incorporation completely suppressed depurination, as the formation of a selenium ylide intermediate requires higher than physiological pH values^22, 31^.

**Table 1.**
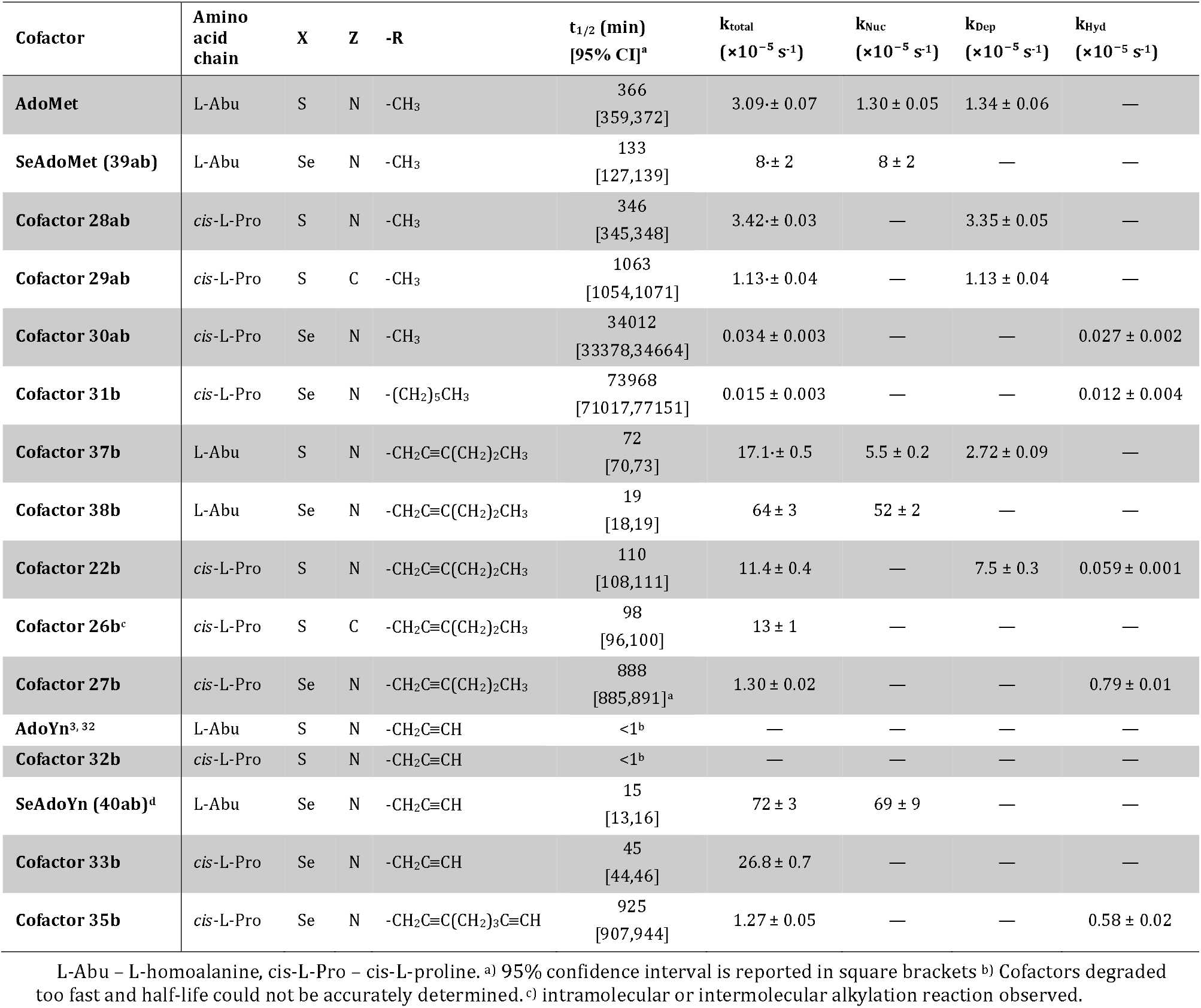
Half-lives and decay rate constants of the studied cofactors.

Replacing the homoalanine side chain with a proline moiety completely inhibited the intramolecular cyclization pathway (Figure S17), but the depurination rate slightly increased and the analog **28ab** showed similar half-life (346 minutes, 5.8 hours) to AdoMet. It is known that the use of a purine isostere, such as 7-deazapurine, can suppress depurination^24^ at physiological pH values due to limited cascade chargeredistribution from formed ylide to carbon at 7 position. Indeed, compound **29ab** exhibited a markedly extended half-life of ~17 hours, with slow depurination as the dominant degradation pathway (Table 1 and Figure 2B).

Remarkably, the selenium-containing cofactor **30ab** displayed complete suppression of both depurination and cyclization (Table 1, Figure 2B and S17), leading to a halflife of ~24 days — 90-fold greater than that of native AdoMet. Here, the only remaining decomposition route was slow hydrolysis of the transferable methyl group.

Next, we measured the stabilities of cofactors bearing activated hex-2-ynyl groups. Half-life of homoalanine containing cofactor **37b** was 72 min with a major degradation by intramolecular cyclization and minor by depurination (Figure 2C, S11 and S16). As expected, the corresponding selenonium cofactor **38b** exhibited markedly accelerated degradation, proceeding predominantly via an intramolecular cyclization pathway with a shorter half-life of 19⍰min (Figures 2C, S11 and S16). The corresponding proline-containing cofactor **22b** exhibited only a marginal increase in half-life compared to its homoalanine counterpart cofactor **37b**, consistent with their similar depurination rate constants of the order of 10^−5^ s^-1^ (Figures S11 and S17, Table 1). Incorporation of a 7-deazapurine (cofactor **26b**) did not improve stability, likely due to increased lability of the hex-2-ynyl group resulting in its intramolecular or intermolecular transfer onto proline nitrogen (Figure S17 and S18). However, the selenonium-proline cofactor **27b** displayed significantly improved stability of ~14 hours, with degradation predominantly occurring through hydrolysis of the transferable group (Figure S17 and S18). This represents a 12-fold increase in half-life compared to the conventional homoalanine-based analog **37b** (72 minutes).

Both the sulfonium proline cofactor **32b** and AdoYn^3, 32^, were highly unstable and rapidly degraded via hydration of the terminal alkyne, with complete decomposition occurring in under 1 min (Table 1). The cofactor **40b** (SeAdoYn) is widely used for tagging proteins, DNA or RNA with a reactive propargyl group for downstream labeling by click chemistry^18, 32–34^. We determined the half-life of this cofactor to be approximately 15 minutes due to very fast intramolecular cyclization (Figure S16 and S18). In comparison, the selenonium-proline cofactor **33b** exhibited improved stability, with a half-life of ~45 minutes (Figures S11 and S17 and Table 1) and degraded via water addition to triple bond, thus significantly slower compared to AdoYn or **32b**. Introduction of proline eliminated intramolecular cyclization pathway, but did not chang susceptibility to water addition at the triple bond. We propose that selenonium salts possess a substantially higher activation barrier to ylide formation at the propargyl substituent, a process that in sulfonium analogs rapidly triggers rearrangement to a highly reactive allene intermediate that is readily attacked by water.

An alternative strategy to overcome propargyl cofactor instability is to position the terminal alkyne distal to the chalcogenonium center^5^. Thus, we designed cofactor **35b**, featuring a transferable n-octadiynyl group—an eight-carbon linker terminating in a terminal alkyne which can be used via click chemistry for 2-step labelling or pulldown applications. This design retained similar halflife of ~15h and degradation kinetics to cofactor **27b**, which contains a shorter hex-2-ynyl side chain, and showed no degradation via water addition to triple bond (Figure S17).

### Enzymatic activity with Methyltransferases TaqI and HhaI Q82A/Y254S/N304A

The efficiency of enzymatic transfer of methyl or triple-bond–activated substituents (R groups) was estimated using a DNA protection assay, by varying the molar ratio of DNA methyltransferase to its target sites. Reactions contained a constant amount of λ DNA and two-fold serial dilutions of M.TaqI, and apparent turnovers were determined based on digestion with R.TaqI followed by agarose gel electrophoresis (Figure 2D, S6 and S19). AdoMet served as a benchmark, showing ~512 h^−1^ turnovers. The analog cofactor **28ab**, bearing a proline side chain, exhibited only 8 h^−1^. This might be caused by multiple reasons, including decreased affinity of the cofactor due to the loss of contacts with the protein, steric hindrance and distorted geometry of the transition state due to conformational rigidity of the proline side chain. Remarkably, the selenonium analogue **30ab** with the same proline side chain reached ~512 h^−1^, suggesting that the enhanced electrophilicity of selenium and markedly increased cofactor stability can effectively counteract all mentioned negative effects. The same trend was observed for triple bond activated cofactor series: the benchmark homoalanine cofactor (**37b**) with hex-2-ynyl transferable group attained ~10 h^−1^, while *cis*-L/D-proline analogues **22b** / **24b** showed reduced activity (~4 h^−1^). The selenonium-proline cofactor **27b** exhibited 32 h^−1^ turnovers, representing a significant improvement over homoalanine cofactor **37b**. Importantly, this effect was not limited to short transferable chains, but was retained in fluorescent cofactors, where the fluorescent dye is linked via long hydrophilic PEG4 linker. Here again, selenonium-proline cofactor **42** significantly outperformed sulfonium-proline cofactor **41** and homoalanine cofactor **44**. In fact, M.TaqI activity with cofactor **42** approached its activity with AdoMet (Figure 2D). Longer alkyl carbon chain substituted cofactors, such as -propyl, without activating π-bonds generally result in minimal (<1h^-1^) transfer efficiencies^3, 35^. Nevertheless, the most stable analogue **31b** (halflife ~50 days), bearing a hexyl transferable group, achieved ~8 h^−1^ turnovers with M.TaqI, underscoring the increased reactivity of selenonium centers even without sp or sp^2^ hybridization at the β-position^29^ (Figures 2D and S19).

Similar trend was observed with M.HhaI Q82A/Y254S/N304A and cofactors with transferable methyl group. Activity of proline cofactor **28ab** decreased compared to AdoMet, which was partially rescued using selenonium-proline cofactor **30ab**. However, this effect was not retained with longer transferable chains: while cofactor **22b** showed low activity, cofactor **27b** was completely inactive (Figure S20). This behavior likely results from the stringent geometric requirements for formation of the intermediate covalent complex in C5-MTases, which cannot simultaneously accommodate the constrained proline side chain and the larger selenium atom.

Next, we confirmed the nature of enzymatic reaction products in the MTase reaction under a single-turnover conditions using duplex DNA oligonucleotide bearing a single MTase site and excess of enzyme and cofactor. The DNA was then hydrolyzed to nucleosides, and LC/MS analysis was used to identify the resulting modifications. Expected identities of modification products by M.TaqI (Figures S21 and S22) and M.HhaI Q82A/Y254S/N304A (Figure S23) were confirmed by mass spectrometry in all the cases. The results corroborate the conclusion that both DNA MTases can use stabilized *cis*-L-proline sulfonium and selenonium cofactors to modify DNA.

Under single turnover conditions, M.TaqI reaction efficiency is unlikely to be limited by affinity towards the cofactor. Nonetheless, sulfonium-proline cofactors **22b, 24b**, and **41** and selenonium-proline cofactors **31b** and **43**, which lack activating π-bonds, fail to modify both DNA strands. This suggests a challenge in accommodating the conformationally rigid cofactor and the bulky modification on the complementary strand of the DNA target site in a productive orientation within the enzyme active site.

### Two-Step and Direct Labeling of DNA with fluorescent dyes

To demonstrate the applicability of proline-based cofactors for DNA labeling, we carried out two-step and single-step sequence-specific labeling of pUC19 DNA (Figure 3). In the two-step approach, proline cofactor **35b**, bearing an n-octadiynyl transferable group, was used. In the first step, the terminal alkyne-containing group was enzymatically transferred to pUC19 DNA by M.TaqI, and complete modification of TCGA sites was confirmed by protection from R.TaqI cleavage (Figure S24). In the second step, the resulting alkyne-tagged DNA was conjugated with Calfluor647-azide^36^ under CuAAC conditions. To assess labeling specificity, the modified DNA was digested with R.MbiI and analyzed by agarose gel electrophoresis. Fluorescence imaging in the Cy5 channel revealed selective labeling of the 1801 bp and 644 bp fragments—each containing two M.TaqI sites and no labeling on 240 bp fragment without M.TaqI sites (Figure 3A).

**Figure 3.**
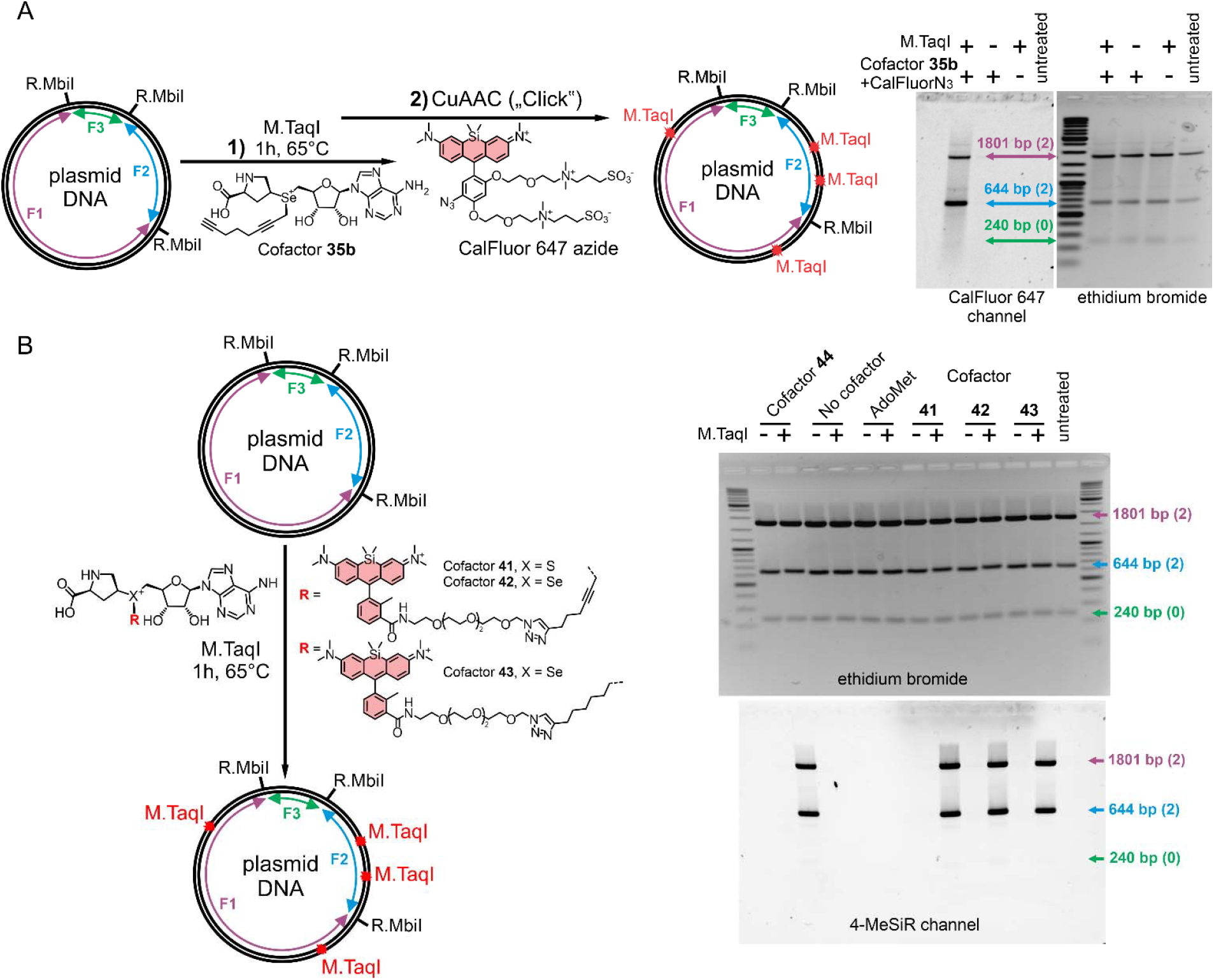
DNA labeling with proline-based AdoMet analogs. Principal scheme of two step (A) and single step (B) labeling of pUC19 plasmid (left) and agarose gel electrophoresis demonstrating site-specific DNA labelling after digestion with R.MbiI.

Upon labeling pUC19 DNA with M.TaqI and fluorescent proline cofactors **41, 42**, and **44** afforded complete protection from R.TaqI digestion, confirming successful fluorophore transfer (Figure S25). The specificity of labeling was validated by digestion with R.MbiI and analyzing fluorescent DNA fragments on agarose gel (Figure 3B). Identity of adenine modifications was confirmed by LC–MS analysis using duplex oligonucleotide under single-turnover reaction conditions (Figure S26).

## CONCLUSIONS

In conclusion, we have introduced a new class of stabilized AdoMet analogs based on a selenonium proline scaffold that effectively overcomes the main degradation pathways. These analogs are compatible with wild-type methyltransferase TaqI, enabling efficient sequencespecific labeling with both small and extended chemical groups. The improved cofactor lifetime significantly enhances the practical utility of methyltransferase-mediated tagging or labeling under physiological conditions or elevated temperatures that is required for thermophilic methyltransferases. This design strategy paves the way for more robust biomolecular labeling tools and we believe it may be extended to other types of AdoMet-dependent enzymatic systems. The modularity of the approach also supports future development of stable and specifically tailored AdoMet analogs for tagging, epigenetic profiling or other applications in diagnostics.

## AUTHOR INFORMATION

### Author Contributions

All authors performed the experiments, analyzed the obtained data and prepared the manuscript.

### Notes

The authors have filed patent application No. PCT/EP2025/056156 covering proline cofactor analogs.

## ACKNOWLEDGMENT

The authors thank the Max Planck Society for supporting this work. The authors are grateful to Dr. Vladimir Belov, Jan Seikowski, Jens Schimpfhauser, and Jürgen Bienert (Facility for Synthetic Chemistry, MPI-NAT) for acquiring NMR spectra. The authors acknowledge Dr. Holm Frauendorf and the central analytics/mass spectrometry team (Institute for Organic and Biomolecular Chemistry, Georg-August University, Göttingen) for recording HRMS spectra. The authors thank Georgij Kostiuk for providing purified M.TaqI.

