## Supplementary information for "Dual Stabilization of S-Adenosylmethionine for Enzymatic DNA Labeling"

### Contents

**Supplementary Figures** ... 3
Figure S1 ... 3
Figure S2 ... 4
Figure S3 ... 5
Figure S4 ... 6
Figure S5 ... 7
Figure S6 ... 8
Figure S7 ... 9
Figure S8 ... 10
Figure S9. ... 11
Figure S10. ... 12
Figure S11 ... 13
Figure S12 ... 14
Figure S13 ... 15
Figure S14 ... 16
Figure S15 ... 17
Figure S16 ... 18
Figure S17 ... 19
Figure S18 ... 20
Figure S19 ... 21
Figure S20 ... 22
Figure S21 ... 23
Figure S22 ... 24
Figure S23 ... 25
Figure S24 ... 26
Figure S25 ... 27
Figure S26 ... 28
**Supplementary Methods** ... 29
HPLC-MS Analysis of Cofactor Decomposition Kinetics and Products: ... 29
DNA protection assay ... 29
Chemo-enzymatic two-step labeling of plasmid DNA: ... 31
Enzymatic single-step labeling of plasmid DNA: ... 32
General chemical experimental information and synthesis methods ... 33
Copies of NMR spectra ... 97
**Supplementary References** ... 137

### Supplementary Figures

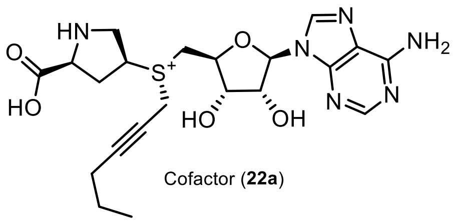

Cofactor (**22a**)

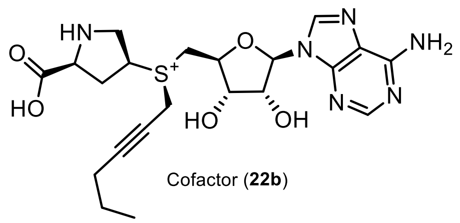

Cofactor (**22b**)

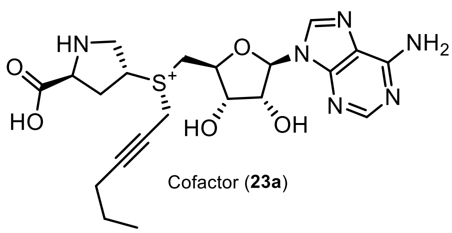

Cofactor (**23a**)

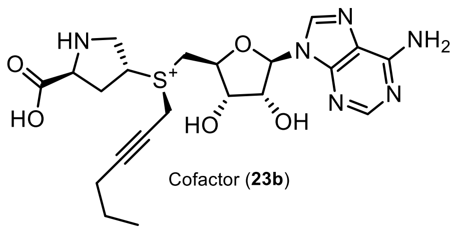

Cofactor (**23b**)

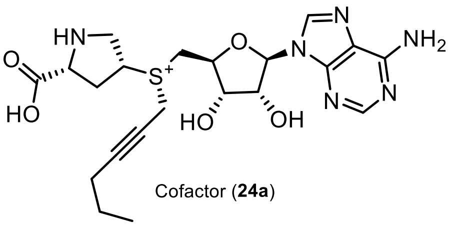

Cofactor (**24a**)

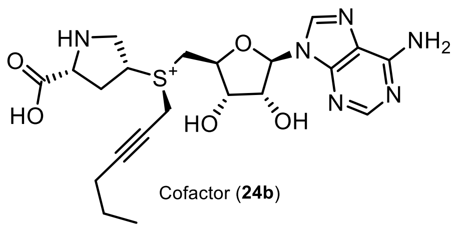

Cofactor (**24b**)

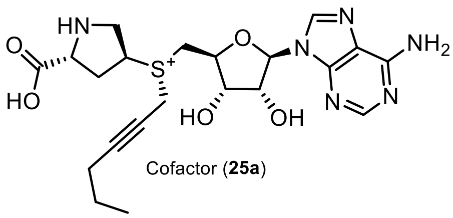

Cofactor (**25a**)

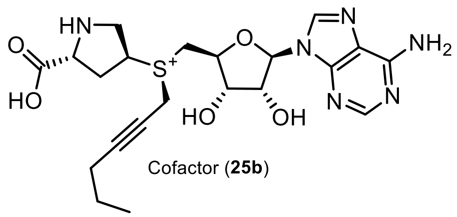

Cofactor (**25b**)

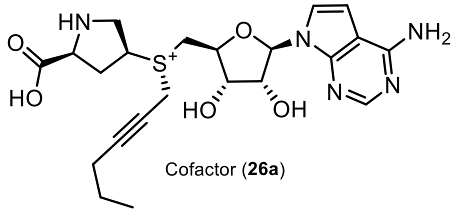

Cofactor (**26a**)

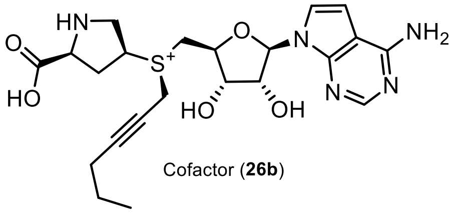

Cofactor (**26b**)

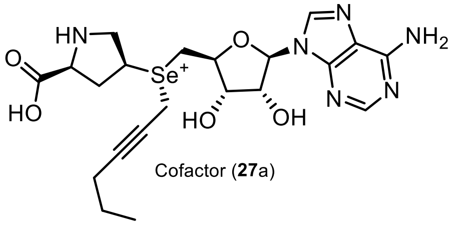

Cofactor (**27**a)

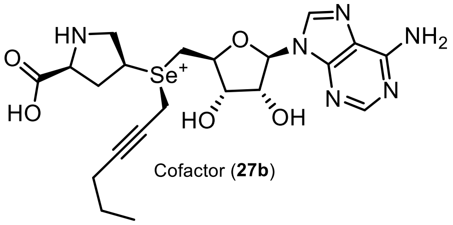

Cofactor (**27b**)

**Figure S1.** Structures of proline cofactors (**22a**-**27a** and **22b**-**27b**) synthesized and used in this study (part 1 of 4).

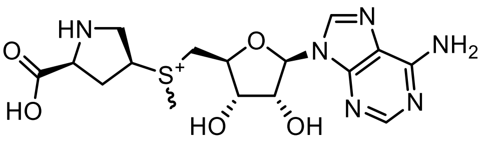

Cofactor (**28ab**)

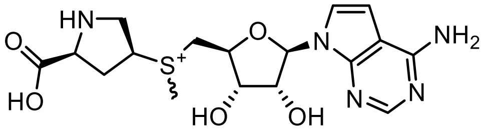

Cofactor(**29ab**)

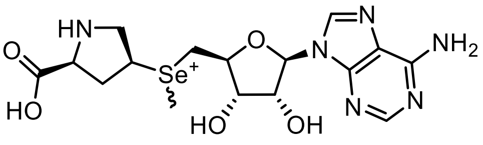

Cofactor (**30ab**)

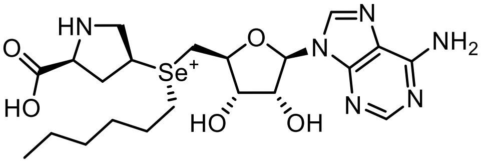

Cofactor (**31a**)

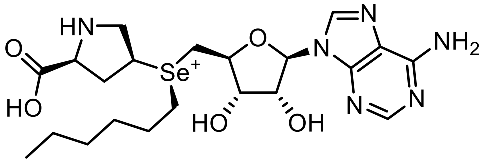

Cofactor (**31b**)

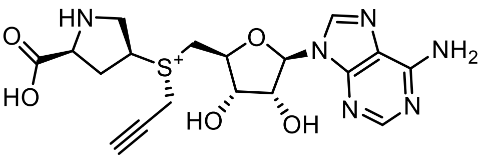

Cofactor **32a**

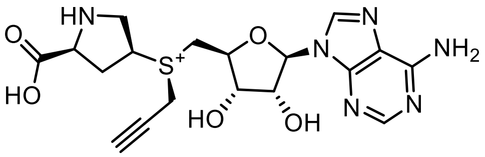

Cofactor (**32b**)

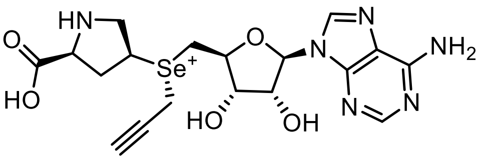

Cofactor (**33a**)

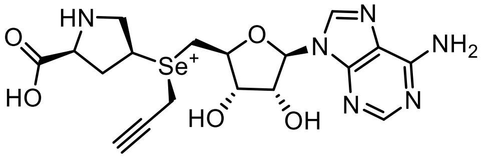

Cofactor (**33b**)

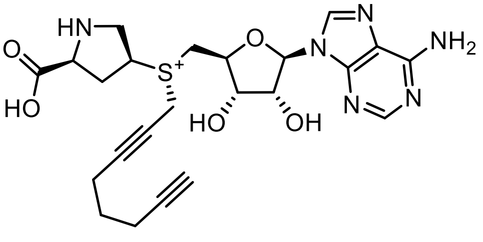

Cofactor (**34a**)

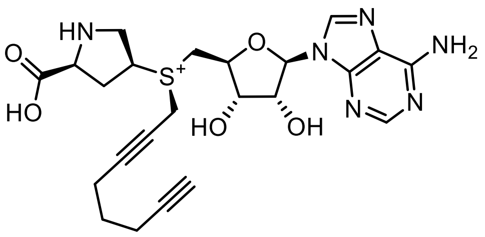

Cofactor (**34b**)

**Figure S2.** Structures of proline cofactors (**28ab**-**30ab**, **31a**-**34a** and **31b**-**34b**) synthesized and used in this study (part 2 of 4).

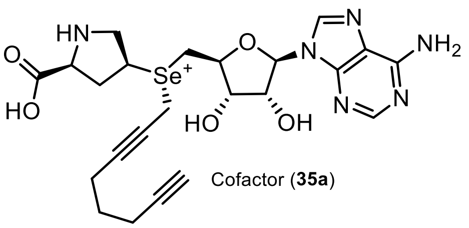

Cofactor (**35a**)

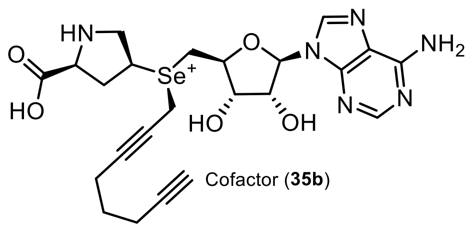

Cofactor (**35b**)

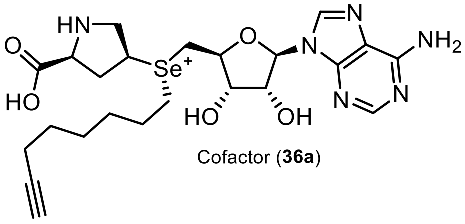

Cofactor (**36a**)

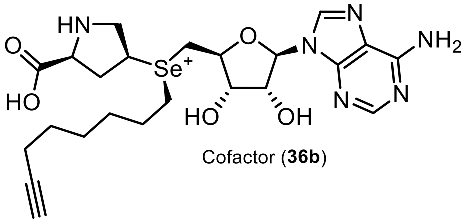

Cofactor (**36b**)

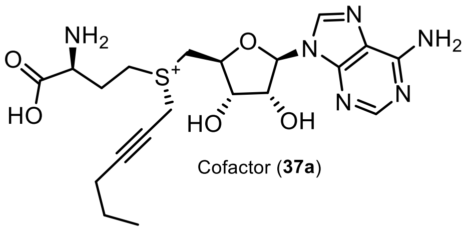

Cofactor (**37a**)

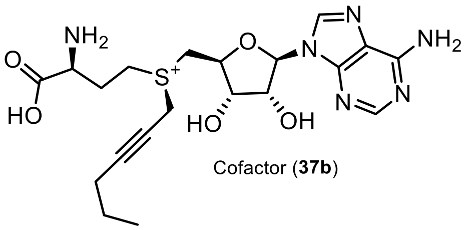

Cofactor (**37b**)

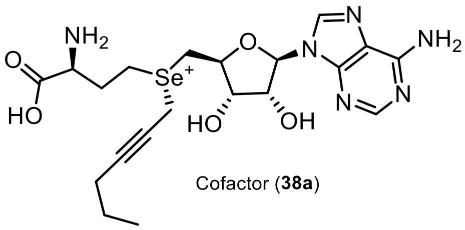

Cofactor (**38a**)

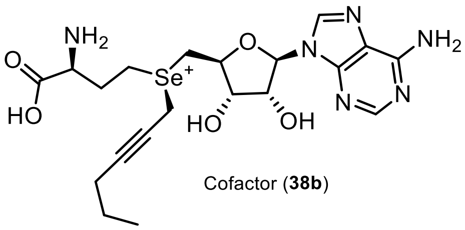

Cofactor (**38b**)

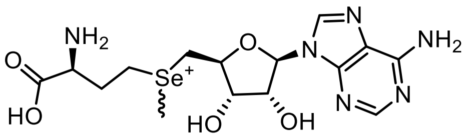

**SeAdoMet**
Cofactor (**39ab**)

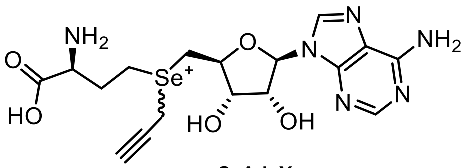

**SeAdoYn**
Cofactor (**40ab**)

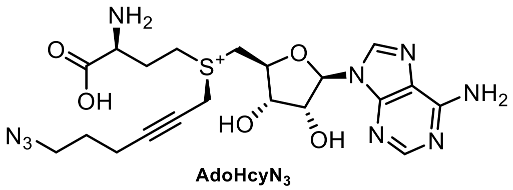

**AdoHcyN$_3$**

**Figure S3.** Structures of proline (**35a**, **35b**, **36a**, **36b**) and homoalanine (**37a**, **37b**, **38a**, **38b**, **39ab**, **40ab** and **AdoHcyN$_3$**) cofactors synthesized and used in this study (part 3 of 4).

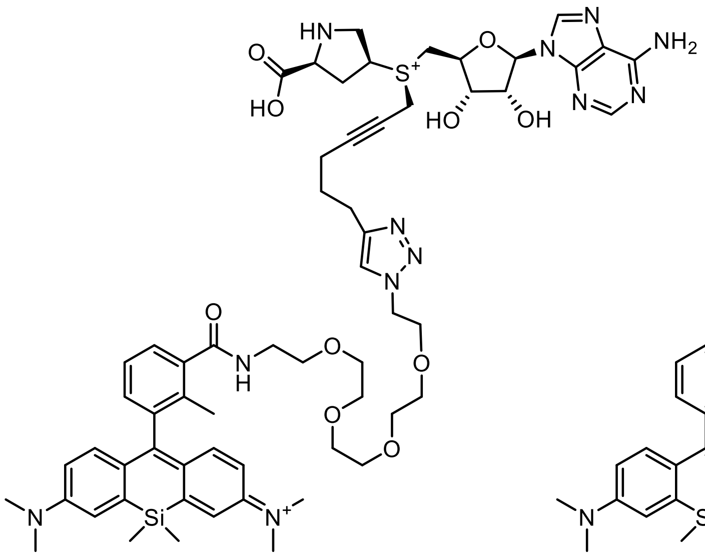

Cofactor (**41**)

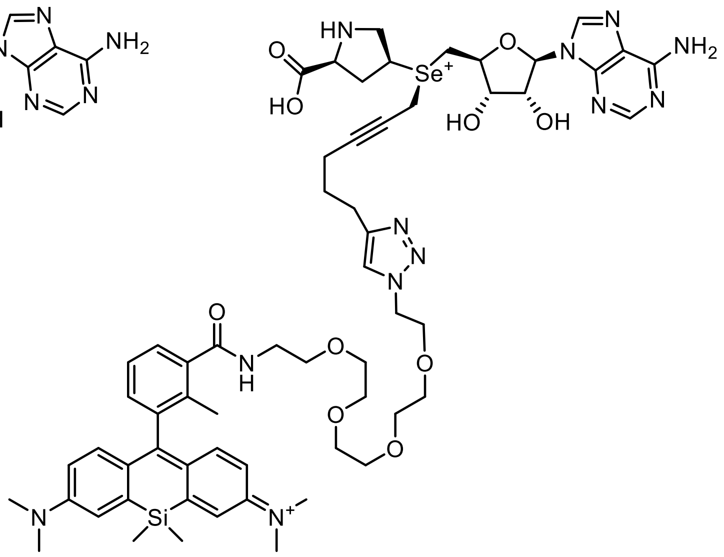

Cofactor (**42**)

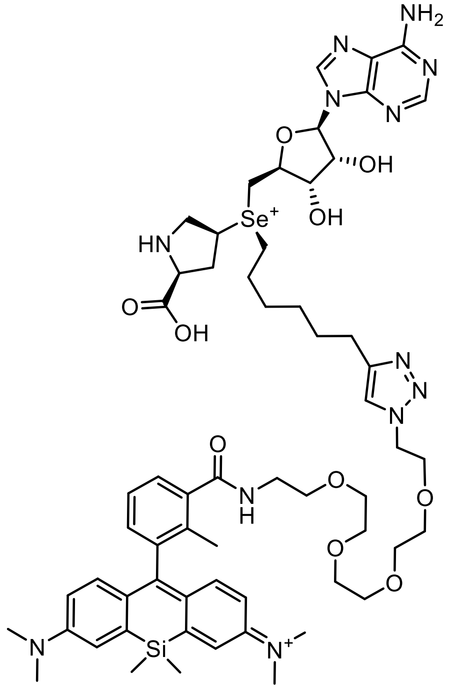

Cofactor (**43**)

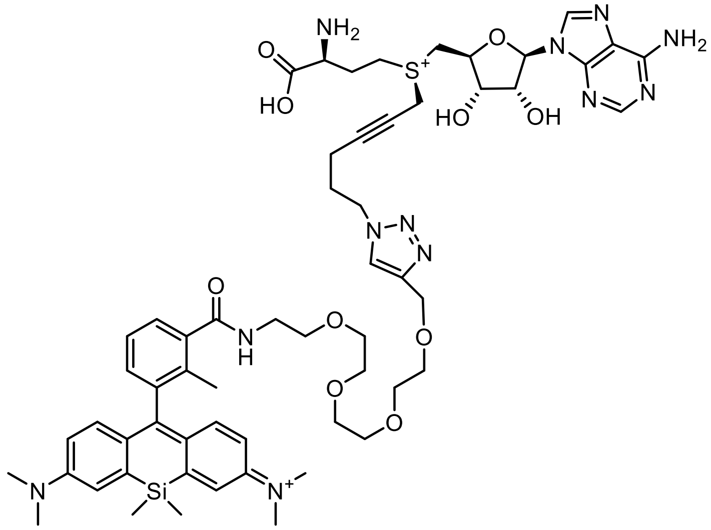

Cofactor (**44**)

**Figure S4.** Structures of fluorescent proline cofactors (**41** – **44**) synthesized and used in this study (part 4 of 4).

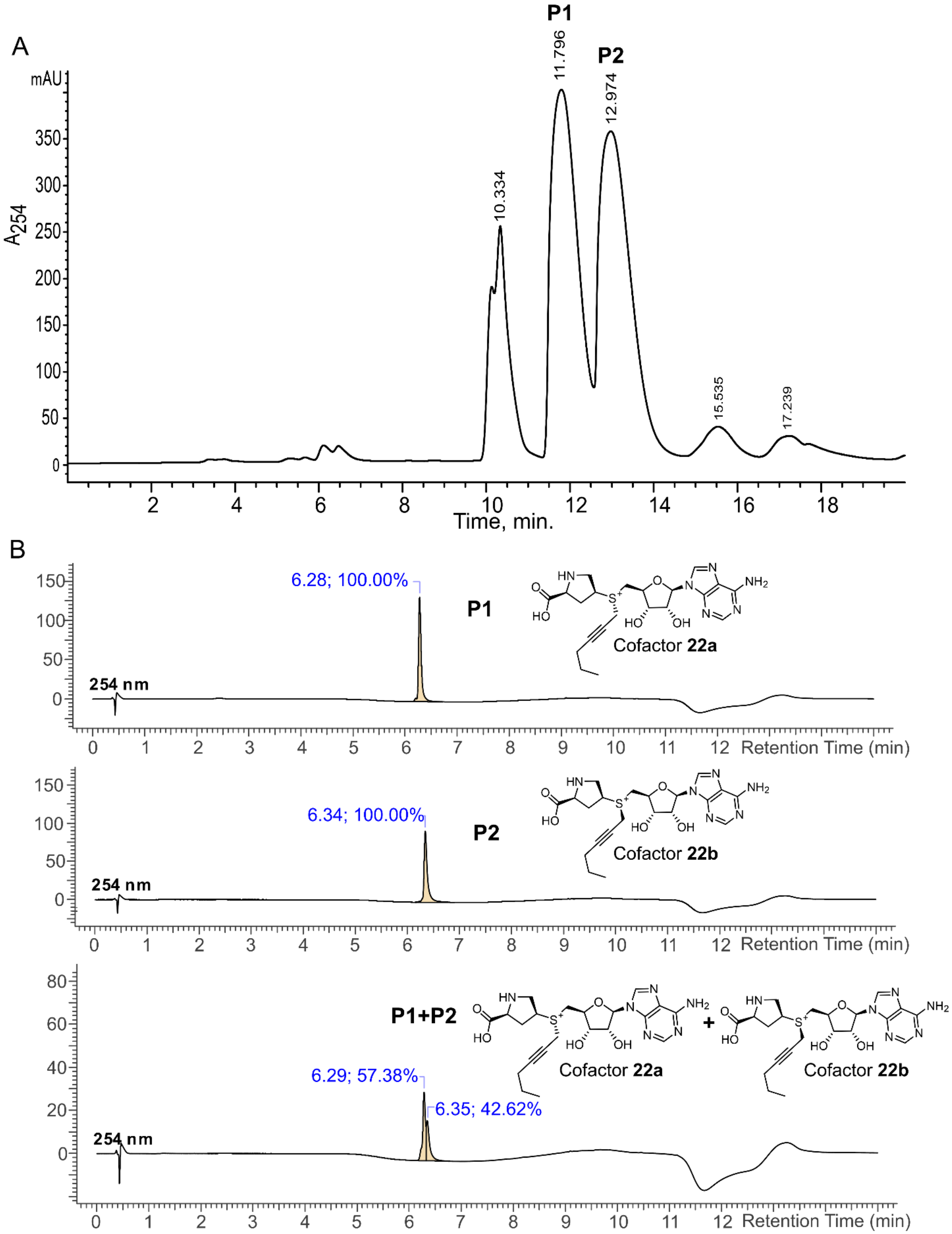


**Figure S5.** Example of separation of diastereomeric mixtures of synthesized cofactors. (A) Preparative HPLC chromatogram of cofactors **22a** and **22b** separation. (B) Analytical HPLC chromatograms of purified diastereomers **22a** and **22b** and a chromatogram of mixture **22a**+**22b**.

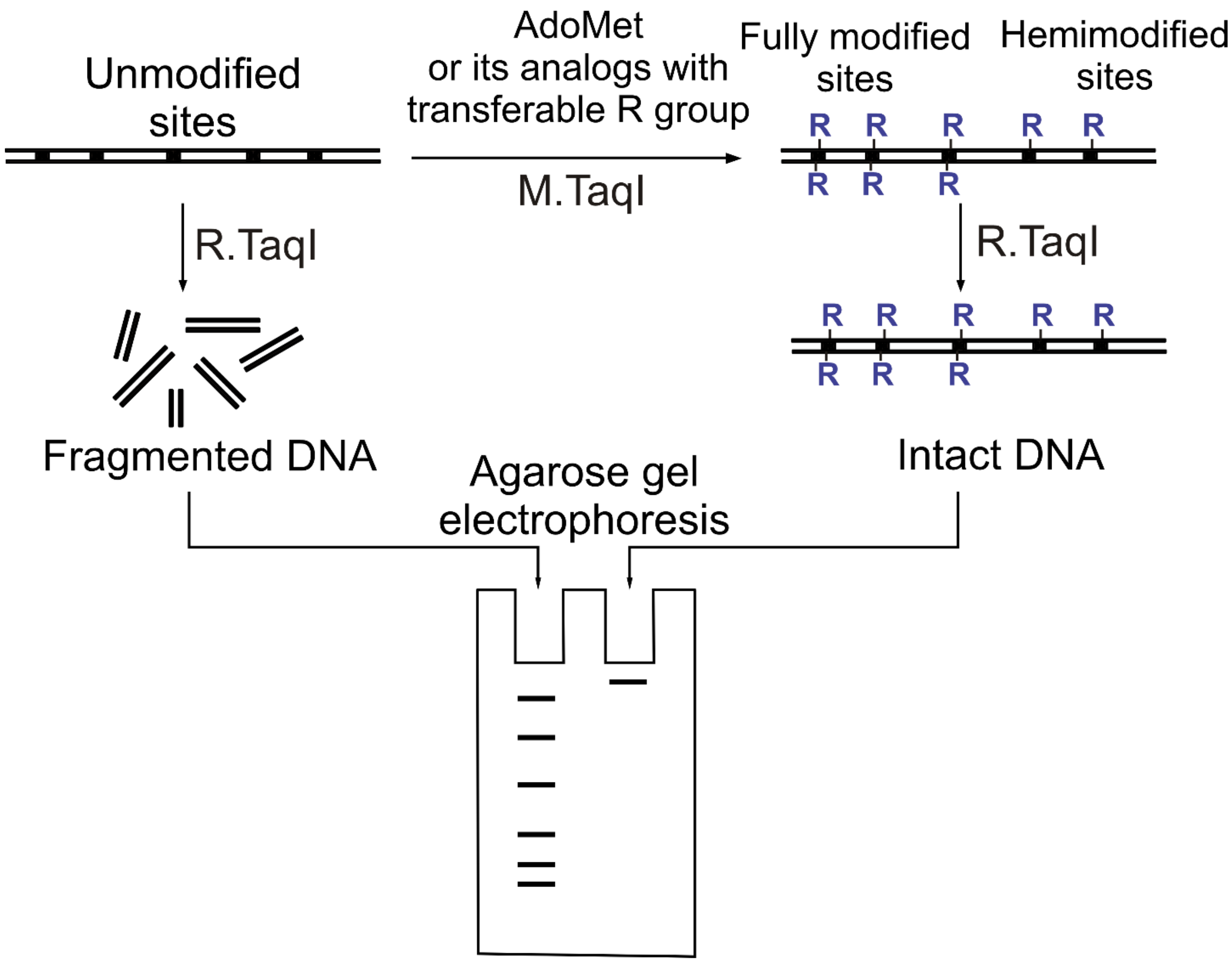


**Figure S6**. Principle of a DNA protection assay. It based on DNA alkylation using M.TaqI and modified DNA hydrolysis with R.TaqI.

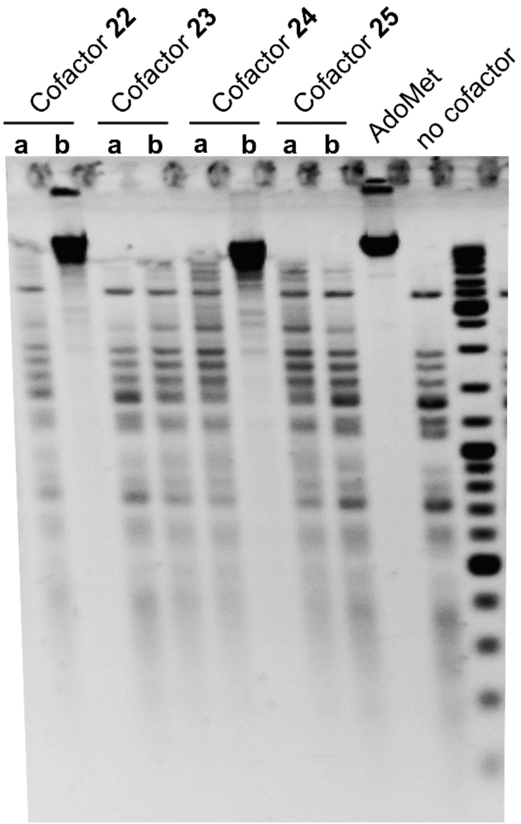


| | Amino acid side chain | Z = | X = | R = |
|---|---|---|---|---|
| AdoMet | L-Abu | N | S | Methyl |
| Cofactor **22** | *cis*-L-Pro | N | S | Hex-2-ynyl |
| Cofactor **23** | *trans*-L-Pro | N | S | Hex-2-ynyl |
| Cofactor **24** | *cis*-D-Pro | N | S | Hex-2-ynyl |
| Cofactor **25** | *trans*-D-Pro | N | S | Hex-2-ynyl |

**Figure S7**. Activity of wild-type M.TaqI with proline-based cofactors **22**–**25** bearing a transferable hex-2-ynyl group. Agarose gel analysis of λ-DNA following treatment with M.TaqI and diastereomeric cofactors demonstrates sequence-specific protection from R.TaqI digestion by the cis-proline stereoisomers 22b and 24b. The modification reaction contained 60 ng/µL λ-DNA, 0.5 U/µL M.TaqI (New England Biolabs), and 100 µM cofactor, and was incubated for 1 h at 65 °C.

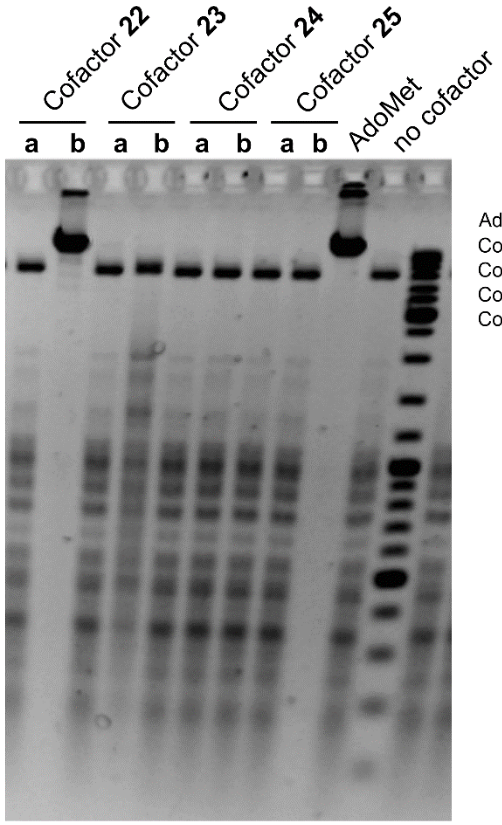


| | Amino acid side chain | Z = | X = | R = |
|---|---|---|---|---|
| AdoMet | L-Abu | N | S | Methyl |
| Cofactor **22** | *cis*-L-Pro | N | S | Hex-2-ynyl |
| Cofactor **23** | *trans*-L-Pro | N | S | Hex-2-ynyl |
| Cofactor **24** | *cis*-D-Pro | N | S | Hex-2-ynyl |
| Cofactor **25** | *trans*-D-Pro | N | S | Hex-2-ynyl |

**Figure S8**. Activity of M.HhaI Q82A/Y254S/N304A mutant with proline-based cofactors **22**–**25** bearing a transferable hex-2-ynyl group. Agarose gel of λ-DNA after treatment with M.HhaI Q82A/Y254S/N304A mutant and diastereomeric cofactors showing sequence-specific protection of λ-DNA from R.Hin6I digestion by single active **22b** cofactor. The modification reaction was incubated for 1 h at 37°C with M.HhaI Q82A/Y254S/N304A:targets ratio of 1:1 in the presence of 100 µM cofactor.

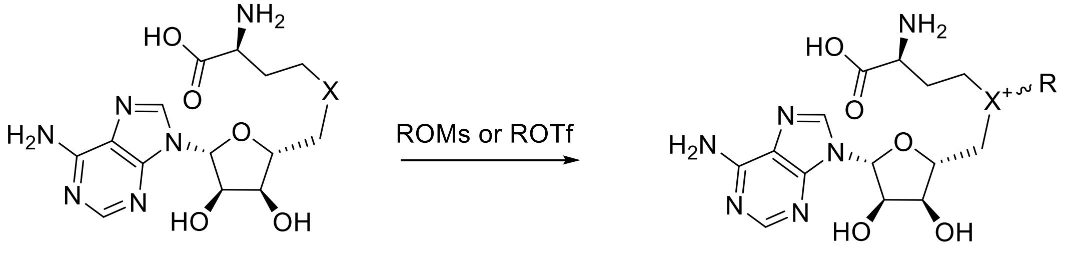


**37ab** (X = Se, R = -$CH_3$)

**38a** and **38b** (X = S, R = $-CH_2C{\equiv}C(CH_2)_2CH_3$)

**39a** and **39b** (X = Se, R = $-CH_2C{\equiv}C(CH_2)_2CH_3$)

**40ab** (X = Se, R = -$CH_2C{\equiv}CH$)

**Figure S9.** Synthesis scheme of homoalanine cofactors **37ab**-**40ab**.

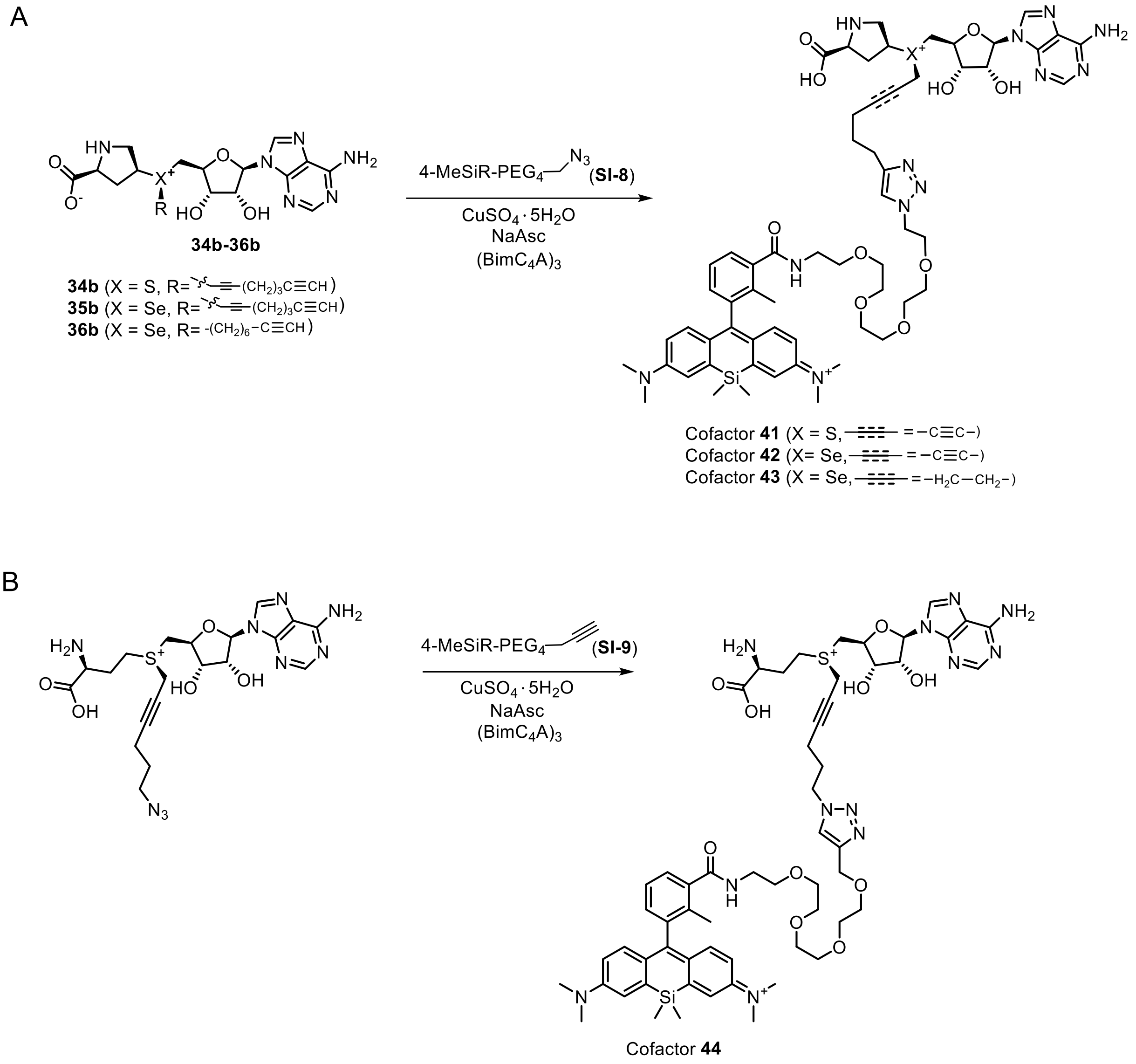


**Figure S10.** Synthesis scheme of fluorescent proline cofactors **41**-**43** (**A**) and fluorescent homoalanine cofactor **44** (**B**).

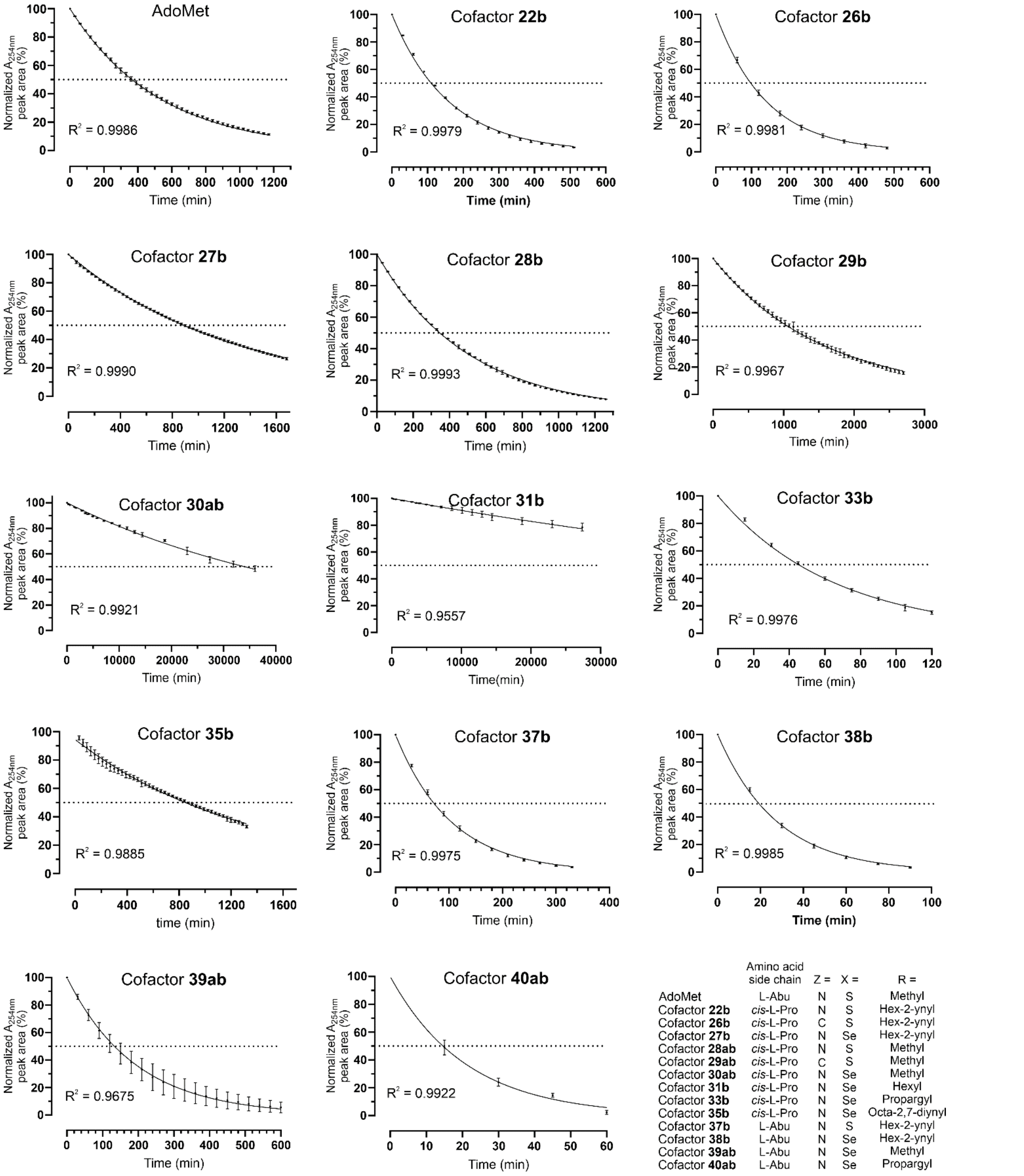


| | Amino acid side chain | Z = | X = | R = |
|---|---|---|---|---|
| AdoMet | L-Abu | N | S | Methyl |
| Cofactor **22b** | *cis*-L-Pro | N | S | Hex-2-ynyl |
| Cofactor **26b** | *cis*-L-Pro | C | S | Hex-2-ynyl |
| Cofactor **27b** | *cis*-L-Pro | N | Se | Hex-2-ynyl |
| Cofactor **28ab** | *cis*-L-Pro | N | S | Methyl |
| Cofactor **29ab** | *cis*-L-Pro | C | S | Methyl |
| Cofactor **30ab** | *cis*-L-Pro | N | Se | Methyl |
| Cofactor **31b** | *cis*-L-Pro | N | Se | Hexyl |
| Cofactor **33b** | *cis*-L-Pro | N | Se | Propargyl |
| Cofactor **35b** | *cis*-L-Pro | N | Se | Octa-2,7-diynyl |
| Cofactor **37b** | L-Abu | N | S | Hex-2-ynyl |
| Cofactor **38b** | L-Abu | N | Se | Hex-2-ynyl |
| Cofactor **39ab** | L-Abu | N | Se | Methyl |
| Cofactor **40ab** | L-Abu | N | Se | Propargyl |

**Figure S11.** Stability profiles of AdoMet or its analogs. Degradation kinetics of AdoMet or its analogs over time in 50 mM Tris-HCl, pH 8.0, 37 °C, monitored by HPLC, N = 3. Fitted to exponential one phase decay as implemented in GraphPad Prism 10.

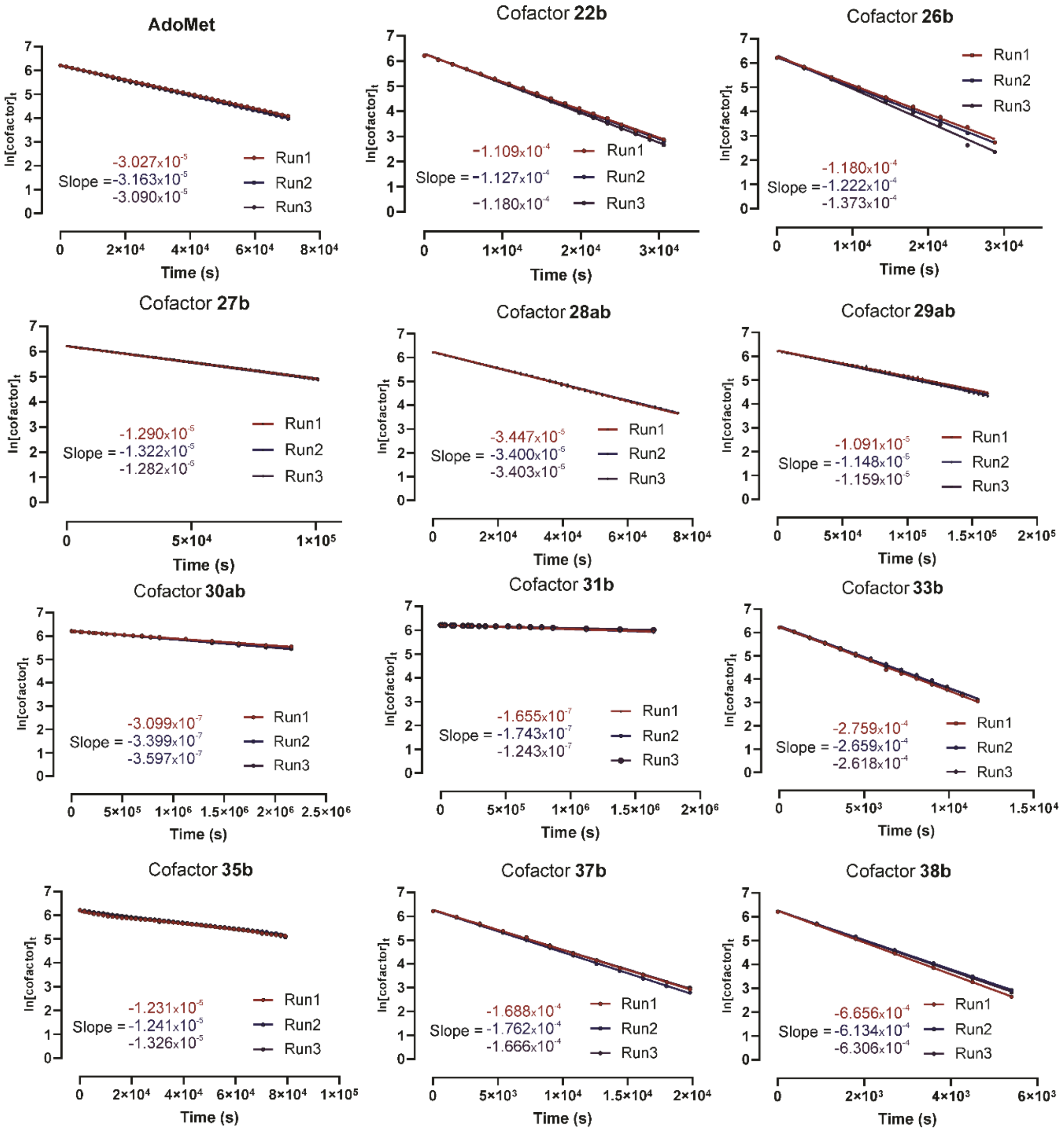


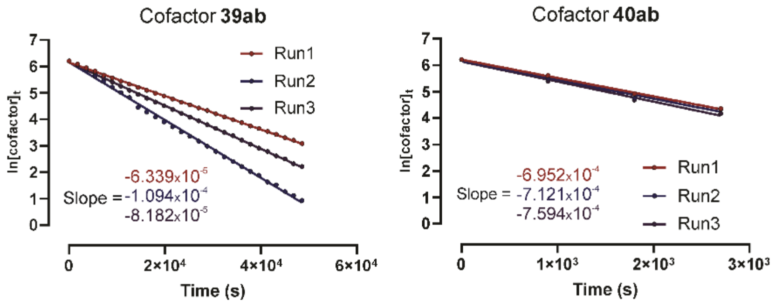


| | Amino acid side chain | Z = | X = | R = |
|---|---|---|---|---|
| AdoMet | L-Abu | N | S | Methyl |
| Cofactor **22b** | *cis*-L-Pro | N | S | Hex-2-ynyl |
| Cofactor **26b** | *cis*-L-Pro | C | S | Hex-2-ynyl |
| Cofactor **27b** | *cis*-L-Pro | N | Se | Hex-2-ynyl |
| Cofactor **28ab** | *cis*-L-Pro | N | S | Methyl |
| Cofactor **29ab** | *cis*-L-Pro | C | S | Methyl |
| Cofactor **30ab** | *cis*-L-Pro | N | Se | Methyl |
| Cofactor **31b** | *cis*-L-Pro | N | Se | Hexyl |
| Cofactor **33b** | *cis*-L-Pro | N | Se | Propargyl |
| Cofactor **35b** | *cis*-L-Pro | N | Se | Octa-2,7-diynyl |
| Cofactor **37b** | L-Abu | N | S | Hex-2-ynyl |
| Cofactor **38b** | L-Abu | N | Se | Hex-2-ynyl |
| Cofactor **39ab** | L-Abu | N | Se | Methyl |
| Cofactor **40ab** | L-Abu | N | Se | Propargyl |

**Figure S12**. Log-linear analysis of cofactor degradation kinetics. Decay curves were natural-log transformed to linearize the exponential time course. The transformed data were fit by linear regression assuming pseudo-first-order kinetics. Pseudo-first-order rate constants ($k_{total}$) were determined from the negative slope of the linear region of the fit.

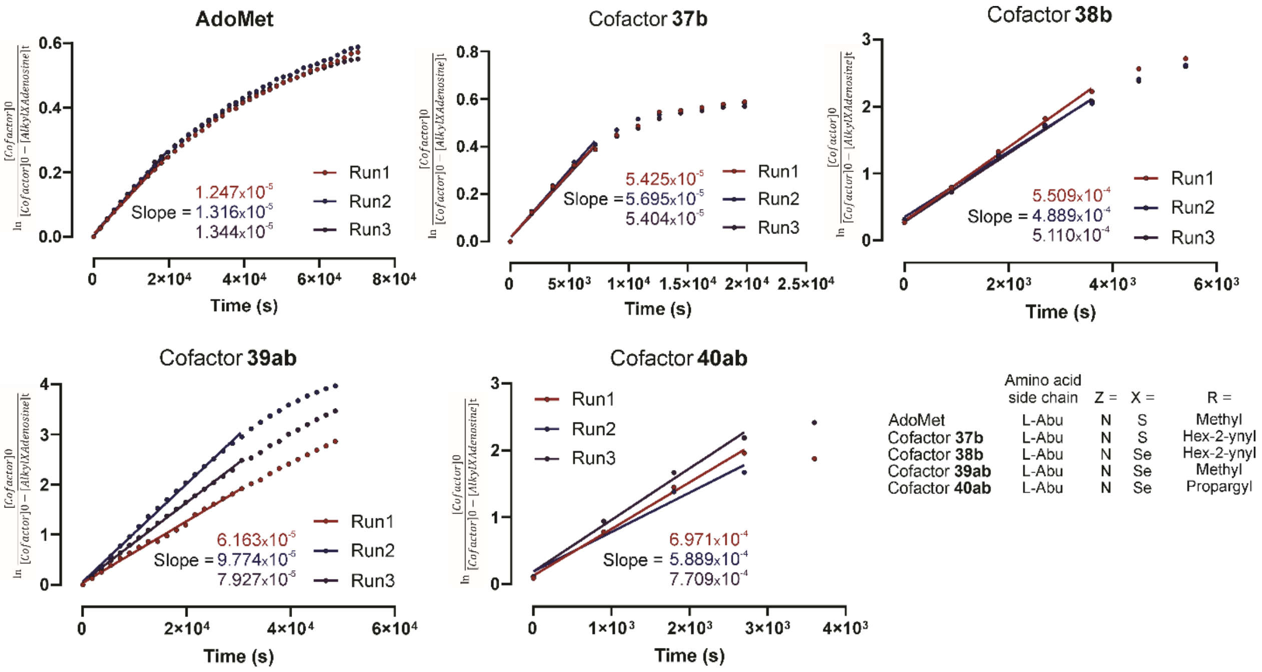


**Figure S13**. Log-linear analysis of intramolecular cyclization product formation kinetics. Product accumulation curves were transformed to linearize the time course and fit by linear regression assuming pseudo first-order kinetics. Pseudo first-order rate constants ($k_{Nuc}$) were determined from the slope of the linear region of the fit.

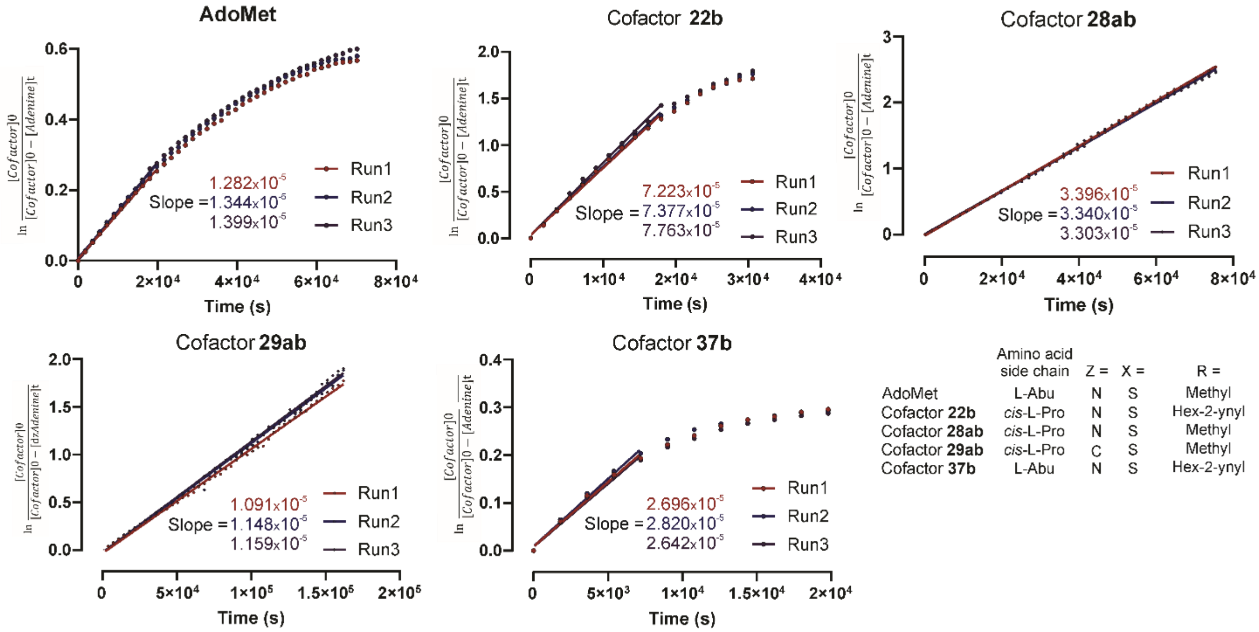


**Figure S14**. Log-linear analysis of depurination product formation kinetics. Adenine or deazaadenine accumulation curves were transformed to linearize the time course and fit by simple linear regression assuming pseudo first-order kinetics. Pseudo first-order rate constants ($k_{Dep}$) were determined from the slope of the linear region of the fit.

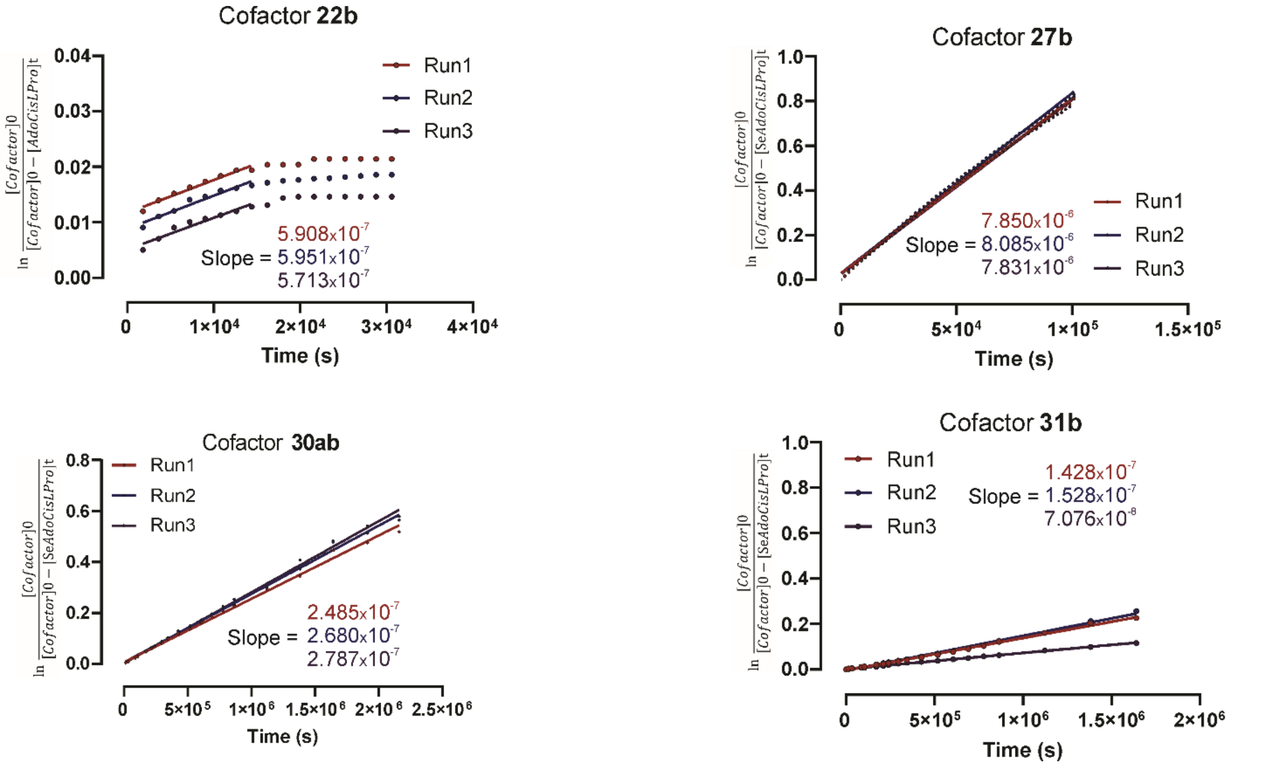


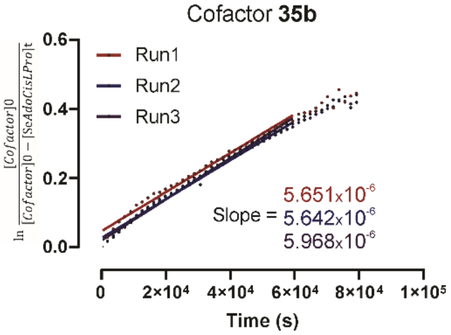


| | Amino acid side chain | Z = | X = | R = |
|---|---|---|---|---|
| Cofactor **22b** | *cis*-L-Pro | N | S | Hex-2-ynyl |
| Cofactor **27b** | *cis*-L-Pro | N | Se | Hex-2-ynyl |
| Cofactor **30ab** | *cis*-L-Pro | N | Se | Methyl |
| Cofactor **31b** | *cis*-L-Pro | N | Se | Hexyl |
| Cofactor **35b** | *cis*-L-Pro | N | Se | Octa-2,7-diynyl |

**Figure S15**. Log-linear analysis of hydrolysis product formation kinetics. Accumulation curves were transformed to linearize the time course and fit by linear regression assuming pseudo first-order kinetics. Pseudo first-order rate constants ($k_{Hyd}$) were determined from the slope of the linear region of the fit.

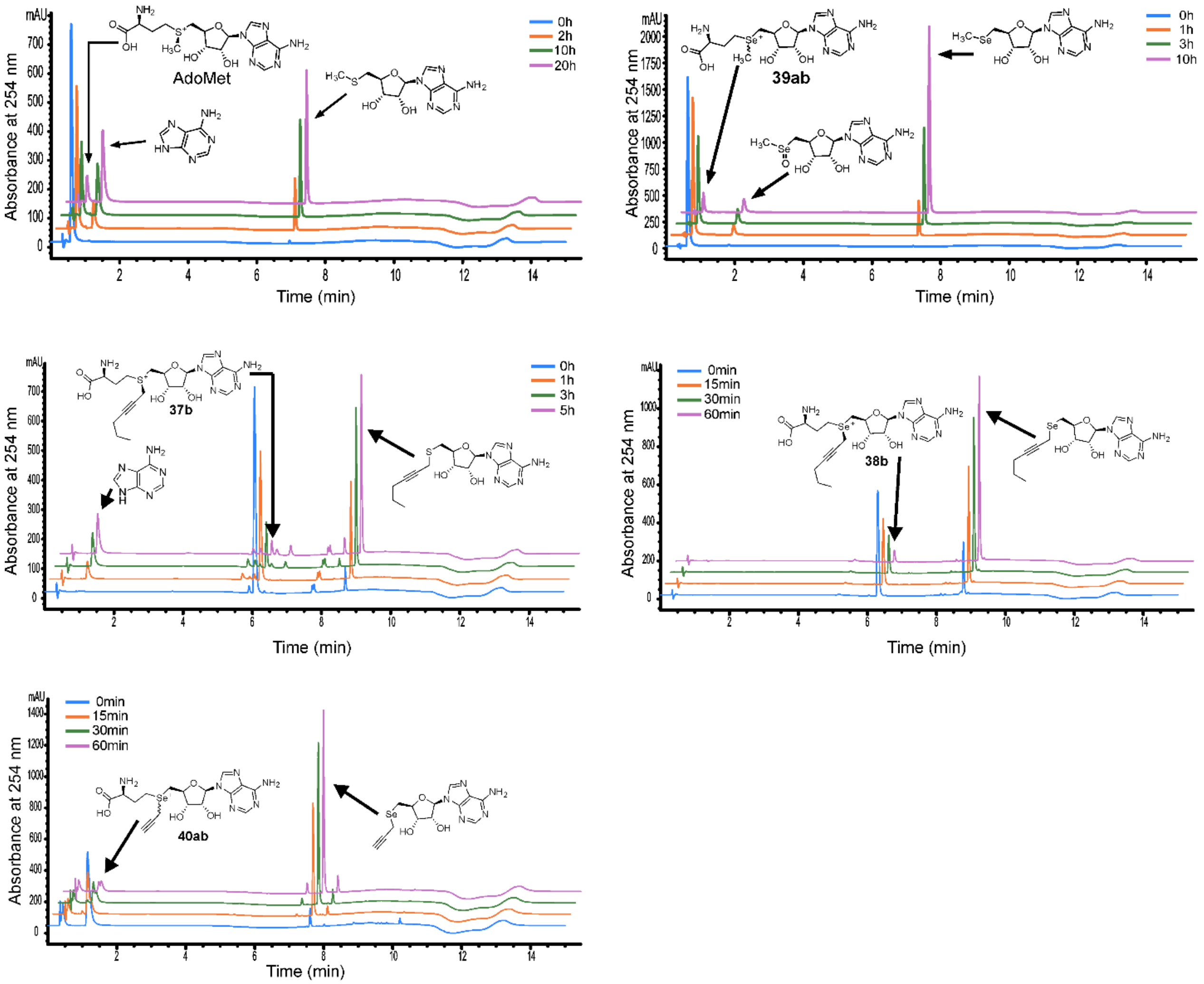


**Figure S16.** Analytical HPLC chromatograms of AdoMet and its analogs **37b**, **38b**, **39ab and 40ab**. Decay reaction products were separated on an Ascentis® Express AQ-C18 UHPLC column (2 µm, 5 cm × 2.1 mm) using a methanol gradient in 25 mM ammonium formate (pH 3.6) aqueous buffer. Peaks detected by absorbance at 254 nm corresponding to the major identified decay products are indicated.

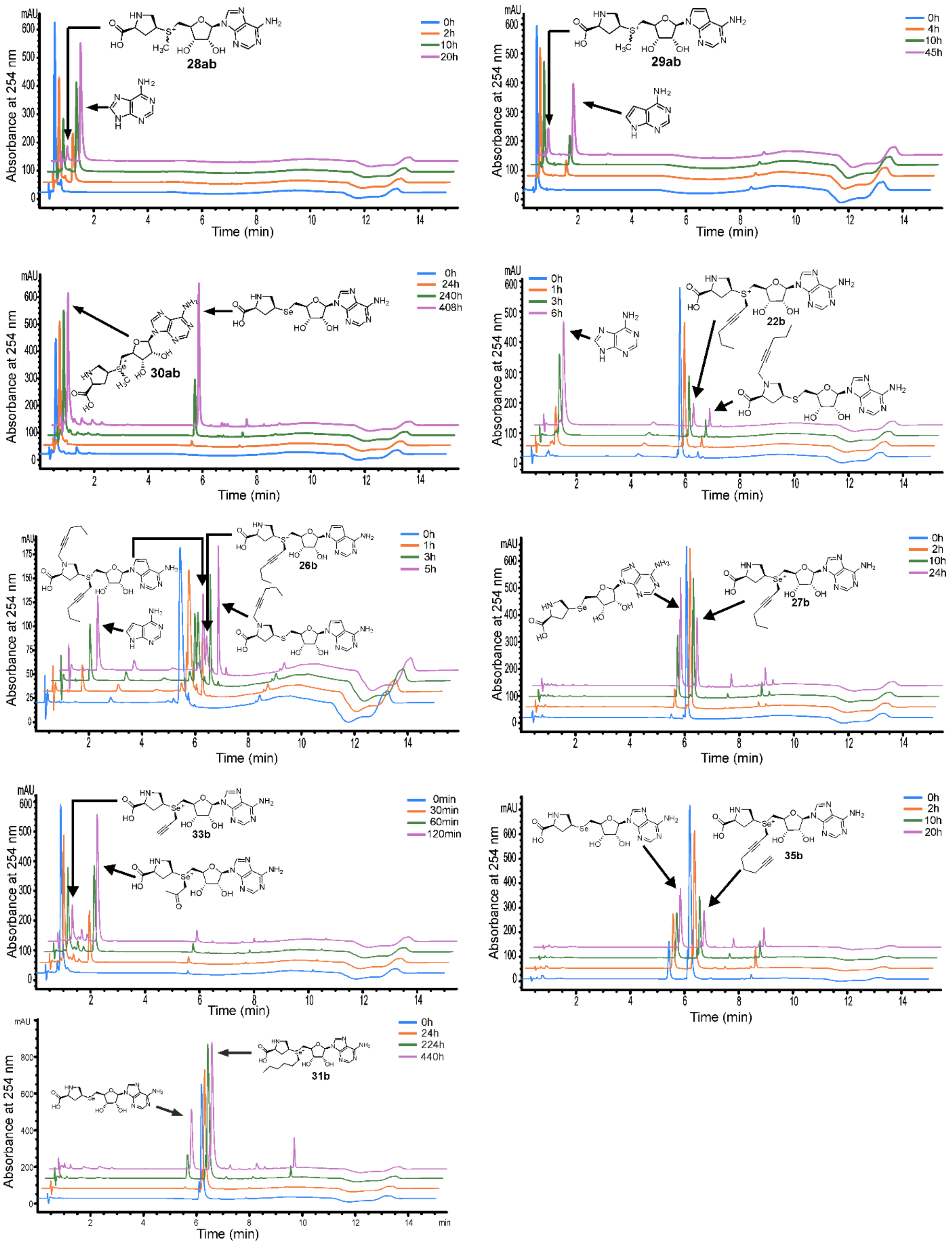


**Figure S17.** Analytical HPLC chromatograms of proline cofactors **22b**, **26b**, **27b**, **28ab**, **29ab**, **30ab**, **31b**, **33b** and **35b**. Decay reaction products were separated on an Ascentis® Express AQ-C18 UHPLC column (2 µm, 5 cm × 2.1 mm) using a methanol gradient in 25 mM ammonium formate (pH 3.6) aqueous buffer. Peaks detected by absorbance at 254 nm corresponding to the major identified decay products are indicated.

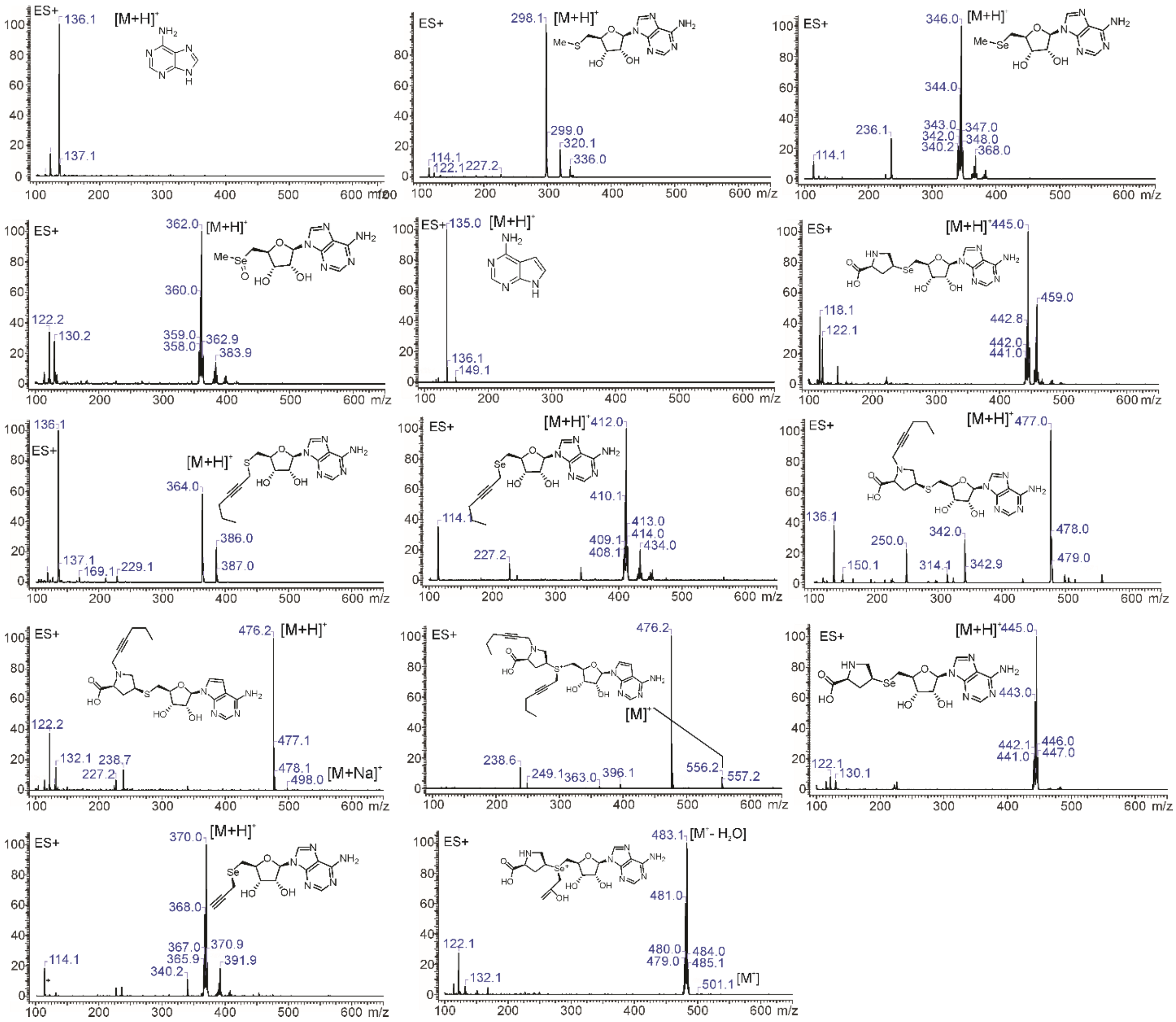

**Figure S18.** MS spectra of identified cofactors' degradation products.

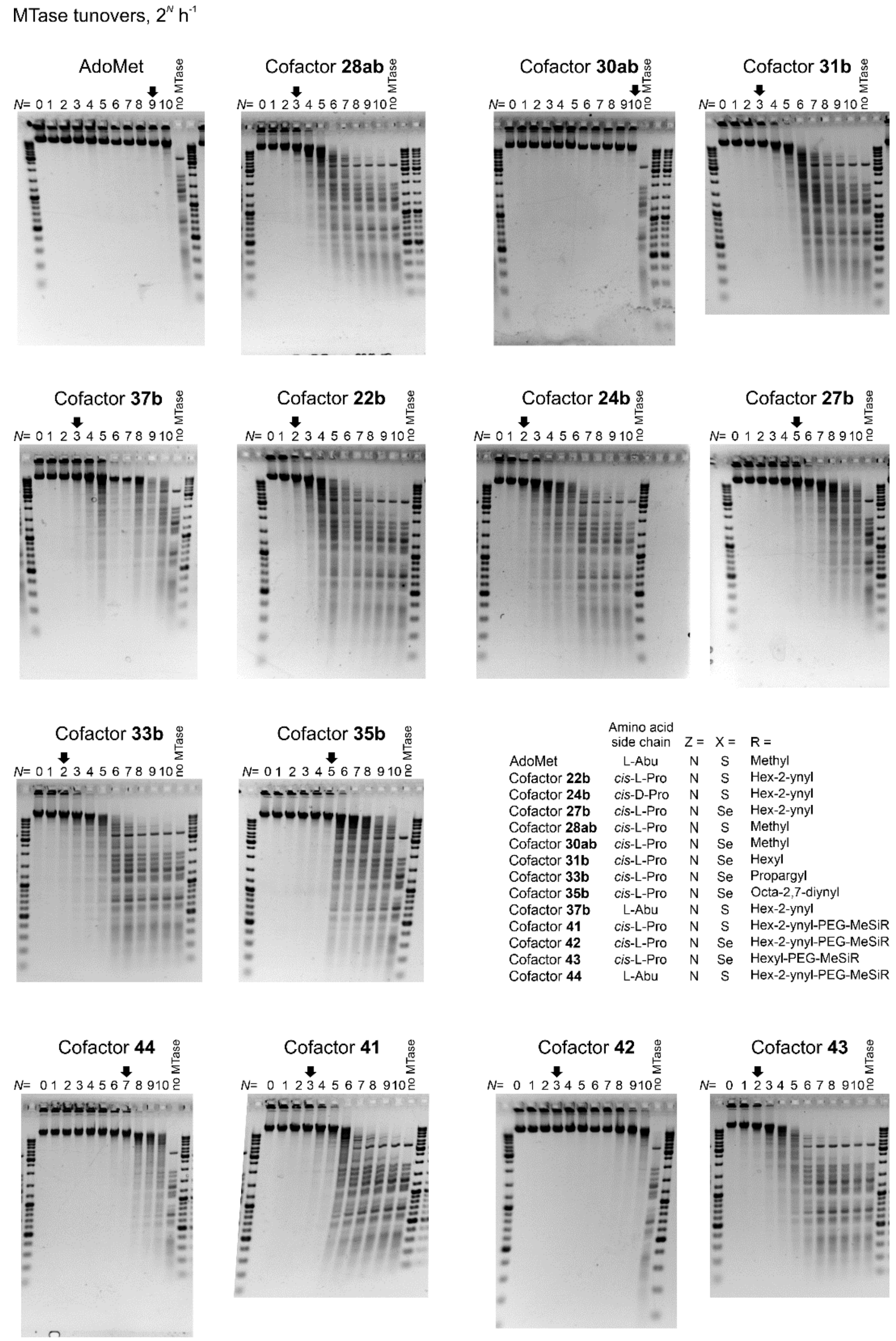


| | Amino acid side chain | Z = | X = | R = |
|---|---|---|---|---|
| AdoMet | L-Abu | N | S | Methyl |
| Cofactor **22b** | *cis*-L-Pro | N | S | Hex-2-ynyl |
| Cofactor **24b** | *cis*-D-Pro | N | S | Hex-2-ynyl |
| Cofactor **27b** | *cis*-L-Pro | N | Se | Hex-2-ynyl |
| Cofactor **28ab** | *cis*-L-Pro | N | S | Methyl |
| Cofactor **30ab** | *cis*-L-Pro | N | Se | Methyl |
| Cofactor **31b** | *cis*-L-Pro | N | Se | Hexyl |
| Cofactor **33b** | *cis*-L-Pro | N | Se | Propargyl |
| Cofactor **35b** | *cis*-L-Pro | N | Se | Octa-2,7-diynyl |
| Cofactor **37b** | L-Abu | N | S | Hex-2-ynyl |
| Cofactor **41** | *cis*-L-Pro | N | S | Hex-2-ynyl-PEG-MeSiR |
| Cofactor **42** | *cis*-L-Pro | N | Se | Hex-2-ynyl-PEG-MeSiR |
| Cofactor **43** | *cis*-L-Pro | N | Se | Hexyl-PEG-MeSiR |
| Cofactor **44** | L-Abu | N | S | Hex-2-ynyl-PEG-MeSiR |

**Figure S19.** Activity of M.TaqI with AdoMet and its analogs. λ DNA was incubated for 1 h at 65°C with 100 µM cofactor and 2-fold serial dilutions of M.TaqI starting from equimolar TCGA sites : M.TaqI ratio and then challenged with R.TaqI. In lane *N*, the ratio of target sites to M.TaqI is $2^N$. Arrows mark the highest *N* at which complete protection from R.TaqI is achieved, indicating that MTase has completed $2^N$ catalytic turnovers. Representative gels from single replicate, N ≥ 3.

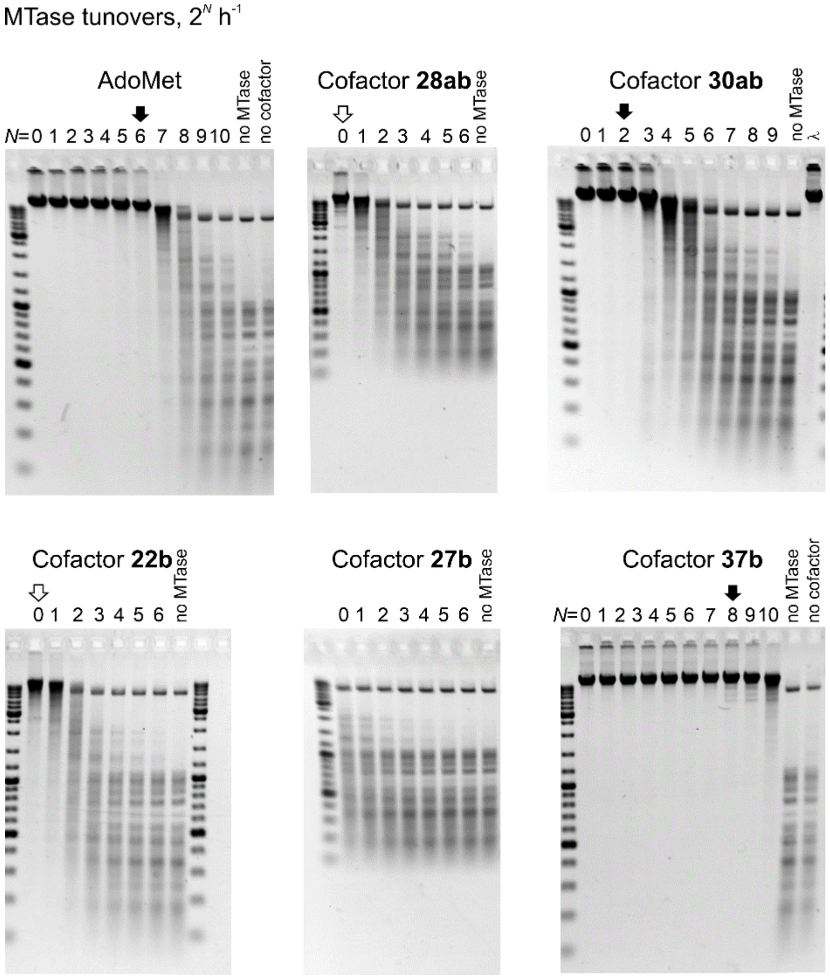


| | Amino acid side chain | Z = | X = | R = |
|---|---|---|---|---|
| AdoMet | L-Abu | N | S | Methyl |
| Cofactor **22b** | *cis*-L-Pro | N | S | Hex-2-ynyl |
| Cofactor **27b** | *cis*-L-Pro | N | Se | Hex-2-ynyl |
| Cofactor **28ab** | *cis*-L-Pro | N | S | Methyl |
| Cofactor **30ab** | *cis*-L-Pro | N | Se | Methyl |
| Cofactor **37b** | L-Abu | N | S | Hex-2-ynyl |

**Figure S20.** Activity of M.HhaI Q82A/Y254A/N304A with AdoMet and its analogs. λ DNA was incubated with 100 µM cofactor and 2-fold serial dilutions of M.HhaI Q82A/Y254A/N304A, starting from equimolar GCGC target sites : MTase ratio. After 1 h at 37°C, MTase was inactivated by heating samples for 10 min. at 80°C and then challenged with R.Hin6I. In lane *N*, the ratio of target sites to M.HhaI Q82A Y254A N304A is $2^N$. Black arrows mark the highest *N* at which complete protection from R.Hin6I is achieved, indicating that MTase has completed $2^N$ catalytic turnovers. Open arrow indicates substantial but incomplete protection even at the highest MTase concentration. Representative gels of a single replicate, N ≥ 3.

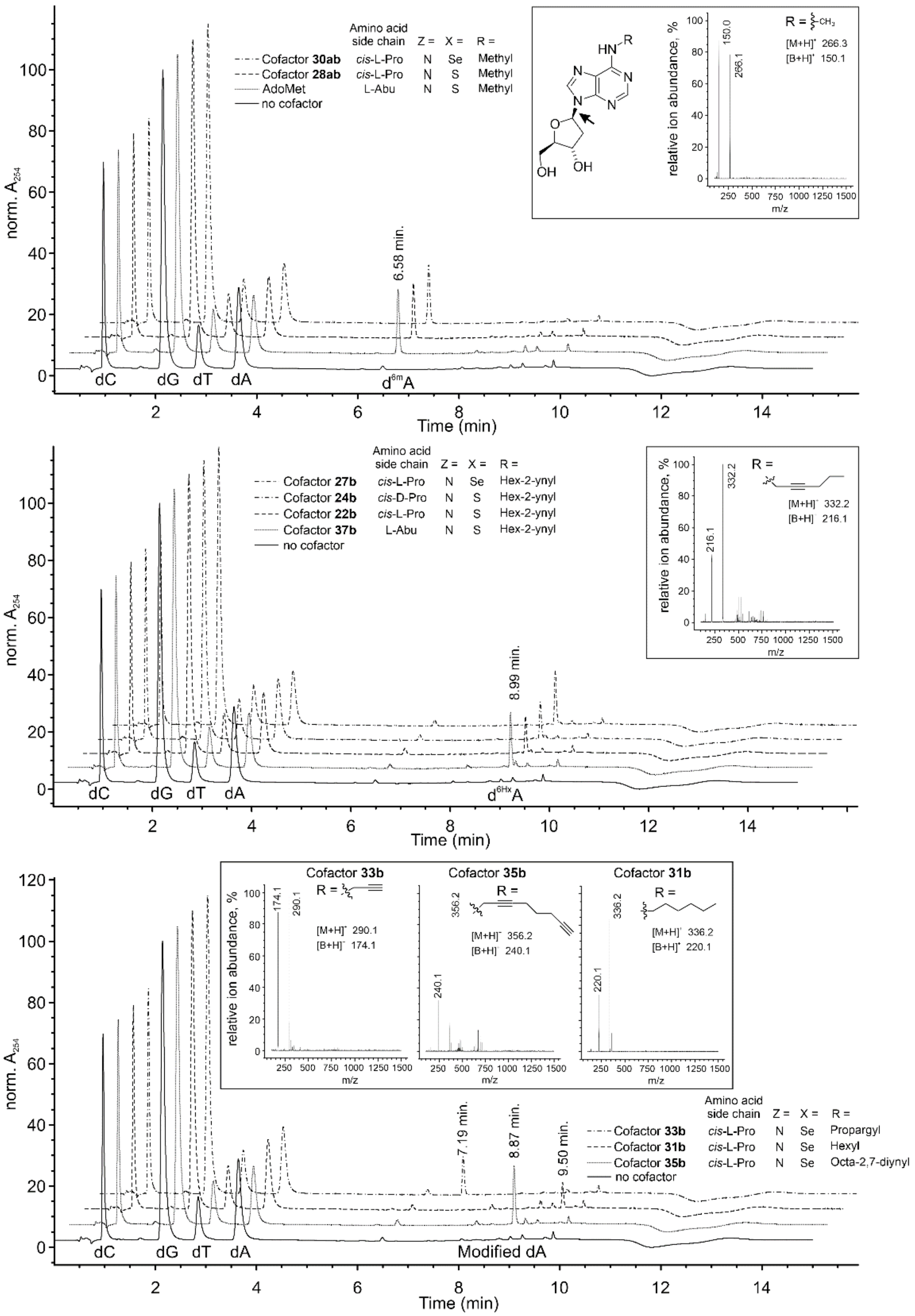


**Figure S21.** LC/MS analysis and identification of products formed in the reaction of M.TaqI with new cofactor analogs. Excess of M.TaqI was incubated with 23-mer duplex DNA substrate and 50 µM AdoMet or its analog for 1 h at 65°C, then DNA was hydrolyzed to deoxynucleosides and the products were analyzed by LC/MS. Retention times of $d^{6X}A$ products are indicated above the peaks. Insets: general structure of modified deoxyadenosine ($d^{6X}A$), where -R – group transferred from the corresponding cofactor analog; black arrow denotes a major site of fragmentation that occurs in mass spectrometer. Experimental mass spectra at the corresponding product peak and theoretical m/z values are shown. $[M+H]^+$ – protonated molecular ion, $[B+H]^+$ – protonated nucleobase.

A 5'-CCCGCCGAAG**TCGA**GCGCCGAAC
GGGCGGCTTC**AGCT**CGCGGCTTG-5'

B

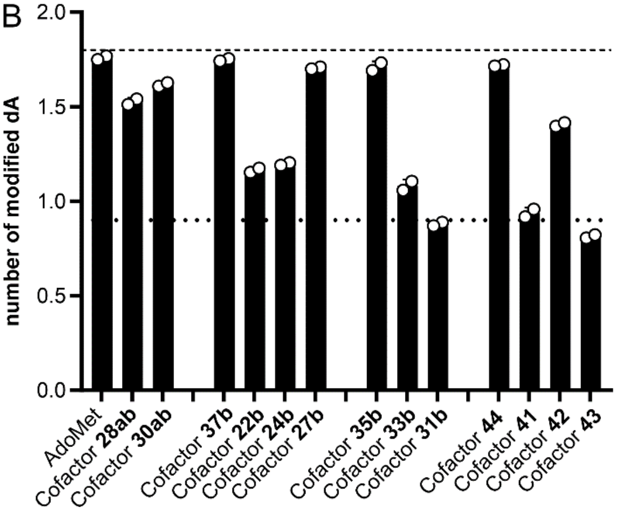


| | Amino acid side chain | Z = | X = | R = |
|---|---|---|---|---|
| AdoMet | L-Abu | N | S | Methyl |
| Cofactor **22b** | *cis*-L-Pro | N | S | Hex-2-ynyl |
| Cofactor **24b** | *cis*-D-Pro | N | S | Hex-2-ynyl |
| Cofactor **27b** | *cis*-L-Pro | N | Se | Hex-2-ynyl |
| Cofactor **28ab** | *cis*-L-Pro | N | S | Methyl |
| Cofactor **30ab** | *cis*-L-Pro | N | Se | Methyl |
| Cofactor **31b** | *cis*-L-Pro | N | Se | Hexyl |
| Cofactor **33b** | *cis*-L-Pro | N | Se | Propargyl |
| Cofactor **35b** | *cis*-L-Pro | N | Se | Octa-2,7-diynyl |
| Cofactor **37b** | L-Abu | N | S | Hex-2-ynyl |
| Cofactor **41** | *cis*-L-Pro | N | S | Hex-2-ynyl-PEG-MeSiR |
| Cofactor **42** | *cis*-L-Pro | N | Se | Hex-2-ynyl-PEG-MeSiR |
| Cofactor **43** | *cis*-L-Pro | N | Se | Hexyl-PEG-MeSiR |
| Cofactor **44** | L-Abu | N | S | Hex-2-ynyl-PEG-MeSiR |

**Figure S22.** Activity of wild-type M.TaqI with cofactor analogs and a single-target-site oligonucleotide duplex DNA substrate under single-turnover conditions. (A) Sequence of 23-mer duplex oligodeoxynucleotide. Target site is highlighted in bold. (B) Calculated number of adenines modified per target site by M.TaqI. M.TaqI (15 µM) was incubated with a 23-mer duplex oligodeoxynucleotide substrate (12.5 µM) containing a single TCGA site and 50 µM AdoMet or its analog for 1 h at 65 °C. Following the reaction, DNA was hydrolyzed to deoxynucleosides and analyzed by LC/MS (N = 2). Complete conversion corresponds to modification of two adenines per substrate molecule.

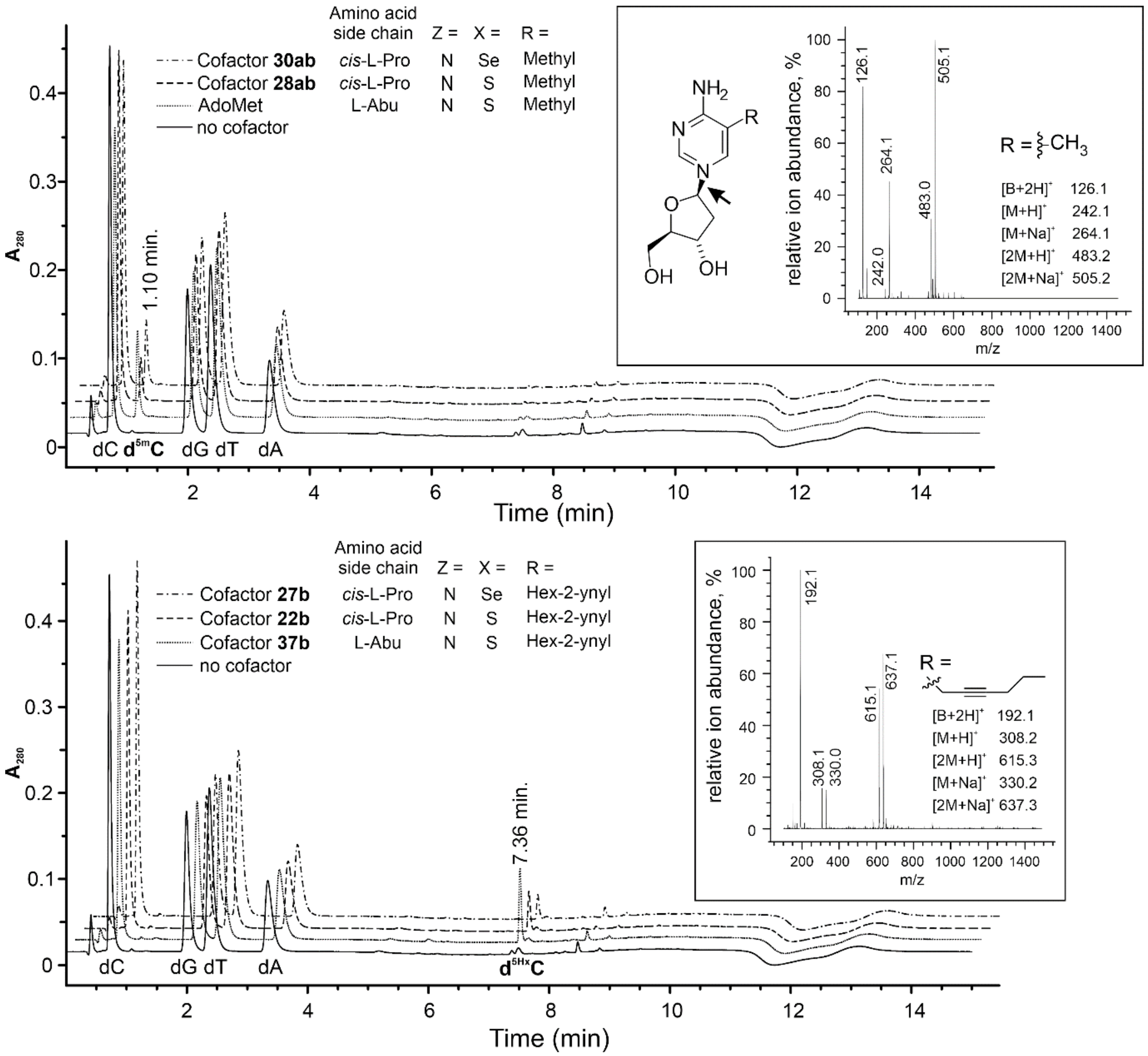


**Figure S23.** LC/MS analysis and identification of products generated by the M.HhaI Q82A/Y254S/N304A variant in reactions containing duplex DNA and AdoMet or its analogs. Excess of M. M.HhaI Q82A/Y254S/N304A enzyme was incubated with 22-mer duplex DNA substrate and 50 µM AdoMet or its analog for 1 h at 37°C, then DNA was hydrolyzed to deoxynucleosides and the products were analyzed by LC/MS. Retention times of $d^{5m}C$ and $d^{5Hx}C$ are indicated above the peaks. Insets show general structure of MTase reaction product and background-subtracted mass spectra at the apex of the corresponding peak. Arrow denotes fragmentation inside the mass spectrometer. Theoretical m/z values are given: $[M+H]^+$ – protonated molecular ion, $[M+Na]^+$ – sodium complex with molecular ion, $[2M+H]^+$ – protonated dimer ion, $[2M+Na]^+$ – sodium complex with dimer ion, $[B+2H]^+$ – protonated nucleobase formed upon fragmentation in the mass spectrometer.

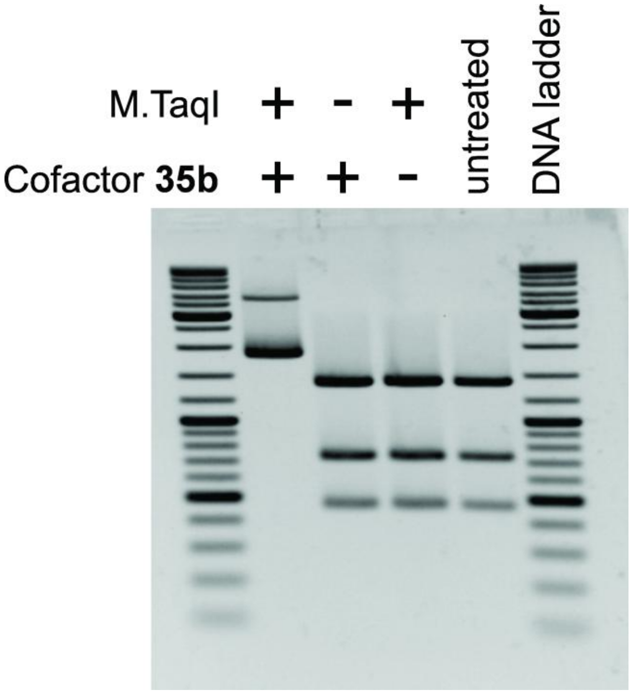


**Figure S24**. Efficient protection of pUC19 DNA from R.TaqI digestion in the reaction with M.TaqI and cofactor **35b**.

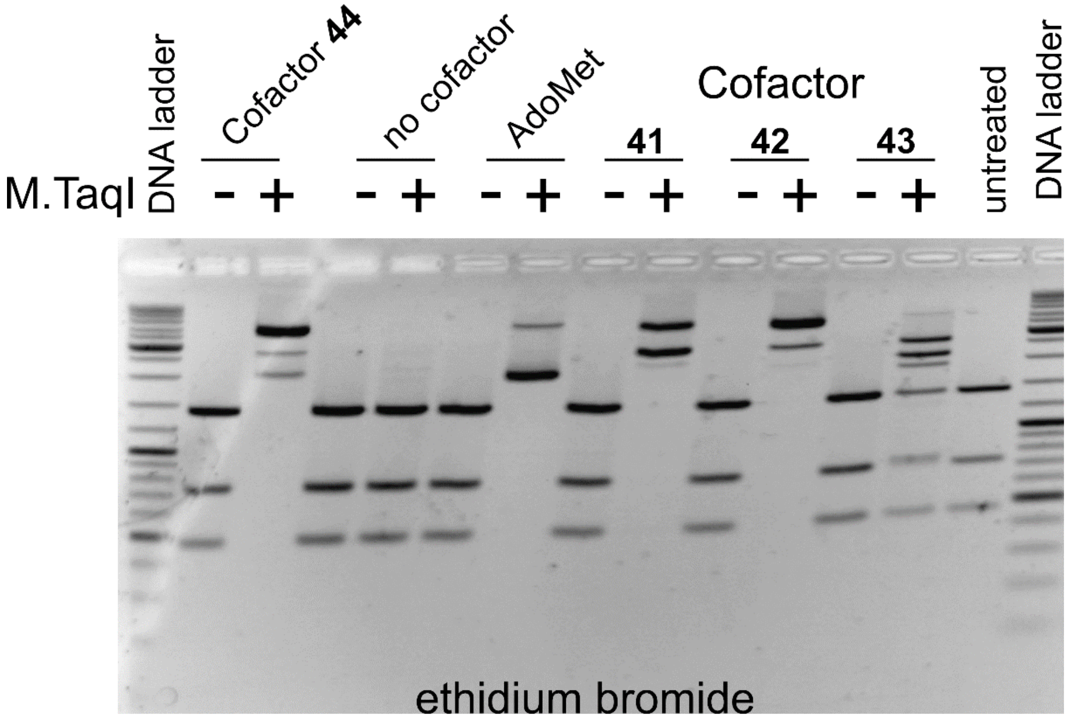


| | Amino acid side chain | Z = | X = | R = |
|---|---|---|---|---|
| AdoMet | L-Abu | N | S | Methyl |
| Cofactor **41** | *cis*-L-Pro | N | S | Hex-2-ynyl-PEG-MeSiR |
| Cofactor **42** | *cis*-L-Pro | N | Se | Hex-2-ynyl-PEG-MeSiR |
| Cofactor **43** | *cis*-L-Pro | N | Se | Hexyl-PEG-MeSiR |
| Cofactor **44** | L-Abu | N | S | Hex-2-ynyl-PEG-MeSiR |

**Figure S25**. Protection of pUC19 DNA from R.TaqI digestion in the reaction with M.TaqI and fluorescent cofactors **41** – **44**. pUC19 DNA was incubated with 25 µM cofactor and M.TaqI for 1 h at 65°C and then digested with R.TaqI. Due to different cofactor activities (Figure 2D) different M.TaqI:DNA ratios were used. [M.TaqI]:TCGA of 1:32 was used for AdoMet and cofactors **42**and **44**; [M.TaqI]:TCGA of 1:4 was used in “no cofactor” control and for cofactors **41** and **43**.

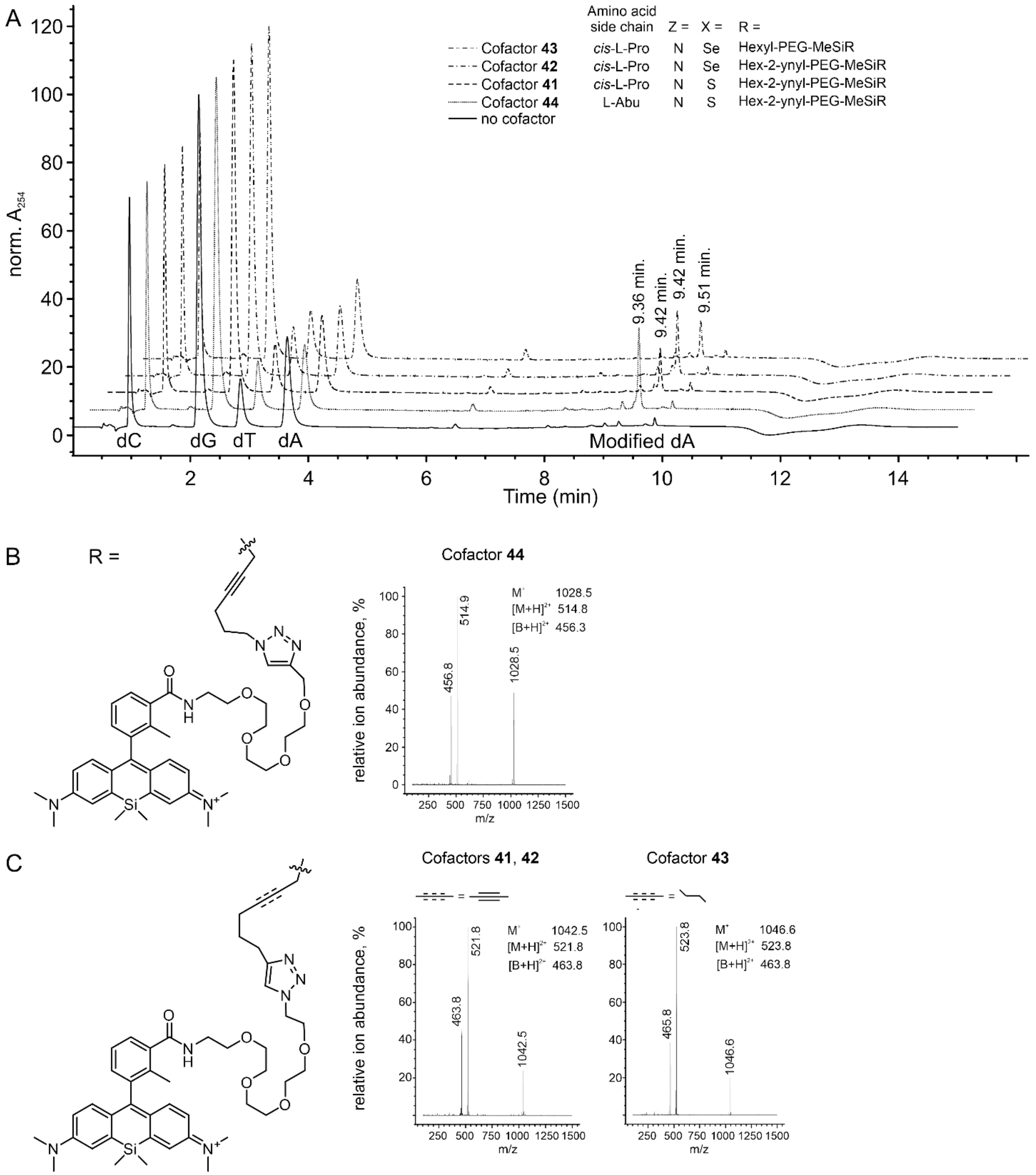


**Figure S26.** LC/MS analysis and identification of products formed in the reaction of M.TaqI and fluorescent cofactors **41** - **44**. (**A**) HPLC separation of deoxyribonucleosides. Retention times of dA modified by M.TaqI are indicated above the peaks. (**B**) structure of transferrable chain in cofactor **44** and mass spectra of the obtained deoxyadenosine derivative. (**C**) Structure of transferrable chain in cofactor **41** - **43** and mass spectra of the obtained deoxyadenosine derivatives. Theoretical m/z values are given. $M^+$ – molecular ion, $[M+H]^{2+}$ – protonated molecular ion, $[B+H]^{2+}$ – protonated nucleobase formed upon fragmentation in the mass spectrometer.

### Supplementary Methods

**HPLC-MS Analysis of Cofactor Decomposition Kinetics and Products:**

The stability of the synthesized AdoMet analogues was evaluated in 50 mM Tris-HCl buffer pH = 8.0 at 37°C at a concentration of 500 µM. The required amount of cofactor was diluted into 37°C preheated 50 mM Tris-HCl buffer and analyzed by HPLC every indicated time point (different time points were used in accordance to the stability of cofactors). Samples were eluted with a gradient of solvents A (10mM $HCOONH_4$, pH 3.6) and B (MeOH) (0-2 min isocratic 100% A, 2-9 min linear gradient from 100% to 0%, 9-10 min linear gradient from 0% to 100% A and 10-15 min isocratic 100% A at 0.4 ml/min flow) and detected by a diode array UV absorbance detector at 254 nm followed by ESI single quadruple mass detection. The half-life times of the cofactors were obtained by fitting to exponential one phase decay fit as implemented in GraphPad 9.0 software:

$$Y=(Y_0 - \text{Plateau})*\exp(-k*t) + \text{Plateau} \qquad (1)$$

where Y- area of cofactor absorbance peak at 254 nm at time point "t", $Y_0$ – area of cofactor absorbance peak at 254 nm at time point "0", equal to 100%; Plateau – area of cofactor absorbance peak at 254 nm at time point "infinity", equal to 0%; t – time, k – decay rate.

Pseudo-first-order rate decomposition constants $k_{total}$ were obtained from the negative value of slope of simple linear regression fit:

$$Y = b - k_{total} \cdot t \qquad (2)$$

where Y= $\ln[\text{cofactor}]_t$, and b = $\ln[\text{cofactor}]_0$, t – time and $k_{total}$ – cofactor decay rate

Pseudo first-order decay rate constants of intramolecular cyclization ($k_{Nuc}$), depurnation ($k_{Dep}$) and hydrolysis ($k_{Hyd}$) were obtained from the slope of the linear part of the linear regression fits:

$k_{Nuc}$ = slope of $ln\frac{[Cofactor]_0}{[Cofactor]_0-[AlkylAdenosine\ or\ Alkyl(Se)Adenosine]_t}$ vs time

$k_{Dep}$ =slope of $ln\frac{[Cofactor]_0}{[Cofactor]_0-[Adenine\ or\ deazaAdenine]_t}$ vs time

$k_{Hyd}$ = slope of $ln\frac{[Cofactor]_0}{[Cofactor]_0-[AdoCisLPro\ or\ (Se)AdoCisLPro]_t}$ vs time

**DNA protection assay**

*M.TaqI activity.* A series of reactions were performed that contained a constant amount of λ DNA (*dam*$^-$ *dcm*$^-$, ThermoFisher) and 2-fold serial dilutions of M.TaqI in the reaction buffer (50 mM Tris, 50 mM MOPS, 0.2 mg/ml BSA, pH 8.0). Reactions were assembled on ice by mixing 10 µl of solution containing

200 µM cofactor and 1.2 µg λ DNA with 10 µl of solution containing M.TaqI, at serial 2-fold dilutions starting from 460 nM. The final reaction mixture contained 230 nM TCGA sites and thereby [target] : [MTase] ratios from 1:1 to 512:1 were probed. The samples were incubated for 1 h at 65°C, then 10 µl of 3× CutSmart buffer (150 mM Potassium Acetate, 60 mM Tris-acetate, 30 mM Magnesium Acetate, 0.3 mg/ml BSA, pH 7.9) containing 2.5 units of R.TaqI-v2 (New England Biolabs) was added, and the incubation continued for 1 h at 65°C. The reaction products were fractionated in 0.8% agarose gel and MTase turnover number ($h^{-1}$) was read as maximal [TCGA] : [M.TaqI] ratio still allowing full protection against fragmentation by R.TaqI-v2.

*M.HhaI Q82A/Y254S/N304A activity.* A series of reactions were performed that contained a constant amount of λ DNA (*dam*$^-$ *dcm*$^-$, ThermoFisher) and serial 2-fold dilutions of M.HhaI Q82A/Y254S/N304A in the reaction buffer (50 mM Tris, 50 mM MOPS, 0.2 mg/ml BSA, 2 mM β-mercaptoethanol, pH 7.5). Reactions were assembled on ice by mixing 10 µl of solution containing 200 µM cofactor and 1.2 µg λ DNA with 10 µl of solution containing M.HhaI Q82A/Y254S/N304A at serial 2-fold dilutions, starting from 430 nM. The final reaction mixture contained 215 nM GCGC sites and thereby [target] : [MTase] ratios from 1:1 to 512:1 were probed. The samples were incubated for 1 h at 37°C, MTase was inactivated by heating at 65°C for 15 min. and 10 µl of 3× Tango buffer (99 mM Tris-acetate (pH 7.9 at 37°C), 30 mM magnesium acetate, 198 mM potassium acetate, 0.3 mg/ml BSA; Thermo Fisher Scientific, #BY5) containing 2.5 units of R.Hin6I (Thermo Fisher Scientific, #ER0481) was added, and the incubation continued for 1 h at 37°C. The reaction products were fractionated in 0.8% agarose gel and MTase turnover number ($h^{-1}$) was read as maximal [GCGC] : [M.HhaI Q82A/Y254S/N304A] ratio still allowing full protection against fragmentation by R.Hin6I .

**Identification of MTase-catalyzed cytosine and adenine modifications by LC/MS**

Desalted oligodeoxyribonucleotides with a single MTase site were synthesized by Sigma. The duplex DNA substrates (Tabel S1) were produced by mixing equal molar amounts of complementary single-stranded oligodeoxynucleotides in water, heating up to 95°C in 500ml water bath and allowing to cool down slowly to room temperature overnight.

Table S1. DNA MTase substrates.

MTase target sites are in bold, modified bases underlined

| | |
|---|---|
| M.TaqI | 5'-CCCGCCGAAG**TCGA**GCGCCGAAC<br>GGGCGGCTTC**AGCT**CGCGGCTTG-5' |
| M.HhaI Q82A/Y254S/N304A | 5'-GCTATTATT**GCGC**TATTATTGC<br>CGATAATAA**CGCG**ATAATAACG-5' |

Enzymatic modifications were performed by incubating duplex oligodeoxynucleotide (12.5 µM) with 50 µM cofactor and 15 µM MTase in 40 µl of corresponding reaction buffer for 1h at 65°C (M.TaqI) or at 37°C (M.HhaI Q82A/Y254S/N304A). After incubation, M.HhaI Q82A/Y254S/N304A was inactivated by heating the sample for 15 min. at 65°C. MTase was digested by proteinase K for 1h at 55°C. Afterwards, the samples were diluted to 100 µl and separated from unreacted cofactor and protein digestion products by passing through MicroSpin™ Sephadex™ G-25 columns (cat. No: 27532501, Cytiva, USA) equilibrated with water. 10 µl of 10× Nucleoside Digestion Mix buffer and 1 µl of Nucleoside Digestion Mix (New England Biolabs, #M0649S) were added and DNA was hydrolyzed to nucleosides for 2 h at 37°C. The resulting nucleoside mixture was analyzed on Agilent 1260 Infinity II LC system using reversed-phase UHPLC column (Supelco Titan™ C18, 1.9 µm, 7.5 cm × 2.1 mm, Sigma-Aldrich, Germany) equipped with a pre-column (Supelco Titan™ C18, 1.9 µm, 0.5 cm × 2.1 mm, Sigma-Aldrich, Germany). Compounds were eluted with methanol (5% for 2 min, followed by linear gradient to 100% in 6 min, hold 100% for 1 min, followed by linear gradient to 5% in 1 min and hold 5% for 5 min) in ammonium formate buffer (25 mM, pH 3.5) at a flow of 0.4 mL/min and detected using absorbance DAD detector set at 254 nm (scanning 210 – 850 nm range) and single quadruple LC/MSD XT mass spectrometer scanning 100 - 1500 m/z range. An extent of deoxyadenosine or deoxycytosine modification was measured as decrease in respective peak areas after normalization to deoxyguanosine peak to account for variations in DNA recovery. Identities of modification products were confirmed by mass spectrometry.

**Chemo-enzymatic two-step labeling of plasmid DNA:**

16 µg of pUC19 DNA (New England Biolabs) was incubated with 144 nM M.TaqI and 100 µM cofactor 42 in 320 µl of reaction buffer for 1 h at 65°C. This corresponds to M.TaqI : targets ratio of 1:8. MTase was degraded by incubation with proteinase K at 50°C for 30 min., followed by extraction with equal volume of phenol : chloroform : isoamyl alcohol (50:49:1) mixture, then with equal volume of chloroform and ethanol precipitation. DNA modification was confirmed by protection from R.TaqI-v2 (New England Biolabs, # R0149S) (Figure S24).

2 µg of modified DNA was incubated in 50 µl of 100 mM sodium phosphate pH 7.0 containing 50 µM CalFluor 647 azide (Click Chemistry Tools, #1372), 2 mM sodium ascorbate and 10 mM BTTES with 2 mM copper (II) acetate for 1 h at room temperature. The reaction mixture was diluted 2-fold, mixed with 500 µl of binding buffer and applied onto MSB® Spin PCRapace column (Invitek, # 1020220400). The columns were washed once with binding buffer and DNA was eluted in 30 µl of water. 200 ng of DNA was digested with Fast Digest MbiI restriction endonuclease (Thermo Fisher) and fractionated on agarose gel. MTase-

dependent labeling of fragments containing M.TaqI sites was confirmed by imaging fluorescence with 630 nm LED excitation. Total DNA was visualized by ethidium bromide staining.

**Enzymatic single-step labeling of plasmid DNA:**

4 µg of pUC19 DNA (New England Biolabs, # N3041L) in 40 µl reaction buffer was incubated with 58 nM (no cofactor, cofactors 52 and 54) or 7.2 nM (cofactor 53) M.TaqI and 25 µM cofactor for 1 h at 65°C. This corresponds to M.TaqI : targets ratio of 1:4 and 1:32. Then, the reaction mixture was diluted to 200 µl with water and incubated with proteinase K for 30 min. at 50°C. DNA was mixed with 1 ml of binding buffer and purified with QIAquick PCR Purification Kit (Qiagen, # 28104). 400 ng of DNA was digested with Fast Digest MbiI restriction endonuclease (Thermo Fisher, # FD1274) and fractionated on agarose gel. MTase-dependent labeling of fragments containing M.TaqI sites was confirmed by imaging fluorescence with 630 nm LED excitation. Total DNA was visualized by ethidium bromide staining. Effectiveness of DNA modification was assessed by protection from R.TaqI digestion (Figure S25).

**General chemical experimental information and synthesis methods**

NMR spectra were recorded at 25 °C with an Agilent 400-MR spectrometer at 400.06 MHz ($^{1}H$) and 100.60 MHz ($^{13}C$) and are reported in ppm. All $^{1}H$ and $^{13}C$ spectra are referenced to tetramethylsilane (δ = 0 ppm) using the residual signals of the solvents according to the values reported in literature[2]. Multiplicities of signals are described as follows: s = singlet, d = doublet, t = triplet, q = quartet, p = pentet, m = multiplet or overlap of non-equivalent resonances; br = broad signal. Coupling constants (*J*) are given in Hz. All NMR spectra were processed with MestRenova 11.0.4 software.

ESI-MS were recorded on a Varian 500-MS spectrometer (Agilent). ESI-HRMS were recorded on a MICROTOF spectrometer (Bruker) equipped with ESI ion source (Apollo) and direct injector with LC autosampler Agilent RR 1200.

Analytical LC–MS analysis was performed on an Agilent 1260 Infinity II LC/MS system equipped with an autosampler, diode array detector WR, fluorescence detector Spectra and Infinity Lab LC/MSD 6100 series quadruple with API electrospray. Analysis was done by using an Ascentis® Express AQ-C18 UHPLC Column 2 µm, 5 cm x 2.1 mm column with A: 25 mM $HCOONH_4$ (pH = 3.6) aqueous buffer and B: MeOH

Preparative HPLC was performed on a combined Agilent1260/1290 Infinity II preparative system equipped with an 1290 Infinity II open-bed sampler (G7169B)/fraction collector (G7159B), 1290 Infinity II preparative binary pump (G7161A), 1260 Infinity II multiple wavelength detector (G7165A) and with Agilent Pursuit 10 $C_{18}$, 10 µm, 250 X 50 mm preparative column.

**1-(tert-butyl) 2-methyl (2S,4R)-4-((methylsulfonyl)oxy)pyrrolidine-1,2-dicarboxylate**

MsO-MeO-Boc-*trans*-*L*-Pro (**1**):

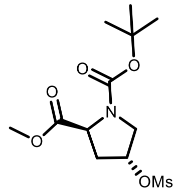


MsO-MeO-Boc-*trans*-*L*-Pro (**1**)

To a solution of N-Boc-trans-4-hydroxy-L-proline methyl ester (1.5 g, 6.1 mmol) in dry DCM (50 mL) DIPEA (1.27 mL, 7.3 mmol) was added and the mixture was cooled to 0$^o$C in an ice bath. Then MsCl (0.565 mL, 7.3 mmol) was added dropwise in the course of 15 min. The reaction mixture was stirred at room temperature for 2h and the end of the reaction was monitored by TLC (EtOAc:DCM 3:7, $R_f$ = 0.6, stained with phosphomolybdic acid). The mixture was washed with 1M HCl, water and brine, the organic phase was dried with $Na_2SO_4$ and the solvent was evaporated to dryness. The residue was purified by flash column chromatography (Teledyne Isco RediSep Rf 40 g; gradient 10% to 80% DCM – EtOAc) to give 1.67 g of yellow oil in a 85% yield.

$^1$H NMR (400 MHz, chloroform-*d*) rotamer mixture (ratio 1:0.7), δ (ppm): 5.29 – 5.22 (m, 1H), 4.46 and 4.40 (t, *J* = 7.7 Hz and t, *J* = 8.0 Hz 1H), 3.88 – 3.72 (m, 5H), 3.05 (s, 3H), 2.71 – 2.50 (m, 1H), 2.30 – 2.21 (m, 1H), 1.46 and 1.41 (s, 9H).

$^{13}$C NMR (101 MHz, chloroform-*d*) rotamer mixture (ratio 1:0.7), major rotamer peaks reported δ (ppm) 172.8, 153.4, 81.1, 78.0, 57.6, 52.4, 52.3, 38.9, 37.6, 28.3.

ESI-MS, positive mode: m/z = 346.1 [M+Na]$^+$.

HRMS (ESI) calculated for $C_{12}H_{22}NO_7S$ [M+H]$^+$ 324.1111, found 324.1124.

**1-(tert-butyl) 2-methyl (2S,4S)-4-((methylsulfonyl)oxy)pyrrolidine-1,2-dicarboxylate**

MsO-MeO-Boc-*cis*-*L*-Pro (**2**):

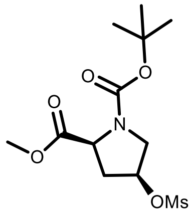


MsO-MeO-Boc-*cis*-*L*-Pro (**2**)

To a solution of N-Boc-cis-4-hydroxy-L-proline methyl ester (1.5 g, 6.1 mmol) in dry DCM (50 mL) DIPEA (1.27 mL, 7.3 mmol) was added and the mixture was cooled to 0$^o$C in an ice bath. Then MsCl (0.565 mL, 7.3 mmol) was added dropwise in the course of 15 min. The reaction mixture was stirred at room temperature for 2h and the end of the reaction was monitored by TLC (EtOAc:DCM 3:7, $R_f$ = 0.6, stained with phosphomolybdic acid). The mixture was washed with 1M HCl, water and brine, organic phase was dried with $Na_2SO_4$ and the solvent was evaporated to dryness. The residue was purified by flash column chromatography (Teledyne Isco RediSep Rf 40 g; gradient 10% to 80% DCM – EtOAc) to give 1.85 g of yellow oil in a 94% yield.

$^{1}H$ NMR (400 MHz, chloroform-*d*) rotamer mixture (ratio 1:0.8), δ (ppm): 5.26 – 5.19 (m, 1H), 4.51 and 4.39 (dd, *J* = 8.3, 3.5 Hz and t, *J* = 6.0 Hz, 1H), 3.81 – 3.72 (m, 5H), 3.00 (s, 3H), 2.57 – 2.42 (m, 2H), 1.47 and 1.42 (s, 9H).

$^{13}C$ NMR (101 MHz, chloroform-*d*) rotamer mixture (ratio 1:0.8), only major rotamer peaks reported δ (ppm) 172.1, 153.5, 80.8, 77.6, 57.5, 52.6, 52.1, 39.0, 37.3, 28.4.

ESI-MS, positive mode: m/z = 346.1 $[M+Na]^+$.

HRMS (ESI) calculated for $C_{12}H_{22}NO_7S$ $[M+H]^+$ 324.1111, found 324.1108.

**1-(tert-butyl) 2-methyl (2R,4S)-4-((methylsulfonyl)oxy)pyrrolidine-1,2-dicarboxylate**

MsO-MeO-Boc-*trans*-*D*-Pro (**3**):

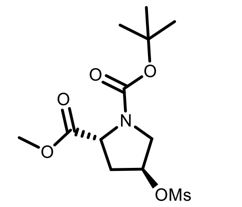

MsO-MeO-Boc-*trans*-*D*-Pro (**3**)

To a solution of N-Boc-trans-4-hydroxy-D-proline methyl ester (1.5 g, 6.1 mmol) in dry DCM (50 mL) DIPEA (1.27 mL, 7.3 mmol) was added and the mixture was cooled to 0°C in an ice bath. Then MsCl (0.565 mL, 7.3 mmol) was added dropwise in the course of 15 min. The reaction mixture was stirred at room temperature for 2h, the end of the reaction was monitored by TLC (EtOAc:DCM 3:7, $R_f$ = 0.6, stained with phosphomolybdic acid). The mixture was washed with 1M HCl, water and brine, the organic phase was dried with $Na_2SO_4$ and the solvent was evaporated to dryness. The residue was purified by flash column chromatography (Teledyne Isco RediSep Rf 40 g; gradient 10% to 80% DCM – EtOAc) to give 1.71 g of yellow oil in a 87% yield.

$^{1}H$ NMR (400 MHz, chloroform-*d*) rotamer mixture (ratio 1:0.7), δ (ppm): 5.27 – 5.22 (m, 1H), 4.45 and 4.39 (t, *J* = 7.7 Hz and t, *J* = 8.0 Hz, 1H), 3.86 – 3.72 (m, 5H), 3.04 (s, 3H), 2.70 – 2.50 (m, 1H), 2.30 – 2.20 (m, 1H), 1.45 and 1.41 (s, 9H).

$^{13}C$ NMR (101 MHz, chloroform-*d*) rotamer mixture (ratio 1:0.7), only major rotamer peaks reported δ (ppm) δ 172.8, 153.4, 81.0, 78.0, 57.6, 52.6, 52.3, 38.9, 37.6, 28.3.

ESI-MS, positive mode: m/z = 346.1 $[M+Na]^+$.

HRMS (ESI) calculated for $C_{12}H_{22}NO_7S$ $[M+H]^+$ 324.1111, found 324.1117.

**1-(tert-butyl) 2-methyl (2R,4R)-4-((methylsulfonyl)oxy)pyrrolidine-1,2-dicarboxylate**

MsO-MeO-Boc-*cis*-*D*-Pro (**4**):

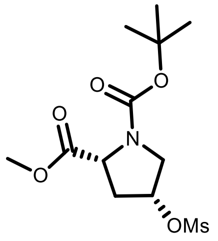

MsO-MeO-Boc-*cis*-*D*-Pro (**4**)

To a solution of N-Boc-cis-4-hydroxy-D-proline methyl ester (1.5 g, 6.1 mmol) in dry DCM (50 mL) DIPEA (1.27 mL, 7.3 mmol) was added and the mixture was cooled to 0°C in an ice bath. Then MsCl (0.565 mL, 7.3 mmol) was added dropwise in the course of 15 min. The reaction mixture was stirred at room temperature for 2h and the end of the reaction was monitored by TLC (EtOAc:DCM 3:7, $R_f$ = 0.6, stained with phosphomolybdic acid). The mixture was washed with 1M HCl, water and brine, organic phase was dried with $Na_2SO_4$ and the solvent was evaporated to dryness. The residue was purified by flash column chromatography (Teledyne Isco RediSep Rf 40 g; gradient 10% to 80% DCM – EtOAc) to give 1.79 g of yellow oil in a 91% yield.

$^1$H NMR (400 MHz, chloroform-*d*) rotamer mixture (ratio 1:0.9), δ (ppm): 5.27 – 5.15 (m, 1H), 4.51 and 4.39 (dd, *J* = 8.3, 3.5 Hz and t, *J* = 6.0 Hz, 1H), 3.80 – 3.73 (m, 5H), 3.00 (s, 3H), 2.56 – 2.46 (m, 2H), 1.47 and 1.42 (s, 9H).

$^{13}$C NMR (101 MHz, chloroform-*d*) rotamer mixture (ratio 1:0.9), only major rotamer peaks reported δ (ppm) 172.0, 153.5, 80.8, 77.6, 57.5, 52.6, 52.4, 39.0, 37.3, 28.4.

ESI-MS, positive mode: m/z = 346.1 $[M+Na]^+$.

HRMS (ESI) calculated for $C_{12}H_{22}NO_7S$ $[M+H]^+$ 324.1111, found 324.1120.

**1-(tert-butyl) 2-methyl (2S,4S)-4-(acetylthio)pyrrolidine-1,2-dicarboxylate**

SAc-MeO-Boc-*cis*-*L*-Pro (**5**):

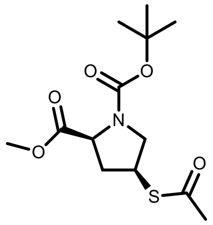

SAc-MeO-Boc-*cis*-*L*-Pro (**5**)

To a solution of **1** (1.6 g, 4.95 mmol ) in dry DMF (40 mL) under an argon atmosphere, potassium thioacetate (1.13 g, 9.9 mmol) was added. The reaction mixture was stirred at 70°C overnight. Then the mixture was diluted with EtoAc (100 mL) and washed with water (3 x 100 mL) and brine (100 mL). The organic phase was dried over anhydrous $Na_2SO_4$ and the solvent was evaporated to dryness. The residue was purified by flash column chromatography (Teledyne Isco RediSep Rf 40 g; gradient 5% to 80% Hexane – EtOAc) to give 1.13 g of orange oil in a 75% yield.

$^1$H NMR (400 MHz, chloroform-*d*) rotamer mixture (ratio 1:0.8), δ (ppm): 4.36 and 4.28 (t, *J* = 7.4 Hz and t, *J* = 7.6 Hz, 1H), 4.03 – 3.90 (m, 2H), 3.73 (s, 3H), 3.39 – 3.27 (m, 1H), 2.79 – 2.63 (m, 1H), 2.32 (s, 3H), 2.01 – 1.90 (m, 1H), 1.45 and 1.40 (s, 9H).

$^{13}C$ NMR (101 MHz, chloroform-*d*) rotamer mixture (ratio 1:0.8), only major rotamer peaks reported δ (ppm) 194.8, 172.9, 153.3, 80.6, 58.7, 52.3, 51.4, 38.8, 37.1, 30.7, 28.4.

ESI-MS, positive mode: m/z = 326.1 $[M+Na]^+$.

HRMS (ESI) calculated for $C_{13}H_{22}NO_5S$ $[M+H]^+$ 304.1213, found 304.1208.

**1-(tert-butyl) 2-methyl (2S,4R)-4-(acetylthio)pyrrolidine-1,2-dicarboxylate**

SAc-MeO-Boc-*trans-L*-Pro (**6**):

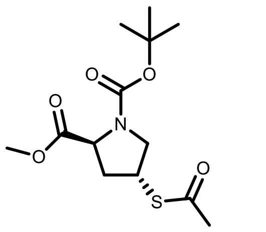

SAc-MeO-Boc-*trans-L*-Pro (**6**)

To a solution of **2** (1.8 g, 5.57 mmol ) in dry DMF (30 mL) under an argon atmosphere, potassium thioacetate (1.27 g, 11.14 mmol) was added. The reaction mixture was stirred at 70°C overnight. Then the mixture was diluted with EtoAc (100 mL) and washed with water (3 x 100 mL) and brine (100 mL). The organic phase was dried over anhydrous $Na_2SO_4$ and the solvent was evaporated to dryness. The residue was purified by flash column chromatography (Teledyne Isco RediSep Rf 40 g; gradient 5% to 80% Hexane – EtOAc) to give 1.15 g of orange oil in a 68% yield.

$^{1}H$ NMR (400 MHz, chloroform-*d*) rotamer mixture (ratio 1:0.8), δ (ppm): 4.39 and 4.29 (dd, *J* = 8.6, 4.2 Hz and dd, *J* = 8.4, 5.4 Hz, 1H), 4.08 – 3.99 (m, 1H), 3.93 (dd, *J* = 11.2, 7.0 Hz, 1H), 3.74 (s, 3H), 3.42 and 3.30 (dd, *J* = 11.3, 5.4 Hz and dd, *J* = 11.0, 6.4 Hz, 1H), 2.49 – 2.34 (m, 1H), 2.32 (s, 3H), 2.29 – 2.14 (m, 1H), 1.44 and 1.4 (s, 9H).

$^{13}C$ NMR (101 MHz, chloroform-*d*) rotamer mixture (ratio 1:0.8), only major rotamer peaks reported δ (ppm) 194.8, 173.0, 153.5, 80.6, 58.6, 52.3, 51.6, 39.6, 37.0, 30.7, 28.4.

ESI-MS, positive mode: m/z = 326.1 $[M+Na]^+$.

HRMS (ESI) calculated for $C_{13}H_{22}NO_5S$ $[M+H]^+$ 304.1213, found 304.1211.

**1-(tert-butyl) 2-methyl (2R,4R)-4-(acetylthio)pyrrolidine-1,2-dicarboxylate**

SAc-MeO-Boc-*cis-D*-Pro (**7**):

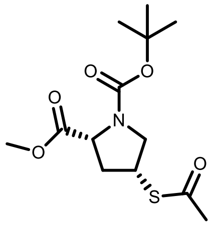

SAc-MeO-Boc-*cis-D*-Pro (**7**)

To a solution of **3** (1.7 g, 5.26 mmol ) in dry DMF (30 mL) under an argon atmosphere, potassium thioacetate (1.2 g, 10.5 mmol) was added. The reaction mixture was stirred at 70°C overnight. Then the mixture was diluted with EtoAc (100 mL) and washed with water (3 x 100 mL) and brine (100 mL). The organic phase was dried over anhydrous $Na_2SO_4$ and the solvent was evaporated to dryness. The residue was purified by flash column chromatography (Teledyne Isco RediSep Rf 40 g; gradient 5% to 80% Hexane – EtOAc) to give 0.98 g of orange oil in a 62% yield.

$^{1}$H NMR (400 MHz, chloroform-*d*) rotamer mixture (ratio 1:0.8), δ (ppm): 4.35 and 4.27 (t, *J* = 7.4 Hz, and t, *J* = 7.6 Hz, 1H), 4.01 – 3.89 (m, 2H), 3.73 (s, 3H), 3.39 – 3.26 (m, 1H), 2.78 – 2.62 (m, 1H), 2.31 (s, 3H), 2.00 – 1.89 (m, 1H), 1.44 and 1.4 (s, 9H).

$^{13}$C NMR (101 MHz, chloroform-*d*) rotamer mixture (ratio 1:0.8), only major rotamer peaks reported δ (ppm) 194.8, 172.9, 153.3, 80.6, 58.7, 52.3, 51.4, 38.8, 37.1, 30.6, 28.4.

ESI-MS, positive mode: m/z = 326.1 $[M+Na]^+$.

HRMS (ESI) calculated for $C_{13}H_{22}NO_5S$ $[M+H]^+$ 304.1213, found 304.1211.

**1-(tert-butyl) 2-methyl (2R,4S)-4-(acetylthio)pyrrolidine-1,2-dicarboxylate**

SAc-MeO-Boc-*trans*-*D*-Pro (**8**):

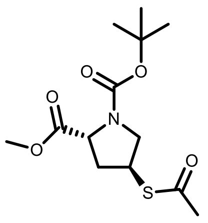


SAc-MeO-Boc-*trans*-*D*-Pro (**8**)

To a solution of **4** (1.75 g, 5.42 mmol ) in dry DMF (30 mL) under an argon atmosphere, potassium thioacetate (1.24 g, 10.84 mmol) was added. The reaction mixture was stirred at 70°C overnight. Then the mixture was diluted with EtoAc (100 mL) and washed with water (3 x 100 mL) and brine (100 mL). The organic phase was dried over anhydrous $Na_2SO_4$ and the solvent was evaporated to dryness. The residue was purified by flash column chromatography (Teledyne Isco RediSep Rf 40 g; gradient 5% to 80% Hexane – EtOAc) to give 1.28g of orange oil in a 78% yield.

$^{1}$H NMR (400 MHz, chloroform-*d*) rotamer mixture (ratio 1:0.8), δ (ppm): 4.38 and 4.29 (dd, *J* = 8.6, 4.3 Hz and dd, *J* = 8.4, 5.3 Hz, 1H), 4.07 – 3.98 (m, 1H), 3.92 (dd, *J* = 11.2, 7.0 Hz, 1H), 3.73 (s, 3H), 3.41 and 3.29 (dd, *J* = 11.3, 5.4 Hz and dd, *J* = 11.0, 6.4 Hz, 1H), 2.43 – 2.33 (m, 1H), 2.31 (s, 3H), 2.29 – 2.13 (m, 1H), 1.44 and 1.39 (s, 9H).

$^{13}$C NMR (101 MHz, chloroform-*d*) rotamer mixture (ratio 1:0.8), only major rotamer peaks reported δ (ppm) 194.8, 173.0, 153.4, 80.6, 58.6, 52.3, 51.6, 39.6, 37.0, 30.7, 28.3.

ESI-MS, positive mode: m/z = 326.1 $[M+Na]^+$.

HRMS (ESI) calculated for $C_{13}H_{22}NO_5S$ $[M+H]^+$ 304.1213, found 304.1212.

**1-(tert-butyl) 2-methyl (2S,4S)-4-selenocyanatopyrrolidine-1,2-dicarboxylate:**

SeCN-MeO-Boc-*cis*-*L*-Pro (**9**):

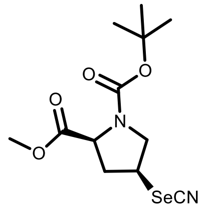


SeCN-MeO-Boc-*cis*-*L*-Pro (**9**)

To a solution of **1** (1.5 g, 4.64 mmol) in dry DMF (40 mL) under an argon atmosphere, KSeCN (1.67 g, 11.6 mmol) was added. The reaction mixture was heated to 90ºC and stirred overnight. Then the mixture was diluted with EtoAc (100 mL) and washed with water (3 x 100 mL) and brine (100 mL). The organic phase was dried over anhydrous $Na_2SO_4$ and the solvent was evaporated to dryness. The residue was purified by flash column chromatography (Teledyne Isco RediSep Rf 40 g; gradient 10% to 80% Hexane – EtOAc and Teledyne Isco RediSep Rf 40 g, gradient DCM:EtOAc 1% to 20%) to give 1.08 g of a light brown oil in a 70% yield.

$^1$H NMR (400 MHz, chloroform-*d*) rotamer mixture (ratio 1:0.7), only major rotamer peaks reported δ (ppm) 4.46 – 4.30 (m, 1H), 4.08 – 3.89 (m, 2H), 3.76 (s, 3H), 3.66 (dd, *J* = 11.7, 6.3 Hz, 1H), 2.95 – 2.79 (m, 1H), 2.41 – 2.25 (m, 1H), 1.41 (s, 9H).

$^{13}$C NMR (101 MHz, chloroform-*d*) rotamer mixture (ratio 1:0.7), only major rotamer peaks reported δ (ppm) 171.2, 153.1, 100.6, 81.2, 58.5, 58.2, 52.9, 38.8, 28.3.

ESI-MS, positive mode: m/z = 357.1 $[M+Na]^+$.

HRMS (ESI) calculated for $C_{12}H_{18}N_2O_4SeNa$ $[M+Na]^+$ 357.0324, found 357.0337.

**1-(tert-butyl) 2-methyl (2S,4S)-4-((((2S,3S,4R,5R)-5-(6-amino-9H-purin-9-yl)-3,4-dihydroxytetrahydro-furan-2-yl)methyl)thio)pyrrolidine-1,2-dicarboxylate**

Ado-MeO-Boc-*cis*-*L*-Pro (**10**):

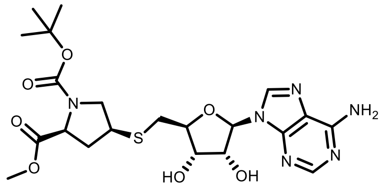


Ado-MeO-Boc-*cis*-*L*-Pro (**10**)

To a solution of **5** (0.75 g, 2.47 mmol) in dry MeOH (15 mL) under an argon atmosphere a solution of NaOMe (680 µL of 25% wt solution in MeOH, 2.47 mmol) was added and the resulting mixture was stirred for 30 min at room temperature. Then a solution of 5'-tosyladenosine (0.84 g, 2 mmol) in a 1:1 mixture of MeOH:MeCN (70 mL) was added and the reaction mixture was stirred under reflux overnight. The solvent was evaporated, the residue was dissolved in a mixture of DCM and MeOH, deposited on celite and purified by flash column chromatography (Interchim Puriflash 30 µm DCM:MeOH 2% to 25%) to afford 0.72 g in a 71% yield.

$^1$H NMR (400 MHz, Methanol-$d_4$) rotamer mixture (ratio 1:0.8), δ (ppm): 8.30 (s, 1H), 8.23 (s, 1H), 6.01 (d, *J* = 5.0 Hz, 1H), 4.84 – 4.79 (m, 1H), 4.36 (dt, *J* = 8.4, 5.1 Hz, 1H), 4.21 (dq, *J* = 9.6, 5.1, 4.6 Hz, 2H),

3.83 (ddd, *J* = 18.6, 10.8, 7.2 Hz, 1H), 3.73 (s, 1H), 3.71 (s, 2H), 3.52 – 3.38 (m, 1H), 3.21 – 3.11 (m, 1H), 3.12 – 2.95 (m, 2H), 2.70 – 2.56 (m, 1H), 1.94 – 1.81 (m, 1H), 1.44 and 1.40 (s, 9H).-$NH_2$ and 2x -OH is not present due to exchange with deuterium from $CD_3OD$.

$^{13}$C NMR (101 MHz, Methanol-*d*$_4$) rotamer mixture (ratio 1:0.8), only major rotamer peaks reported δ (ppm) 174.6, 157.3, 155.1, 153.9, 150.6, 141.5, 120.6, 90.3, 85.8, 81.8, 74.6, 74.0, 60.2, 53.8, 52.7, 42.4, 38.8, 35.0, 28.5.

ESI-MS, positive mode: m/z = 511.2 $[M+H]^+$.

HRMS (ESI) calculated for $C_{21}H_{31}N_6O_7S$ $[M+H]^+$ 511.1969, found 511.1977.

**1-(tert-butyl) 2-methyl (2S,4R)-4-((((2S,3S,4R,5R)-5-(6-amino-9H-purin-9-yl)-3,4-dihydroxytetrahydrofuran-2-yl)methyl)thio)pyrrolidine-1,2-dicarboxylate**

Ado-MeO-Boc-*trans*-*L*-Pro (**11**):

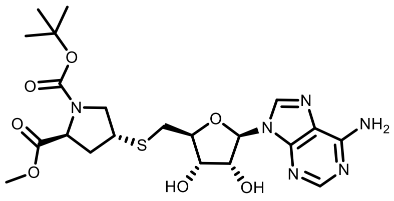


Ado-MeO-Boc-*trans*-*L*-Pro (**11**)

To a solution of **6** (0.8 g, 2.64 mmol) in dry MeOH (15 mL) under an argon atmosphere a solution of NaOMe (726 µL of 25% wt solution in MeOH, 2.64 mmol) was added and the resulting mixture was stirred for 30 min at room temperature. Then a solution of 5'-tosyladenosine (0.88 g, 2.1 mmol) in a 1:1 mixture of MeOH:MeCN (70 mL) was added and the reaction mixture was stirred under reflux overnight. The solvent was evaporated, the residue was dissolved in a mixture of DCM and MeOH, deposited on celite and purified by flash column chromatography (Interchim Puriflash 30 µm DCM:MeOH 2% to 25%) to afford 0.67 g in a 63% yield.

$^1$H NMR (400 MHz, DMSO-*d*$_6$) rotamer mixture (ratio 1:0.6), δ (ppm): 8.34 (s, 1H), 8.15 (s, 1H), 7.28 (s, 2H), 5.89 (d, *J* = 5.4 Hz, 1H), 5.51 (d, *J* = 6.0 Hz, 1H), 5.33 (t, *J* = 5.1 Hz, 1H), 4.73 (q, *J* = 5.6 Hz, 1H), 4.25 – 4.12 (m, 2H), 4.00 (p, *J* = 5.6, 4.9 Hz, 1H), 3.69 (td, *J* = 10.7, 6.8 Hz, 1H), 3.61 and 3.57 (s, 3H), 3.46 (h, *J* = 5.9, 5.4 Hz, 1H), 3.18 – 3.12 (m, 1H), 3.02 – 2.91 (m, 2H), 2.21 – 2.07 (m, 2H), 1.35 and 1.31 (s, 9H).

$^{13}$C NMR (101 MHz, DMSO-*d*$_6$) rotamer mixture (ratio 1:0.6), only major rotamer peaks reported δ (ppm): 172.6, 156.1, 153.2, 152.6, 149.4, 139.9, 119.2, 87.5, 84.0, 79.3, 72.6, 58.1, 52.7, 52.0, 41.0, 40.2, 36.6, 33.3, 27.8.

ESI-MS, positive mode: m/z = 511.2 $[M+H]^+$.

HRMS (ESI) calculated for $C_{21}H_{31}N_6O_7S$ $[M+H]^+$ 511.1969, found 511.1974.

**1-(tert-butyl) 2-methyl (2R,4R)-4-((((2S,3S,4R,5R)-5-(6-amino-9H-purin-9-yl)-3,4-dihydroxytetrahydro-furan-2-yl)methyl)thio)pyrrolidine-1,2-dicarboxylate**

Ado-MeO-Boc-*cis-D*-Pro (**12**):

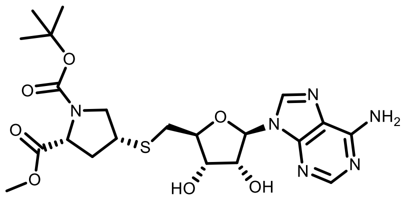

Ado-MeO-Boc-*cis-D*-Pro (**12**)

To a solution of **7** (0.9 g, 2.97 mmol) in dry MeOH (15 mL) under an argon atmosphere a solution of NaOMe (817 µL of 25% wt solution in MeOH, 2.97 mmol) was added and the resulting mixture was stirred for 30 min at room temperature. Then a solution of 5'-tosyladenosine (1.0 g, 2.37 mmol) in a 1:1 mixture of MeOH:MeCN (100 mL) was added and the reaction mixture was stirred under reflux overnight. The solvent was evaporated, the residue was dissolved in a mixture of DCM and MeOH, deposited on celite and purified by flash column chromatography (Interchim Puriflash 30 µm DCM:MeOH 2% to 25%) to afford 0.83 g in a 69% yield.

$^1$H NMR (400 MHz, Methanol-$d_4$) rotamer mixture (ratio 1:0.65), δ (ppm): 8.28 (s, 1H), 8.22 (s, 1H), 5.99 (d, *J* = 4.7 Hz, 1H), 4.84 – 4.78 (m, 1H), 4.35 (dt, *J* = 8.1, 5.0 Hz, 1H), 4.23 – 4.12 (m, 2H), 3.85 (dt, *J* = 10.8, 7.3 Hz, 1H), 3.71 and 3.70 (s, 3H), 3.40 (tt, *J* = 9.2, 7.0 Hz, 1H), 3.14 (dd, *J* = 10.7, 8.7 Hz, 1H), 3.07 – 2.95 (m, 2H), 2.59 – 2.49 (m, 1H), 1.79 (ddd, *J* = 12.8, 9.5, 7.9 Hz, 1H), 1.41 and 1.37 (s, 9H). ).-$NH_2$ and 2x -OH is not present due to exchange with deuterium from $CD_3OD$.

$^{13}$C NMR (101 MHz, Methanol-$d_4$) rotamer mixture (ratio 1:0.65), only major rotamer peaks reported δ (ppm): 174.5, 157.3, 155.1, 153.9, 150.6, 141.6, 120.6, 90.4, 85.7, 81.8, 74.6, 73.9, 60.2, 53.9, 52.8, 42.4, 38.7, 34.9, 28.5.

ESI-MS, positive mode: m/z = 511.2 $[M+H]^+$.

HRMS (ESI) calculated for $C_{21}H_{31}N_6O_7S$ $[M+H]^+$ 511.1969, found 511.1977.

**1-(tert-butyl) 2-methyl (2R,4S)-4-((((2S,3S,4R,5R)-5-(6-amino-9H-purin-9-yl)-3,4-dihydroxytetrahydro-furan-2-yl)methyl)thio)pyrrolidine-1,2-dicarboxylate**

Ado-MeO-Boc-*trans-D*-Pro (**13**):

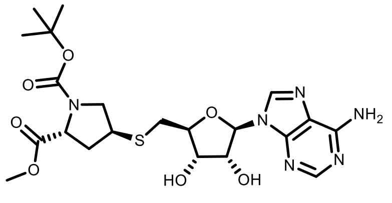

Ado-MeO-Boc-*trans-D*-Pro (**13**)

To a solution of **8** (0.85 g, 2.8 mmol) in dry MeOH (15 mL) under an argon atmosphere a solution of NaOMe (770 µL of 25% wt solution in MeOH, 2.8 mmol) was added and the resulting mixture was stirred for 30 min at room temperature. Then a solution of 5'-tosyladenosine (0.94 g, 2.24mmol) in a 1:1 mixture of MeOH:MeCN (100 mL) was added and the reaction mixture was stirred under reflux overnight. The solvent was evaporated, the residue was dissolved in a mixture of DCM and MeOH, deposited on celite and purified by flash column chromatography (Interchim Puriflash 30 µm $CHCl_3$:MeOH 2% to 30% gradient) to afford 0.8 g in a 56% yield.

$^1$H NMR (400 MHz, Methanol-$d_4$) rotamer mixture (ratio 1:0.7), δ (ppm): 8.28 (s, 1H), 8.22 (s, 1H), 5.98 (d, *J* = 4.9 Hz, 1H), 4.84 – 4.79 (m, 1H), 4.38 – 4.27 (m, 2H), 4.19 (dt, *J* = 6.7, 4.9 Hz, 1H), 3.71 (ddd, *J* = 13.5, 11.0, 6.8 Hz, 1H), 3.65 and 3.63 (s, 3H), 3.56 – 3.47 (m, 1H), 3.30 – 3.20 (m, 1H), 3.08 – 2.97 (m, 2H), 2.29 – 2.14 (m, 2H), 1.43 and 1.38 (s, 9H).-$NH_2$ and 2x -OH is not present due to exchange with deuterium from $CD_3OD$.

$^{13}$C NMR (101 MHz, Methanol-$d_4$) rotamer mixture (ratio 1:0.7), only major rotamer peaks reported δ (ppm): 174.6, 157.3, 155.4, 154.0, 150.6, 141.5, 120.7, 90.3, 86.0, 81.8, 74.6, 74.1, 60.0, 53.6, 52.7, 42.5, 38.3, 34.8, 28.5.

ESI-MS, positive mode: m/z = 511.2 $[M+H]^+$.

HRMS (ESI) calculated for $C_{21}H_{31}N_6O_7S$ $[M+H]^+$ 511.1969, found 511.1979.

**1-(tert-butyl) 2-methyl (2S,4R)-4-((((2S,3S,4R,5R)-5-(4-amino-7H-pyrrolo[2,3-d]pyrimidin-7-yl)-3,4-dihydroxytetrahydrofuran-2-yl)methyl)thio)pyrrolidine-1,2-dicarboxylate:**

dzAdo-MeO-Boc-cis-L-Pro (**14**)

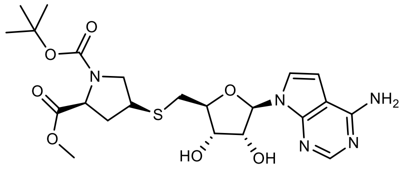


dzAdo-MeO-Boc-*cis*-*L*-Pro (**14**)

To a solution of **5** (640 mg, 2.1 mmol)in dry MeOH (15 mL) under an argon atmosphere a solution of NaOMe (576 µL of 25% wt solution in MeOH, 2.1 mmol) was added and the resulting mixture was stirred for 30 min at room temperature. Then a solution of 5'-chloro-7-deazaadenosine (300 mg, 1.06 mmol) in a 1:1 mixture of MeOH:MeCN (50 mL) was added and the reaction mixture was stirred under reflux overnight. The solvent was evaporated, the residue was dissolved in a mixture of DCM and MeOH, deposited on celite and purified by reverse phase flash column chromatography (C18 60g Biotage SNAP ULTRA 25 µm, MeOH:$H_2O$+0.2% formic acid 10% to 80% gradient) to afford 435 mg in a 81% yield.

$^1$H NMR (400 MHz, Methanol-$d_4$) rotamer mixture (ratio 1 : 0.65) δ 8.13 (s, 1H), 7.37 (d, *J* = 3.7 Hz, 1H), 6.71 (d, *J* = 3.7 Hz, 1H), 6.17 (d, *J* = 4.8 Hz, 1H), 4.55 – 4.47 (m, 1H), 4.28 – 4.12 (m, 3H), 3.80 (dd, *J* = 10.6, 7.3 Hz, 1H), 3.70 (s, 3H), 3.49 – 3.36 (m, 1H), 3.18 – 3.10 (m, 1H), 3.05 – 2.98 (m, 1H), 2.95 – 2.88 (m, 1H), 2.66 – 2.57 (m, 1H), 1.92 – 1.81 (m, 1H), 1.38 (s, 9H).

$^{13}C$ NMR (101 MHz, Methanol-$d_4$) rotamer mixture (ratio 1 : 0.65), only major rotamer peaks reported δ 174.6, 157.7, 155.1, 150.9, 150.8, 150.5, 123.9, 104.6, 101.9, 89.6, 85.0, 81.8, 75.4, 73.9, 60.2, 53.8, 52.7, 42.5, 38.8, 35.0, 28.5.

ESI-MS, positive mode: m/z = 510.2 $[M+H]^+$.

HRMS (ESI) calculated for $C_{22}H_{32}N_5O_7S$ $[M+H]^+$ 510.2017, found 510.2031.

**1-(tert-butyl) 2-methyl (2S,4S)-4-((((2S,3S,4R,5R)-5-(6-amino-9H-purin-9-yl)-3,4-dihydroxytetrahydrofuran-2-yl)methyl)selenyl)pyrrolidine-1,2-dicarboxylate**

SeAdo-MeO-Boc-*cis-L*-Pro (**15**):

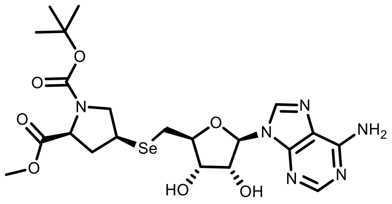


SeAdo-MeO-Boc-*cis-L*-Pro (**15**)

To a solution of **9** (520 mg, 1.56 mmol) in dry MeOH (15 mL) under an argon atmosphere $NaBH_4$ (120 mg, 3.12 mmol) was added in small portions over a period of 15 min. Once the addition was complete the resulting mixture was stirred at room temperature for additional 15 minutes and then a solution of 5'-tosyladenosine (550 mg, 1.3 mmol) in a 1:1 mixture of MeOH:MeCN (80 mL) was added and the reaction mixture was stirred under reflux for 4-6h. The solvent was evaporated and the residue was deposited on celite and purified by flash column chromatography (60g C18 Biotage SNAP ULTRA, $H_2O$ + 0.2% HCOOH : MeCN 10% to 80% gradient). The fractions containing the product were collected, solvents were evaporated to the minimum volume and then the sample was frozen and lyophilized to obtain 550 mg of a white powder in 63% yield.

$^1H$ NMR (400 MHz, Methanol-$d_4$) rotamer mixture (ratio 1:0.7) δ (ppm) 8.29 (s, 1H), 8.21 (s, 1H), 5.99 (dd, *J* = 4.8, 2.7 Hz, 1H), 4.83 – 4.79 (m, 1H), 4.31 (q, *J* = 4.9 Hz, 1H), 4.23 (dt, *J* = 6.7, 5.0 Hz, 1H), 4.16 (dt, *J* = 15.6, 7.7 Hz, 1H), 3.90 – 3.77 (m, 1H), 3.70 and 3.69 (s, 3H), 3.52 – 3.39 (m, 1H), 3.23 (dd, *J* = 10.8, 8.9 Hz, 1H), 3.12 – 3.01 (m, 2H), 2.70 – 2.61 (m, 1H), 1.97 – 1.87 (m, 1H), 1.42 and 1.38 (s, 9H).

$^{13}C$ NMR (101 MHz, Methanol-$d_4$) rotamer mixture (ratio 1:0.7), only major rotamer peaks reported δ (ppm) δ 174.5, 157.3, 155.0, 153.9, 150.6, 141.5, 120.6, 90.2, 86.0, 81.8, 74.8, 74.7, 60.5, 54.5, 52.7, 39.7, 34.3, 28.5, 27.1.

ESI-MS, positive mode: m/z = 559.2 $[M+H]^+$.

HRMS (ESI) calculated for $C_{21}H_{30}N_6O_7SeNa$ $[M+Na]^+$ 581.1235, found 581.1262.

**(2S,4S)-4-((((2S,3S,4R,5R)-5-(6-amino-9H-purin-9-yl)-3,4-dihydroxytetrahydrofuran-2-yl)methyl)thio)pyrrolidine-2-carboxylic acid**

Ado-*cis*-*L*-Pro (**16**):

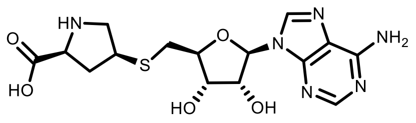
Ado-*cis*-*L*-Pro (**16**)

To a solution of **10** (600 mg, 1.17 mmol) in MeOH (15 mL) a solution of 1M KOH (5mL, 5 mmol) was added. The obtained mixture was stirred for 4-6h until completion of the reaction, and the reaction course was monitored by LC/MS analysis. Then solvents were evaporated to dryness and the residue was dissolved in MeCN (15 mL) and TFA was added (6 mL, 7.8 mmol). The mixture was stirred at room temperature overnight or until completion, and the reaction course was monitored by analytical LC/MS. Then solvents were evaporated to dryness, the residue was dissolved in water and purified by reverse phase flash column chromatography (60g C18 Biotage SNAP ULTRA, $H_2O$ + 0.2% HCOOH : MeOH 2% to 20% gradient). The fractions containing the product were collected, solvents were evaporated to the minimum volume and the sample was frozen and then lyophilized to obtain 300 mg of white powder in 65% yield.

$^1$H NMR (400 MHz, DMSO-$d_6$) δ 8.36 (s, 1H), 8.16 (s, 1H), 7.31 (s, 2H), 5.90 (d, *J* = 5.6 Hz, 1H), 4.73 (t, *J* = 5.4 Hz, 1H), 4.15 (t, *J* = 4.4 Hz, 1H), 4.03 (td, *J* = 6.3, 3.8 Hz, 1H), 3.80 (t, *J* = 8.2 Hz, 1H), 3.52 – 3.34 (m, 2H), 3.09 – 2.86 (m, 3H), 2.55 (t, *J* = 6.5 Hz, 1H), 1.78 (dt, *J* = 13.3, 7.9 Hz, 1H).2x-OH, proline-NH and -COOH protons are not visible due to fast exchange with formic acid protons.

$^{13}$C NMR (101 MHz, DMSO-$d_6$) δ 169.8, 156.1, 152.7, 149.4, 139.9, 119.1, 87.5, 83.8, 72.7, 72.6, 60.1, 50.5, 41.1, 35.9, 33.5.

ESI-MS, positive mode: m/z = 397.1 $[M+H]^+$.

HRMS (ESI) calculated for $C_{15}H_{21}N_6O_5S$ $[M+H]^+$ 397.1289, found 397.1298.

**(2S,4R)-4-((((2S,3S,4R,5R)-5-(6-amino-9H-purin-9-yl)-3,4-dihydroxytetrahydrofuran-2-yl)methyl)thio)pyrrolidine-2-carboxylic acid**

Ado-*trans*-*L*-Pro (**17**):

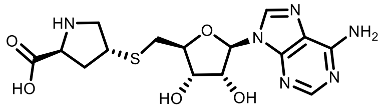
Ado-*trans*-*L*-Pro (**17**)

To a solution of **11** (300 mg, 0.59 mmol) in MeOH (10 mL) a solution of 1M KOH (2.5 mL, 2.5 mmol) was added. The obtained mixture was stirred for 4-6h until completion of the reaction, and the reaction course was monitored by LC/MS analysis. Then solvents were evaporated to dryness and the residue was dissolved in MeCN (10 mL) and TFA was added (3 mL, 3.9

mmol). The mixture was stirred at room temperature overnight or until completion, and the reaction course was monitored by LC/MS analysis. Then solvents were evaporated to dryness, the residue was dissolved in water and purified by reverse phase flash column chromatography (60g C18 Biotage SNAP ULTRA, $H_2O$ + 0.2% HCOOH : MeOH 2% to 20% gradient). The fractions containing the product were collected, solvents were evaporated to the minimum volume and the sample was frozen and then lyophilized to obtain 175 mg of white powder in 75% yield.

$^1$H NMR (400 MHz, DMSO-$d_6$) δ 8.35 (s, 1H), 8.15 (s, 1H), 7.32 (s, 2H), 5.90 (d, *J* = 5.7 Hz, 1H), 4.72 (t, *J* = 5.4 Hz, 1H), 4.14 (dd, *J* = 5.1, 3.9 Hz, 1H), 4.07 – 3.96 (m, 1H), 3.90 (dd, *J* = 8.6, 6.3 Hz, 1H), 3.56 (dd, *J* = 11.7, 6.8 Hz, 1H), 3.42 (p, *J* = 6.8 Hz, 1H), 3.06 – 2.89 (m, 3H), 2.28 (dt, *J* = 13.0, 6.5 Hz, 1H), 2.09 – 1.97 (m, 1H). 2x-OH, proline-NH and -COOH protons are not visible due to fast exchange with formic acid protons.

$^{13}$C NMR (101 MHz, DMSO-$d_6$) δ 169.9, 156.1, 152.7, 149.4, 139.8, 119.1, 87.5, 83.8, 72.8, 72.7, 59.9, 50.9, 41.2, 36.0, 33.8.

ESI-MS, positive mode: m/z = 397.1 $[M+H]^+$.

HRMS (ESI) calculated for $C_{15}H_{21}N_6O_5S$ $[M+H]^+$ 397.1289, found 397.1300.

**(2R,4R)-4-((((2S,3S,4R,5R)-5-(6-amino-9H-purin-9-yl)-3,4-dihydroxytetrahydrofuran-2-yl)methyl)thio)pyrrolidine-2-carboxylic acid**

Ado-*cis-D*-Pro (**18**):

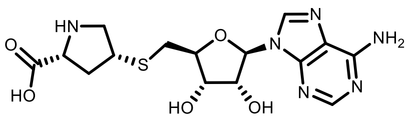


Ado-*cis-D*-Pro (**18**)

To a solution of **12** (200 mg, 0.39 mmol) in MeOH (10 mL) a solution of 1M KOH (2.5 mL, 2.5 mmol) was added. The obtained mixture was stirred for 4-6h until completion of the reaction, and the reaction course was monitored by LC/MS analysis. Then solvents were evaporated to dryness and the residue was dissolved in MeCN (10 mL) and TFA was added (3 mL, 3.9 mmol). The mixture was stirred at room temperature overnight or until completion, and the reaction course was monitored by LC/MS analysis. Then solvents were evaporated to dryness, the residue was dissolved in water and purified by reverse phase flash column chromatography (60g C18 Biotage SNAP ULTRA, $H_2O$ + 0.2% HCOOH : MeOH 2% to 20% gradient). The fractions containing the product were collected, solvents were evaporated to the minimum volume and the sample was frozen and then lyophilized to obtain 105 mg of white powder in 68% yield.

$^1$H NMR (400 MHz, DMSO-$d_6$) δ 8.36 (s, 1H), 8.16 (s, 1H), 7.31 (s, 2H), 5.89 (d, *J* = 5.5 Hz, 1H), 4.71 (t, *J* = 5.4 Hz, 1H), 4.15 (t, *J* = 4.6 Hz, 1H), 4.03 – 3.96 (m, 1H), 3.77 (t, *J* = 8.3 Hz, 1H), 3.49 – 3.39 (m, 2H),

3.05 – 2.95 (m, 2H), 2.90 (dd, *J* = 13.9, 7.1 Hz, 1H), 2.55 – 2.50 (m, 1H), 1.76 (dt, *J* = 13.5, 8.0 Hz, 1H). 2x-OH, proline-NH and -COOH protons are not visible due to fast exchange with formic acid protons.

$^{13}$C NMR (101 MHz, DMSO-$d_6$) δ 169.9, 156.1, 152.7, 149.4, 139.8, 119.1, 87.5, 83.8, 72.7, 72.5, 60.2, 50.7, 41.3, 35.9, 33.6.

ESI-MS, positive mode: m/z = 397.1 [M+H]$^+$.

HRMS (ESI) calculated for $C_{15}H_{21}N_6O_5S$ [M+H]$^+$ 397.1289, found 397.1305.

**(2R,4S)-4-((((2S,3S,4R,5R)-5-(6-amino-9H-purin-9-yl)-3,4-dihydroxytetrahydrofuran-2-yl)methyl)thio)pyrrolidine-2-carboxylic acid**

Ado-*trans-D*-Pro (**19**):

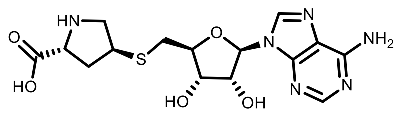

Ado-*trans-D*-Pro (**19**)

To a solution of **13** (200 mg, 0.39 mmol) in MeOH (10 mL) a solution of 1M KOH (2.5 mL, 2.5 mmol) was added. The obtained mixture was stirred for 4-6h until completion of the reaction, and the reaction course was monitored by LC/MS analysis. Then solvents were evaporated to dryness and the residue was dissolved in MeCN (10 mL) and TFA was added (3 mL, 3.9 mmol). The mixture was stirred at room temperature overnight or until completion, and the reaction course was monitored by LC/MS analysis. Then solvents were evaporated to dryness, the residue was dissolved in water and purified by reverse phase flash column chromatography (60g C18 Biotage SNAP ULTRA, $H_2O$ + 0.2% HCOOH : MeOH 2% to 20% gradient). The fractions containing the product were collected, solvents were evaporated to the minimum volume and the sample was frozen and then lyophilized to obtain 90 mg of white powder in 58% yield.

$^1$H NMR (400 MHz, DMSO-$d_6$) δ 8.35 (s, 1H), 8.16 (s, 1H), 7.30 (s, 2H), 5.89 (d, *J* = 5.7 Hz, 1H), 4.73 (t, *J* = 5.3 Hz, 1H), 4.15 (t, *J* = 4.5 Hz, 1H), 4.07 – 3.96 (m, 1H), 3.92 – 3.81 (m, 1H), 3.53 (dd, *J* = 11.6, 6.7 Hz, 1H), 3.40 (p, *J* = 6.7 Hz, 1H), 3.06 – 2.87 (m, 3H), 2.31 (dt, *J* = 13.1, 6.5 Hz, 1H), 2.05 – 1.92 (m, 1H). 2x-OH, proline-NH and -COOH protons are not visible due to fast exchange with formic acid protons.

$^{13}$C NMR (101 MHz, DMSO-$d_6$) δ 169.7, 156.1, 152.7, 149.4, 139.9, 119.2, 87.6, 83.6, 72.7, 72.6, 60.0, 50.7, 40.9, 36.1, 33.6.

ESI-MS, positive mode: m/z = 397.1 [M+H]$^+$.

HRMS (ESI) calculated for $C_{15}H_{21}N_6O_5S$ [M+H]$^+$ 397.1289, found 397.1299.

**(2S,4S)-4-((((2S,3S,4R,5R)-5-(4-amino-7H-pyrrolo[2,3-d]pyrimidin-7-yl)-3,4-dihydroxytetrahydrofuran-2-yl)methyl)thio)pyrrolidine-2-carboxylic acid:**

dzAdo-cis-L-Pro **20**

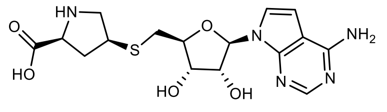

dzAdo-*cis*-L-Pro (**20**)

To a solution of **14** (200 mg, 0.39mmol) in MeOH (20 mL) a solution of 1M KOH (4 mL, 4 mmol) was added. The obtained mixture was stirred for 4-6h until completion of the reaction, and the reaction course was monitored by LC/MS analysis. Then solvents were evaporated to dryness and the residue was dissolved in MeCN (20 mL) and TFA was added (7.5 mL, 9.75 mmol). The mixture was stirred at room temperature overnight or until completion, and the reaction course was monitored by LC/MS analysis. Then solvents were evaporated to dryness, the residue was dissolved in water and purified by reverse phase flash column chromatography (60g C18 Biotage SNAP ULTRA, $H_2O$ + 0.2% HCOOH : MeOH 0% to 15% gradient). The fractions containing the product were collected, solvents were evaporated to the minimum volume and the sample was frozen and then lyophilized to obtain 115 mg of white powder in 75% yield.

$^1$H NMR (400 MHz, DMSO-$d_6$) δ 8.07 (s, 1H), 7.32 (d, *J* = 3.7 Hz, 1H), 7.13 (s, 2H), 6.63 (d, *J* = 3.6 Hz, 1H), 6.06 (d, *J* = 5.7 Hz, 1H), 4.43 (t, *J* = 5.5 Hz, 1H), 4.05 (t, *J* = 4.3 Hz, 1H), 3.96 (q, *J* = 5.9, 5.4 Hz, 1H), 3.89 (t, *J* = 8.2 Hz, 1H), 3.53 – 3.38 (m, 2H), 3.04 (dd, *J* = 11.2, 6.8 Hz, 1H), 2.98 – 2.80 (m, 2H), 2.61 – 2.51 (m, 1H), 1.80 (dt, *J* = 13.2, 8.0 Hz, 1H).

$^{13}$C NMR (101 MHz, DMSO-$d_6$) δ 170.01, 157.29, 151.54, 150.31, 121.73, 102.79, 100.28, 87.02, 82.80, 73.23, 72.56, 59.92, 50.54, 41.06, 35.80, 33.74.

ESI-MS, positive mode: m/z = 396.1 $[M+H]^+$.

HRMS (ESI) calculated for $C_{16}H_{22}N_5O_5S$ $[M+H]^+$ 396.1336, found 396.1355.

**(2S,4S)-4-((((2S,3S,4R,5R)-5-(6-amino-9H-purin-9-yl)-3,4-dihydroxytetrahydrofuran-2-yl)methyl)selenyl)pyrrolidine-2-carboxylic acid:**

SeAdo-*cis*-*L*-Pro (**21**):

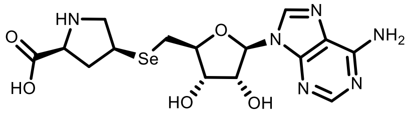

SeAdo-*cis*-*L*-Pro (**21**)

To a solution of **15** (400 mg, 0.72 mmol) in MeOH (20 mL) a solution of 1M KOH (4 mL, 4 mmol) was added. The obtained mixture was stirred for 4-6h until completion of the reaction, and the reaction course was monitored by LC/MS analysis. Then solvents were evaporated to dryness and the residue was dissolved in MeCN (20 mL) and TFA was added

(7.5 mL, 9.75 mmol). The mixture was stirred at room temperature overnight or until completion, and the reaction course was monitored by LC/MS analysis. Then solvents were evaporated to dryness, the residue was dissolved in water and purified by reverse phase flash column chromatography (60g C18 Biotage SNAP ULTRA, $H_2O$ + 0.2% HCOOH : MeOH 2% to 20% gradient). The fractions containing the product were collected, solvents were evaporated to the minimum volume and the sample was frozen and then lyophilized to obtain 207 mg of white powder in 65% yield.

$^{1}$H NMR (400 MHz, DMSO-$d_6$) δ 8.35 (s, 1H), 8.15 (s, 1H), 7.29 (s, 2H), 5.89 (d, J = 5.6 Hz, 1H), 4.72 (t, J = 5.4 Hz, 1H), 4.14 (t, J = 4.5 Hz, 1H), 4.10 – 4.02 (m, 1H), 3.67 (t, J = 8.2 Hz, 1H), 3.52 – 3.37 (m, 2H), 3.08 – 2.91 (m, 3H), 2.59 – 2.51 (m, 1H), 1.88 – 1.73 (m, 1H). 2x-OH, proline-NH and -COOH protons are not visible due to fast exchange with formic acid protons.

$^{13}$C NMR (101 MHz, DMSO-$d_6$) δ 156.1, 152.7, 149.4, 139.8, 119.1, 87.5, 84.1, 73.1, 72.9, 60.5, 51.4, 48.6, 36.9, 33.4, 26.0.

ESI-MS, positive mode: m/z = 445.1 $[M+H]^+$.

HRMS (ESI) calculated for $C_{15}H_{21}N_6O_5Se$ $[M+H]^+$ 445.0734, found 445.0743.

Cofactors **22a** / **22b**:

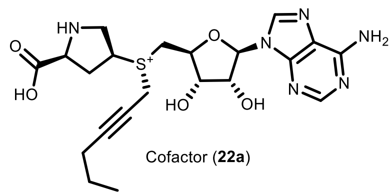

Cofactor (**22a**)

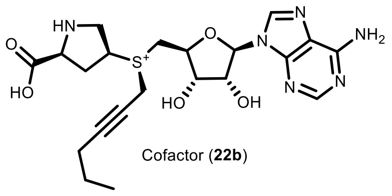

Cofactor (**22b**)

Ado-cis-L-Pro (**16**) (20 mg, 0.051 mmol) was dissolved in (S)-2-chloropropanoic acid (200 µL) and hex-2-yn-1-yl methanesulfonate (**SI-1**) (90 µL, 0.51 mmol) was added to the solution and the obtained mixture was stirred at 40°C for 24h and the course of the reaction was monitored by LC/MS. Then the resulting mixture was poured into an ammonium formate buffer (25 mL, 10mM, pH = 3.5) and washed with $Et_2O$ (3x40mL). The aqueous layer was taken and concentrated to 1-3 mL and the target compound was purified by preparative HPLC (preparative column: Pursuit 10 C18 10 µm, 250×50.0 mm, Agilent, flow rate: 150 mL/min, solvent A: MeOH, solvent B: $H_2O$ (ammonium formate buffer 10 mM, pH = 3.5); temperature 25 °C, gradient A:B - 2 min 2:98 isocratic, 2-20 min 2:98 to 20:80 gradient and 20-22min 20:80 isocratic). Collected fractions of pure stereoisomers were concentrated and the concentrations of the obtained solutions were determined by their UV absorption ($\varepsilon_{260}$ = 15400 $L \cdot mol^{-1} \cdot cm^{-1}$). Obtained 1.2 mL, 9.2 mM of peak-1 (**22a**) in 22% yield, and 1.5 mL 8.5 mM of peak-2 (**22b**) in 25% yield. The peak-2 with longer retention time was determined to be the enzymatically active *R*-stereoisomer (**22b**) and the stereocenter configuration at the sulfur atom was assigned accordingly to active AdoMet stereoisomer. The following analytical data were collected for cofactor **22b**:

$^1$H NMR (400 MHz, deuterium oxide) δ 8.24 (s, 1H), 8.23 (s, 1H), 6.08 (d, *J* = 2.5 Hz, 1H), 4.94 (dd, *J* = 5.1, 2.5 Hz, 1H), 4.69 (dd, *J* = 6.9, 5.1 Hz, 1H), 4.56 (ddd, *J* = 10.6, 6.9, 2.0 Hz, 1H), 4.47 – 4.19 (m, 4H), 3.98 – 3.80 (m, 3H), 3.72 (dd, *J* = 13.6, 10.6 Hz, 1H), 2.89 (dt, *J* = 14.6, 7.9 Hz, 1H), 2.46 (dt, *J* = 14.0, 6.7 Hz, 1H), 2.05 – 1.92 (m, 2H), 1.28 (h, *J* = 6.9 Hz, 2H), 0.77 (t, *J* = 7.4 Hz, 3H).

ESI-MS, positive mode: m/z = 477.2 $[M]^+$. HRMS (ESI) calculated for $C_{21}H_{29}N_6O_5S$ $[M]^+$ 477.1915, found 477.1925.

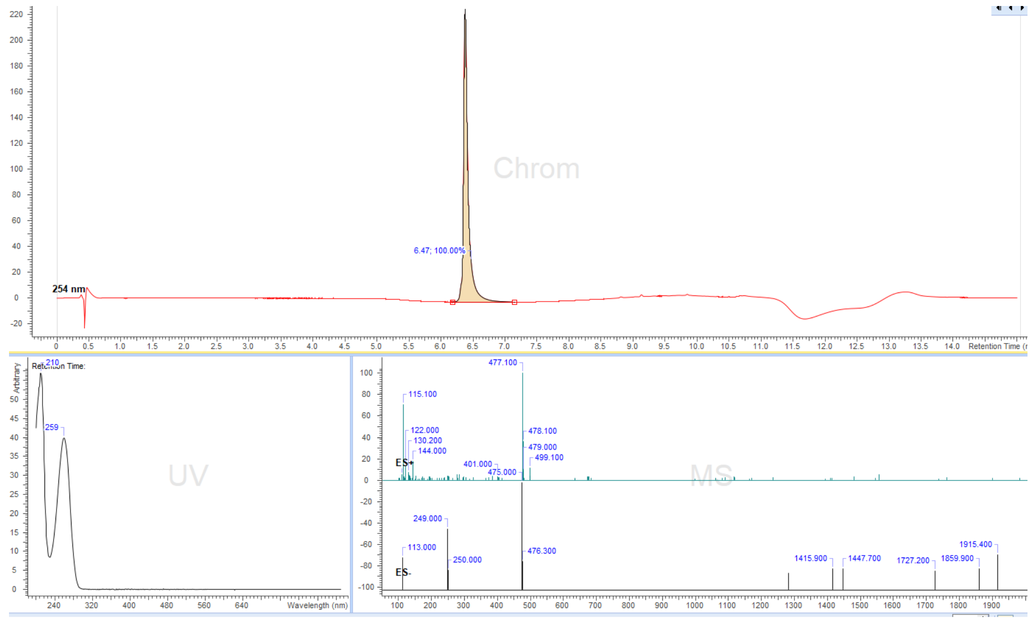


**Figure S24.** LC/MS analysis profile of compound **22b.**

Cofactors **23a** / **23b**:

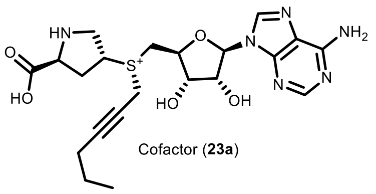

Cofactor (**23a**)

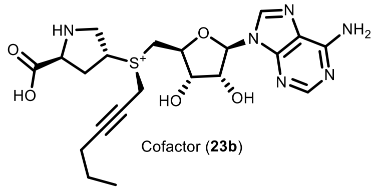

Cofactor (**23b**)

Ado-trans-L-Pro (**17**) (20 mg, 0.051 mmol) was dissolved in (S)-2-chloropropanoic acid (200 µL) and Hex-2-yn-1-yl methanesulfonate (**SI-1**) (90 µL, 0.51 mmol) was added to the solution and the obtained mixture was stirred at 40°C for 24h and the course of the reaction was monitored by LC/MS. Then the resulting mixture was poured into an ammonium formate buffer (25 mL, 10mM, pH = 3.5) and washed with $Et_2O$ (3x40mL). The aqueous layer was taken and concentrated to 1-3 mL and the target compound was purified by preparative HPLC (preparative column: Pursuit 10 C18 10 µm, 250×50.0 mm, Agilent, flow rate: 150 mL/min, solvent A: MeOH, solvent B: $H_2O$ (ammonium formate buffer 10 mM, pH = 3.5); temperature 25 °C, gradient A:B - 2 min 2:98 isocratic, 2-20 min 2:98 to 20:80 gradient and 20-22min 20:80 isocratic). Collected fractions of pure stereoisomers were concentrated and the concentrations of the obtained solutions were determined by their UV absorption ($\varepsilon_{260}$ = 15400 $L{\cdot}mol^{-1}{\cdot}cm^{-1}$). 1.1 mL, 9.2 mM of peak-1 (**23a**) in 20%, and 1.0 mL 9.0 mM of peak-2 (**23b**) in 18% were obtained. The stereocenter configuration at the sulfur atom was assigned accordingly to the example of the cofactor **22b**. The following analytical data were collected for cofactor **23b**:

$^1$H NMR (400 MHz, deuterium oxide) δ 8.26 (s, 1H), 8.24 (s, 1H), 6.09 (d, *J* = 2.6 Hz, 1H), 4.96 (dd, *J* = 5.1, 2.6 Hz, 1H), 4.72 – 4.71 (m, 1H, overlapped with $H_2O$), 4.59 (ddd, *J* = 10.3, 6.7, 1.9 Hz, 1H), 4.55 – 4.08 (m, 4H), 4.01 – 3.92 (m, 2H), 3.80 – 3.67 (m, 2H), 2.86 – 2.67 (m, 2H), 2.06 – 1.90 (m, 2H), 1.32 – 1.25 (m, 2H), 0.77 (t, *J* = 7.4 Hz, 3H).

ESI-MS, positive mode: m/z = 477.2 $[M]^+$. HRMS (ESI) calculated for $C_{21}H_{29}N_6O_5S$ $[M]^+$ 477.1915, found 477.1918.

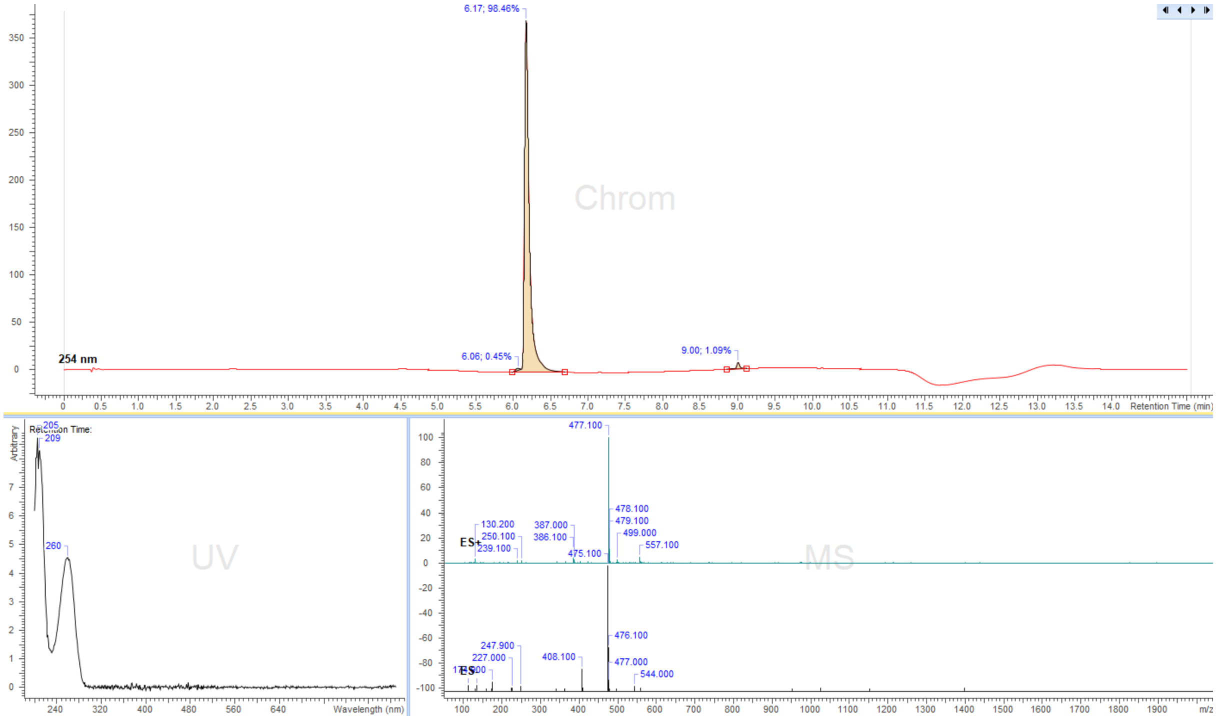


**Figure S25.** LC/MS analysis profile of compound **23b.**

Cofactors **24a** / **24b**:

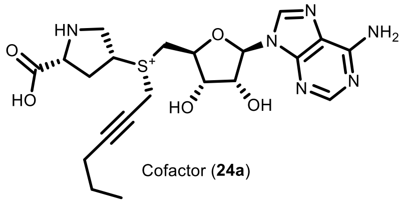

Cofactor (**24a**)

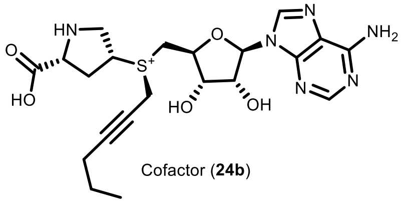

Cofactor (**24b**)

Ado-cis-D-Pro (**18**) (20 mg, 0.051 mmol) was dissolved in (S)-2-chloropropanoic acid (200 µL) and hex-2-yn-1-yl methanesulfonate (**SI-1**) (90 µL, 0.51 mmol) was added to the solution and the obtained mixture was stirred at 40°C for 24h and the course of the reaction was monitored by LC/MS. Then the resulting mixture was poured into an ammonium formate buffer (25 mL, 10mM, pH = 3.5) and washed with $Et_2O$ (3x40mL). The aqueous layer was taken and concentrated to 1-3 mL and the target compound was purified by preparative HPLC (preparative column: Pursuit 10 C18 10 µm, 250×50.0 mm, Agilent, flow rate: 150 mL/min, solvent A: MeOH, solvent B: $H_2O$ (ammonium formate buffer 10 mM, pH = 3.5); temperature 25 °C, gradient A:B - 2 min 2:98 isocratic, 2-20 min 2:98 to 20:80 gradient and 20-22min 20:80 isocratic). Collected fractions of pure stereoisomers were concentrated and the concentrations of the obtained solutions were determined by their UV absorption ($\varepsilon_{260}$ = 15400 L·mol$^{-1}$·cm$^{-1}$). 1.2 mL, 9.7 mM of peak-1 (**24a**) in 23%, and 1.0 mL 8.2 mM of peak-2 (**24b**) in 16% were obtained. The peak-2 with longer retention time was determined to be the enzymatically active *R*-stereoisomer (**24b**) and the stereocenter configuration at the sulfur atom was assigned accordingly to active AdoMet stereoisomer, following analytical data were collected for **24b**:

$^1$H NMR (400 MHz, deuterium oxide) δ 8.25 (s, 1H), 8.23 (s, 1H), 6.08 (d, *J* = 2.6 Hz, 1H), 4.95 (dd, *J* = 5.2, 2.6 Hz, 1H), 4.70 – 4.68 (m, 1H, overlapped with $H_2O$), 4.56 (t, *J* = 8.2 Hz, 1H), 4.46 – 4.22 (m, 4H), 4.01 – 3.64 (m, 4H), 2.93 (dt, *J* = 14.8, 7.5 Hz, 1H), 2.57 (dt, *J* = 13.9, 6.9 Hz, 1H), 2.05 – 1.90 (m, 2H), 1.27 (h, *J* = 7.2 Hz, 2H), 0.77 (t, *J* = 7.4 Hz, 3H).

ESI-MS, positive mode: m/z = 477.2 $[M]^+$. HRMS (ESI) calculated for $C_{21}H_{29}N_6O_5S$ $[M]^+$ 477.1915, found 477.1927.

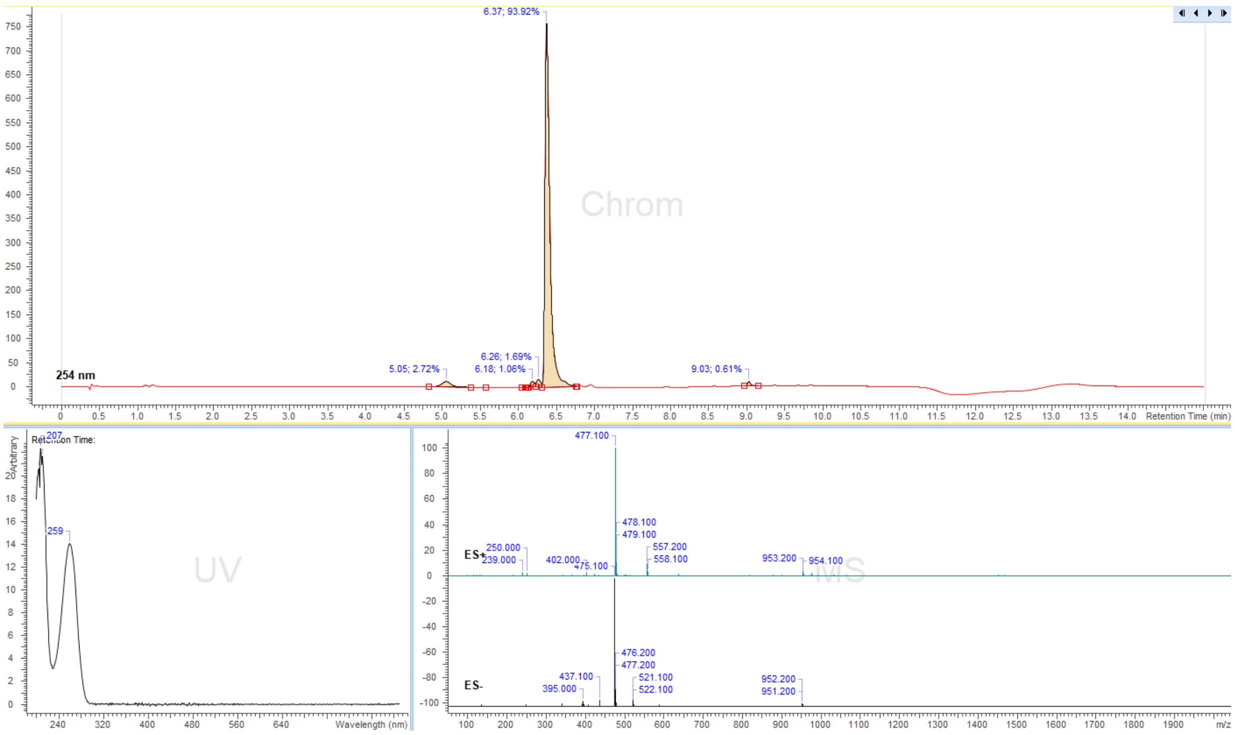


**Figure S26**. LC/MS analysis profile of compound **24b.**

Cofactors **25a** / **25b**:

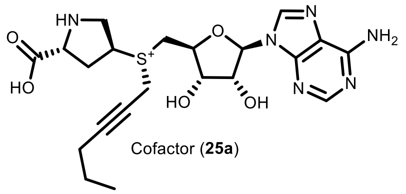

Cofactor (**25a**)

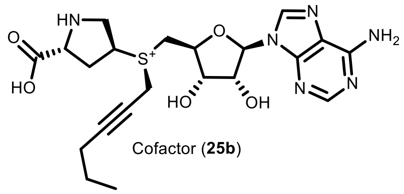

Cofactor (**25b**)

Ado-trans-D-Pro (**19**) (20 mg, 0.051 mmol) was dissolved in (S)-2-chloropropanoic acid (200 µL) and hex-2-yn-1-yl methanesulfonate (**SI-1**) (90 µL, 0.51 mmol) was added to the solution and the obtained mixture was stirred at 40°C for 24h and the course of the reaction was monitored by LC/MS. Then the resulting mixture was poured into an ammonium formate buffer (25 mL, 10mM, pH = 3.5) and washed with $Et_2O$ (3x40mL). The aqueous layer was taken and concentrated to 1-3 mL and the target compound was purified by preparative HPLC (preparative column: Pursuit 10 C18 10 µm, 250×50.0 mm, Agilent, flow rate: 150 mL/min, solvent A: MeOH, solvent B: $H_2O$ (ammonium formate buffer 10 mM, pH = 3.5); temperature 25 °C, gradient A:B - 2 min 2:98 isocratic, 2-20 min 2:98 to 20:80 gradient and 20-22min 20:80 isocratic). Collected fractions of pure stereoisomers were concentrated and the concentrations of the obtained solutions were determined by their UV absorption ($\varepsilon_{260}$ = 15400 $L{\cdot}mol^{-1}{\cdot}cm^{-1}$). 1.2 mL, 7.1 mM of peak-1 (**25a**) in 17%, and 1.35 mL 7.9 mM of peak-2 (**25b**) in 21% were obtained. The stereocenter configuration at the sulfur atom was assigned accordingly to the cofactor **22b** example and the following analytical data were collected for **25b**:

$^1$H NMR (400 MHz, deuterium oxide) δ 8.24 (s, 1H), 8.23 (s, 1H), 6.08 (d, *J* = 2.6 Hz, 1H), 4.95 (dd, *J* = 5.1, 2.6 Hz, 1H), 4.71 – 4.69 (m, 1H, overlapped with $H_2O$), 4.56 (ddd, *J* = 10.7, 6.8, 2.2 Hz, 1H), 4.43 – 4.39 (m, 2H), 4.26 (dd, *J* = 8.4, 5.8 Hz, 1H), 4.17 (p, *J* = 7.3 Hz, 1H), 3.98 – 3.87 (m, 2H), 3.77 – 3.66 (m, 2H), 2.70 – 2.58 (m, 2H), 2.04 – 1.92 (m, 2H), 1.28 (h, *J* = 7.4 Hz, 2H), 0.77 (t, *J* = 7.4 Hz, 3H).

ESI-MS, positive mode: m/z = 477.2 $[M]^+$. HRMS (ESI) calculated for $C_{21}H_{29}N_6O_5S$ $[M]^+$ 477.1915, found 477.1923.

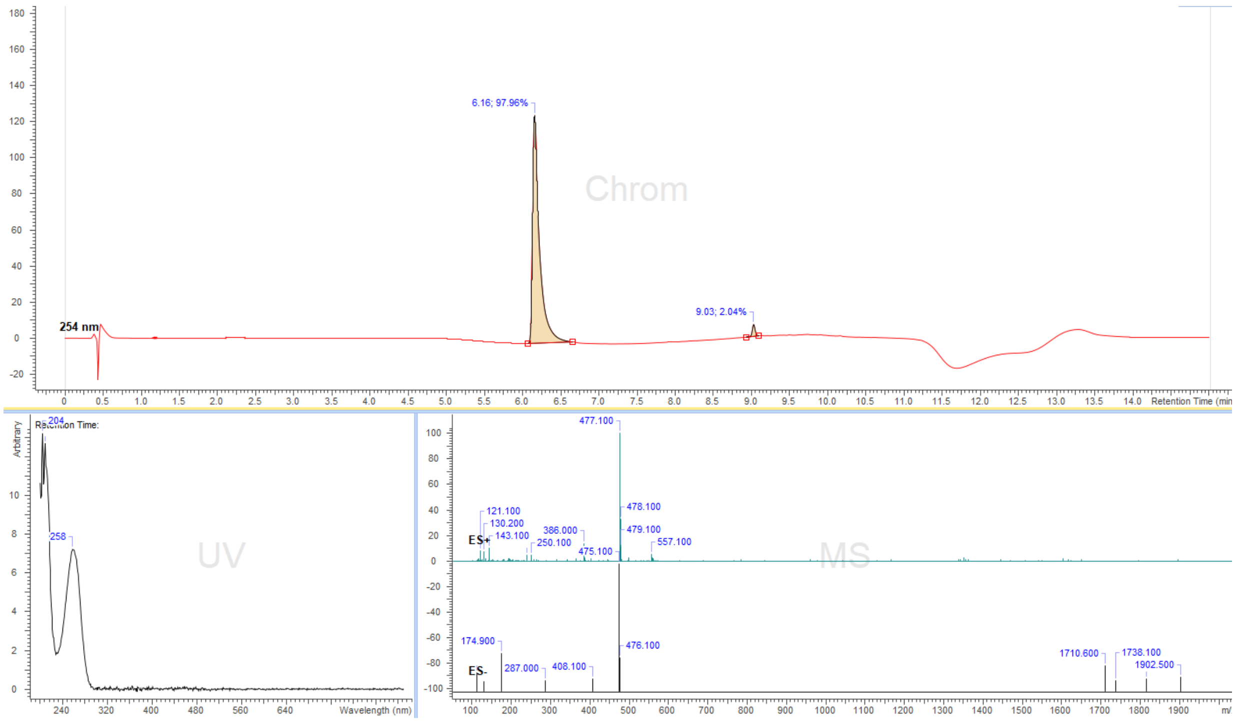


**Figure S27.** LC/MS analysis profile of compound **25b.**

**Cofactors 26a / 26b**

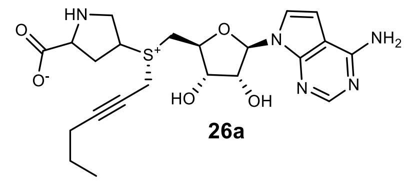


**26a**

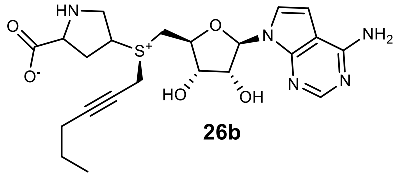


**26b**

dzAdo-cis-L-Pro (**20**) (10 mg, 0.025 mmol) was dissolved in (S)-2-chloropropanoic acid (200 µL) and hex-2-yn-1-yl methanesulfonate (**SI-1**) (90 µL, 0.51 mmol) was added to the solution and the obtained mixture was stirred at 40°C for 24h and the course of the reaction was monitored by LC/MS. Then the resulting mixture was poured into an ammonium formate buffer (25 mL, 10mM, pH = 3.5) and washed with $Et_2O$ (3x40mL). The aqueous layer was taken and concentrated to 1-3 mL and the target compound was purified by preparative HPLC (preparative column: Pursuit 10 C18 10 µm, 250×50.0 mm, Agilent, flow rate: 150 mL/min, solvent A: MeOH, solvent B: $H_2O$ (ammonium formate buffer 10 mM, pH = 3.5); temperature 25 °C, gradient A:B - 2 min 2:98 isocratic, 2-20 min 2:98 to 20:80 gradient and 20-22min 20:80 isocratic). Collected fractions of pure stereoisomers were concentrated and the concentrations of the obtained solutions were determined by their UV absorption ($\varepsilon_{260}$ = 15400 $L \cdot mol^{-1} \cdot cm^{-1}$). Obtained 0.8 mL, 9.2 mM of peak-1 **26a** in 29% yield, and 0.9 mL 8.5 mM of peak-2 **26b** in 30% yield. The peak-2 with longer retention time was determined to be the enzymatically active *R*-stereoisomer (**26b**) and the stereocenter configuration at the sulfur atom was assigned accordingly to active AdoMet stereoisomer. The following analytical data were collected for cofactor **26b**:

$^1H$ NMR (400 MHz, Deuterium Oxide) δ 8.19 (s, 1H), 7.34 (d, *J* = 3.8 Hz, 1H), 6.75 (d, *J* = 3.7 Hz, 1H), 6.19 (d, *J* = 2.9 Hz, 1H), 4.82 – 4.81 (m, 1H), 4.71 – 4.19 (m, 6H), 3.95 – 3.66 (m, 4H), 2.90 (dt, *J* = 14.6, 7.8 Hz, 1H), 2.45 (dt, *J* = 14.1, 6.9 Hz, 1H), 2.14 – 2.07 (m, 2H), 1.37 (p, *J* = 7.2 Hz, 2H), 0.85 (t, *J* = 7.4 Hz, 3H).

ESI-MS, positive mode: m/z = 476.2 $[M]^+$.

HRMS (ESI) calculated for $C_{22}H_{31}N_5O_5S$ $[M]^+$ 476.1962, found 476.1964.

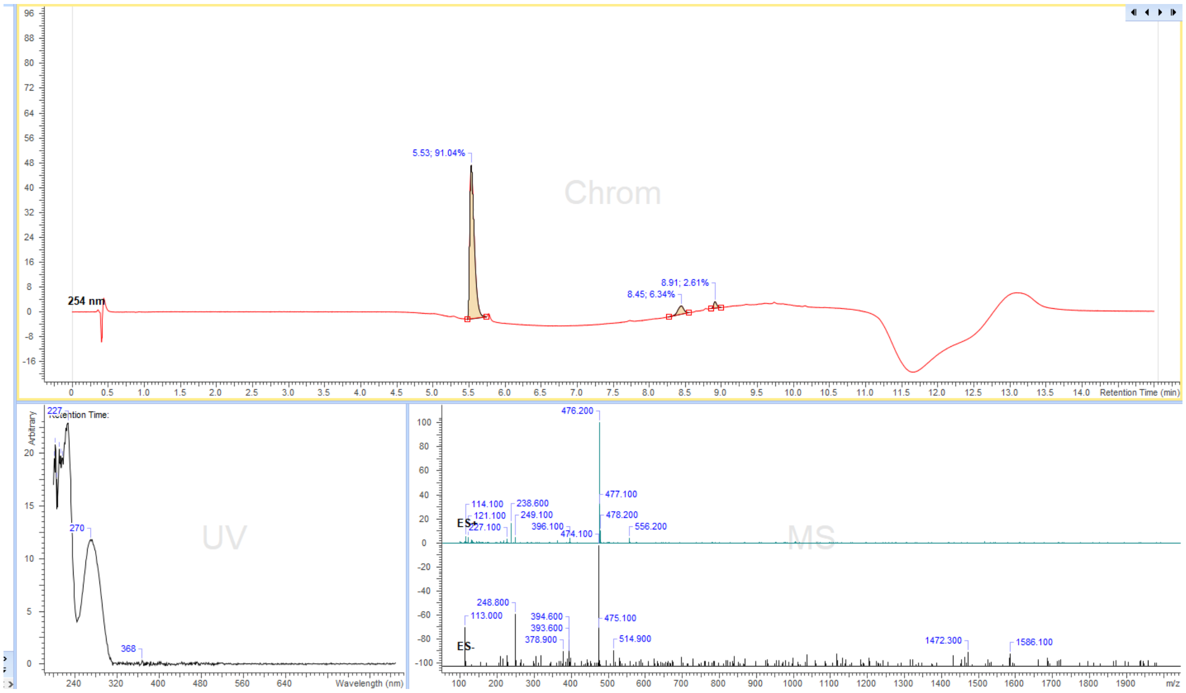


**Figure S28.** LC/MS analysis profile of compound **26b.**

Cofactors **27a** / **27b**:

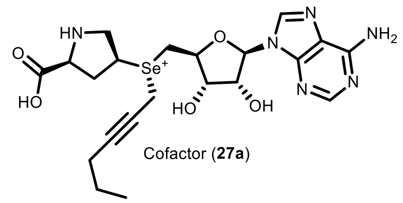

Cofactor (**27a**)

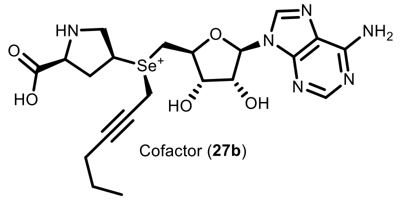

Cofactor (**27b**)

SeAdo-cis-L-Pro (**21**) (20 mg, 0.045 mmol) was dissolved in (S)-2-chloropropanoic acid (200 µL) and hex-2-yn-1-yl methanesulfonate (**SI-1**) (80 µL, 0.45 mmol) was added to the solution and the obtained mixture was stirred at room temperature for 24h-36h and the course of the reaction was monitored by LC/MS. Then the resulting mixture was poured into an ammonium formate buffer (25 mL, 10mM, pH = 3.5) and washed with $Et_2O$ (3x40mL). The aqueous layer was taken and concentrated to 1-3 mL and the target compound was purified by preparative HPLC (preparative column: Pursuit 10 C18 10 µm, 250×50.0 mm, Agilent, flow rate: 150 mL/min, solvent A: MeOH, solvent B: $H_2O$ (ammonium formate buffer 10 mM, pH = 3.5); temperature 25 °C, gradient A:B - 2 min 2:98 isocratic, 2-20 min 2:98 to 20:80 gradient and 20-22min 20:80 isocratic). Collected fractions of pure stereoisomers were concentrated and the concentrations of the obtained solutions were determined by their UV absorption ($\varepsilon_{260}$ = 15400 $L \cdot mol^{-1} \cdot cm^{-1}$). 1.0 mL, 6.7 mM of peak-1 (**27a**) in 15% yield, and 0.8 mL 10.2 mM of peak-2 (**27b**) in 18% yield were obtained. The peak-2 (**27b**) with longer retention time was determined to be the enzymatically active stereoisomer and the stereocenter configuration at the selenium atom was assigned accordingly to the example of the active AdoMet stereoisomer. The following analytical data were collected for cofactor **27b**:

$^1$H NMR (400 MHz, deuterium oxide) δ 8.26 (s, 1H), 8.23 (s, 1H), 6.07 (d, *J* = 2.6 Hz, 1H), 4.96 (dd, *J* = 5.0, 2.7 Hz, 1H), 4.66 – 4.62 (m, 1H), 4.55 (ddd, *J* = 10.7, 6.8, 2.5 Hz, 1H), 4.25 (dd, *J* = 8.0, 6.5 Hz, 2H), 4.17 (t, *J* = 2.3 Hz, 2H), 3.97 – 3.82 (m, 3H), 3.68 (dd, *J* = 12.3, 10.7 Hz, 1H), 2.90 (dt, *J* = 14.8, 7.9 Hz, 1H), 2.51 (dt, *J* = 14.8, 6.4 Hz, 1H), 2.04 (tdd, *J* = 7.8, 7.0, 6.3, 2.4 Hz, 2H), 1.33 (h, *J* = 7.3 Hz, 2H), 0.81 (t, *J* = 7.4 Hz, 3H).

ESI-MS, positive mode: m/z = 525.2 $[M]^+$. HRMS (ESI) calculated for $C_{21}H_{29}N_6O_5Se$ $[M]^+$ 525.1360, found 525.1368.

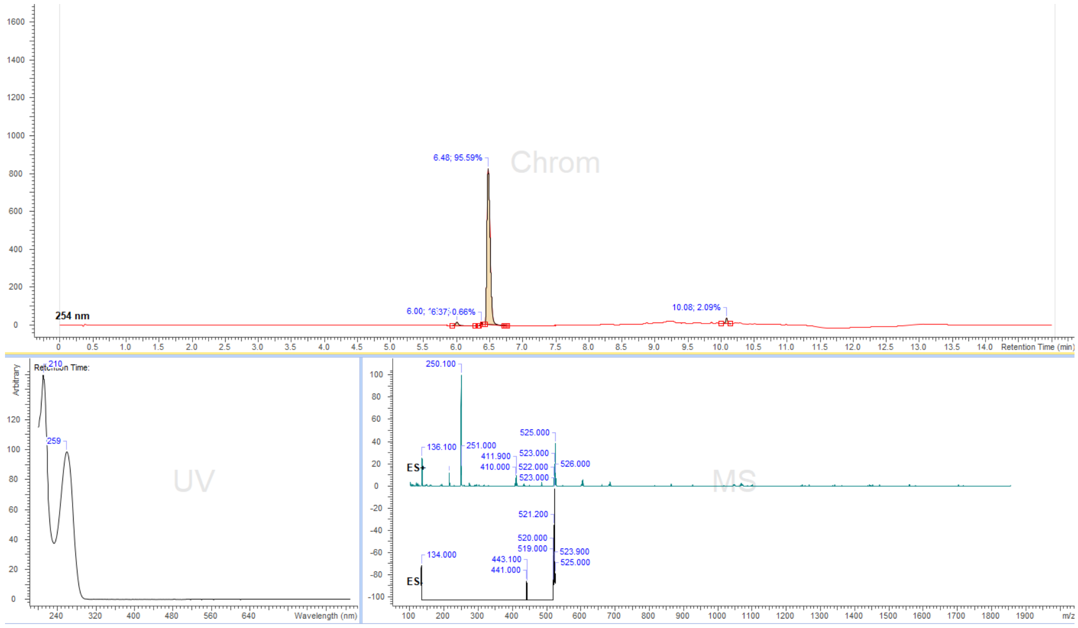


**Figure S29.** LC/MS analysis profile of compound **27b.**

Cofactor **28ab**:

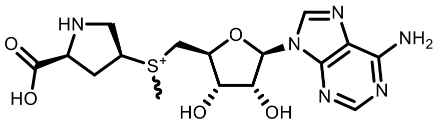

Cofactor (**28ab**)

Ado-cis-L-Pro (**16**) (20 mg, 0.051 mmol) was dissolved in (S)-2-chloropropanoic acid (200 µL) and methyl trifluoromethanesulfonate (30 µL, 0.25 mmol) was added to the solution, the obtained mixture was stirred at room temperature for 3h and the course of the reaction was monitored by LC/MS. Then the resulting mixture was poured into an ammonium formate buffer (25 mL, 10mM, pH = 3.5) and washed with $Et_2O$ (3x40mL). The aqueous layer was taken and concentrated to 1-3 mL and the target compound was purified by preparative HPLC (preparative column: Pursuit 10 C18 10 µm, 250×50.0 mm, Agilent, flow rate: 150 mL/min, solvent A: MeOH, solvent B: $H_2O$ (ammonium formate buffer 10 mM, pH = 3.5); temperature 25 °C, gradient A:B - 2 min 2:98 isocratic, 2-20 min 2:98 to 20:80 gradient and 20-22min 20:80 isocratic). The collected fraction of the mixture of both stereoisomers was concentrated and the concentration of the obtained solution was determined by its UV absorption ($\varepsilon_{260}$ = 15400 $L\cdot mol^{-1}\cdot cm^{-1}$). 1.5 mL, 13.9 mM of a mixture of both stereoisomers in 41% yield were obtained.

$^1$H NMR (400 MHz, deuterium oxide) δ 8.27 – 8.20 (m, 2H), 6.07 (dd, *J* = 4.4, 3.3 Hz, 1H), 4.92 (ddd, *J* = 10.2, 5.0, 4.3 Hz, 1H), 4.72 – 4.30 (m, 4H), 4.28 – 4.23 (m, 1H), 4.16 – 4.04 (m, 1H), 3.97 (dd, *J* = 14.1, 2.1 Hz, 1H), 3.91 – 3.77 (m, 1H), 3.73 (d, *J* = 6.9 Hz, 1H), 3.02 (d, *J* = 3.5 Hz, 3H), 2.96 – 2.87 (m, 1H), 2.46 (ddd, *J* = 14.2, 7.5, 6.6 Hz, 1H).

ESI-MS, positive mode: m/z = 411.2 $[M]^+$. HRMS (ESI) calculated for $C_{16}H_{23}N_6O_5S$ $[M]^+$ 411.1445, found 411.1448.

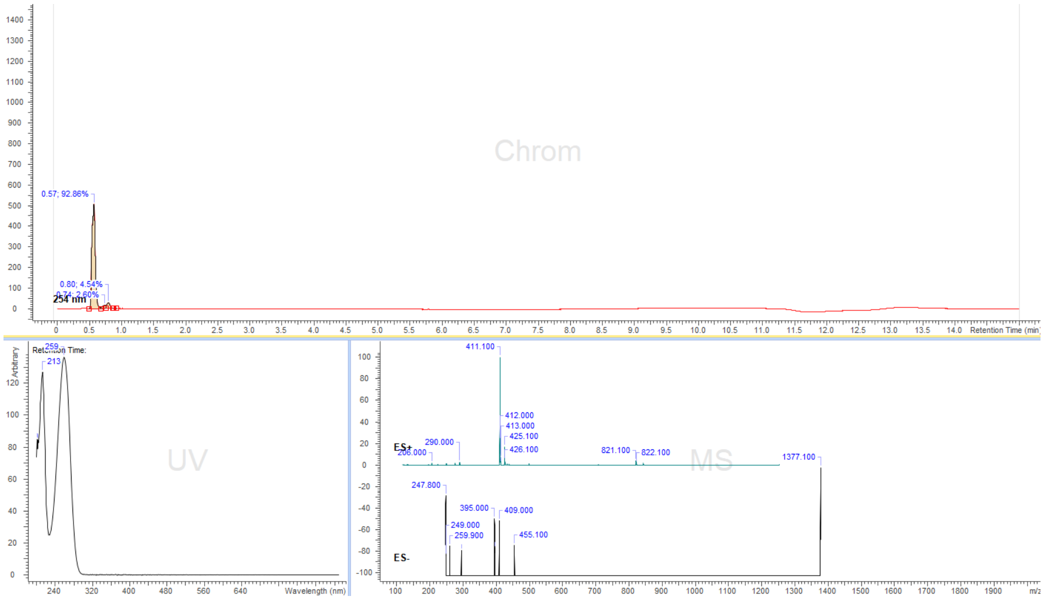


**Figure S30.** LC/MS analysis profile of compound **28ab.**

Cofactor **29ab**

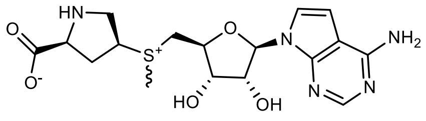


Cofactor **29ab**
dzAdo-cis-L-ProMet

dzAdo-cis-L-Pro (**20**) (10 mg, 0.025 mmol) was dissolved in (S)-2-chloropropanoic acid (200 µL) and methyl trifluoromethanesulfonate (30 µL, 0.25 mmol) was added to the solution, the obtained mixture was stirred at room temperature for 3h and the course of the reaction was monitored by LC/MS. Then the resulting mixture was poured into an ammonium formate buffer (25 mL, 10mM, pH = 3.5) and washed with $Et_2O$ (3x40mL). The aqueous layer was taken and concentrated to 1-3 mL and the target compound was purified by preparative HPLC (preparative column: Pursuit 10 C18 10 µm, 250×50.0 mm, Agilent, flow rate: 150 mL/min, solvent A: MeOH, solvent B: $H_2O$ (ammonium formate buffer 10 mM, pH = 3.5); temperature 25 °C, gradient A:B - 2 min 2:98 isocratic, 2-20 min 2:98 to 20:80 gradient and 20-22min 20:80 isocratic). The collected fraction of the mixture of both stereoisomers was concentrated and the concentration of the obtained solution was determined by its UV absorption ($\varepsilon_{260}$ = 15400 $L \cdot mol^{-1} \cdot cm^{-1}$). 1.2 mL, 7 mM of a mixture of both stereoisomers in 34% yield were obtained.

$^1$H NMR (400 MHz, Deuterium Oxide) δ 8.23 (d, *J* = 2.0 Hz, 1H), 7.41 (d, *J* = 3.9 Hz, 1H), 6.8 1 (dd, *J* = 3.8, 1.5 Hz, 1H), 6.21 (dd, *J* = 6.7, 4.6 Hz, 1H), 4.90 – 4.80 (m, 1H), 4.60 – 4.44 (m, 3H), 4.36 – 4.20 (m, 1H), 4.07 – 3.77 (m, 3H), 3.71 (d, *J* = 6.9 Hz, 1H), 3.05 (dd, *J* = 8.8, 1.9 Hz, 3H), 2.99 – 2.68 (m, 1H), 2.69 – 2.36 (m, 1H).

ESI-MS, positive mode: m/z = 410.2 $[M]^+$.

HRMS (ESI) calculated for $C_{16}H_{22}N_5O_5S$ $[M+H]^+$ 410.1490, found 410.1493.

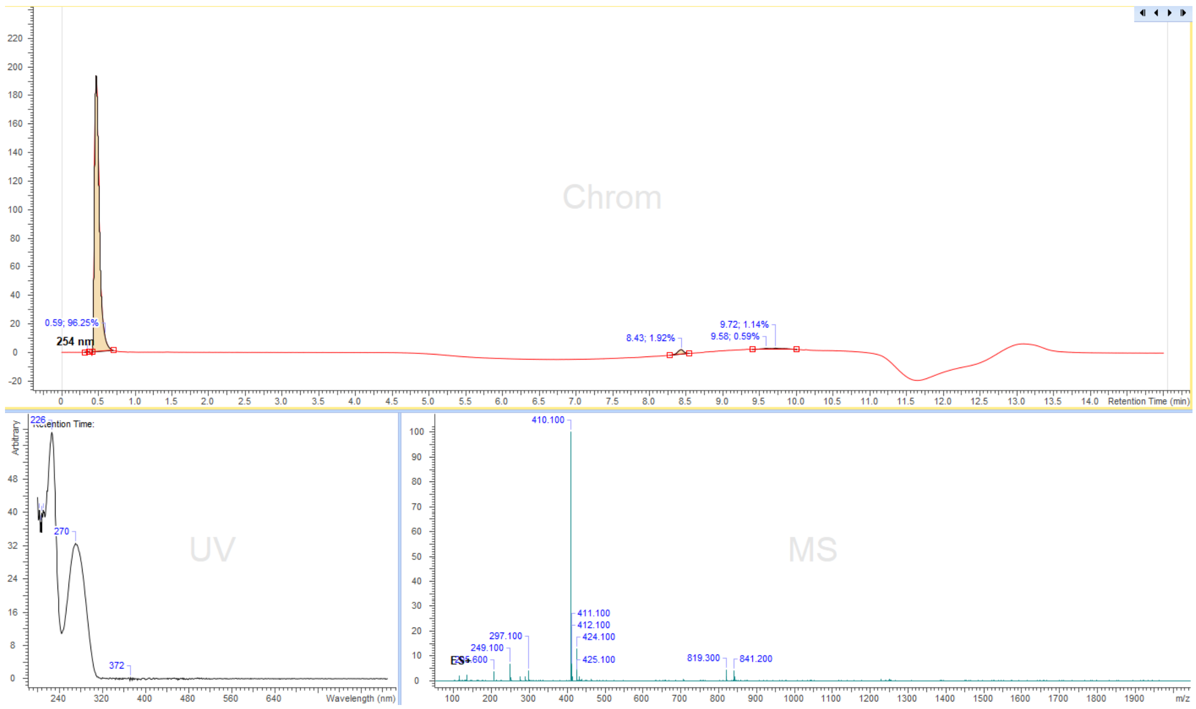


**Figure S31.** LC/MS analysis profile of compound **29ab.**

Cofactor **30ab**:

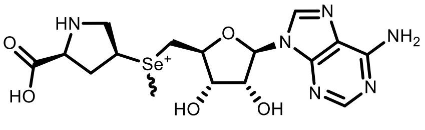

Cofactor (**30ab**)

SeAdo-cis-L-Pro (**21**) (20 mg, 0.045 mmol) was dissolved in (S)-2-chloropropanoic acid (200 µL) and methyl trifluoromethanesulfonate (30 µL, 0.25 mmol) was added to the solution, the obtained mixture was stirred at room temperature for 3h and the course of the reaction was monitored by LC/MS. Then the resulting mixture was poured into an ammonium formate buffer (25 mL, 10mM, pH = 3.5) and washed with $Et_2O$ (3x40mL). The aqueous layer was taken and concentrated to 1-3 mL and the target compound was purified by preparative HPLC (preparative column: Pursuit 10 C18 10 µm, 250×50.0 mm, Agilent, flow rate: 150 mL/min, solvent A: MeOH, solvent B: $H_2O$ (ammonium formate buffer 10 mM, pH = 3.5); temperature 25 °C, gradient A:B - 2 min 2:98 isocratic, 2-20 min 2:98 to 20:80 gradient and 20-22min 20:80 isocratic). The collected fraction of the mixture of both stereoisomers was concentrated and the concentration of the obtained solution was determined by its UV absorption ($\varepsilon_{260}$ = 15400 L·mol$^{-1}$·cm$^{-1}$). 1.7 mL, 10.0 mM of a mixture of both stereoisomers in 38% yield were obtained.

$^1$H NMR (400 MHz, deuterium oxide) δ 8.26 (s, 1H), 8.24 (d, *J* = 1.3 Hz, 1H), 6.06 (dd, *J* = 4.4, 3.2 Hz, 1H), 4.91 (ddd, *J* = 5.3, 4.4, 2.7 Hz, 1H), 4.59 – 4.49 (m, 2H), 4.38 (dt, *J* = 9.9, 6.9 Hz, 1H), 4.24 – 4.10 (m, 1H), 4.09 – 3.90 (m, 2H), 3.91 – 3.51 (m, 2H), 2.92 – 2.63 (m, 4H), 2.46 – 2.34 (m, 1H).

ESI-MS, positive mode: m/z = 459.1 [M]$^+$. HRMS (ESI) calculated for $C_{16}H_{23}N_6O_5Se$ [M]$^+$ 459.0890, found 459.0899.

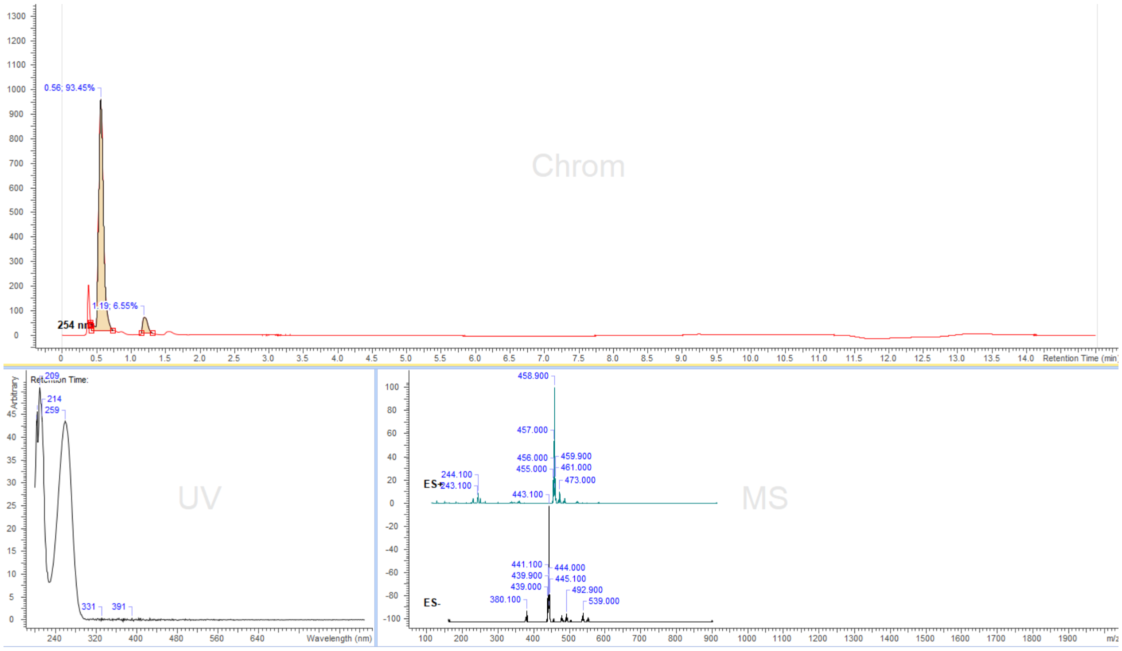


**Figure S32.** LC/MS analysis profile of compound **30ab.**

Cofactors **31a** / **31b**:

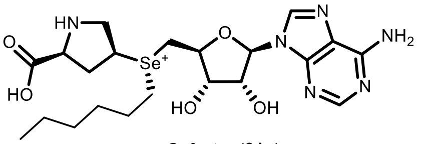

Cofactor (**31a**)

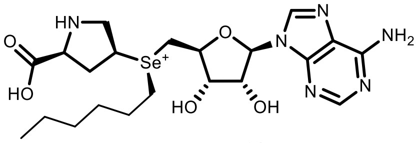

Cofactor (**31b**)

SeAdo-cis-L-Pro (**21**) (20 mg, 0.045 mmol) was dissolved in (S)-2-chloropropanoic acid (200 µL) and hexyl trifluoromethanesulfonate (**SI-4**) (50 µL, 0.25 mmol) was added to the solution, the obtained mixture was stirred at 40°C for 12h and the course of the reaction was monitored by LC/MS. Then the resulting mixture was poured into an ammonium formate buffer (25 mL, 10mM, pH = 3.5) and washed with $Et_2O$ (3x40mL). The aqueous layer was taken and concentrated to 1-3 mL and the target compound was purified by preparative HPLC (preparative column: Pursuit 10 C18 10 µm, 250×50.0 mm, Agilent, flow rate: 150 mL/min, solvent A: MeOH, solvent B: $H_2O$ (ammonium formate buffer 10 mM, pH = 3.5); temperature 25 °C, gradient A:B - 2 min 2:98 isocratic, 2-20 min 2:98 to 20:80 gradient and 20-22min 20:80 isocratic). Collected fractions of pure stereoisomers were concentrated and the concentrations of the obtained solutions were determined by their UV absorption ($\varepsilon_{260}$ = 15400 $L\cdot mol^{-1}\cdot cm^{-1}$). 0.8 mL, 9.0 mM of peak-1 (**31a**) in 16% yield, and 1.0 mL 5.8 mM of peak-2 (**31b**) in 13% yield were obtained. The selenium stereocenter configuration of peak-1 (**31a**) and peak-2 (**31b**) was assigned according to previous examples. The following analytical data were collected for cofactor **31b**:

$^1$H NMR (400 MHz, deuterium oxide) δ 8.30 (s, 1H), 8.25 (s, 1H), 6.05 (d, *J* = 4.8 Hz, 1H), 5.00 (t, *J* = 5.0 Hz, 1H), 4.58 – 4.51 (m, 2H), 4.44 (p, *J* = 6.8 Hz, 1H), 4.25 (dd, *J* = 8.2, 6.7 Hz, 1H), 3.99 – 3.77 (m, 4H), 3.37 – 3.32 (m, 2H), 2.91 (dt, *J* = 15.3, 7.8 Hz, 1H), 2.44 (dt, *J* = 14.6, 6.5 Hz, 1H), 1.49 (p, *J* = 8.0 Hz, 2H), 0.99 – 0.85 (m, 4H), 0.77 – 0.67 (m, 2H), 0.65 (t, *J* = 7.3 Hz, 3H).

ESI-MS, positive mode: m/z = 529.2 $[M]^+$. HRMS (ESI) calculated for $C_{21}H_{33}N_6O_5Se$ $[M]^+$ 529.1673, found 529.1671.

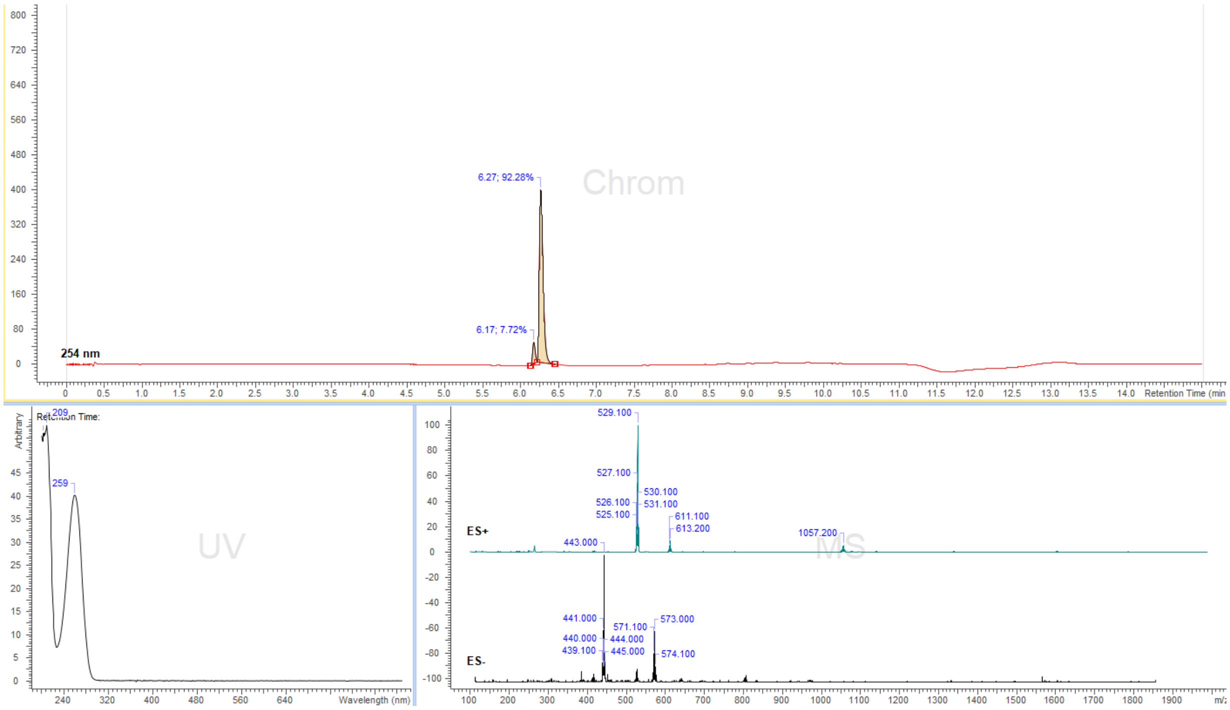


**Figure S33.** LC/MS analysis profile of compound **31b.**

Cofactors **32a** / **32b**:

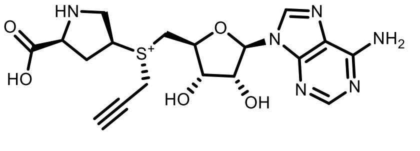

Cofactor (**32a**)

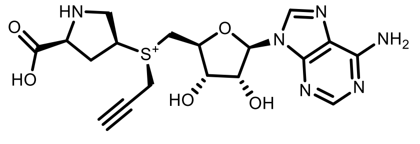

Cofactor (**32b**)

Ado-cis-L-Pro (**16**) (20 mg, 0.051 mmol) was dissolved in (S)-2-chloropropanoic acid (200 µL) and prop-2-yn-1-yl methanesulfonate (**SI-3**) (100 µL, 0.76 mmol) was added to the solution and the obtained mixture was stirred at 40$^{o}$C for 6-8h and the course of the reaction was monitored by LC/MS. Then the resulting mixture was poured into an ammonium formate buffer (25 mL, 10mM, pH = 3.5) and washed with $Et_2O$ (3x40mL). The aqueous layer was taken and concentrated to 1-3 mL and the target compound was purified by preparative HPLC (preparative column: Pursuit 10 C18 10 µm, 250×50.0 mm, Agilent, flow rate: 150 mL/min, solvent A: MeOH, solvent B: $H_2O$ (ammonium formate buffer 10 mM, pH = 3.5); temperature 25 °C, gradient A:B - 2 min 2:98 isocratic, 2-20 min 2:98 to 20:80 gradient and 20-22min 20:80 isocratic). Collected fractions of pure stereoisomers were concentrated and the concentrations of the obtained solutions were determined by their UV absorption ($\varepsilon_{260}$ = 15400 $L\cdot mol^{-1}\cdot cm^{-1}$). 0.7 mL, 2.9 mM of peak-1 (**32a**) in 4% yield, and 1.1 mL, 3.7 mM of peak-2 (**32b**) in 8% were obtained. The peak-2 (**32b**) with longer retention time was determined to be the enzymatically active R-stereoisomer and the stereocenter configuration at the sulfur atom was assigned accordingly to the example of the active AdoMet stereoisomer. The following analytical data were collected for cofactor **32b**:

ESI-MS, positive mode: m/z = 435.2 $[M]^+$. HRMS (ESI) calculated for $C_{18}H_{23}N_6O_5S$ $[M]^+$ 435.1445, found 435.1440.

The compound was not stable enough to collect $^1H$ NMR spectra or to reliably evaluate its stability in Tris buffer.

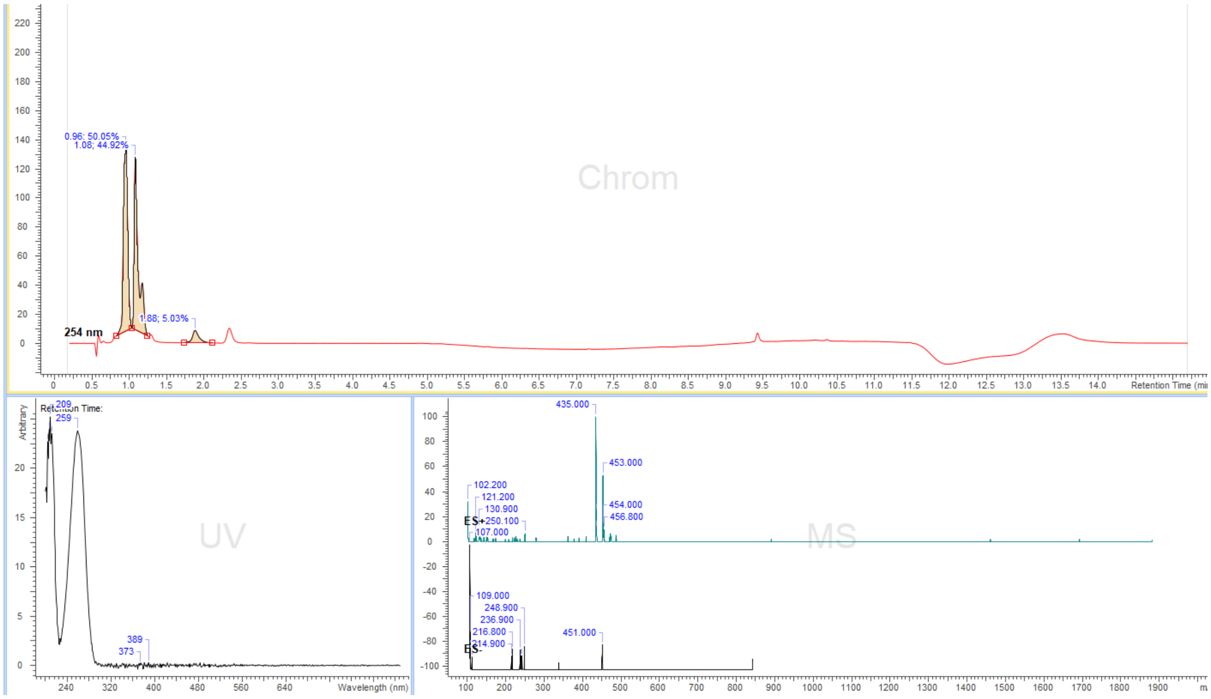


**Figure S34.** LC/MS analysis profile of compound **32b.**

Cofactors **33a** / **33b**:

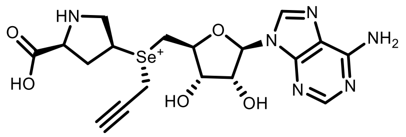

Cofactor (**33a**)

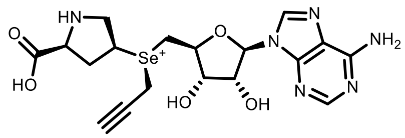

Cofactor (**33b**)

SeAdo-cis-L-Pro (**21**) (20 mg, 0.045 mmol) was dissolved in (S)-2-chloropropanoic acid (200 µL) and prop-2-yn-1-yl methanesulfonate (**SI-3**) (90 µL, 0.67 mmol) was added to the solution and the obtained mixture was stirred at room temperature for 24h and the course of the reaction was monitored by LC/MS. Then the resulting mixture was poured into an ammonium formate buffer (25 mL, 10mM, pH = 3.5) and washed with $Et_2O$ (3x40mL). The aqueous layer was taken and concentrated to 1-3 mL and the target compound was purified by preparative HPLC (preparative column: Pursuit 10 C18 10 µm, 250×50.0 mm, Agilent, flow rate: 150 mL/min, solvent A: MeOH, solvent B: $H_2O$ (ammonium formate buffer 10 mM, pH = 3.5); temperature 25 °C, gradient A:B - 2 min 2:98 isocratic, 2-20 min 2:98 to 20:80 gradient and 20-22min 20:80 isocratic). Collected fractions of pure stereoisomers were concentrated and the concentrations of the obtained solutions were determined by their UV absorption ($\varepsilon_{260}$ = 15400 $L{\cdot}mol^{-1}{\cdot}cm^{-1}$). 0.9 mL, 7.6 mM of peak-1 (**33a**) in 15% yield, and 1.0 mL 8.5 mM of peak-2 (**33b**) in 19% yield were obtained. The peak-2 (**33b**) with the longer retention time was determined to be the enzymatically active R-stereoisomer and the stereocenter configuration at the selenium atom was assigned accordingly to the example of the active AdoMet stereoisomer. The following analytical data were collected for cofactor **33b**:

$^1$H NMR (400 MHz, deuterium oxide) δ 8.26 (s, 1H), 8.25 (s, 1H), 6.05 (d, *J* = 4.3 Hz, 1H), 4.92 (dd, *J* = 5.6, 4.3 Hz, 1H), 4.61 (t, *J* = 5.7 Hz, 1H), 4.58 – 4.51 (m, 1H), 4.48 (t, *J* = 6.7 Hz, 1H), 4.29 (d, *J* = 2.7 Hz, 2H), 4.19 – 4.05 (m, 3H), 3.96 – 3.84 (m, 2H), 3.25 (t, *J* = 2.6 Hz, 1H), 2.73 (dt, *J* = 15.6, 7.9 Hz, 1H), 2.52 (dt, *J* = 15.0, 6.0 Hz, 1H).

ESI-MS, positive mode: m/z = 483.1 $[M]^+$. HRMS (ESI) calculated for $C_{18}H_{23}N_6O_5Se$ $[M]^+$ 483.0891, found 483.0903.

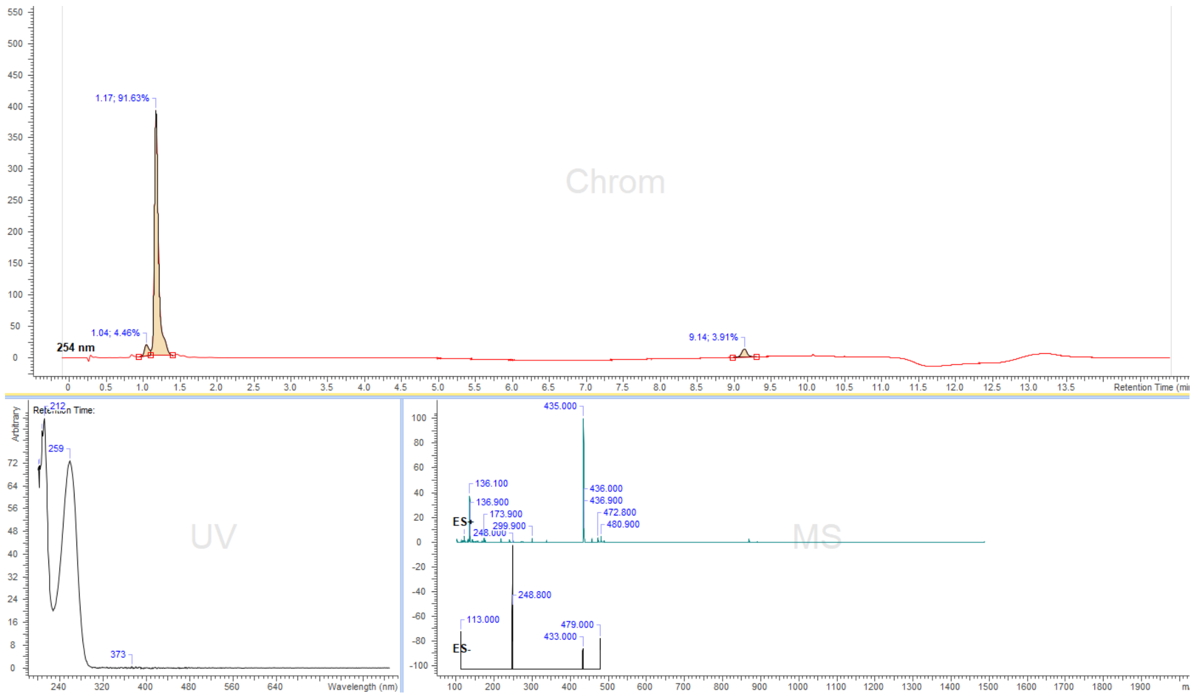


**Figure S35.** LC/MS analysis profile of compound **33b.**

Cofactors **34a** / **34b**:

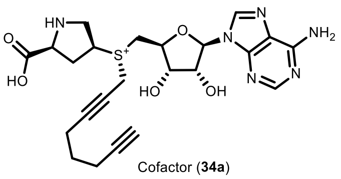

Cofactor (**34a**)

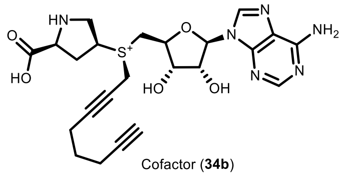

Cofactor (**34b**)

Ado-cis-L-Pro (**16**) (20 mg, 0.051 mmol) was dissolved in (S)-2-chloropropanoic acid (200 µL) and octa-2,7-diyn-1-yl methanesulfonate (**SI-2**) (100 µL, 0.51 mmol) was added to the solution and the obtained mixture was stirred at 40°C for 36h and the course of the reaction was monitored by LC/MS. Then the resulting mixture was poured into an ammonium formate buffer (25 mL, 10mM, pH = 3.5) and washed with $Et_2O$ (3x40mL). The aqueous layer was taken and concentrated to 1-3 mL and the target compound was purified by preparative HPLC (preparative column: Pursuit 10 C18 10 µm, 250×50.0 mm, Agilent, flow rate: 150 mL/min, solvent A: MeOH, solvent B: $H_2O$ (ammonium formate buffer 10 mM, pH = 3.5); temperature 25 °C, gradient A:B - 2 min 2:98 isocratic, 2-20 min 2:98 to 20:80 gradient and 20-22min 20:80 isocratic). Collected fractions of pure stereoisomers were concentrated and the concentrations of the obtained solutions were determined by their UV absorption ($\varepsilon_{260}$ = 15400 $L \cdot mol^{-1} \cdot cm^{-1}$). 1.5 mL, 7.5 mM of peak-1 (**34a**) in 22% yield, and 1.3 mL, 9.4 mM of peak-2 (**34b**) in 24% were obtained. The peak-2 (**34b**) with longer retention time was determined to be the enzymatically active stereoisomer and the stereocenter configuration at the sulfur atom was assigned accordingly to the example of the active AdoMet stereoisomer. The following analytical data were collected for cofactor **34b**:

$^1$H NMR (400 MHz, deuterium oxide) δ 8.40 (s, 1H), 8.40 (s, 1H), 6.12 (d, *J* = 4.2 Hz, 1H), 4.90 (dd, *J* = 5.6, 4.2 Hz, 1H), 4.64 (t, *J* = 5.7 Hz, 1H), 4.55 – 4.50 (m, 3H), 4.34 (dd, *J* = 8.2, 6.6 Hz, 1H), 4.14 – 4.07 (m, 2H), 4.03 – 3.87 (m, 2H), 2.98 (dt, *J* = 15.3, 7.9 Hz, 1H), 2.59 (dt, *J* = 14.7, 6.2 Hz, 1H), 2.45 (t, *J* = 7.1 Hz, 2H), 2.36 (t, *J* = 2.7 Hz, 1H), 2.28 (td, *J* = 7.0, 2.7 Hz, 2H), 2.02 (s, 1H), 1.71 (p, *J* = 7.0 Hz, 2H).

ESI-MS, positive mode: m/z = 501.2 $[M]^+$. HRMS (ESI) calculated for $C_{23}H_{29}N_6O_5S$ $[M]^+$ 501.1915, found 501.1931.

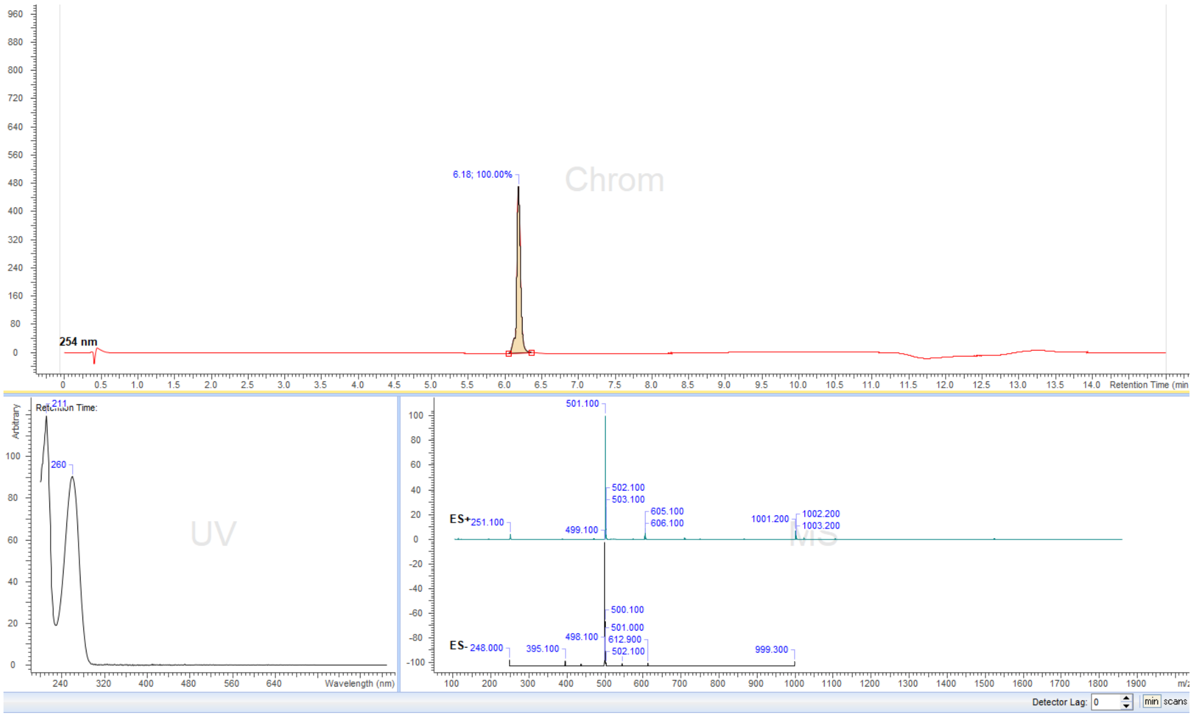


**FigureS36.** LC/MS analysis profile of compound **34b.**

Cofactors **35a** / **35b**:

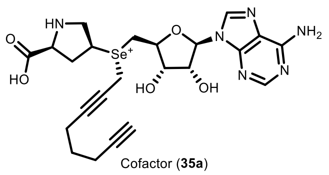

Cofactor (**35a**)

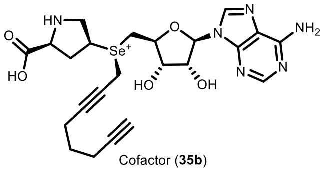

Cofactor (**35b**)

SeAdo-cis-L-Pro (**21**) (20 mg, 0.045 mmol) was dissolved in (S)-2-chloropropanoic acid (200 µL) and octa-2,7-diyn-1-yl methanesulfonate (**SI-2**) (90 µL, 0.45 mmol) was added to the solution and the obtained mixture was stirred at room temperature for 24h-48h until all starting compound was consumed and the course of the reaction was monitored by LC/MS. Then the resulting mixture was poured into an ammonium formate buffer (25 mL, 10mM, pH = 3.5) and washed with $Et_2O$ (3x40mL). The aqueous layer was taken and concentrated to 1-3 mL and the target compound was purified by preparative HPLC (preparative column: Pursuit 10 C18 10 µm, 250×50.0 mm, Agilent, flow rate: 150 mL/min, solvent A: MeOH, solvent B: $H_2O$ (ammonium formate buffer 10 mM, pH = 3.5); temperature 25 °C, gradient A:B - 2 min 2:98 isocratic, 2-20 min 2:98 to 20:80 gradient and 20-22min 20:80 isocratic). Collected fractions of pure stereoisomers were concentrated and the concentrations of the obtained solutions were determined by their UV absorption ($\varepsilon_{260}$ = 15400 L·mol$^{-1}$·cm$^{-1}$). 1.2 mL, 7.4 mM of peak-1 (**35a**) in 20% yield, and 1.5 mL 7.8 mM of peak-2 (**35b**) in 26% yield were obtained. The peak-2 (**35b**) with longer retention time was determined to be the enzymatically active R-stereoisomer and the stereocenter configuration at the selenium atom was assigned accordingly to the example of the active AdoMet stereoisomer. The following analytical data were collected for cofactor **35b**:

$^1$H NMR (400 MHz, deuterium oxide) δ 8.26 (s, 1H), 8.24 (s, 1H), 6.07 (d, *J* = 2.8 Hz, 1H), 4.97 (dd, *J* = 5.1, 2.8 Hz, 1H), 4.64 (dd, *J* = 6.8, 5.1 Hz, 1H), 4.59 – 4.52 (m, 1H), 4.31 – 4.21 (m, 2H), 4.21 – 4.15 (m, 2H), 3.93 (dd, *J* = 12.2, 2.5 Hz, 1H), 3.86 (d, *J* = 6.6 Hz, 2H), 3.68 (dd, *J* = 12.2, 10.7 Hz, 1H), 2.90 (dt, *J* = 14.8, 7.9 Hz, 1H), 2.50 (dt, *J* = 14.8, 6.4 Hz, 1H), 2.35 (t, *J* = 2.6 Hz, 1H), 2.27 – 2.13 (m, 4H), 1.59 – 1.49 (m, 2H).

ESI-MS, positive mode: m/z = 549.2 $[M]^+$. HRMS (ESI) calculated for $C_{23}H_{29}N_6O_5Se$ $[M]^+$ 549.1360, found 549.1377.

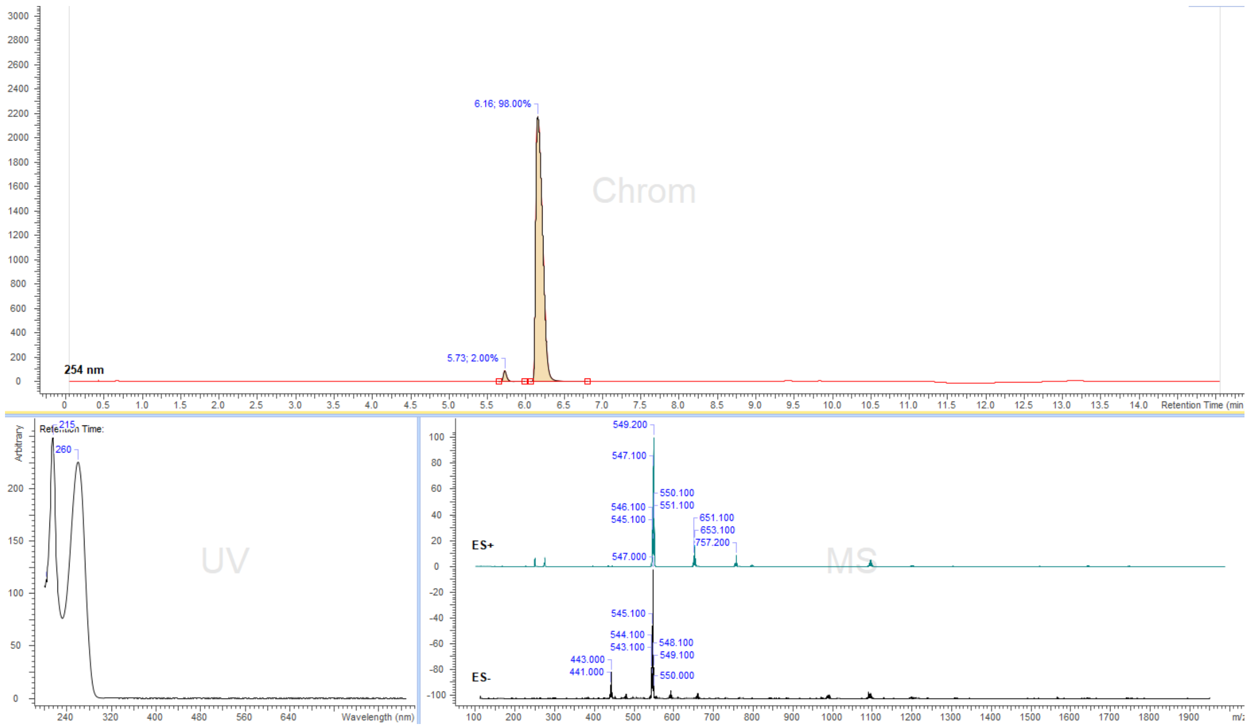


**Figure S37.** LC/MS analysis profile of compound **35b.**

Cofactors **36a** / **36b**:

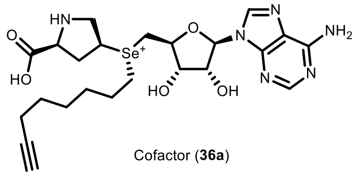

Cofactor (**36a**)

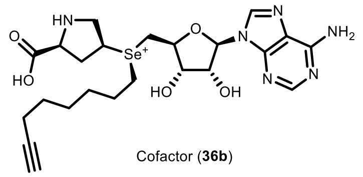

Cofactor (**36b**)

SeAdo-cis-L-Pro (**21**) (20 mg, 0.045 mmol) was dissolved in (S)-2-chloropropanoic acid (200 µL) and oct-7-yn-1-yl trifluoromethanesulfonate (**SI-5**) (45 µL, 0.25 mmol) was added to the solution, the obtained mixture was stirred at 40°C for 12h, the course of the reaction was monitored by LC/MS. Then the resulting mixture was poured into an ammonium formate buffer (25 mL, 10mM, pH = 3.5) and washed with $Et_2O$ (3x40mL). The aqueous layer was taken and concentrated to 1-3 mL and the target compound was purified by preparative HPLC (preparative column: Pursuit 10 C18 10 µm, 250×50.0 mm, Agilent, flow rate: 150 mL/min, solvent A: MeOH, solvent B: $H_2O$ (ammonium formate buffer 10 mM, pH = 3.5); temperature 25 °C, gradient A:B - 2 min 2:98 isocratic, 2-20 min 2:98 to 20:80 gradient and 20-22 min 20:80 isocratic). Collected fractions of pure stereoisomers were concentrated and the concentrations of the obtained solutions were determined by their UV absorption ($\varepsilon_{260}$ = 15400 $L\cdot mol^{-1}\cdot cm^{-1}$). 1.0 mL, 6.6 mM of peak-1 (**36a**) in 14% yield, and 0.8 mL 10.7 mM of peak-2 (**36b**) in 18% yield were obtained. The selenium stereocenter configuration of peak-1 (**36a**) and peak-2 (**36b**) was assigned according to previous examples. The following analytical data were collected for cofactor **36b**:

$^1H$ NMR (400 MHz, deuterium oxide) δ 8.31 (s, 1H), 8.25 (s, 1H), 6.05 (d, *J* = 4.9 Hz, 1H), 4.99 (t, *J* = 5.1 Hz, 1H), 4.57 – 4.51 (m, 2H), 4.45 (p, *J* = 7.1 Hz, 1H), 4.25 (dd, *J* = 8.3, 6.7 Hz, 1H), 4.00 – 3.77 (m, 4H), 3.39 – 3.31 (m, 2H), 2.91 (dt, *J* = 14.5, 7.8 Hz, 1H), 2.47 – 2.39 (m, 1H), 2.28 (t, *J* = 2.6 Hz, 1H), 1.98 (td, *J* = 7.1, 2.6 Hz, 2H), 1.52 (p, *J* = 7.8 Hz, 2H), 1.14 (p, *J* = 7.2 Hz, 2H), 0.97 – 0.82 (m, 4H).

ESI-MS, positive mode: m/z = 553.2 $[M]^+$. HRMS (ESI) calculated for $C_{23}H_{33}N_6O_5Se$ $[M]^+$ 553.1673, found 553.1676.

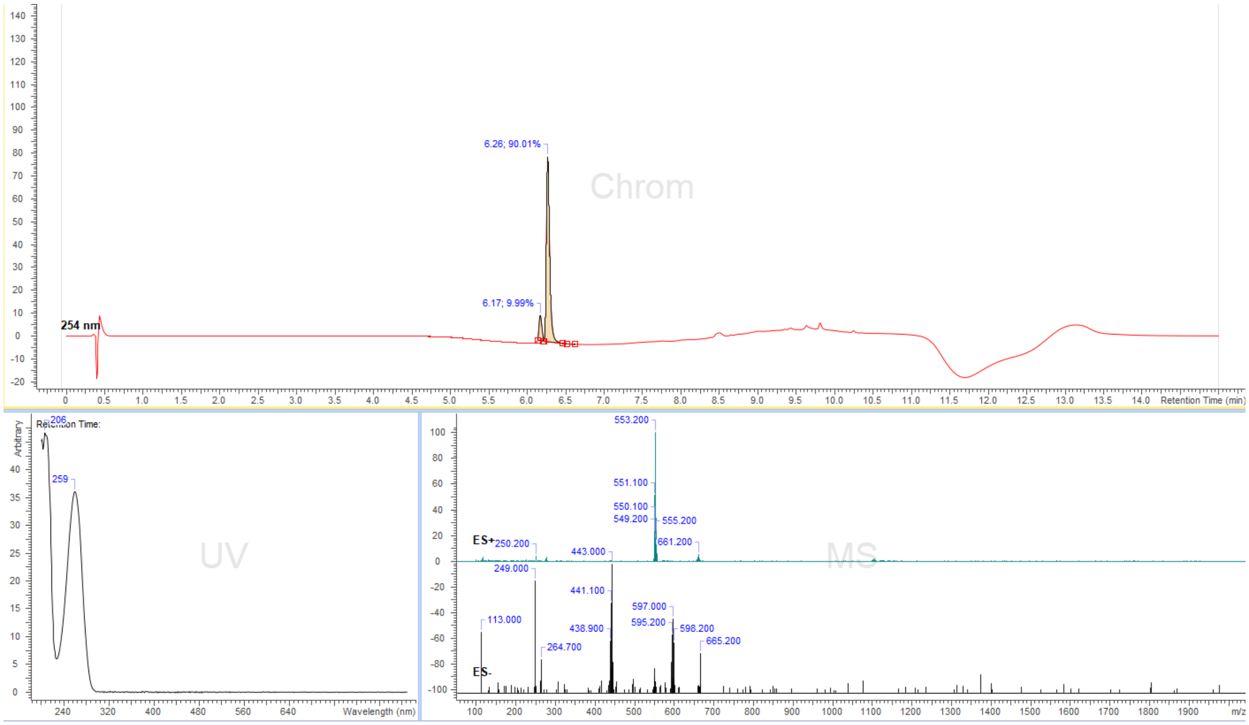


**Figure S38.** LC/MS analysis profile of compound **36b.**

Cofactors **37a** and **37b**

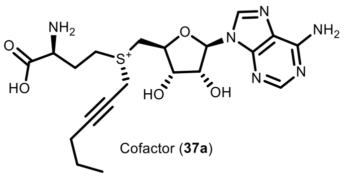

Cofactor (**37a**)

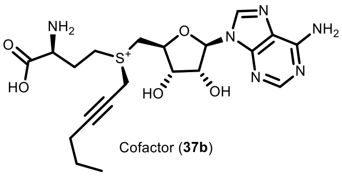

Cofactor (**37b**)

S-Adenosyl-L-homocysteine (10 mg, 0.026 mmol) was dissolved in a (S)-2-chloropropanoic acid (200 µL) and hex-2-yn-1-yl methanesulfonate (**SI-I**) (60 µL, 0.34 mmol) was added to the solution and the obtained mixture was stirred at 40°C for 24h, wherein the course of the reaction was monitored by LC/MS. Then the resulting mixture was poured into an ammonium formate buffer (25 mL, 10mM, pH = 3.5) and washed with $Et_2O$ (3x25 mL). The aqueous layer was taken and concentrated to 1-3 mL and the target compound was purified by preparative HPLC (preparative column: Pursuit 10 C18 10 µm, 250×50.0 mm, Agilent, flow rate: 150 mL/min, solvent A: MeOH, solvent B: $H_2O$ (ammonium formate buffer 10 mM, pH = 3.5); temperature 25 °C, gradient A:B - 2 min 2:98 isocratic, 2-20 min 2:98 to 20:80 gradient and 20-22min 20:80 isocratic). Collected fractions of pure stereoisomers were concentrated and the concentrations of the obtained solutions were determined by their UV absorption ($\varepsilon_{260}$ = 15400 $L \cdot mol^{-1} \cdot cm^{-1}$). 0.8 mL, 5.8 mM of peak-1 (**37a**) in 18% yield, and 0.8 mL 5.2 mM of peak-2 (**37b**) in 16% yield were obtained. The peak-2 (**37b**) with longer retention time was determined to be the enzymatically active stereoisomer and the stereocenter configuration at the selenium atom was assigned accordingly to the example of the active AdoMet stereoisomer. The following analytical data were collected for cofactor **37b**:

$^1$H NMR (400 MHz, deuterium oxide) δ 8.29 (s, 1H), 8.28 (s, 1H), 6.10 (d, J = 3.9 Hz, 1H), 4.97 (dd, J = 5.5, 3.9 Hz, 1H), 4.68 (d, J = 5.9 Hz, 1H), 4.59 – 4.53 (m, 1H), 4.35 (t, J = 2.3 Hz, 2H), 4.03 – 3.89 (m, 2H), 3.80 (t, J = 6.5 Hz, 1H), 3.65 (dt, J = 13.2, 7.8 Hz, 1H), 3.51 (dt, J = 13.5, 7.6 Hz, 1H), 2.39 – 2.29 (m, 2H), 2.23 (tt, J = 6.5, 2.1 Hz, 2H), 1.52 – 1.42 (m, 2H), 0.89 (t, J = 7.4 Hz, 3H).

ESI-MS, positive mode: m/z = 465.2 $[M]^+$.

HRMS (ESI) calculated. for $C_{20}H_{29}N_6O_5S$ $[M]^+$ 465.1915, found 465.1924.

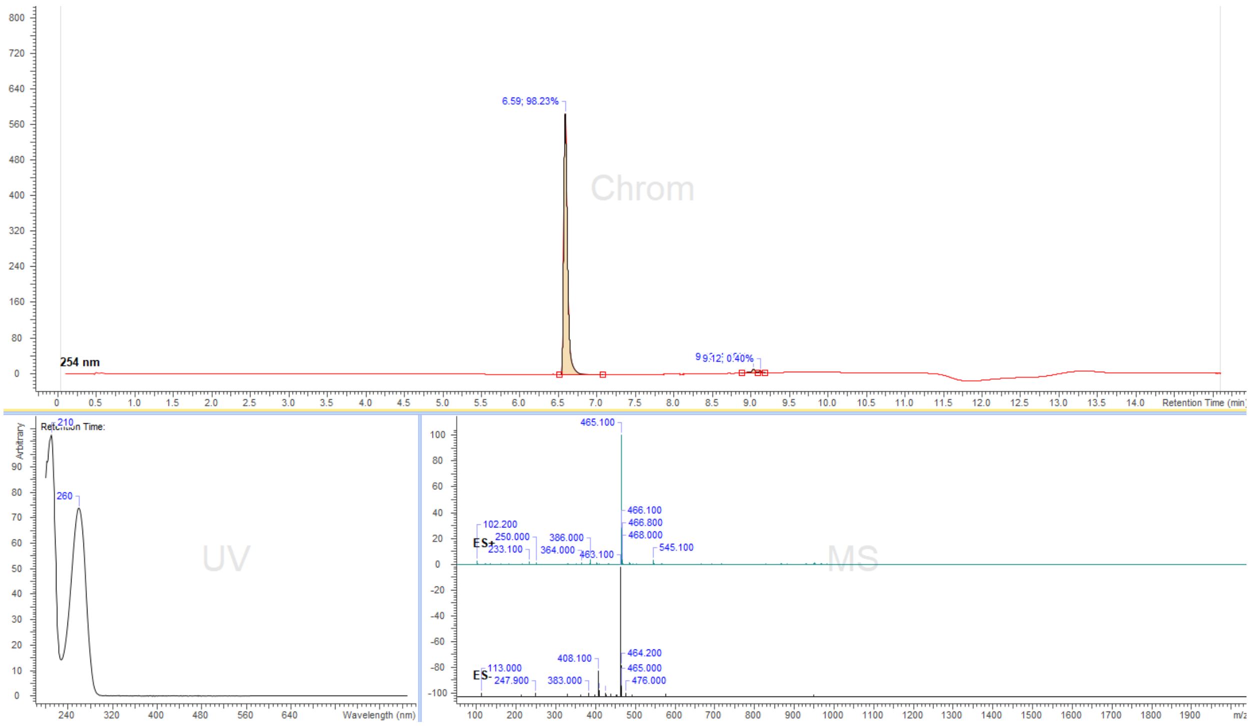


**Figure S39.** LC/MS analysis profile of compound **37b.**

Cofactors **38a** and **38b**:

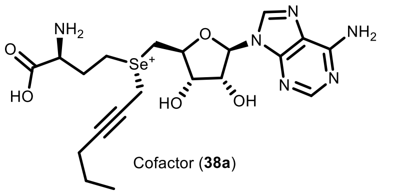


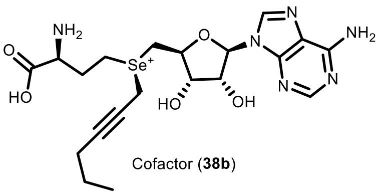


SeAdoHcy (**SI-10**) (10 mg, 0.023 mmol) was dissolved in a (S)-2-chloropropanoic acid (200 µL) and hex-2-yn-1-yl methanesulfonate (**SI-1**) (60 µL, 0.34 mmol) was added to the solution and the obtained mixture was stirred at room temperature for 24h, wherein the course of the reaction was monitored by LC/MS. Then the resulting mixture was poured into water + 0.1% TFA (25 mL) and washed with $Et_2O$ (3x25mL). The aqueous layer was taken and concentrated to 1-3 mL and the target compound was purified by preparative HPLC (preparative column: Pursuit 10 C18 10 µm, 250×50.0 mm, Agilent, flow rate: 150 mL/min, solvent A: MeOH, solvent B: $H_2O$ (+ 0.1 % TFA); temperature 25 °C, gradient A:B - 2 min 2:98 isocratic, 2-20 min 2:98 to 20:80 gradient and 20-22min 20:80 isocratic). Collected fractions of pure stereoisomers were concentrated and the concentrations of the obtained solutions were determined by their UV absorption ($\varepsilon_{260}$ = 15400 L·mol$^{-1}$·cm$^{-1}$). 1.2 mL, 3.8 mM of peak-1 (**38a**) in 19% yield, and 1.0 mL 4.0 mM of peak-2 (**38b**) in 17% yield were obtained. The peak-2 (**38b**) with the longer retention time was determined to be the enzymatically active stereoisomer and the stereocenter configuration at the selenium atom was assigned accordingly to the example of the active AdoMet stereoisomer. The following analytical data were collected for cofactor **38b**:

$^1$H NMR (400 MHz, $D_2O$) δ 8.24 (s, 1H), 8.23 (s, 1H), 5.92 (d, J = 3.6 Hz, 1H), 4.61 – 4.58 (m, 1H), 4.45 – 4.40 (m, 1H), 4.36 – 4.30 (m, 1H), 4.00 (t, J = 2.2 Hz, 2H), 3.96 – 3.88 (m, 1H), 3.73 (dd, J = 12.3, 2.8 Hz, 1H), 3.63 (dd, J = 12.6, 9.0 Hz, 1H), 3.41 – 3.23 (m, 2H), 2.36 – 2.16 (m, 2H), 2.10 – 2.03 (m, 2H), 1.28 (h, J = 7.0 Hz, 2H), 0.69 (t, J = 7.4 Hz, 3H).

ESI-MS, positive mode: m/z = 513.1 $[M]^+$.

HRMS (ESI) calculated for $C_{20}H_{29}N_6O_5Se$ $[M]^+$ 513.1360, found 513.1359

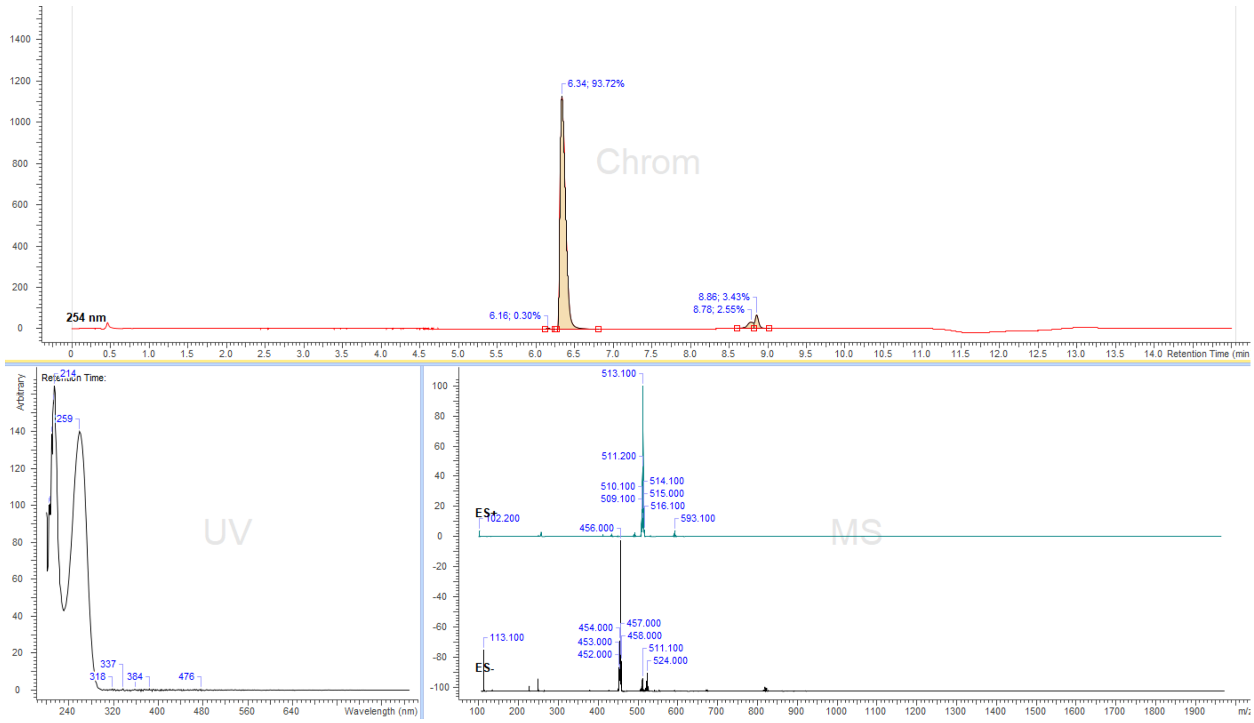


**Figure S40.** LC/MS analysis profile of compound **38b.**

**SeAdoMet (39ab):**

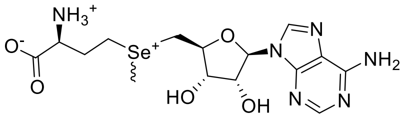

SeAdoMet (**39ab**)

SeAdoHcy (**SI-10**) (5 mg, 0.012 mmol) was dissolved in (S)-2-chloropropanoic acid (100 µL) and methyl trifluoromethanesulfonate (6.6 µL, 0.6 mmol) was added to the solution and the obtained mixture was stirred at room temperature for 2h, the course of the reaction was monitored by LC/MS. Then the resulting mixture was poured into an ammonium formate buffer (25 mL, 10 mM, pH = 3.5) and washed with $Et_2O$ (3 x 40mL). The aqueous layer was taken and concentrated to 1-3 mL and the target compound was purified by preparative HPLC (preparative column: Pursuit 10 C18 10 µm, 250 × 50.0 mm, Agilent, flow rate: 150 mL/min, solvent A: MeOH, solvent B: $H_2O$ + 0.1% TFA; temperature 25 °C, gradient A:B - 2 min 2:98 isocratic, 2-20 min 2:98 to 20:80 gradient and 20-22min 20:80 isocratic). Collected fractions of mixture of stereoisomers were concentrated and the concentration of the obtained solution was determined by the UV absorption ($\varepsilon_{260}$ = 15400 L·mol$^{-1}$·cm$^{-1}$). 0.8 mL, 4.8 mM of SeAdoMet (**39ab**) were obtained in 32 % yield.

$^1$H NMR (400 MHz, $D_2O$) δ 7.80 (s, 1H), 7.79 (s, 1H), 5.47 (d, J = 3.8 Hz, 1H), 4.15 (ddd, J = 10.9, 5.6, 3.9 Hz, 1H), 3.95 (t, J = 5.9 Hz, 1H), 3.92 – 3.84 (m, 1H), 3.47 (ddd, J = 25.0, 7.5, 5.5 Hz, 1H), 3.30 – 3.17 (m, 2H), 3.01 – 2.75 (m, 2H), 2.22 – 2.13 (m, 3H), 1.90 – 1.69 (m, 2H).

ESI-MS, positive mode: m/z = 447.1 $[M]^+$.

HRMS (ESI) calculated for $C_{15}H_{23}N_6O_5Se$ $[M]^+$ 447.0890, found 447.0894.

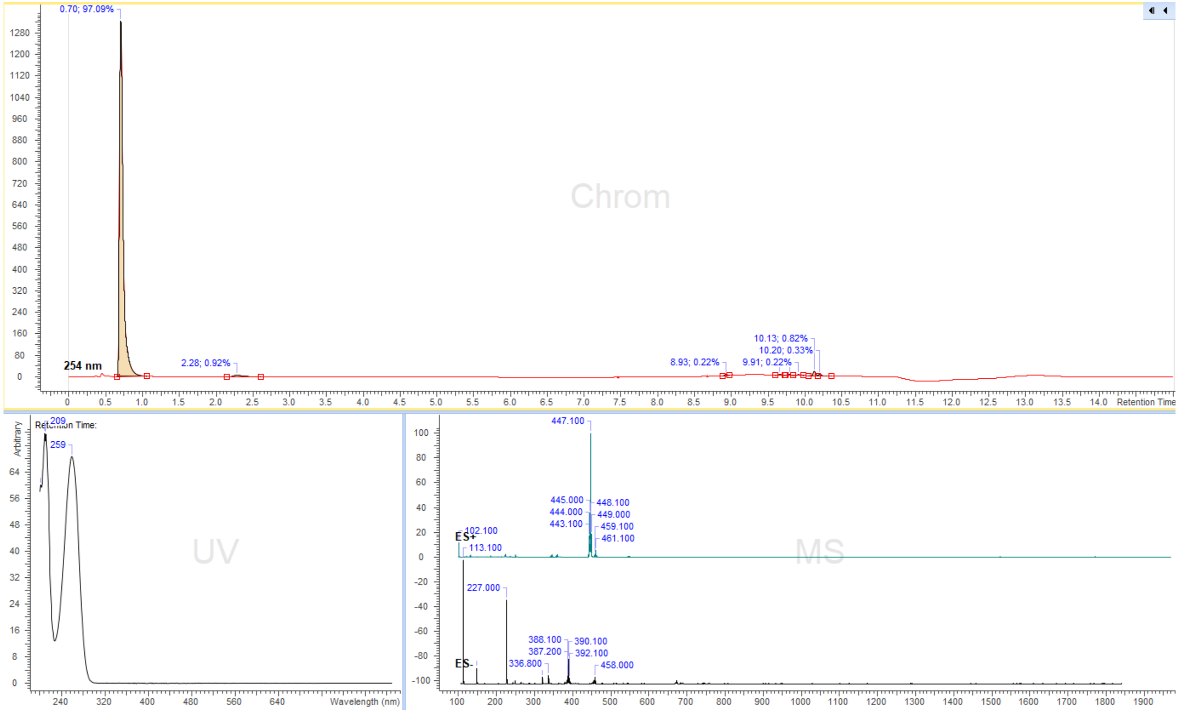


**Figure S41.** LC/MS analysis profile of compound **39ab.**

SeAdoYn (**40ab**):

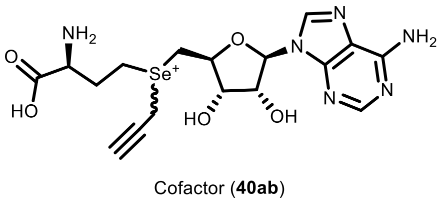


SeAdoHcy (**SI-10**) (10 mg, 0.023 mmol) was dissolved in (S)-2-chloropropanoic acid (200 µL) and propargyl mesylate (**SI-3**) (22 µL, 0.23 mmol) was added to the solution and the obtained mixture was stirred at room temperature for 14h, wherein the course of the reaction was monitored by LC/MS. Then the resulting mixture was poured into an ammonium formate buffer (25 mL, 10mM, pH = 3.5) and washed with $Et_2O$ (3x40mL). The aqueous layer was taken and concentrated to 1-3 mL and the target compound was purified by preparative HPLC (preparative column: Pursuit 10 C18 10 µm, 250×50.0 mm, Agilent, flow rate: 150 mL/min, solvent A: MeOH, solvent B: $H_2O$ + 0.1% TFA; temperature 25 °C, gradient A:B - 2 min 2:98 isocratic, 2-20 min 2:98 to 20:80 gradient and 20-22min 20:80 isocratic). The collected fraction of a mixture of both stereoisomers was concentrated and concentration of the obtained solution was determined by their UV absorption ($\varepsilon_{260}$ = 15400 $L\cdot mol^{-1}\cdot cm^{-1}$). 1.3 mL, 5.1 mM of a mixture of both stereoisomers in 37% yield were obtained.

$^1H$ NMR (400 MHz, $D_2O$) δ 8.07 – 8.02 (m, 2H), 5.72 (dd, J = 7.4, 3.5 Hz, 1H), 4.43 – 4.37 (m, 1H), 4.26 – 4.19 (m, 1H), 4.19 – 4.11 (m, 1H), 3.82 (d, J = 2.7 Hz, 1H), 3.80 – 3.36 (m, 4H), 3.28 – 3.10 (m, 2H), 2.81 – 2.70 (m, 1H), 2.16 – 1.96 (m, 2H).

ESI-MS, positive mode: m/z = 471.1$[M]^+$.

HRMS (ESI) calculated for $C_{17}H_{23}N_6O_5Se$ $[M]^+$ 471.0890, found 471.0880.

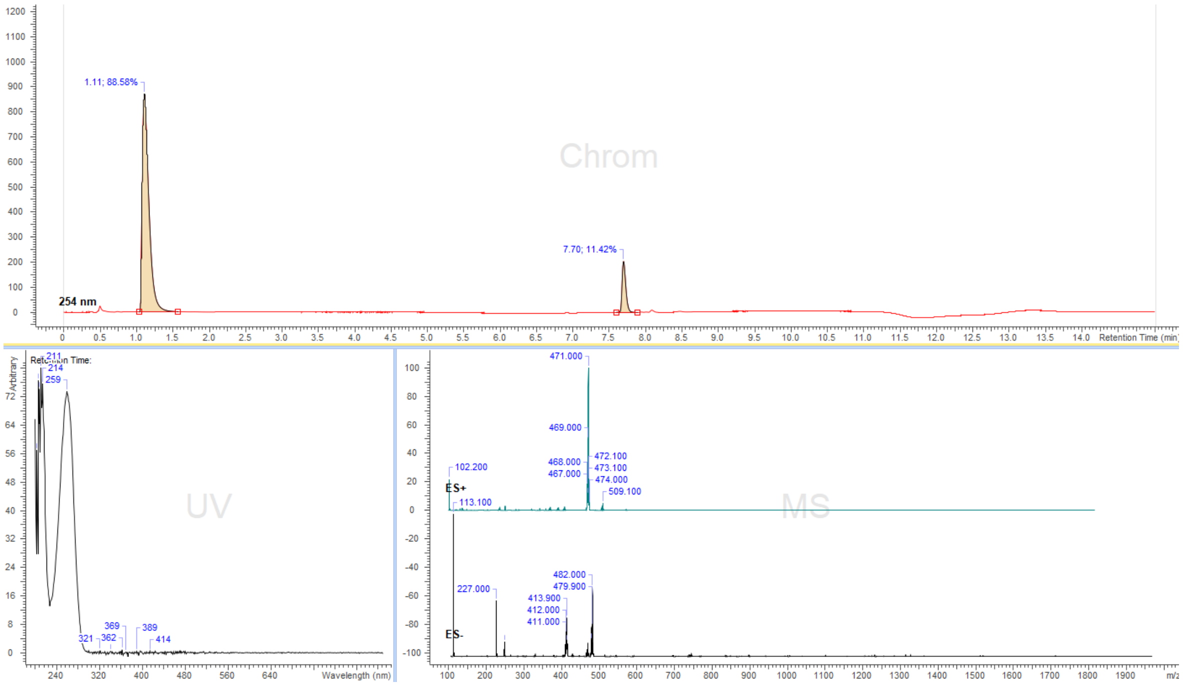


**Figure S42.** LC/MS analysis profile of compound **40ab.**

Cofactor **41**:

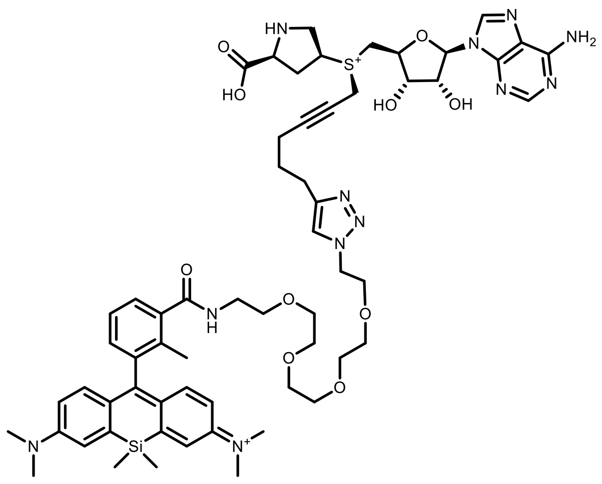

Cofactor (**41**)

A cofactor **34b** solution (250 µL of 10.2 mM in 10mM pH 3.5 ammonium formate buffer) was mixed with a solution of 4-MeSiR-PEG$_4$-N$_3$ (**SI-8**) (500 µL of 10 mM in 10mM ammonium formate buffer pH = 3.5). Fresh solutions of $CuSO_4 \cdot 5H_2O$ (10 mg in 100 µL of AF buffer), ascorbic acid (10 mg in 100 µL of AF buffer) and (BimC$_4$A)$_3$ (1 mg in 10 µL of water) were prepared. 10 µL of the $CuSO_4$ solution were mixed with 5 µL of the (BimC$_4$A)$_3$ solution and sonicated for 1 minute, then 30 µL of the ascorbic acid solution were added (an immediate color change was observed) and again sonicated for 1 minute (it was noticed that $CuSO_4$ facilitates solvolysis of starting compound **34b** and solvolysis of product **41**. Therefore, it is advised to use as little as possible of $CuSO_4$). Then 20 µL of the obtained mixture were transferred to the reaction vial containing the solution of **34b** and **SI-8**. The reaction vial was sonicated for 2 min and then left at room temperature for 60 min. The progress of the reaction was monitored by LC/MS every 20 min. Once the conversion reached >70%, the reaction was diluted with water to a volume of 2 mL and the product was purified by preparative HPLC (preparative column: Pursuit 10 C18 10 µm, 250×50.0 mm, Agilent, flow rate: 150 mL/min, solvent A: MeOH, solvent B: $H_2O$ (ammonium formate buffer 10 mM, pH = 3.5), temperature 25 °C, gradient A:B - 2 min 30:70 isocratic, 2-20 min 30:70 to 100:0 gradient and 20-22min 100:0 isocratic). Collected fractions of pure product were concentrated and the concentrations of the obtained solutions were determined by their UV absorption ($\varepsilon_{640}$ = 150 000 $L \cdot mol^{-1} \cdot cm^{-1}$). 1.2 mL, 0.8 mM in 38% yield were obtained.

ESI-MS, positive mode: m/z = 594.3 $[M]^{2+}$. HRMS (ESI) calculated for $C_{60}H_{80}N_{12}O_{10}SSi$ $[M]^{2+}$ 594.2800, found 594.2801.

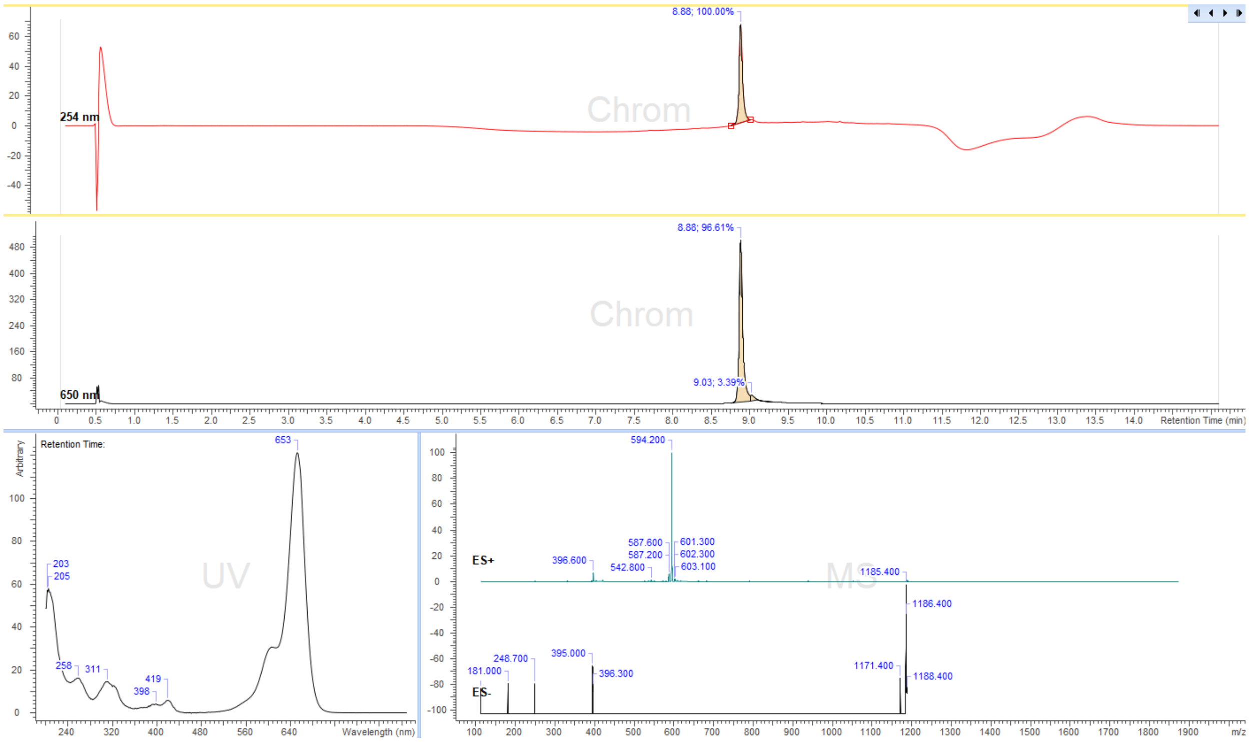


**Figure S43.** LC/MS analysis profile of compound **41.**

Cofactor **42**:

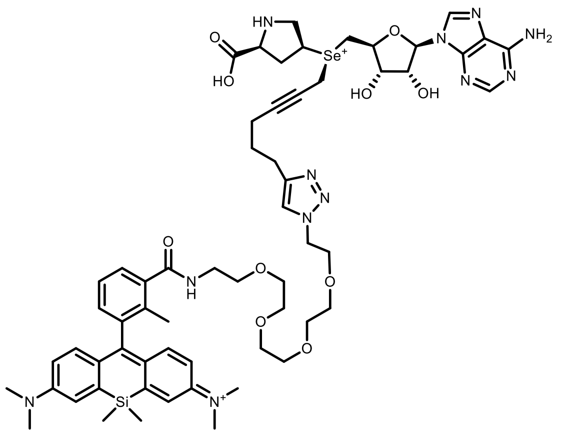

Cofactor (**42**)

A cofactor **35b** solution (250 µL of 7.8 mM in 10mM pH 3.5 ammonium formate buffer) was mixed with a solution of 4-MeSiR-PEG$_4$-N$_3$ (**SI-8**) (500 µL of 10 mM in 10mM ammonium formate buffer pH = 3.5). Fresh solutions of $CuSO_4 \cdot 5H_2O$ (10 mg in 100 µL of AF buffer), ascorbic acid (10 mg in 100 µL of AF buffer) and (BimC$_4$A)$_3$ (1 mg in 10 µL of water) were prepared. 10 µL of the $CuSO_4$ solution were mixed with 5 µL of (BimC$_4$A)$_3$ solution and sonicated for 1 minute, then 30 µL of the ascorbic acid solution were added (an immediate color change was observed) and again sonicated for 1 minute (it was noticed that $CuSO_4$ facilitates solvolysis of starting compound **35b** and solvolysis of product **42**. Therefore, it is advised to use as little as possible of $CuSO_4$). Then 10 µL of the obtained mixture were transferred to the reaction vial containing the solution of **35b** and **SI-8**. The reaction vial was sonicated for 2 min and then left at room temperature for 60 min. The progress of the reaction was monitored by LC/MS every 20 min. Once the conversion reached >70% the reaction was diluted with water to a volume of 2 mL and the product was purified by preparative HPLC (preparative column: Pursuit 10 C18 10 µm, 250×50.0 mm, Agilent, flow rate: 150 mL/min, solvent A: MeOH, solvent B: $H_2O$ (ammonium formate buffer 10 mM, pH = 3.5), temperature 25 °C, gradient A:B - 2 min 30:70 isocratic, 2-20 min 30:70 to 100:0 gradient and 20-22min 100:0 isocratic). Collected fractions of pure product were concentrated and the concentrations of the obtained solutions were determined by their UV absorption ($\varepsilon_{640}$ = 150 000 L·mol$^{-1}$·cm$^{-1}$). 1.2 mL, 0.5 mM in 30% yield were obtained.

ESI-MS, positive mode: m/z = 618.3 [M]$^{2+}$. HRMS (ESI) calculated for $C_{60}H_{80}N_{12}O_{10}SeSi$ [M]$^{2+}$ 618.2526, found 618.2564.

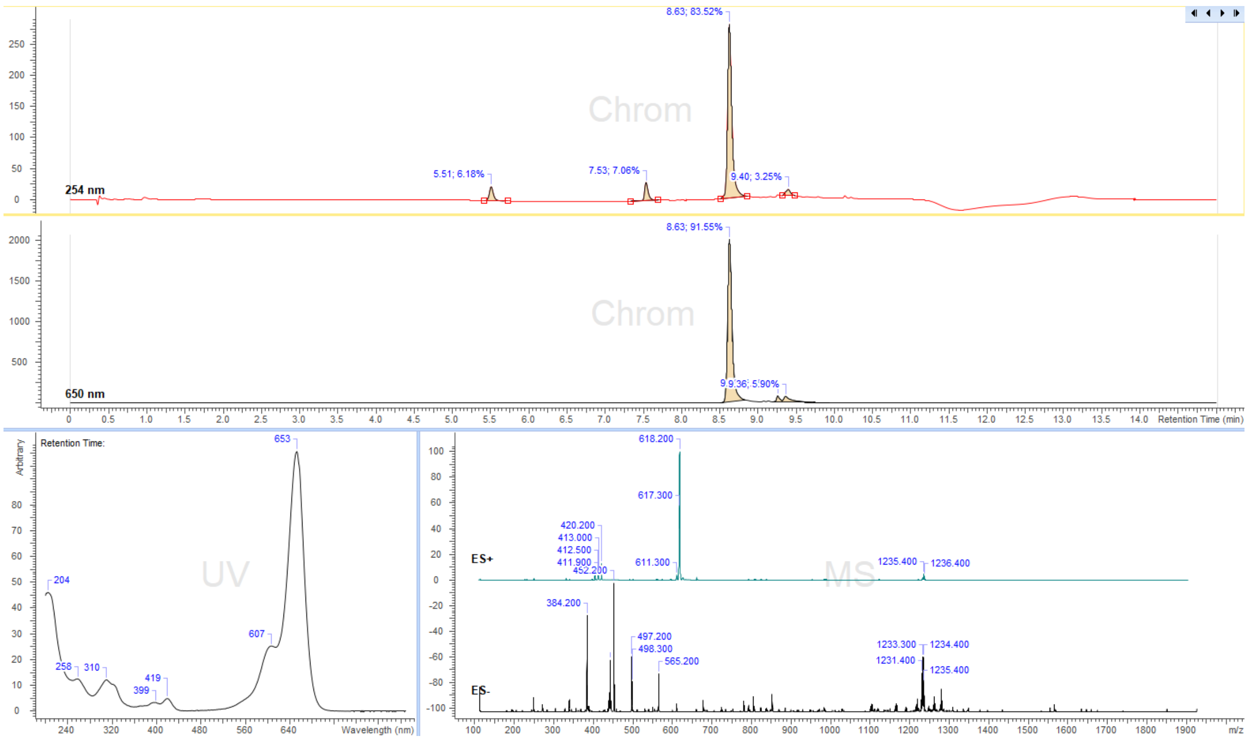


**Figure S44.** LC/MS analysis profile of compound **42.**

Cofactor **43**:

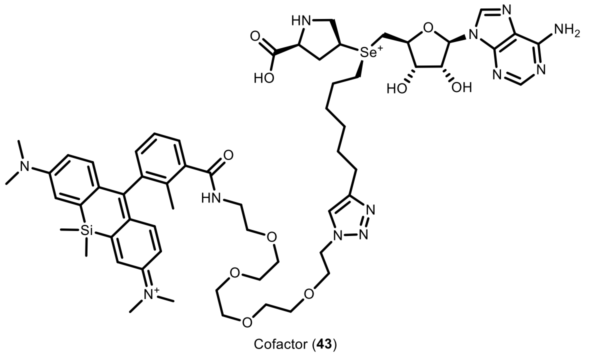

Cofactor (**43**)

A cofactor **36b** solution (250 µL of 10.7 mM in 10 mM pH 3.5 ammonium formate buffer) was mixed with a solution of 4-MeSiR-PEG$_4$-N$_3$ (**SI-8**) (500 µL of 10 mM in 10mM ammonium formate buffer pH = 3.5). Fresh solutions of $CuSO_4{\cdot}5H_2O$ (10 mg in 100 µL of AF buffer), ascorbic acid (10 mg in 100 µL of AF buffer) and (BimC$_4$A)$_3$ (1 mg in 10 µL of water) were prepared. 10 µL of the $CuSO_4$ solution were mixed with 5 µL of the (BimC$_4$A)$_3$ solution and sonicated for 1 minute, then 30 µL of the ascorbic acid solution were added (an immediate color change was observed) and again sonicated for 1 minute (no $CuSO_4$ induced solvolysis of reactant **38b** or product **43** was observed). Then 30 µL of the obtained mixture were transferred to the reaction vial containing the solution of **38b** and **SI-8**. The reaction vial was sonicated for 2 min and then left at room temperature for 120 min. The progress of the reaction was monitored by LC/MS every 40 min. Once the conversion reached >90%, the reaction was diluted with water to a volume of 2 mL and the product was purified by preparative HPLC (preparative column: Pursuit 10 C18 10 µm, 250×50.0 mm, Agilent, flow rate: 150 mL/min, solvent A: MeOH, solvent B: $H_2O$ (ammonium formate buffer 10 mM, pH = 3.5), temperature 25 °C, gradient A:B - 2 min 30:70 isocratic, 2-20 min 30:70 to 100:0 gradient and 20-22min 100:0 isocratic). Collected fractions of pure product were concentrated and the concentrations of the obtained solutions were determined by their UV absorption ($\varepsilon_{640}$ = 150 000 L·mol$^{-1}$·cm$^{-1}$). 0.9 mL, 1.7 mM in 57% yield were obtained.

ESI-MS, positive mode: m/z = 620.3 $[M]^{2+}$. HRMS (ESI) calculated for $C_{60}H_{84}N_{12}O_{10}SeSi$ $[M]^{2+}$ 620.2683, found 620.2707.

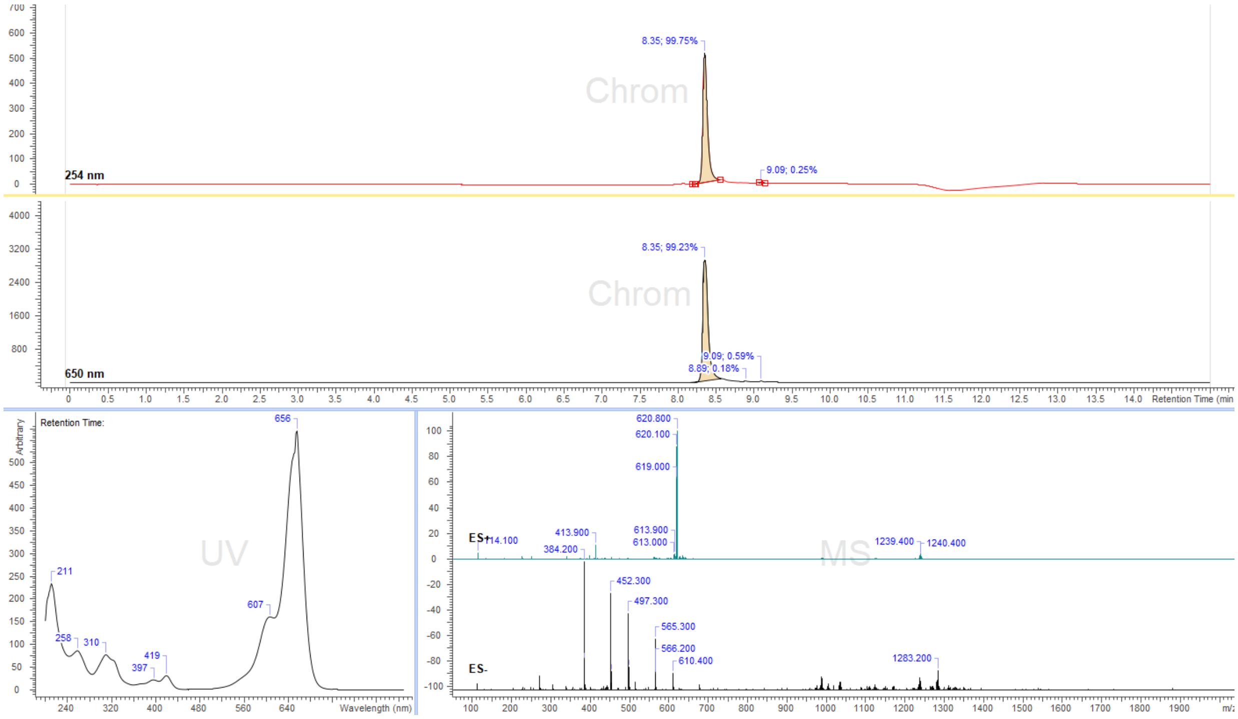


**Figure S45.** LC/MS analysis profile of compound **43.**

AdoHcyMeSiR (**44**)

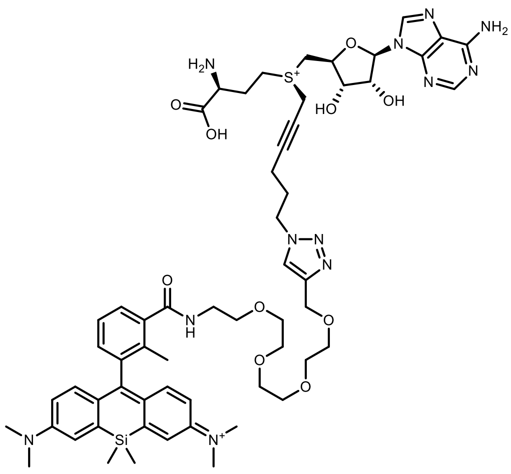

Cofactor (**44**)

A cofactor **AdoHcyN$_3$**[3] solution (500 µL of 6.7 mM in 10mM pH 3.5 ammonium formate buffer) was mixed with a solution of 4-MeSiR-PEG$_4$-Alkyne (**SI-9**) (500 µL of 10 mM in 10mM ammonium formate buffer pH = 3.5). Fresh solutions of $CuSO_4 \cdot 5H_2O$ (10 mg in 100 µL of AF buffer), ascorbic acid (10 mg in 100 µL of AF buffer) and $(BimC_4A)_3$ (1 mg in 10 µL of water) were prepared. 10 µL of the $CuSO_4$ solution were mixed with 5 µL of the $(BimC_4A)_3$ solution and sonicated for 1 minute, then 30 µL of the ascorbic acid solution were added (an immediate color change was observed) and again sonicated for 1 minute. Then 20 µL of the obtained mixture were transferred to the reaction vial containing the solution of **AdoHcyN$_3$** and **SI-9**. The reaction vial was sonicated for 2 min and then left at room temperature for 90 min. The progress of the reaction was monitored by LC/MS every 20 min. Once the conversion reached >90%, the reaction was diluted with water to a volume of 2 mL and the product was purified by preparative HPLC (preparative column: Pursuit 10 C18 10 µm, 250×50.0 mm, Agilent, flow rate: 150 mL/min, solvent A: MeOH, solvent B: $H_2O$ (ammonium formate buffer 10 mM, pH = 3.5), temperature 25 °C, gradient A:B - 2 min 30:70 isocratic, 2-20 min 30:70 to 100:0 gradient and 20-22 min 100:0 isocratic). Collected fractions of pure product were concentrated and the concentrations of the obtained solutions were determined by their UV absorption ($\varepsilon_{640}$ = 150 000 $L \cdot mol^{-1} \cdot cm^{-1}$). 0.8 mL, 2.8 mM in 67% yield were obtained.

ESI-MS, positive mode: m/z = 581.3 $[M]^{2+}$.

HRMS (ESI) calculated for $C_{58}H_{78}N_{12}O_{10}SSi$ $[M]^{2+}$ 581.2721, found 581.2730.

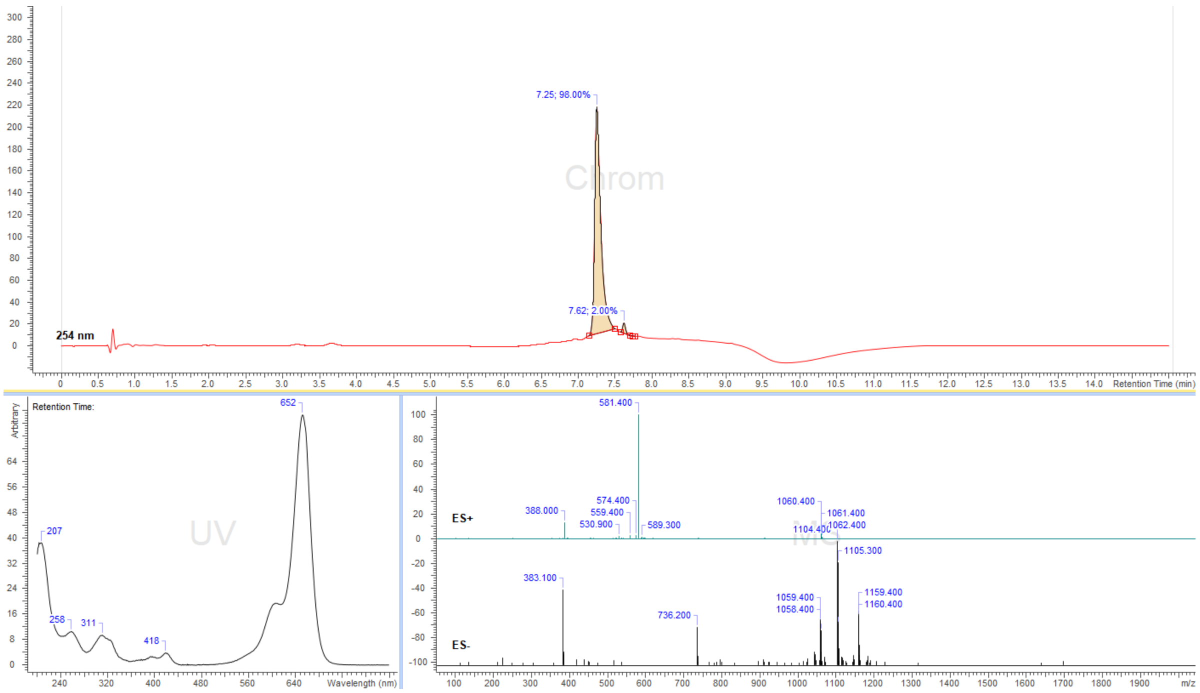


**Figure S46.** LC/MS analysis profile of compound **44.**

**Hex-2-yn-1-yl methanesulfonate** (**SI-1**):

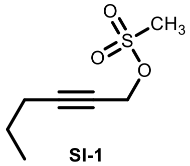
SI-1

To a solution of 2-hexyn-1-ol (1.0 g, 10.2 mmol) in dry DCM (25 mL) DIPEA (2 mL, 12.2 mmol) was added and the mixture was cooled in an ice/water bath. Then MsCl (0.8 mL, 10.2 mmol) was added dropwise over a period of 10 min. Once the addition was complete, the mixture was removed from the ice/water bath and stirred at room temperature for additional 2h. Then the mixture was washed with 1M HCl (1x25 mL), water (1x25mL) and brine (1x25mL). The solvent was evaporated, the residue was deposited on celite and the product was purified by flash column chromatography (Interchim Puriflash 30 µm Hexane:DCM 20% to 100% gradient, stained with $KMnO_4$ staining solution) to afford 1.6 g of a transparent liquid in 89% yield.

$^{1}$H NMR (400 MHz, chloroform-*d*) δ 4.86 (t, *J* = 2.2 Hz, 2H), 3.12 (s, 3H), 2.24 (tt, *J* = 7.0, 2.2 Hz, 2H), 1.55 (p, *J* = 7.3 Hz, 2H), 1.00 (t, *J* = 7.3 Hz, 3H).

$^{13}$C NMR (101 MHz, chloroform-*d*) δ 91.0, 72.5, 58.7, 39.1, 21.8, 20.8, 13.5.

**Octa-2,7-diyn-1-yl methanesulfonate** (**SI-2**):

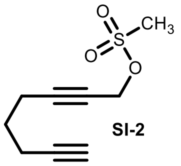
SI-2

To a solution of octa-2,7-diyn-1-ol (1.0 g, 8.2 mmol) in dry DCM (25 mL) DIPEA (2 mL, 12.2 mmol) was added and the mixture was cooled in an ice/water bath. Then MsCl (0.64 mL, 8.2 mmol) was added dropwise over a period of 10 min. Once the addition was complete, the mixture was removed from the ice/water bath and stirred at room temperature for additional 2h. Then the mixture was washed with 1M HCl (1x25 mL), water (1x25mL) and brine (1x25mL). The solvent was evaporated, the residue was deposited on celite and the product was purified by flash column chromatography (Interchim Puriflash 30 µm DCM:EtOAc 0% to 20% gradient, stained with $KMnO_4$ staining solution) to afford 1.06 g of a yellowish transparent liquid in 65% yield.

$^{1}$H NMR (400 MHz, chloroform-*d*) δ 4.83 (t, *J* = 2.2 Hz, 2H), 3.10 (s, 3H), 2.40 (tt, *J* = 7.0, 2.2 Hz, 2H), 2.30 (td, *J* = 6.9, 2.6 Hz, 2H), 1.97 (t, *J* = 2.6 Hz, 1H), 1.74 (p, *J* = 7.0 Hz, 2H).

$^{13}$C NMR (101 MHz, chloroform-*d*) δ 89.8, 83.0, 73.1, 69.4, 58.4, 39.1, 27.0, 17.8, 17.6.

**Prop-2-yn-1-yl methanesulfonate** (**SI-3**):

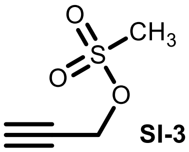
SI-3

To a solution of 2-propyn-1-ol (0.5 g, 8.92 mmol) in dry DCM (25 mL) DIPEA (2 mL, 12.2 mmol) was added and the mixture was cooled in an ice/water bath. Then MsCl (0.69 mL, 8.92 mmol) was added dropwise over a period of 10 min. Once the addition was complete, the mixture was removed from the ice/water bath and stirred at room temperature for additional 2h. Then the mixture was washed with 1M HCl (1x25 mL), water (1x25mL) and brine (1x25mL). The solvent was evaporated, the residue was deposited on celite and the product was purified by flash column chromatography (Interchim Puriflash 30 µm DCM:EtOAc 0% to 10% gradient, stained with $KMnO_4$ staining solution) to afford 0.9 g of a yellowish transparent liquid in 75% yield.

$^{1}$H NMR (400 MHz, chloroform-*d*) δ 4.85 (d, *J* = 2.5 Hz, 2H), 3.13 (s, 3H), 2.70 (t, *J* = 2.4 Hz, 1H).

$^{13}$C NMR (101 MHz, chloroform-*d*) δ 78.0, 75.9, 57.3, 39.2.

**Trifluoromethanesulfonate** (**SI-4**):

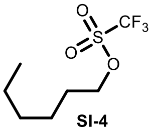


**SI-4**

To a solution of 1-hexanol (0.5 g, 4.9 mmol) in dry DCM (25 mL) DIPEA (2 mL, 12.2 mmol) was added and the mixture was cooled in an ice/water bath. Then $TfO_2$ (0.8 mL, 4.9 mmol) was added dropwise over a period of 10 min. Once the addition was complete, the mixture was removed from the ice/water bath and stirred at room temperature for additional 2h. Then the mixture was washed with 1M HCl (1x25 mL), water (1x25mL) and brine (1x25mL). The solvent was evaporated, the residue was deposited on celite and the product was purified by flash column chromatography (Interchim Puriflash 30 µm DCM:EtOAc 0% to 10% gradient, stained with $KMnO_4$ staining solution) to afford 0.97 g of a transparent liquid in 85% yield.

$^{1}$H NMR (400 MHz, chloroform-*d*) δ 4.54 (t, *J* = 6.6 Hz, 2H), 1.90 – 1.73 (m, 2H), 1.52 – 1.24 (m, 6H), 0.97 – 0.83 (m, 3H).

$^{13}$C NMR (101 MHz, chloroform-*d*) δ 118.8 (q, $^{1}J_{C\text{-}F}$ = 318 Hz), 77.9, 31.1, 29.3, 24.9, 22.5, 14.0.

**Oct-7-yn-1-yl trifluoromethanesulfonate** (**SI-5**):

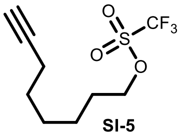


**SI-5**

To a solution of 7-Octyn-1-ol (0.5 g, 4.0 mmol) in dry DCM (25 mL) DIPEA (2 mL, 12.2 mmol) was added and the mixture was cooled in an ice/water bath. Then $TfO_2$ (0.66 mL, 4.0 mmol) was added dropwise over a period of 5 min. Once the addition was complete, the mixture was removed from the ice/water bath and stirred at room temperature for additional 2h. Then the mixture was washed with 1M HCl (1x25 mL), water (1x25mL) and brine (1x25mL). The solvent was evaporated, the residue was deposited on celite and the product was purified by flash column chromatography (Interchim Puriflash 30 µm DCM:EtOAc 0% to 10% gradient, stained with $KMnO_4$ staining solution) to afford 0.63 g of transparent liquid in 61% yield.

$^{1}$H NMR (400 MHz, chloroform-*d*) δ 4.54 (t, *J* = 6.4 Hz, 2H), 2.21 (td, *J* = 6.8, 2.6 Hz, 2H), 1.95 (t, *J* = 2.7 Hz, 1H), 1.88 – 1.81 (m, 2H), 1.57 – 1.44 (m, 6H).

$^{13}$C NMR (101 MHz, chloroform-*d*) δ 118.8 (q, $^{1}J_{C-F}$ = 318 Hz), 84.3, 77.6, 68.6, 29.3, 28.2, 28.0, 24.8, 18.4.

**2-(3-Bromo-2-methylphenyl)-4,4-dimethyl-4,5-dihydrooxazole** (**SI-6**):

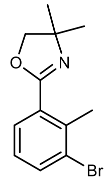

**SI-6**

3-bromo-2-methylbenzoic acid (3 g, 13.9 mmol) was suspended in $SOCl_2$ (15 mL) and 3 drops of DMF were added. The resulting mixture was refluxed for 3 hours and thionyl chloride was evaporated. The residue was dissolved in DCM (40 mL) and was added dropwise to the mixture of 2-amino-2-methyl-1-propanol (2 mL, 20.9 mmol) and DIPEA (3 mL, 17.2 mmol) in DCM (50 mL) whilst cooled in ice bath. Once the addition was complete, the mixture was stirred at room temperature overnight. Then the solvent was evaporated and a solution of $NaHCO_3$ (30mL) was added and the product was extracted with EtOAc (3x30mL). The combined organic layers were washed with water, brine and dried over $Na_2SO_4$. The solvent was evaporated and the residue was suspended in DCM (50mL), then DBU (3.1 mL, 21 mmol) was added and the mixture became clear. The obtained mixture was cooled in an ice bath and NfF (3.3 mL, 18.2 mmol) was added dropwise over a period of 5 min. Once the addition was complete, the ice bath was removed and the reaction mixture was stirred at room temperature for 2h, quenched with $NaHCO_3$ (50 mL) and extracted with DCM (50 mL), washed with water and brine and was dried over $Na_2SO_4$. The solvent was evaporated and the residue was deposited on celite and purified by flash column chromatography (Redisep 80g, DCM:EtOAc 5% to 50% gradient) to afford 3.3 g of **SI-6**in a 88% yield.

$^{1}$H NMR (400 MHz, chloroform-*d*) δ 7.64 – 7.61 (m, 1H), 7.61 – 7.59 (m, 1H), 7.06 (td, *J* = 7.9, 0.7 Hz, 1H), 4.09 (s, 2H), 2.60 (s, 3H), 1.40 (s, 6H).

$^{13}$C NMR (101 MHz, chloroform-*d*) δ 162.5, 137.9, 134.7, 130.2, 129.0, 126.8, 79.1, 68.2, 28.5, 20.9.

ESI-MS, positive mode: m/z = 268.0 $[M+H]^+$. HRMS (ESI) calculated for $C_{12}H_{15}BrNO$ $[M+H]^+$ 268.0332, found 268.0337.

**4-MeSiR-COOH** (**SI-7**):

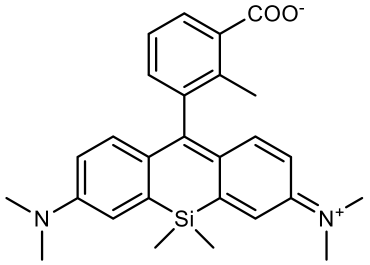


**SI-7**

In a vacuum dried 100 mL flask flushed with argon compound **SI-6** (1.0g, 3.73 mmol) was dissolved in dry THF (20 mL). The solution was cooled to -78°C in a dry ice/acetone bath and *s*-BuLi (3.2 mL of 1.4M, 4.47 mmol) was added by a syringe in 10 minutes. After addition was complete the reaction was stirred for 1h at -78°C and the solution of silaxanthone [4] (0.5 g, 1.54 mmol) in dry THF (40 mL) was slowly introduced via syringe. The cooling bath was removed and the reaction mixture was left to warm up to room temperature and stirred for additional 2h. Then glacial acetic acid (1 mL) was added and the mixture turned blue. Solvents were evaporated on a rotary evaporator and the crude mixture was dissolved in 6N HCl (30 mL) and stirred at 80°C overnight. Then it was cooled to room temperature, poured into a beaker and the pH was carefully adjusted to 2-3 by slow addition of a saturated $NaHCO_3$ solution. The product was extracted with DCM:MeOH (5x40 mL, 85:15 mixture), dried over $Na_2SO_4$ and purified by preparative HPLC (preparative column: Eurospher II 100-5 C18 5 µm, 250×20.0 mm, Article No.: 25PE181E2J, Knauer, flow rate: 25 mL/min, solvent A: acetonitrile, solvent B: $H_2O$ + 0.1% v/v TFA; temperature 25 °C, gradient A:B - 5 min 30:70 isocratic, 5-30 min 30:70 to 100:0 gradient) and lyophilized from an acetonitrile/water mixture to obtain 430mg of 4-MeSiR-COOH (**SI-7**) in 65% yield.

$^1$H NMR (400 MHz, Methanol-$d_4$) δ 8.04 (dd, *J* = 7.8, 1.5 Hz, 1H), 7.47 (td, *J* = 7.7, 0.7 Hz, 1H), 7.37 (d, *J* = 2.9 Hz, 2H), 7.29 (dd, *J* = 7.6, 1.6 Hz, 1H), 7.05 (d, *J* = 9.7 Hz, 2H), 6.79 (dd, *J* = 9.7, 2.9 Hz, 2H), 3.35 (s, 12H), 2.26 (s, 3H), 0.62 (s, 3H), 0.61 (s, 3H).

$^{13}$C NMR (101 MHz, Methanol-d4) δ 170.8, 169.9, 155.8, 149.5, 142.2, 141.7, 137.9, 133.5, 133.4, 131.9, 128.6, 126.7, 122.3, 115.4, 40.9, 18.1, -1.1, -1.3.

ESI-MS, positive mode: m/z = 443.2 $[M]^+$. HRMS (ESI) calculated for $C_{27}H_{31}N_2O_2Si$ $[M]^+$ 443.2149, found 443.2160.

**4-MeSiR-PEG$_4$-N$_3$** (**SI-8**):

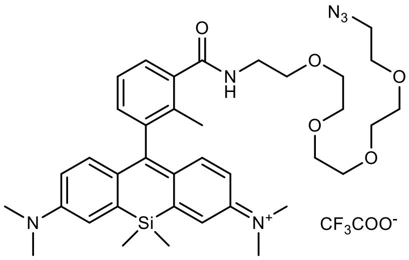


**SI-8**

4-MeSiR-COOH (**SI-7**) (25 mg, 0.056 mmol) was dissolved in dry DMSO (300 µL) and DIPEA (20 µL) and then a solution of HATU (23 mg, 0.06 mmol ) in DMSO (300 µL) was added and the reaction mixture was stirred at room temperature for 5 min. Then a solution of $NH_2$-PEG$_4$-$N_3$ (21 mg, 0.08 mmol) in DMSO (200 µL) was added and the mixture was stirred for 2h at room temperature. Then the reaction was quenched by adding formic acid (50 µL), diluted with water to a volume of 2 mL and the product was purified by preparative HPLC (preparative column: Pursuit 10 C18 10 µm, 250×50.0 mm, Agilent, flow rate: 150 mL/min, solvent A: MeOH, solvent B: $H_2O$ +0.1% TFA; temperature 25 °C, gradient A:B - 2 min 30:70 isocratic, 2-20 min 30:70 to 100:0 gradient and 20-22min 100:0 isocratic). Fractions containing the product were evaporated and dissolved in a 1:2 mixture of MeCN/water and lyophilized. 26 mg (68% yield) of a dark blue solid were obtained.

$^{1}$H NMR (400 MHz, Methanol-$d_4$) δ 7.55 (dd, $J$ = 7.7, 1.4 Hz, 1H), 7.45 (td, $J$ = 7.6, 0.7 Hz, 1H), 7.37 (d, $J$ = 2.8 Hz, 2H), 7.21 (dd, $J$ = 7.6, 1.4 Hz, 1H), 7.11 (d, $J$ = 9.6 Hz, 2H), 6.77 (dd, $J$ = 9.6, 2.8 Hz, 2H), 3.70 – 3.58 (m, 18H), 3.36 (s, 12H), 3.34 – 3.32 (m, 2H), 2.07 (s, 3H), 0.62 (s, 3H), 0.61 (s, 3H).

$^{13}$C NMR (101 MHz, Methanol-$d_4$) δ 172.6, 169.9, 155.8, 149.5, 142.3, 141.2, 139.5, 134.3, 131.5, 128.52, 128.49, 126.8, 122.2, 115.2, 71.61, 71.58, 71.54, 71.52, 71.51, 71.2, 71.1, 70.4, 51.8, 40.9, 40.8, 17.0, -1.1, -1.3.

ESI-MS, positive mode: m/z = 687.4 $[M]^+$. HRMS (ESI) calculated for $C_{37}H_{51}N_6O_5Si$ $[M]^+$ 687.3685, found 687.3687.

**MeSiR-PEG4-alkyne** (**SI-9**)

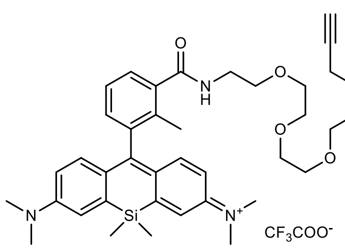

**SI-9**

4-MeSiR-COOH (**SI-7**) **(**25 mg, 0.056 mmol) was dissolved in dry DMSO (300 µL) and DIPEA (20 µL) and then a solution of HATU (23 mg, 0.06 mmol ) in DMSO (300 µL) was added and the reaction mixture was stirred at room temperature for 5 min. Then solution of $NH_2$-$PEG_4$-Alkyne (19 mg, 0.08 mmol) in DMSO (200 µL) was added and the mixture was stirred for 2h at room temperature. Then the reaction was quenched by adding formic acid (50 µL), diluted with water to 2 mL volume and the product was purified by preparative HPLC (preparative column: Pursuit 10 C18 10 µm, 250×50.0 mm, Agilent, flow rate: 150 mL/min, solvent A: MeOH, solvent B: $H_2O$ +0.1% TFA; temperature 25 °C, gradient A:B - 2 min 30:70 isocratic, 2-20 min 30:70 to 100:0 gradient and 20-22 min 100:0 isocratic). Fractions containing the product were evaporated, dissolved in a MeCN/water 1:2 mixture and lyophilized. 30 mg (70% yield) of a dark blue solid were obtained.

$^{1}$H NMR (400 MHz, Methanol-d4) δ 7.55 (d, J = 6.9 Hz, 1H), 7.45 (t, J = 7.6 Hz, 1H), 7.37 (d, J = 2.8 Hz, 2H), 7.21 (d, J = 7.2 Hz, 1H), 7.11 (d, J = 9.6 Hz, 2H), 6.77 (dd, J = 9.6, 2.8 Hz, 2H), 4.15 (d, J = 2.4 Hz, 2H), 3.70 – 3.57 (m, 16H), 3.36 (s, 12H), 2.84 (t, J = 2.3 Hz, 1H), 2.07 (s, 3H), 0.62 (s, 3H), 0.61 (s, 3H).

$^{13}C$ NMR (101 MHz, $CD_3OD$) δ 172.6, 169.9, 155.8, 149.5, 142.3, 141.2, 139.5, 134.3, 131.5, 128.53, 128.49, 126.8, 122.2, 115.2, 80.6, 76.0, 71.62, 71.48, 71.46, 71.31, 71.25, 70.4, 70.1, 59.0, 40.94, 40.86, 17.0, -1.1, -1.3.

ESI-MS, positive mode: m/z = 656.4 $[M]^+$.

HRMS (ESI) calculated for $C_{38}H_{50}N_3O_5Si$ $[M]^+$ 656.3539, found 656.3514.

**SeAdoHcy** (**SI-10**)

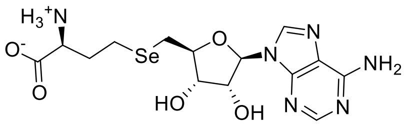

SeAdoHcy (**SI-10**)

15 mL of dry $NH_3$ gas was condensed in a 25 mL argon flushed flask equipped with a dry ice condenser and a MeCN/dry ice cooling bath. Selenomethionine (327 mg, 1.67 mmol) was added to liquid $NH_3$ and stirred until it dissolved. Then three small pieces of sodium (114 mg, 4.95 mmol) were added carefully. The reaction mixture colour changed to dark blue and after 30 min the reaction was quenched with $NH_4Cl$ (250 mg, 4.67 mmol). The reaction mixture was left to slowly warm to room temperature and $NH_3$ was allowed to slowly evaporate. The remaining solid was suspended in 5 mL of 1M NaOH solution and 5′-Tosyl adenosine (630 mg, 1.5 mmol) was added to the mixture, which was then stirred at 80 °C for 3 hours. Then the mixture was cooled and 1 mL of glacial acetic acid was added, water was evaporated and the remaining solid was dissolved in 40 mL of hot MeOH and the insoluble material was filtered. The filtrate was concentrated to a volume of 8-10 mL and 20 mL of *i*-ProH were added, the product precipitated and was filtered and dried. 436 mg of SeAdoHcy (**SI-10**) were obtained in 60% yield.

$^1H$ NMR (400 MHz, Methanol-d4) δ 8.50 (s, 1H), 8.40 (s, 1H), 6.06 (d, J = 4.8 Hz, 1H), 4.74 (t, J = 5.0 Hz, 1H), 4.33 – 4.25 (m, 2H), 4.10 – 4.03 (m, 1H), 3.10 – 2.97 (m, 2H), 2.81 – 2.69 (m, 2H), 2.36 – 2.23 (m, 1H), 2.21 – 2.10 (m, 1H).

$^{13}C$ NMR (101 MHz, Methanol-d4) δ 171.4, 152.5, 150.1, 146.1, 143.9, 120.7, 90.6, 86.1, 75.3, 74.7, 53.7, 32.5, 26.7, 19.9.

ESI-MS, positive mode: m/z = 433.1 $[M]^+$.

HRMS (ESI) calculated for $C_{14}H_{20}N_6O_5Se$ $[M]^+$ 433.0734, found 433.0741.

#### Copies of NMR spectra

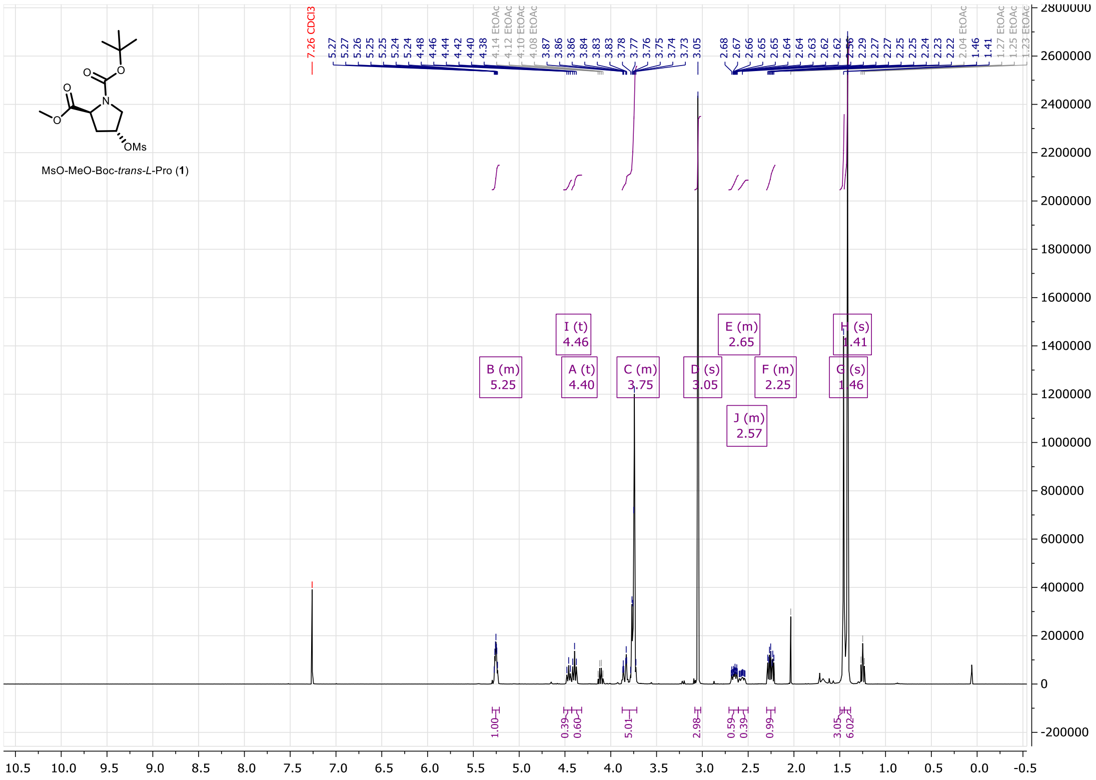

**Figure S47.** $^{1}H$ NMR of compound **1** (400MHz, $CDCl_3$).

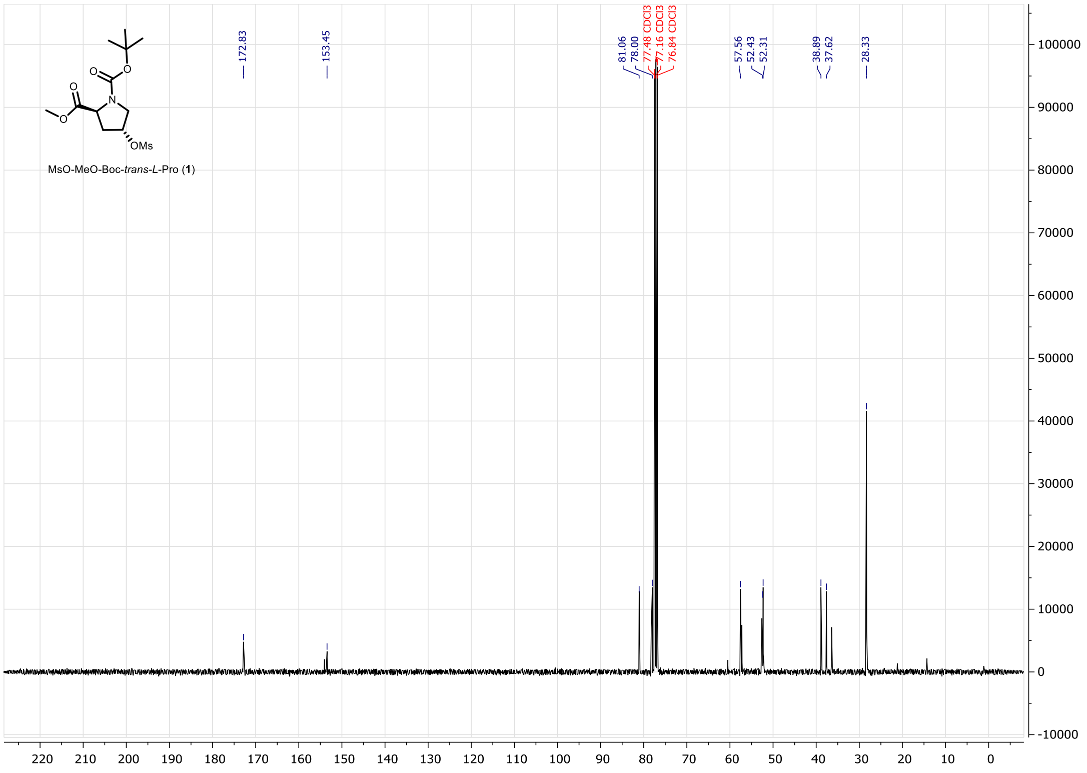

**Figure S48.** $^{13}C$ NMR of compound **1** (100MHz, $CDCl_3$).

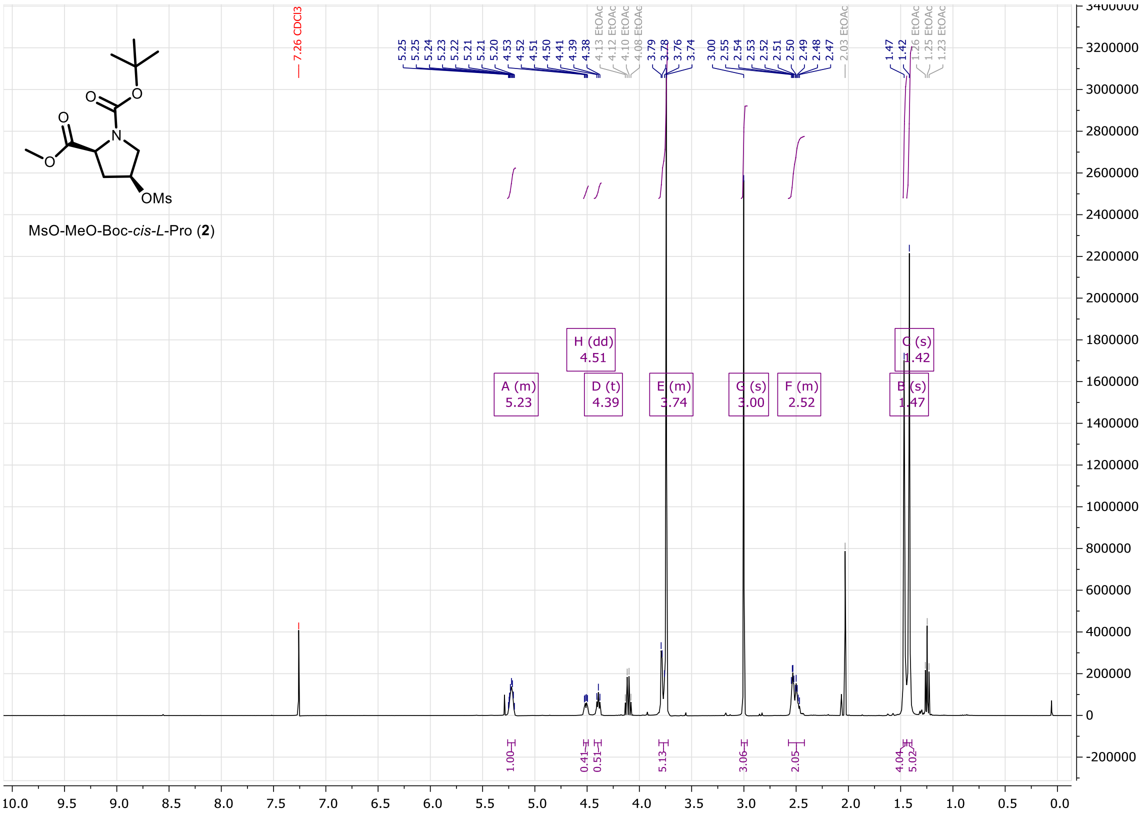


**Figure S49.** $^{1}$H NMR of compound **2** (400MHz, $CDCl_3$).

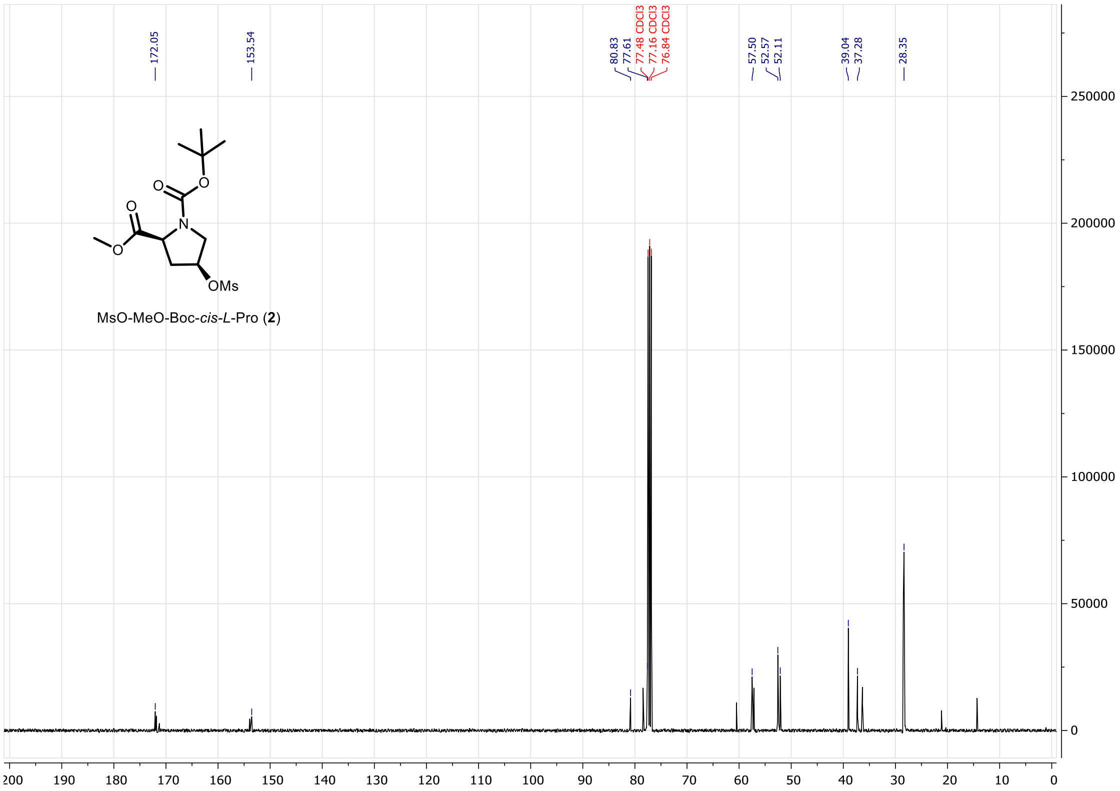

**Figure S50.** $^{13}$C NMR of compound **2** (100MHz, $CDCl_3$).

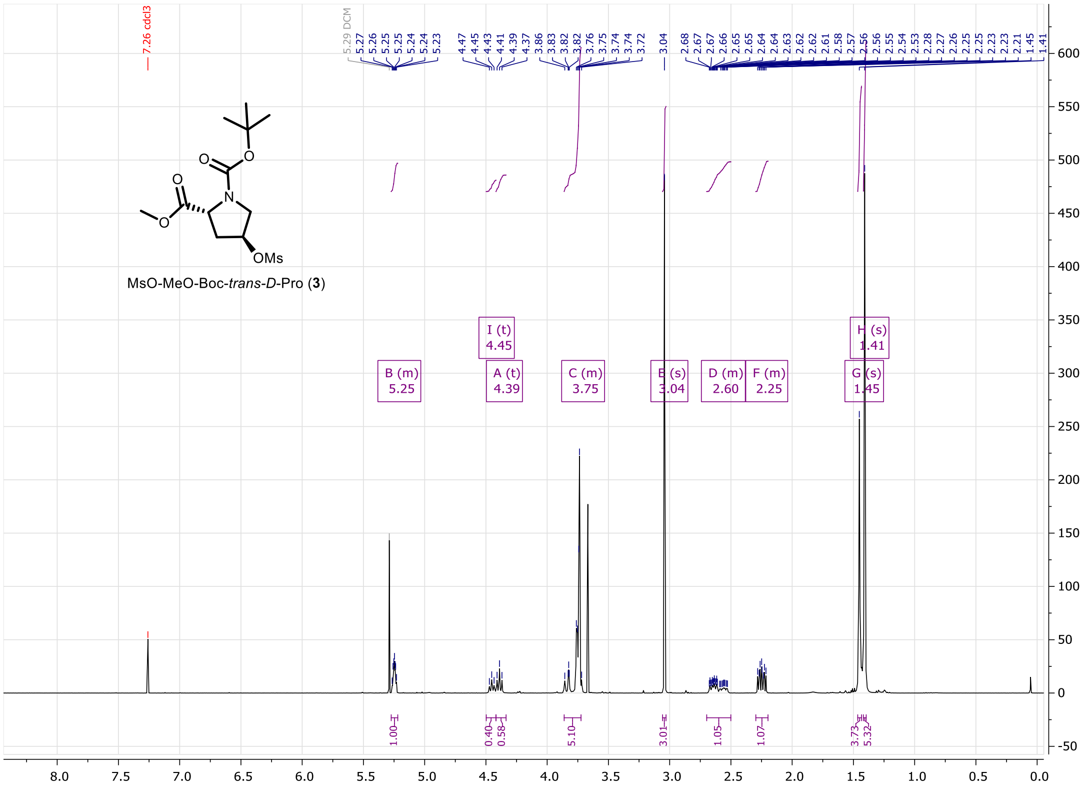


**Figure S51.** $^{1}$H NMR of compound **3** (400MHz, $CDCl_3$).

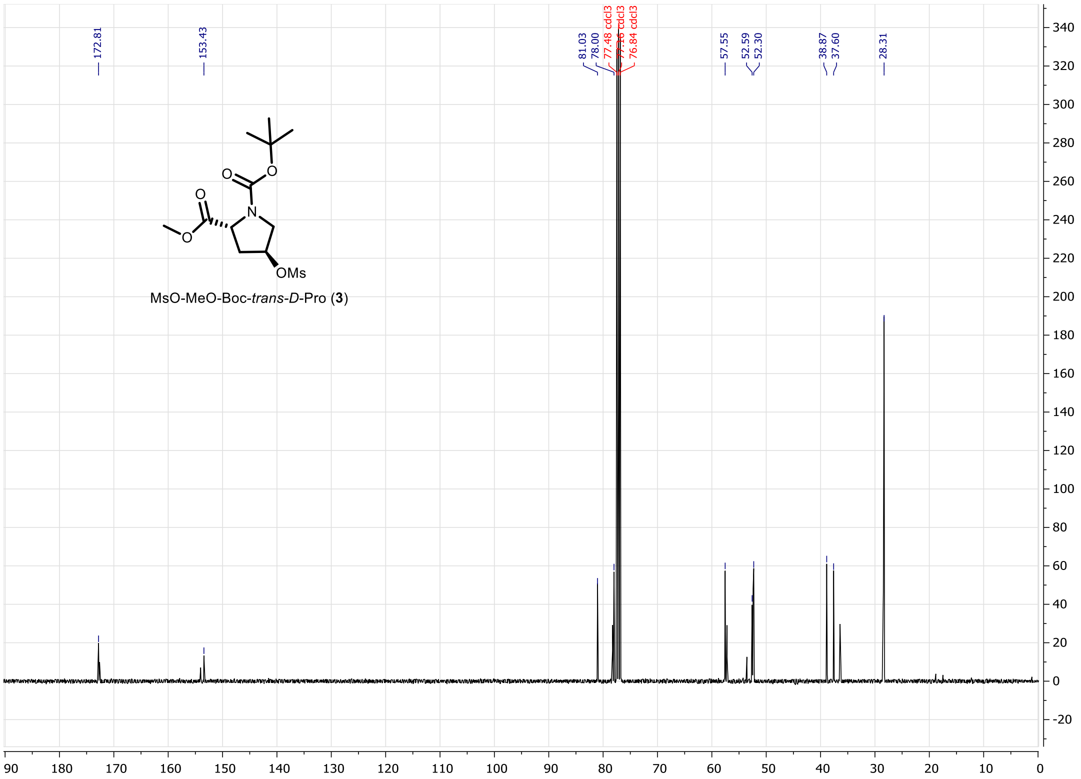


**Figure S52.** $^{13}$C NMR of compound **3** (100MHz, $CDCl_3$).

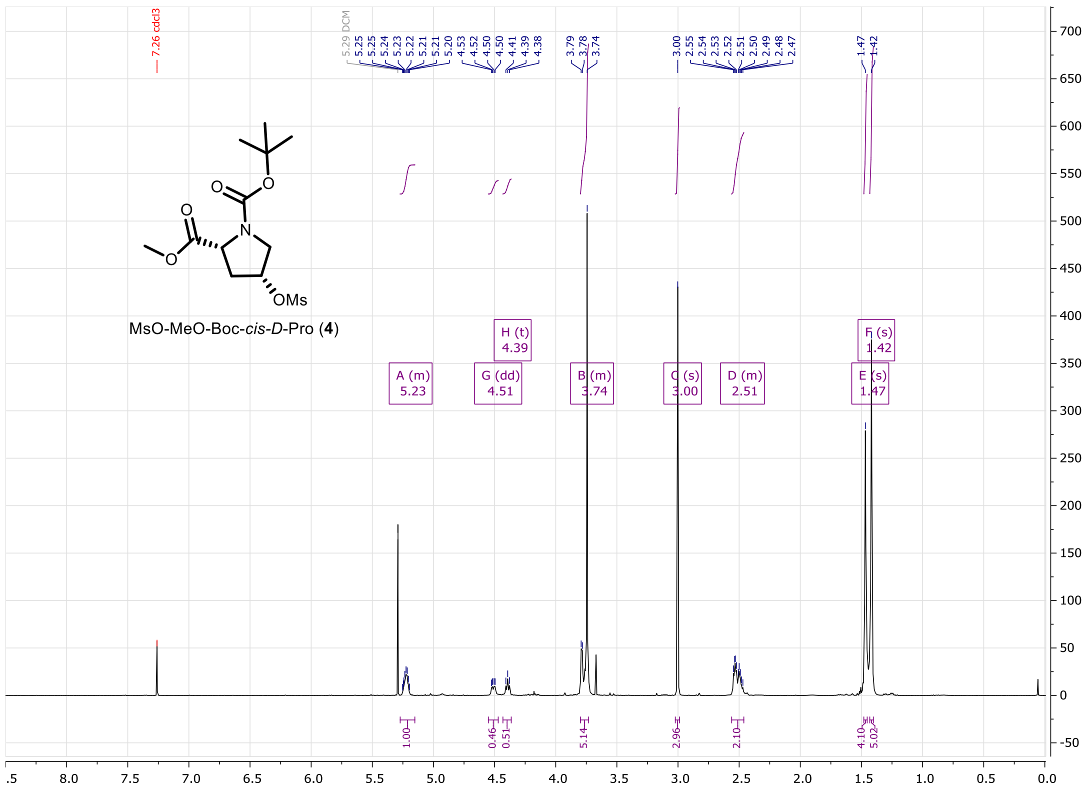

**Figure S53.** $^{1}$H NMR of compound **4** (400MHz, $CDCl_3$).

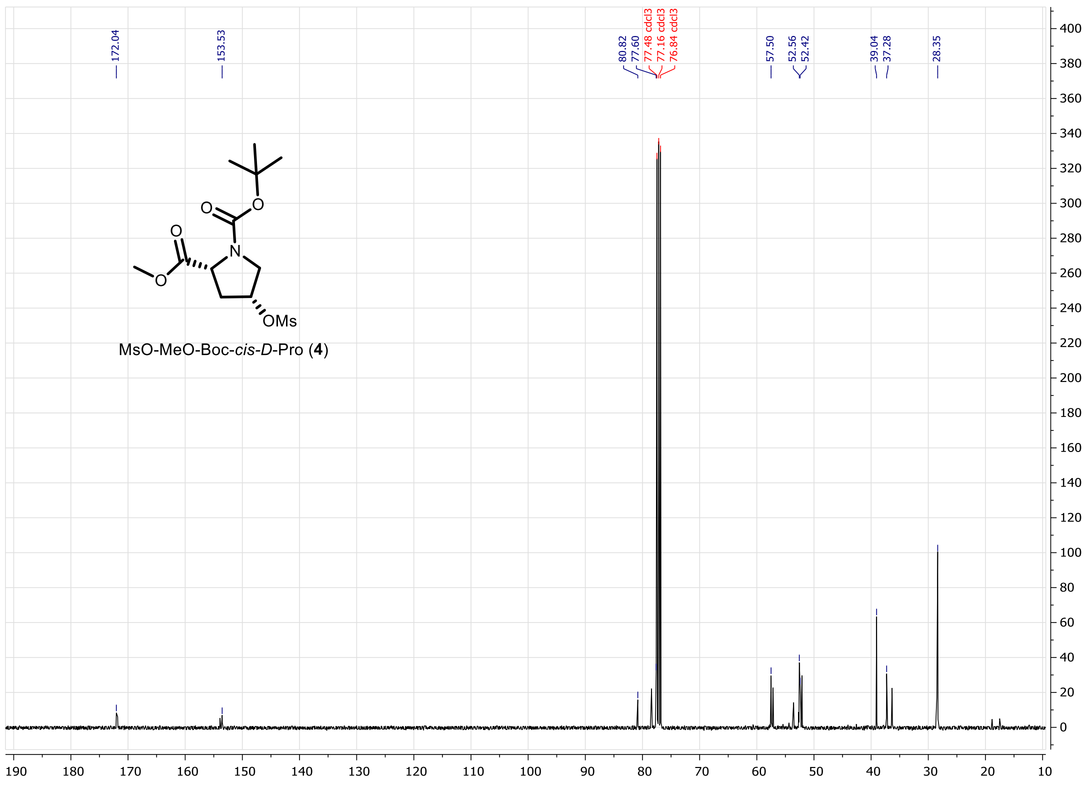

**Figure S54.** $^{13}$C NMR of compound **4** (100MHz, $CDCl_3$).

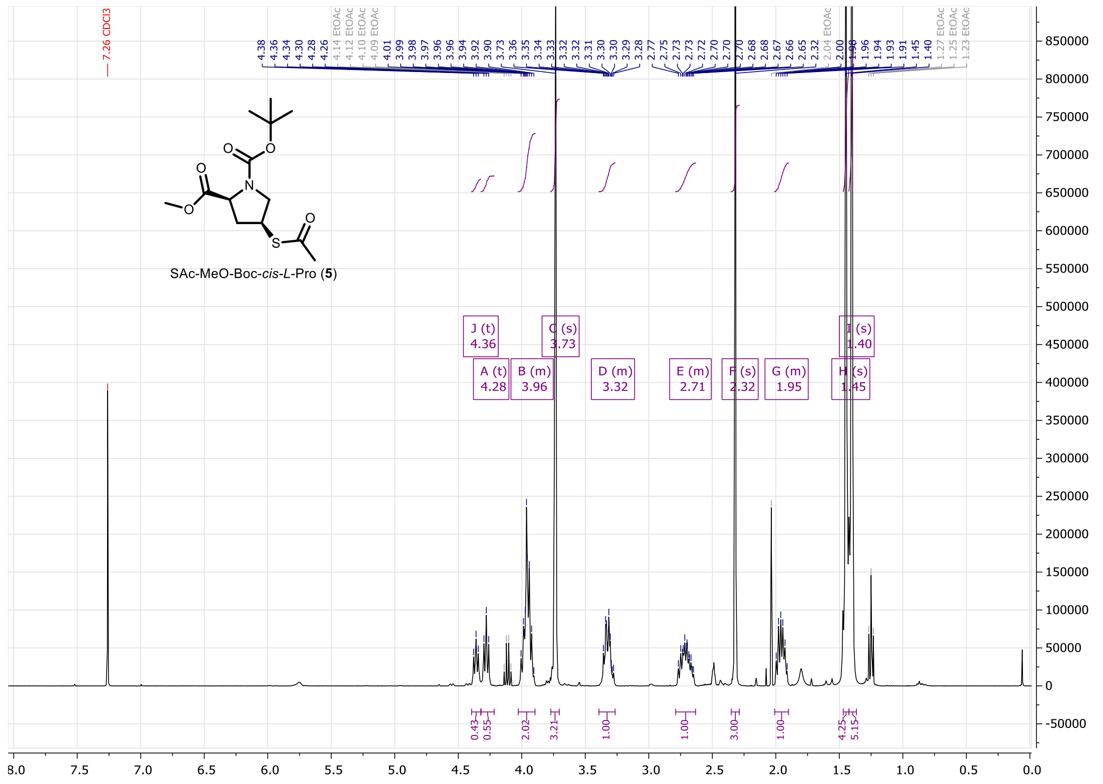

**Figure S55.** $^{1}$H NMR of compound **5** (400MHz, $CDCl_3$).

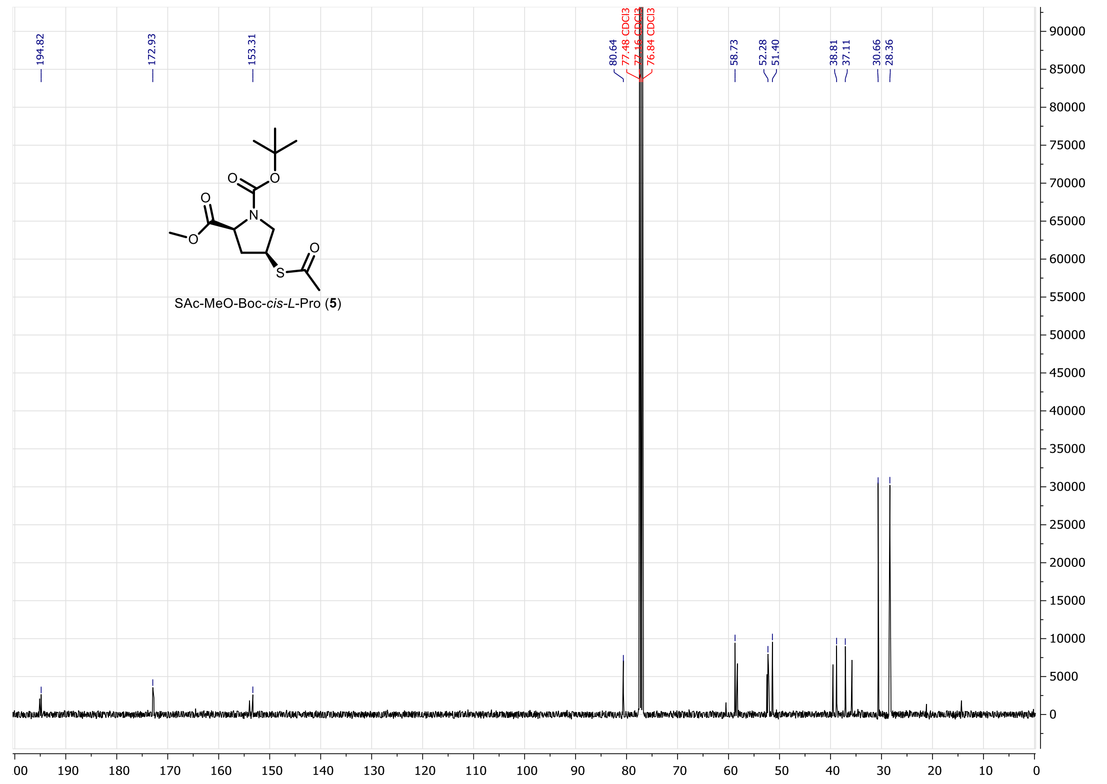

**Figure S56.** $^{13}$C NMR of compound **5** (100MHz, $CDCl_3$).

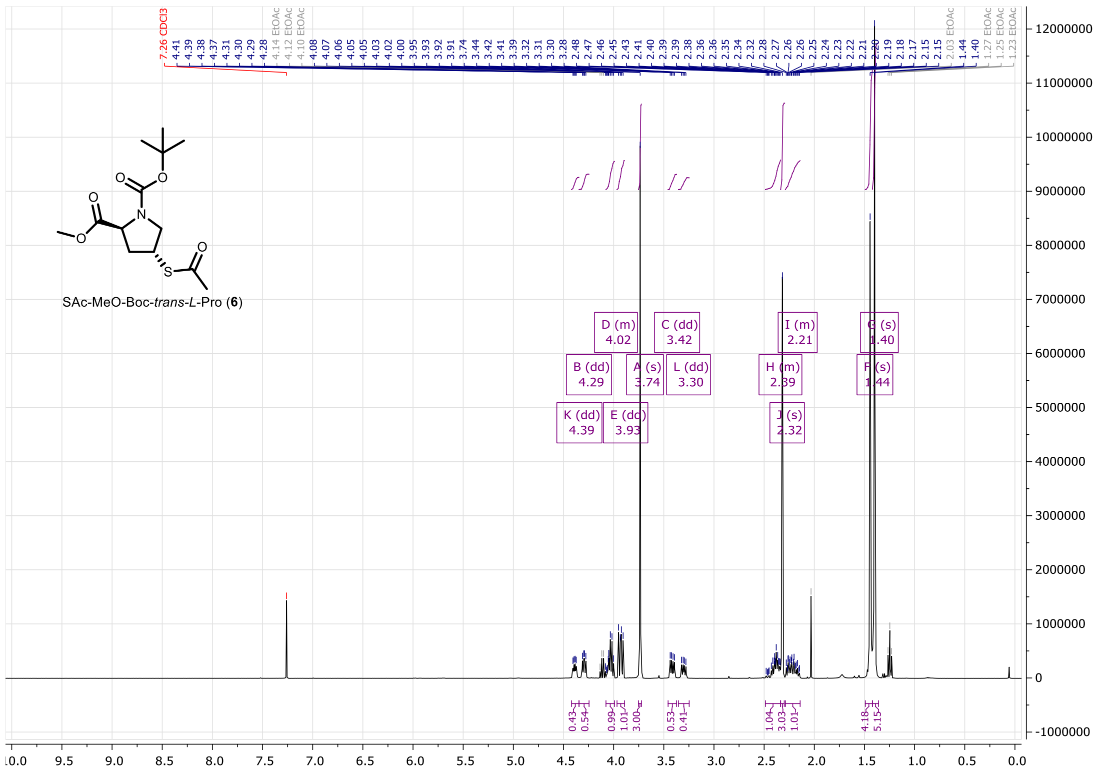

**Figure S57.** $^{1}$H NMR of compound **6** (400MHz, $CDCl_3$).

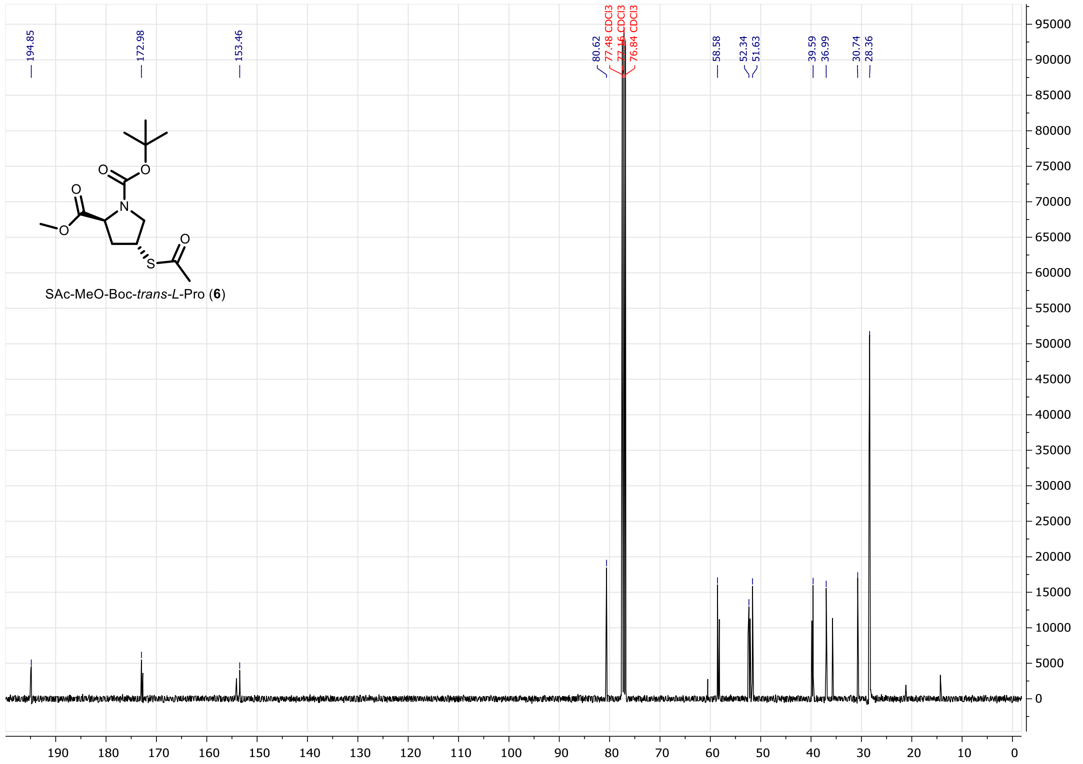

**Figure S58.** $^{13}$C NMR of compound **6** (100MHz, $CDCl_3$).

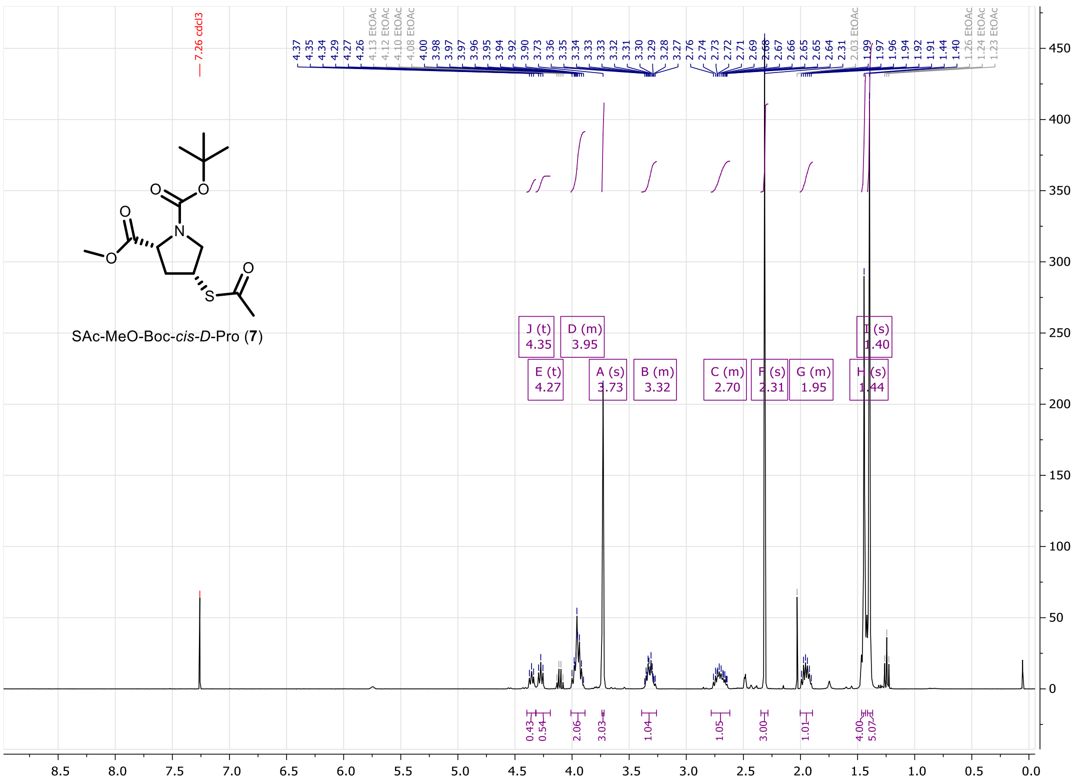


**Figure S59.** $^{1}$H NMR of compound **7** (400MHz, $CDCl_3$).

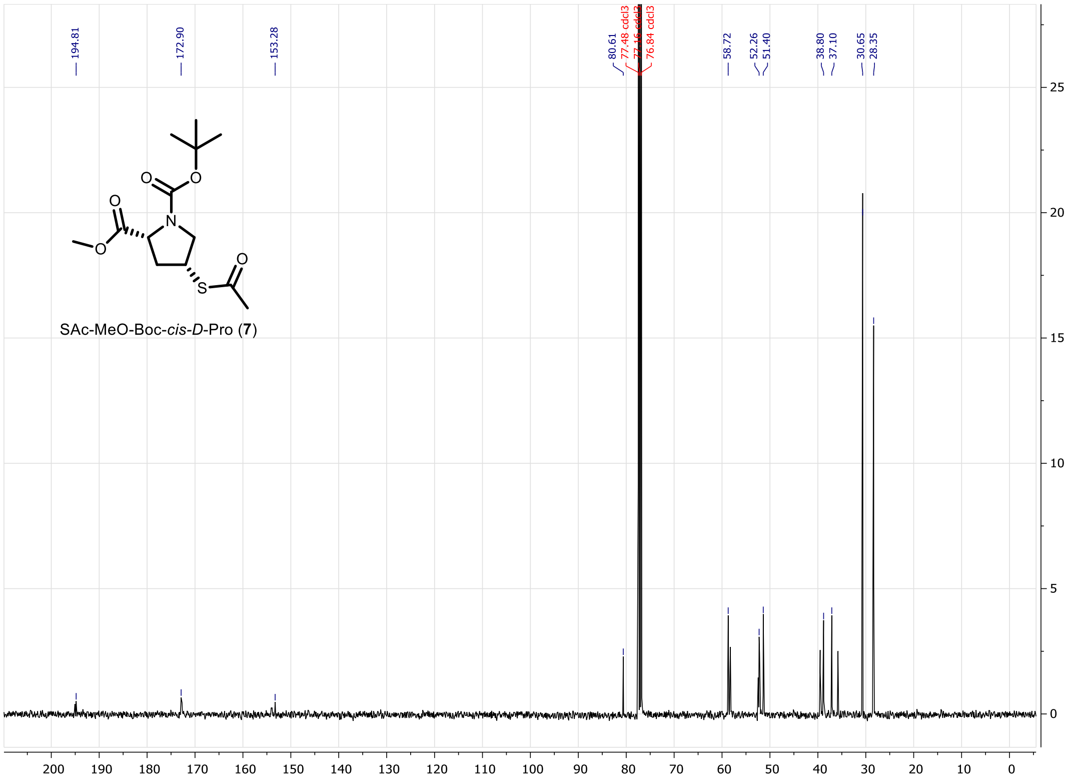


**Figure S60.** $^{13}$C NMR of compound **7** (100MHz, $CDCl_3$).

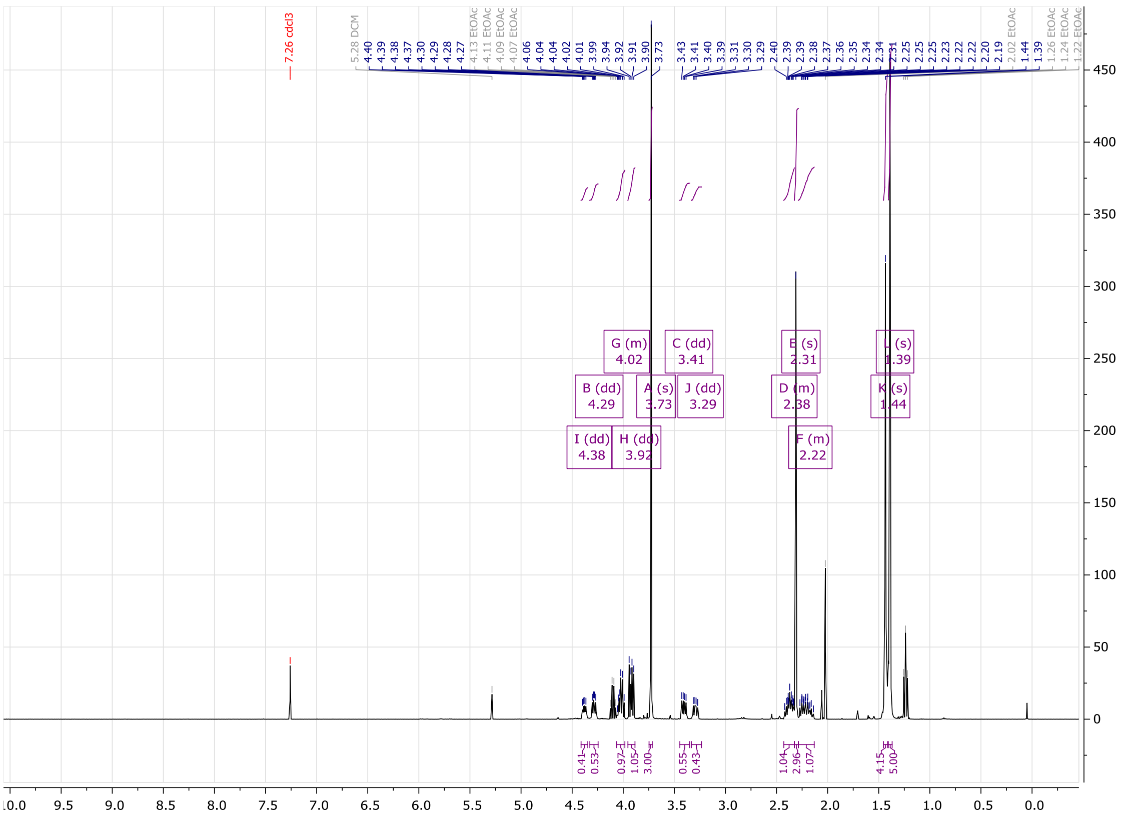

**Figure S61.** $^{1}H$ NMR of compound **8** (400MHz, $CDCl_3$).

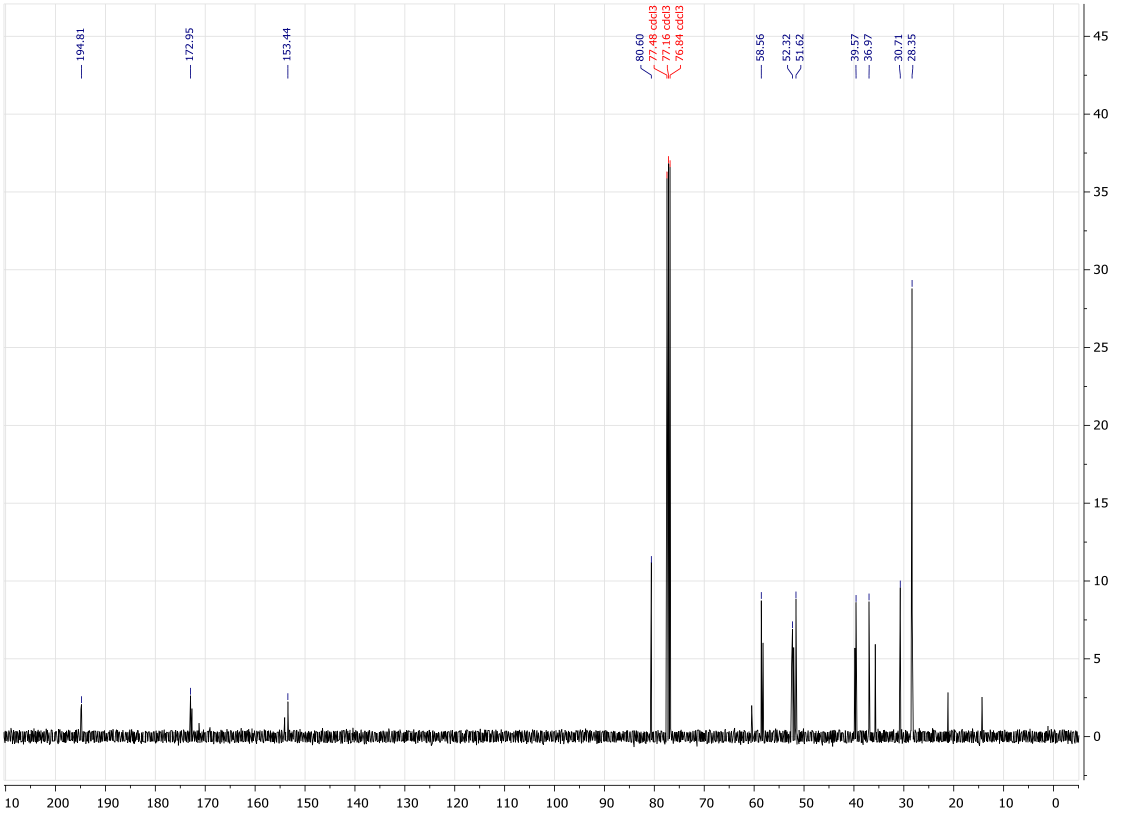

**Figure S62.** $^{13}C$ NMR of compound **8** (100MHz, $CDCl_3$).

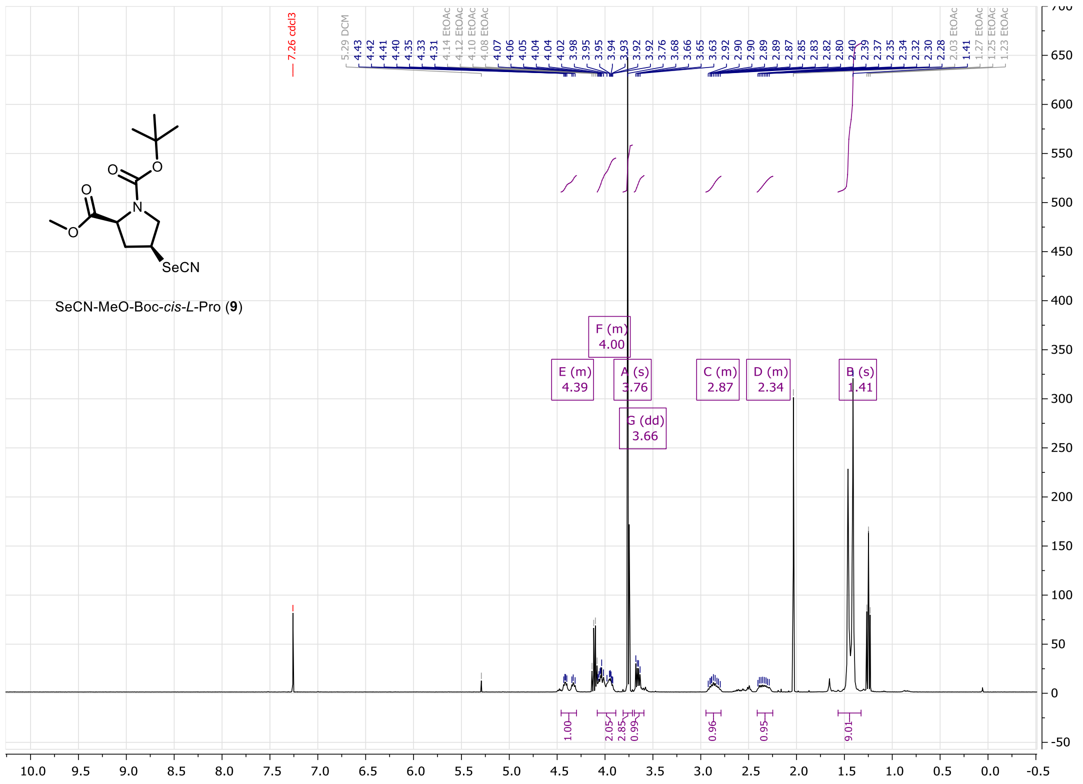

**Figure S63.** $^{1}H$ NMR of compound **9** (400MHz, $CDCl_3$).

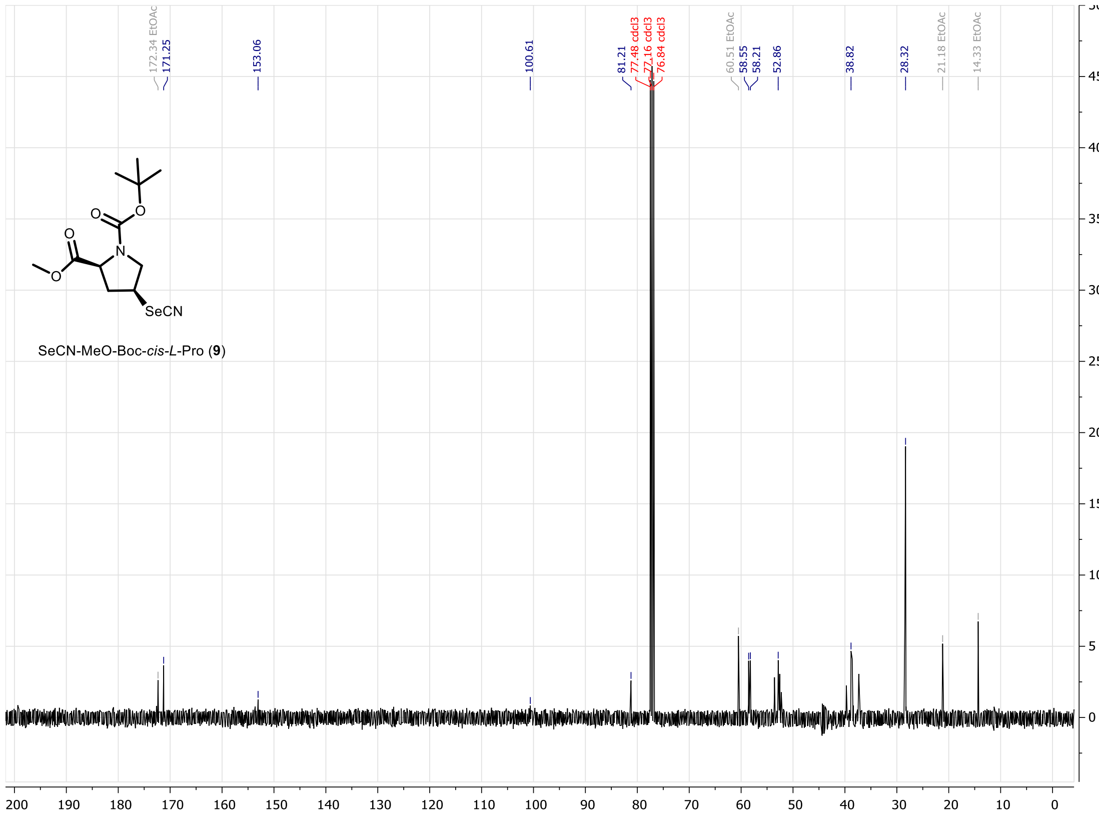

**Figure S64.** $^{13}C$ NMR of compound **9** (100MHz, $CDCl_3$).

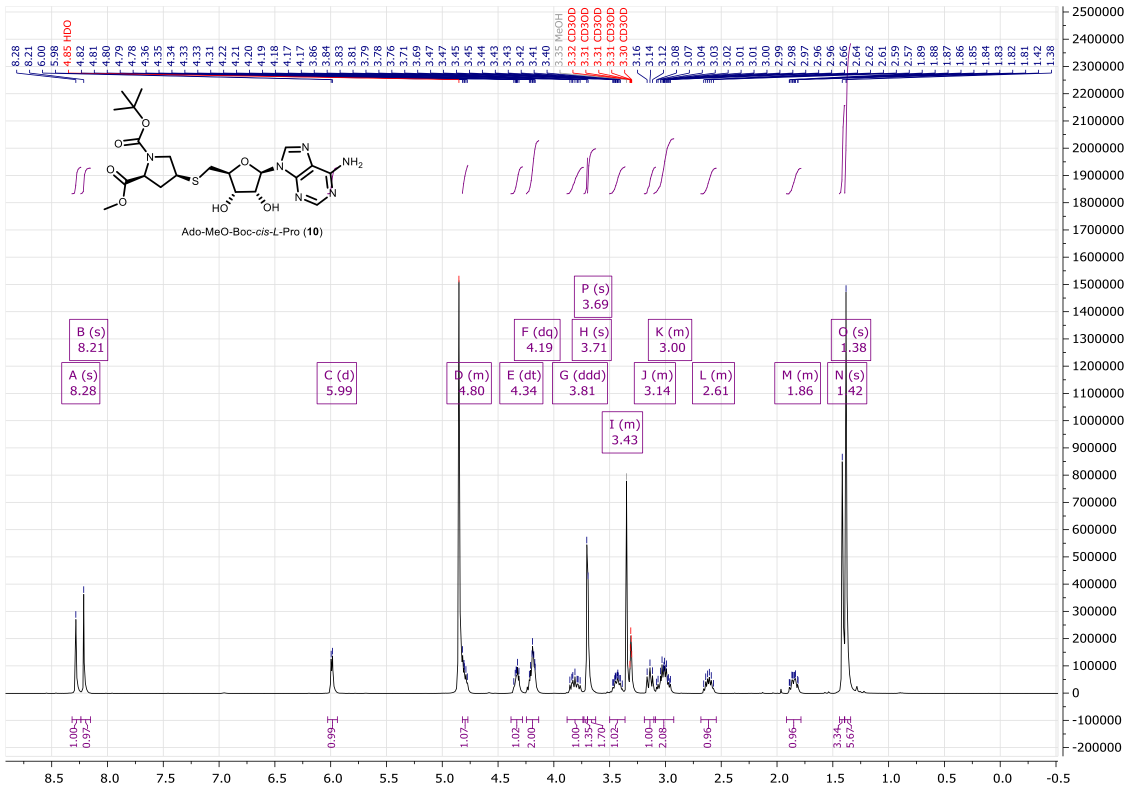


**Figure S65.** $^{1}$H NMR of compound **10** (400MHz, $CD_3OD$).

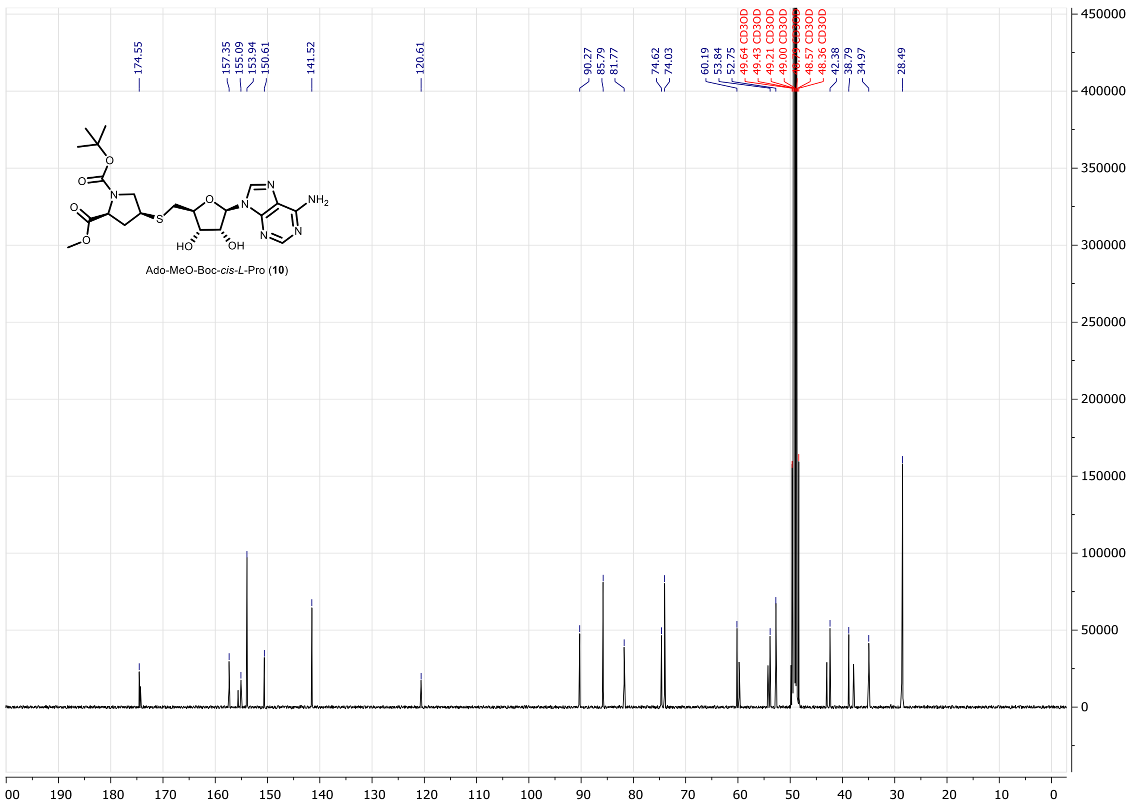


**Figure S66.** $^{13}$C NMR of compound **10** (100MHz, $CD_3OD$).

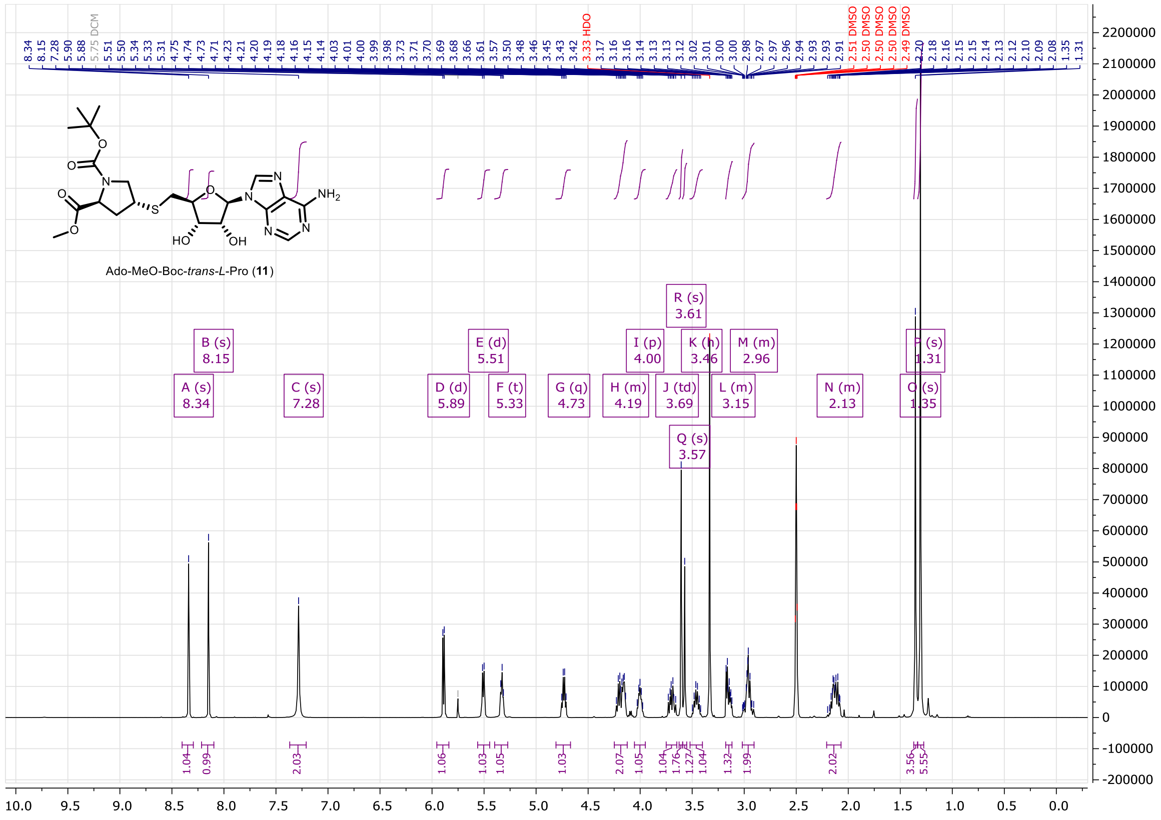

**Figure S67.** $^{1}H$ NMR of compound **11** (400MHz, $d_6$-DMSO).

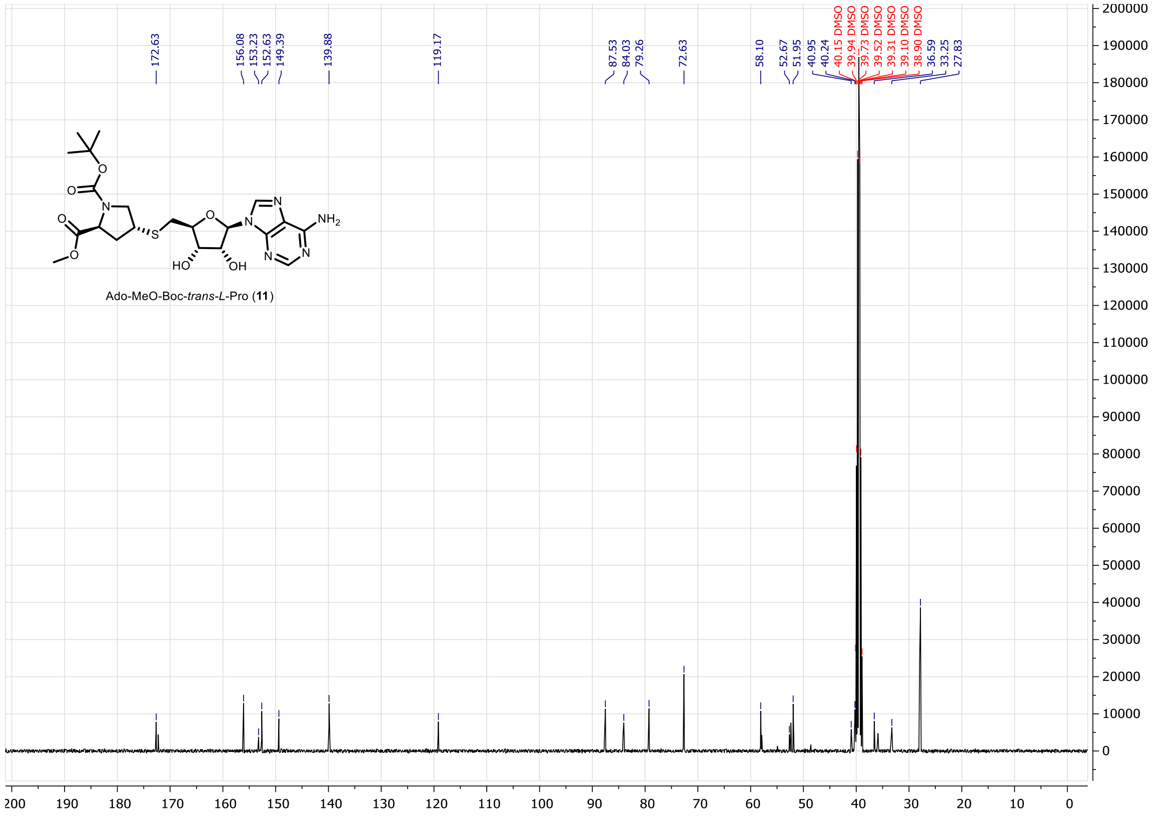

**Figure S68.** $^{13}C$ NMR of compound **11** (100MHz, $d_6$-DMSO).

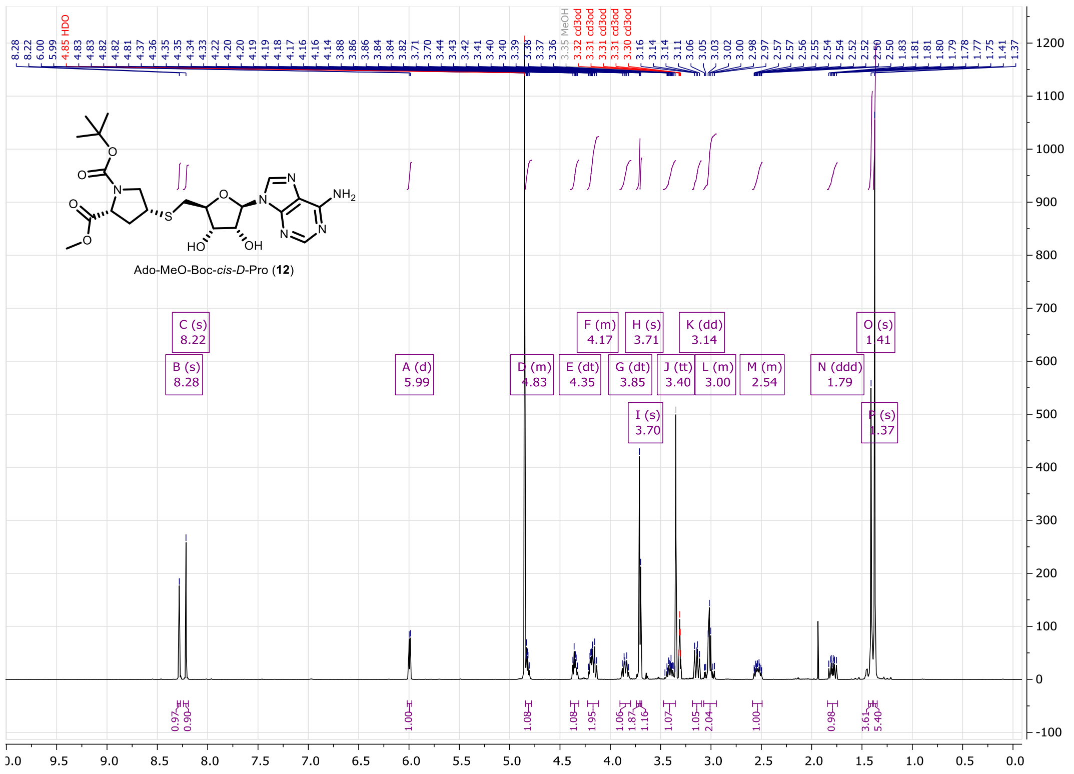

**Figure S69.** $^{1}$H NMR of compound **12** (400MHz, $CD_3OD$).

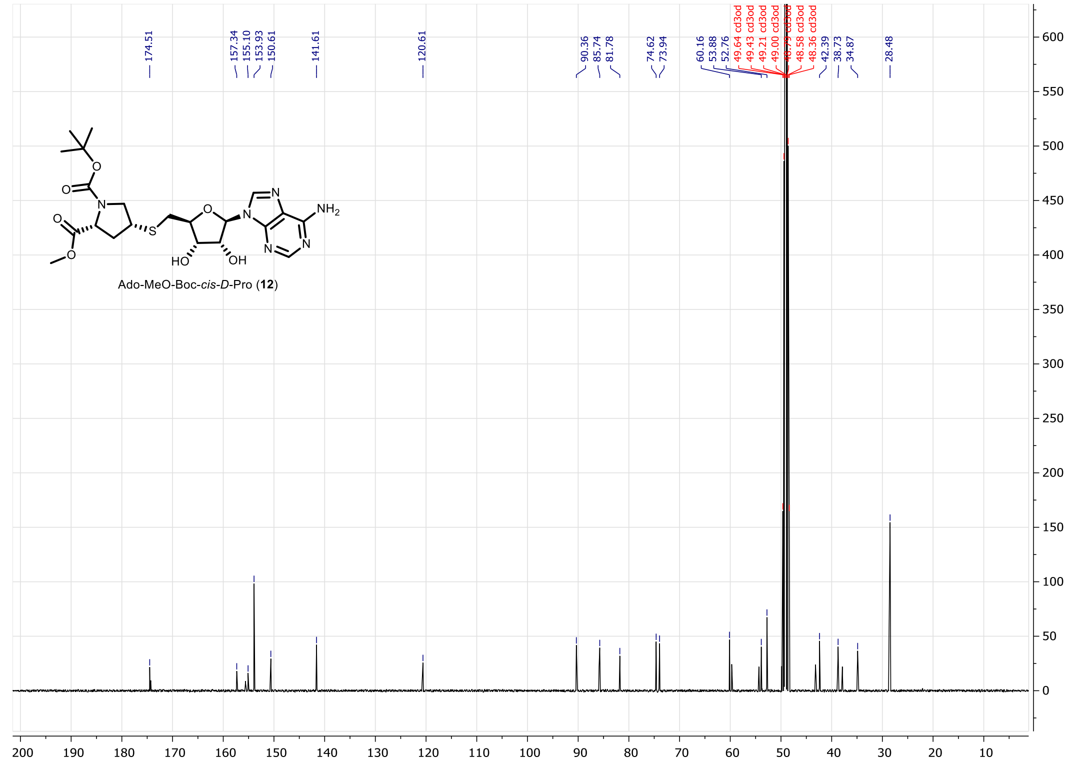

**Figure S70.** $^{13}$C NMR of compound **12** (100MHz, $CD_3OD$).

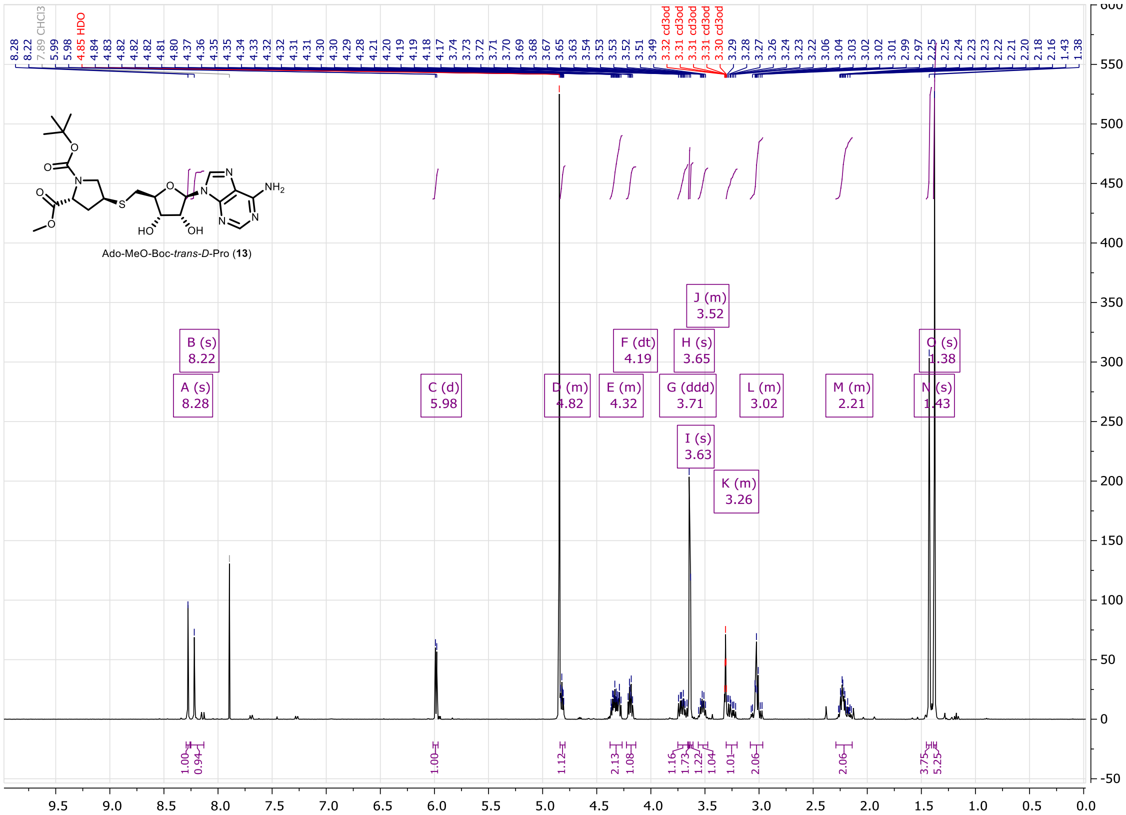


**Figure S71.** $^{1}H$ NMR of compound **13** (400MHz, $CD_3OD$).

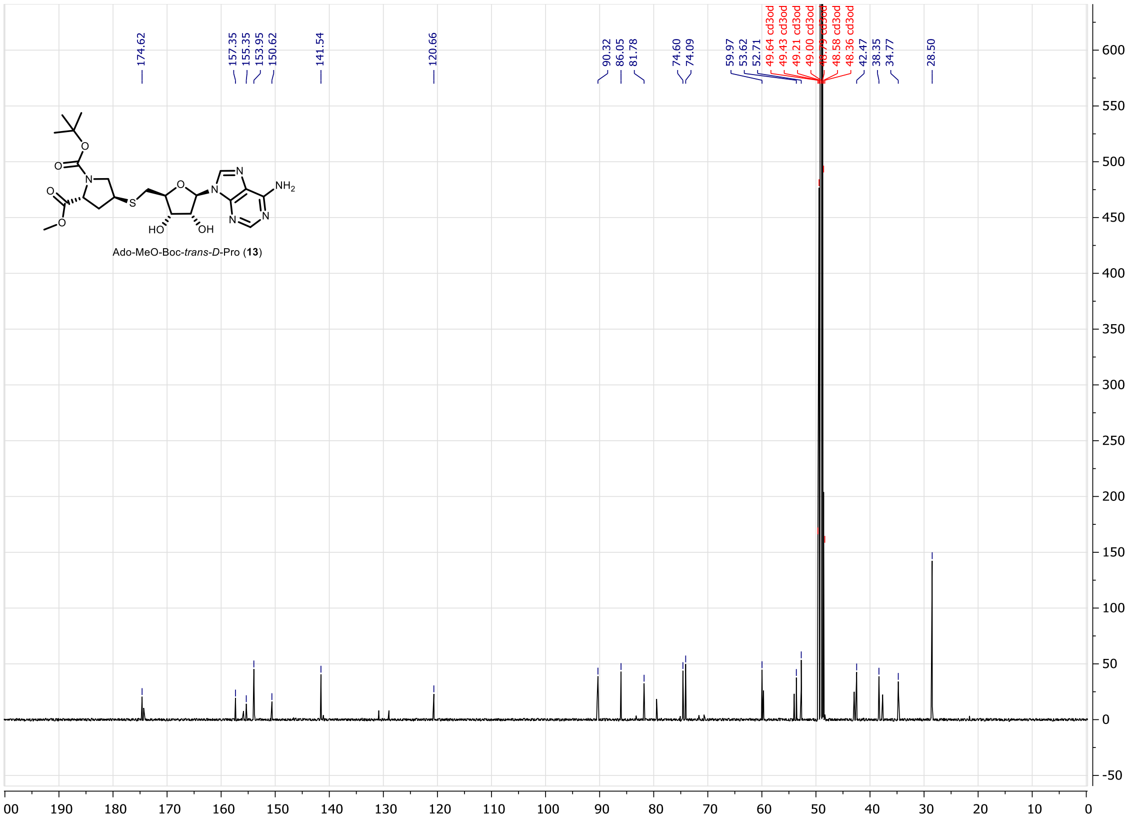


**Figure S72.** $^{13}C$ NMR of compound **13** (100MHz, $CD_3OD$).

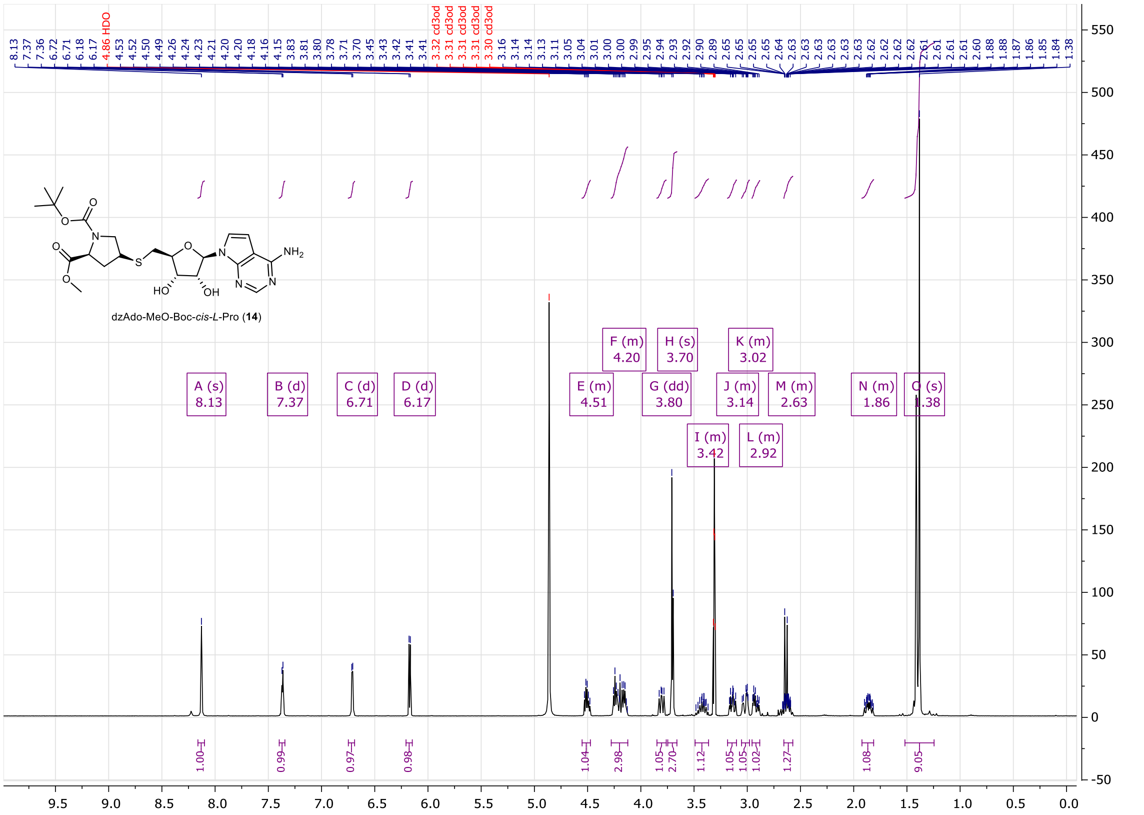


**Figure S73.** $^{1}H$ NMR of compound **14** (400MHz, $CD_3OD$).

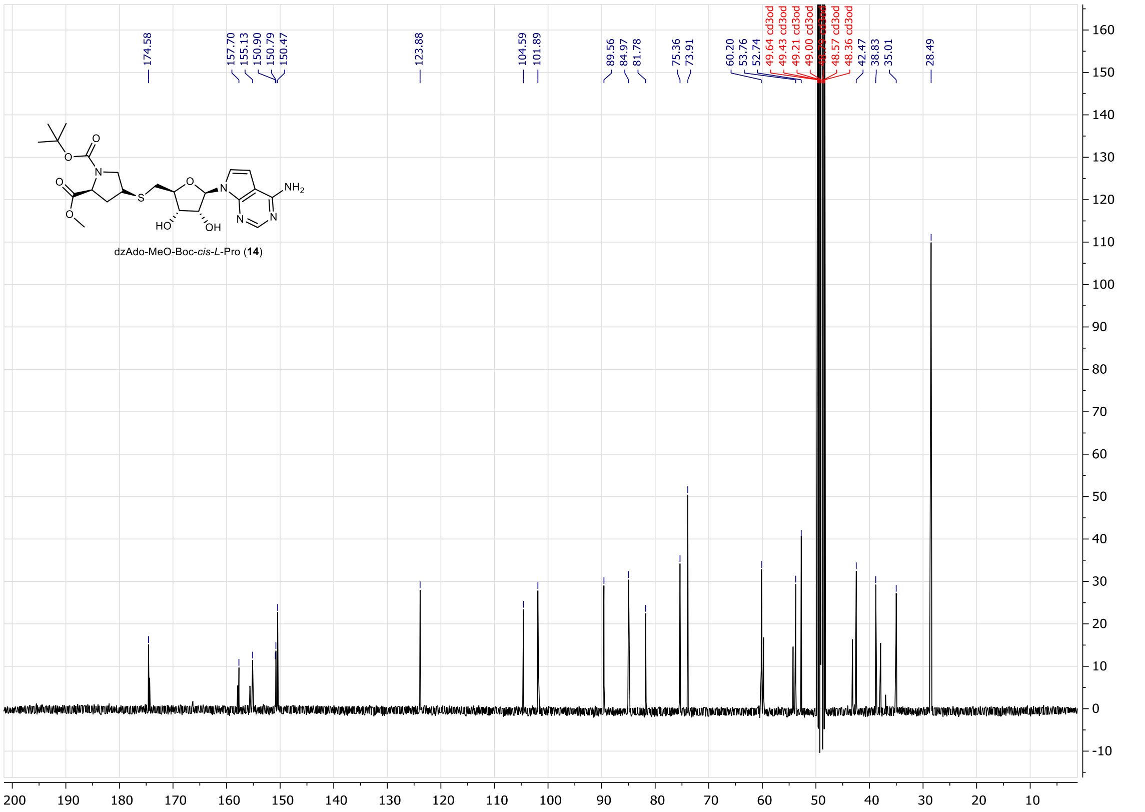


**Figure S74.** $^{13}C$ NMR of compound **14** (100MHz, $CD_3OD$).

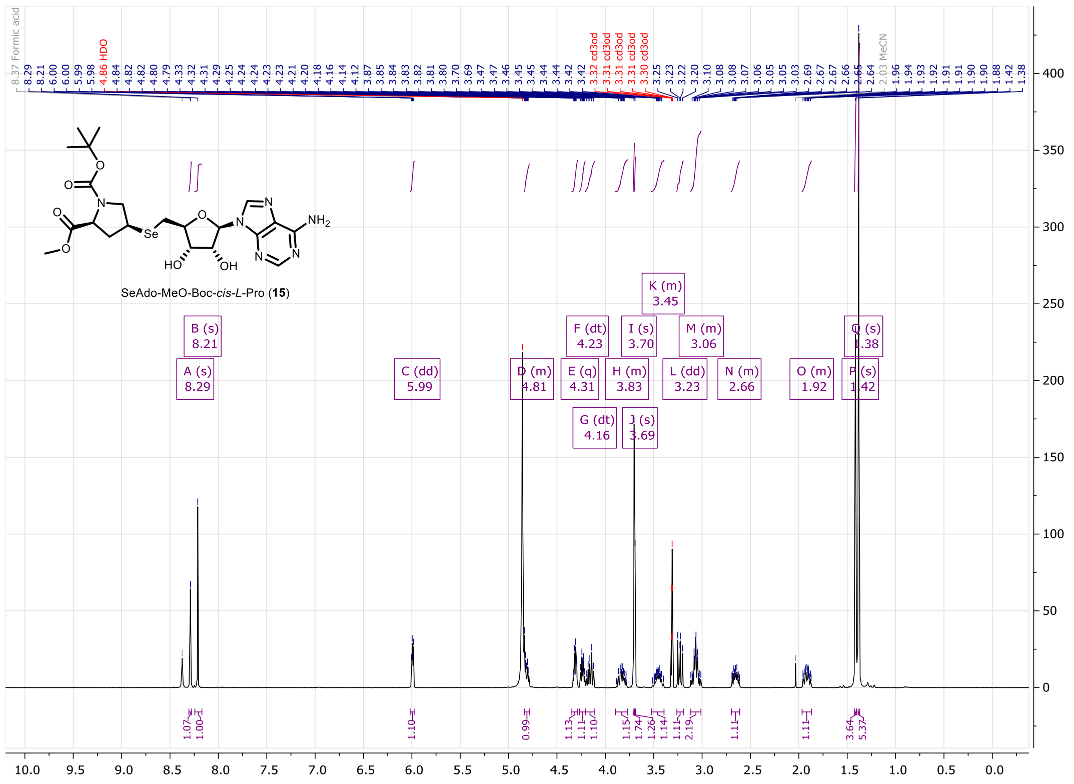

**Figure S75.** $^{1}H$ NMR of compound **15** (400MHz, $CD_3OD$).

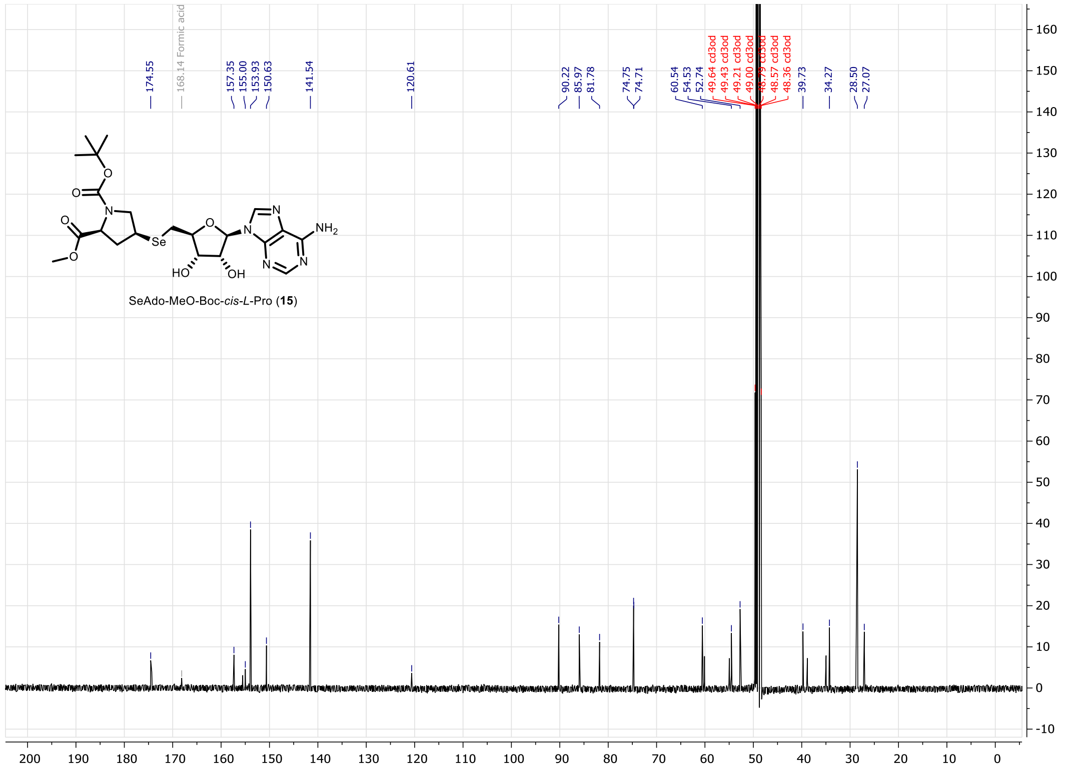

**Figure S76.** $^{13}C$ NMR of compound **15** (100MHz, $CD_3OD$).

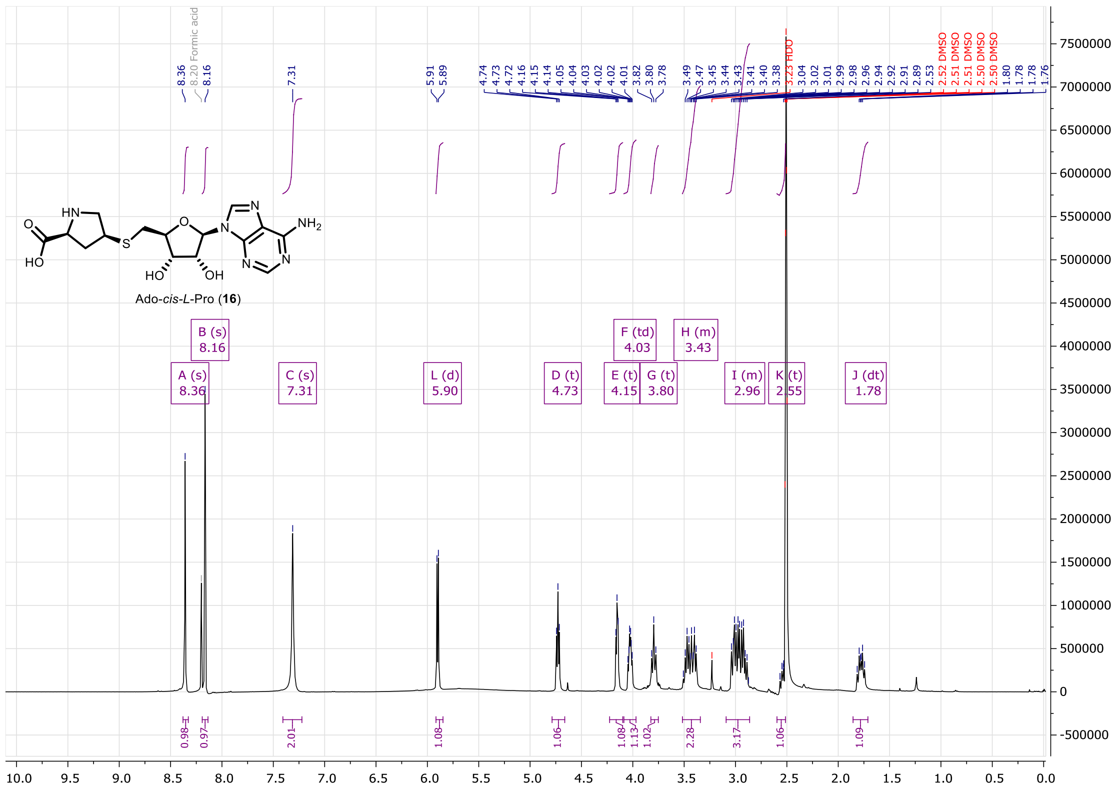

**Figure S77.** $^{1}H$ NMR of compound **16** (400MHz, $d_6$-DMSO).

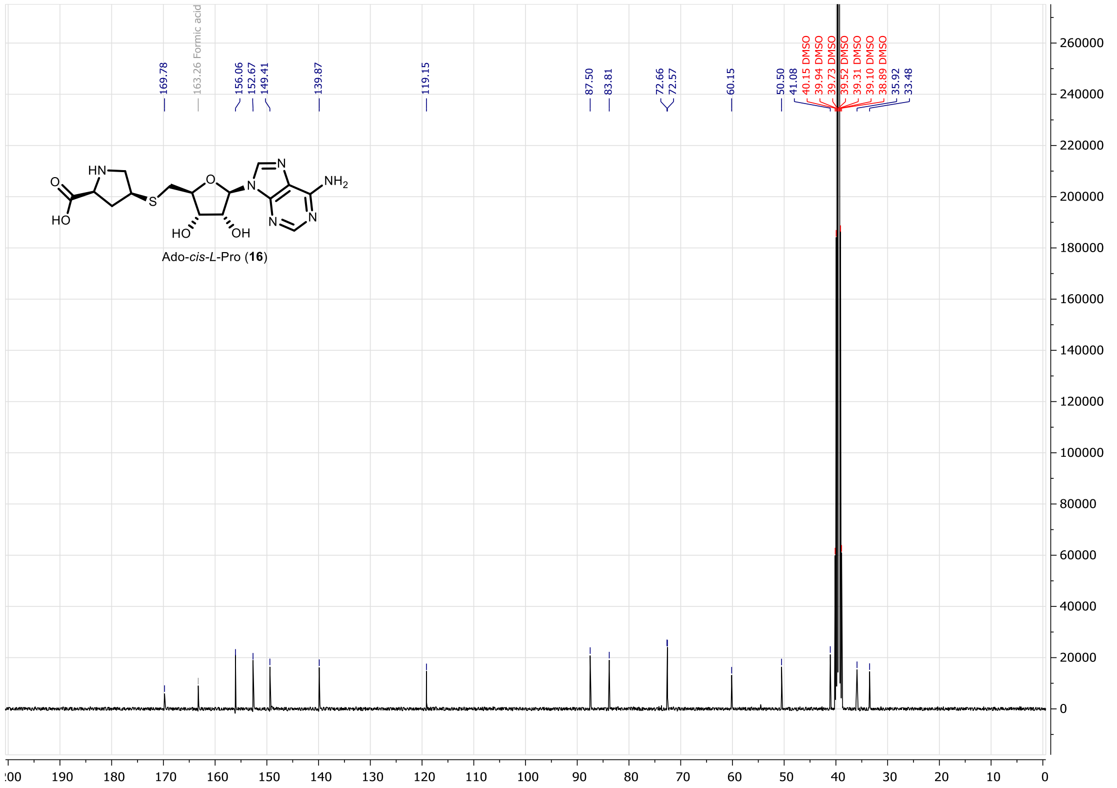

**Figure S78.** $^{13}C$ NMR of compound **16** (100MHz, $d_6$-DMSO).

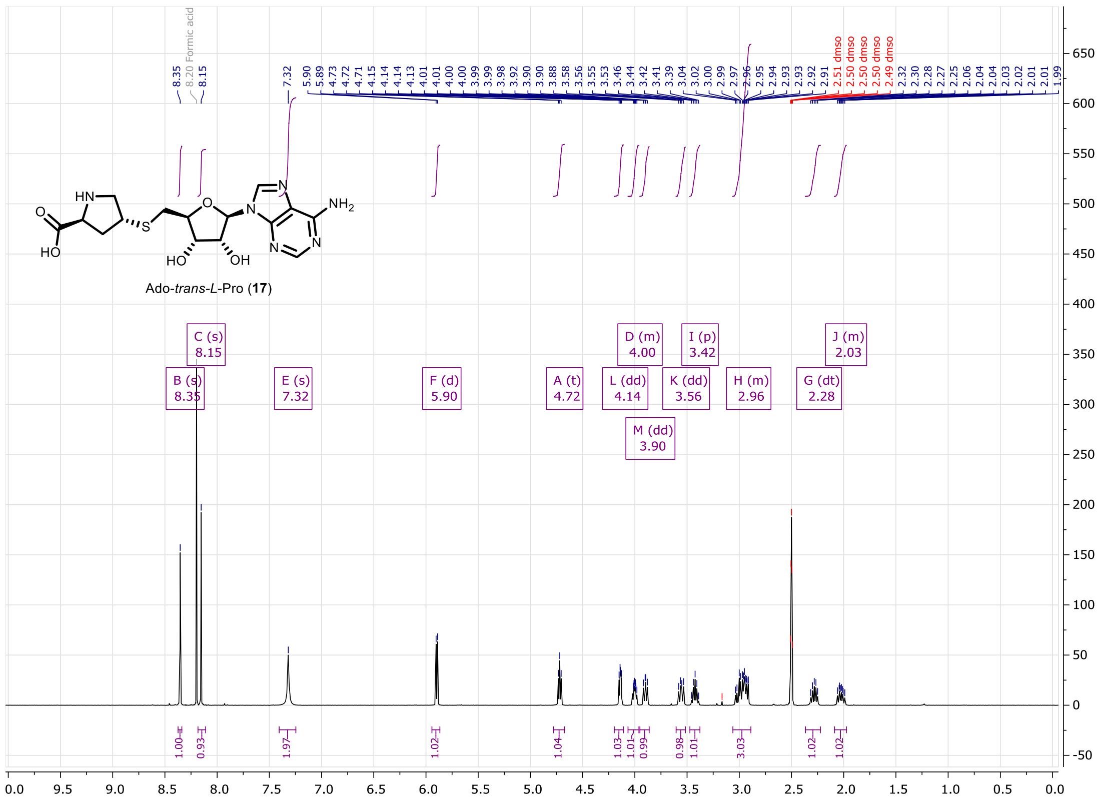

**Figure S79.** $^{1}H$ NMR of compound **17** (400MHz, $d_6$-DMSO).

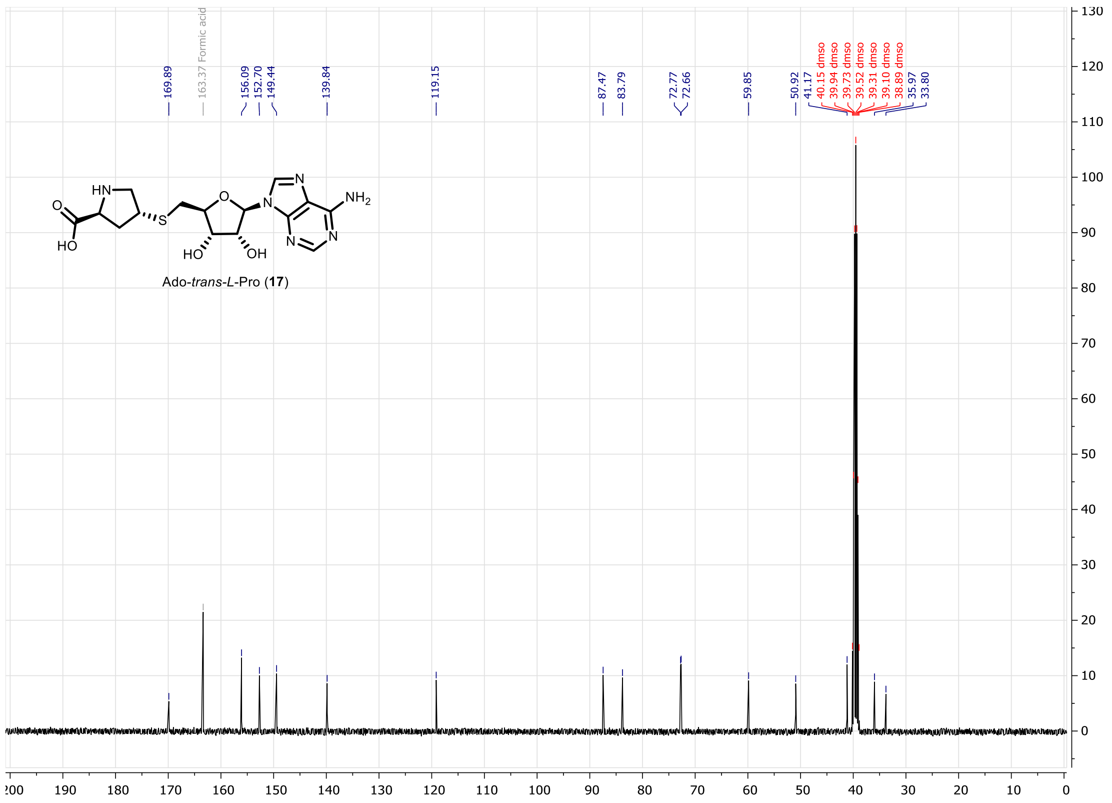

**Figure S80.** $^{13}C$ NMR of compound **17** (100MHz, $d_6$-DMSO).

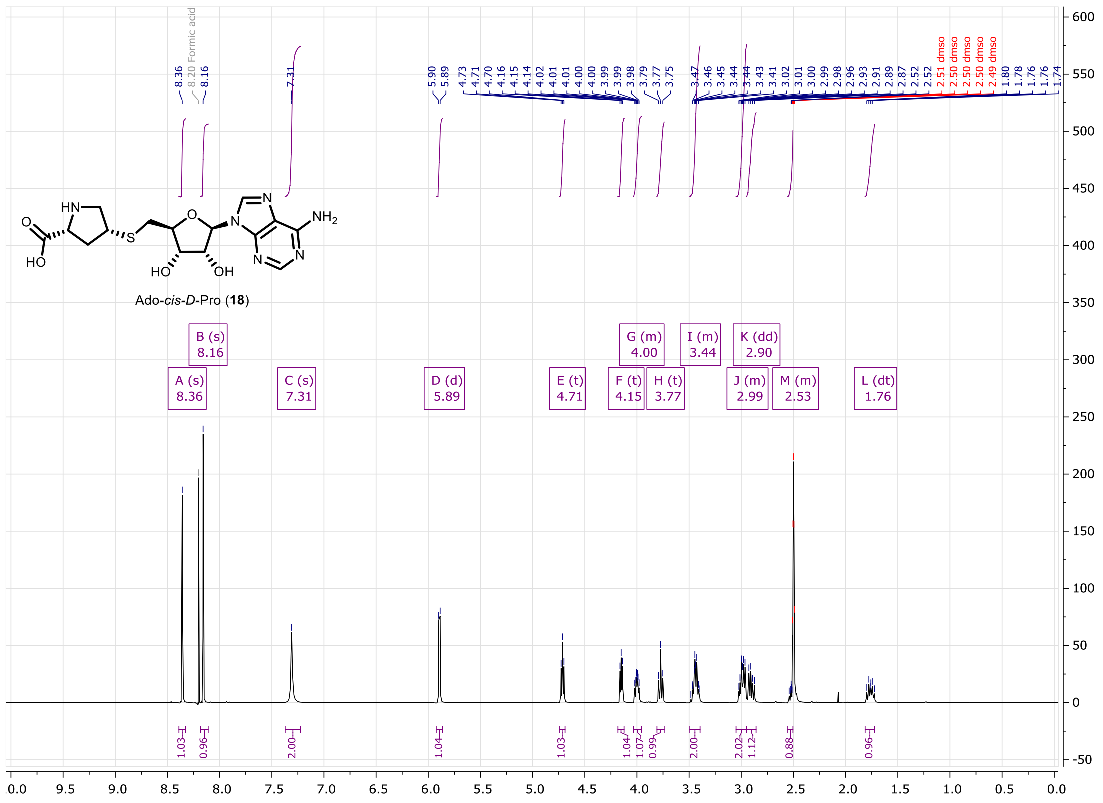

**Figure S81.** $^{1}H$ NMR of compound **18** (400MHz, $d_6$-DMSO).

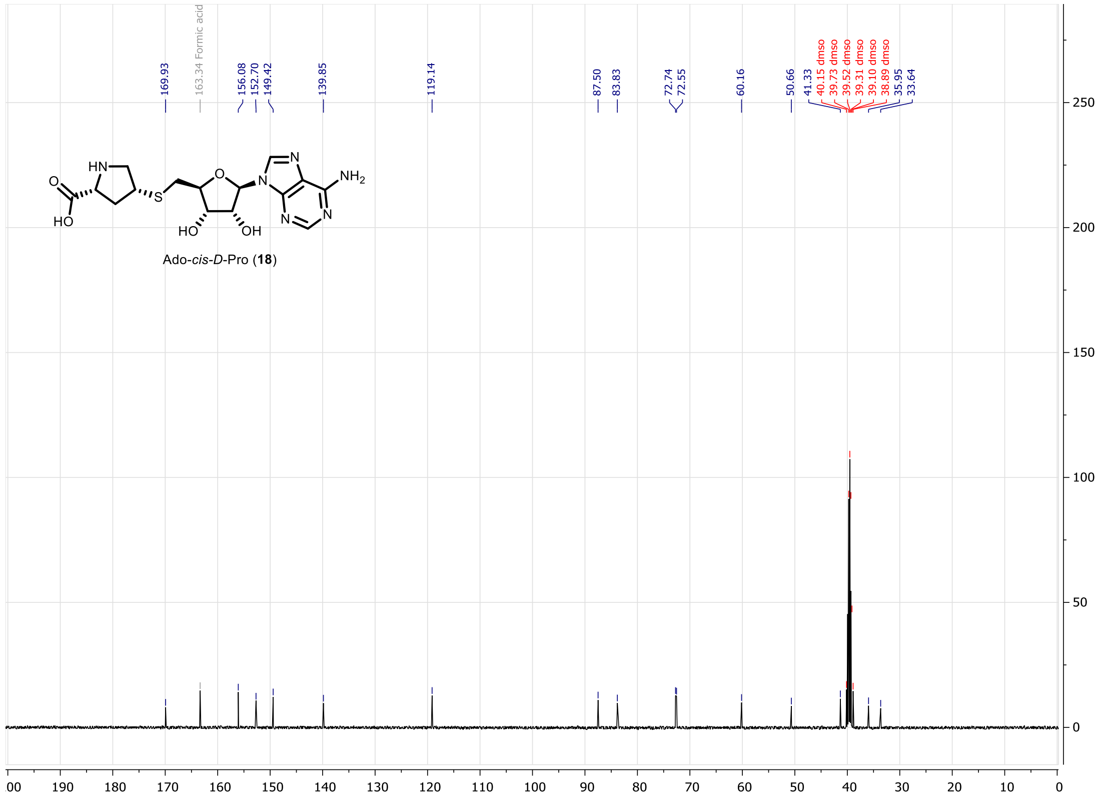

**Figure S82.** $^{13}C$ NMR of compound **18** (100MHz, $d_6$-DMSO).

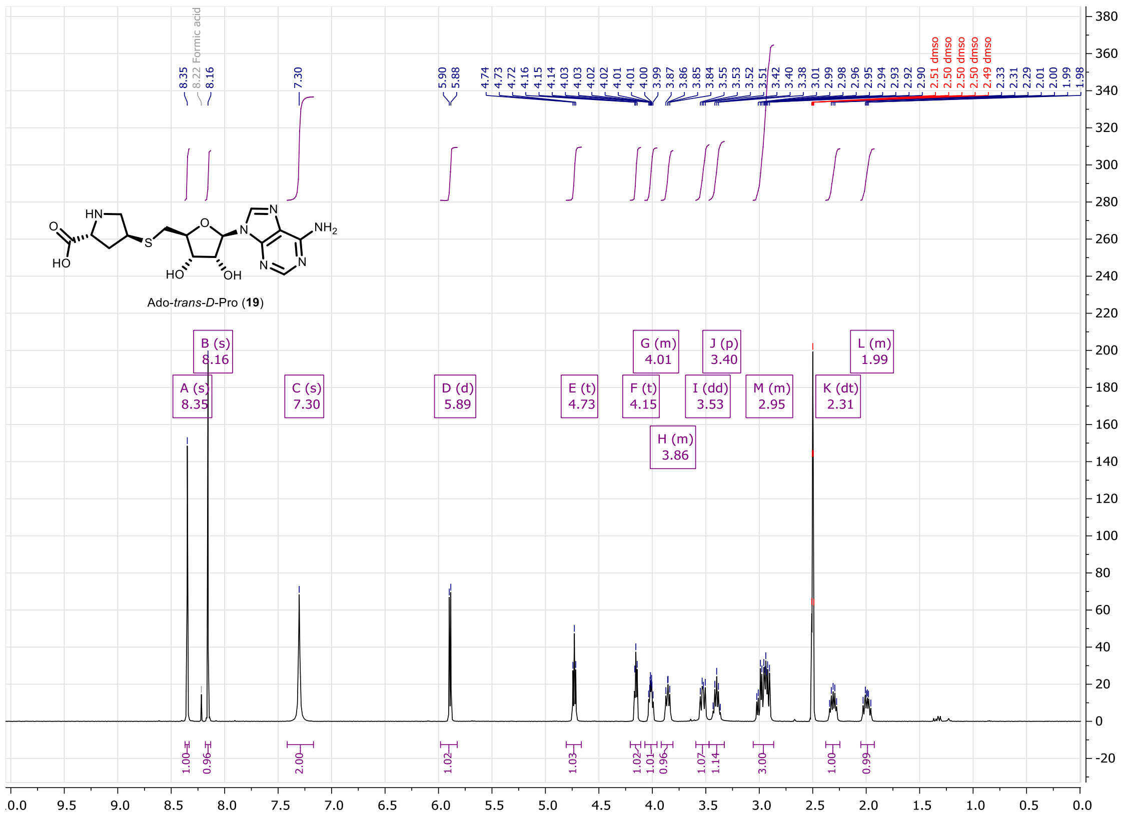

**Figure S83.** $^{1}$H NMR of compound **19** (400MHz, $d_6$-DMSO).

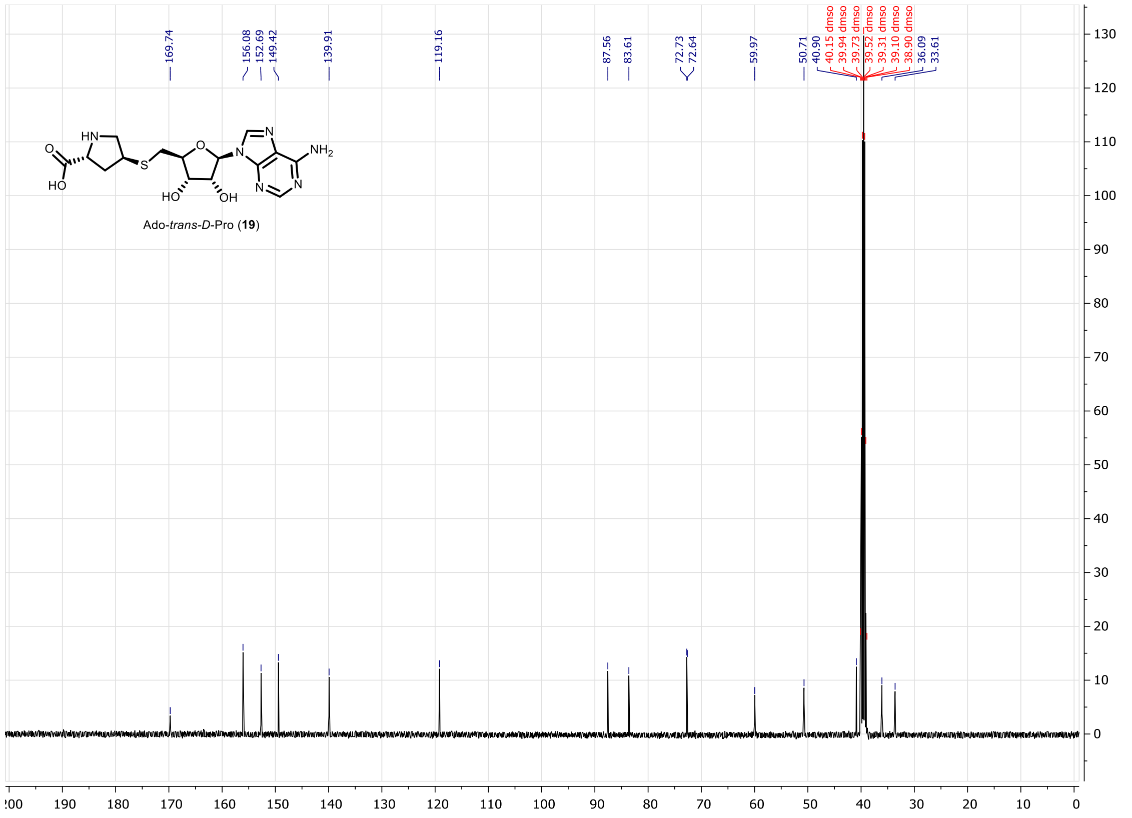

**Figure S84.** $^{13}$C NMR of compound **19** (100MHz, $d_6$-DMSO).

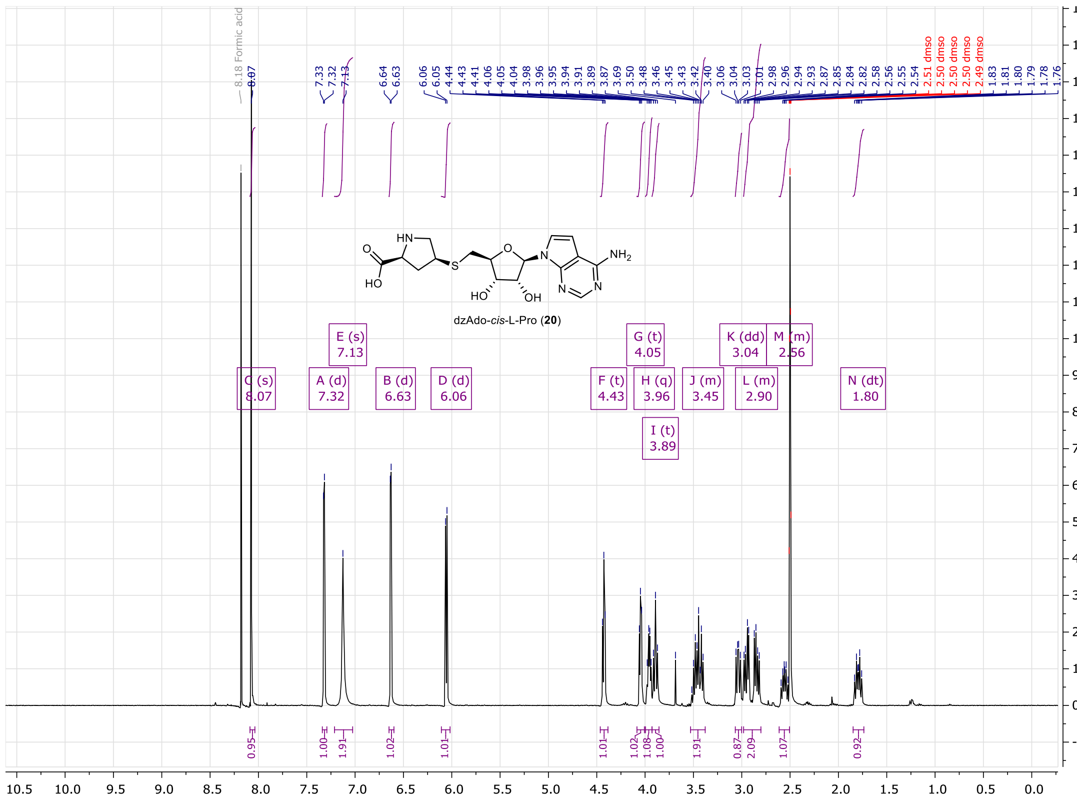

**Figure S85.** $^{1}$H NMR of compound **20** (400MHz, $d_6$-DMSO).

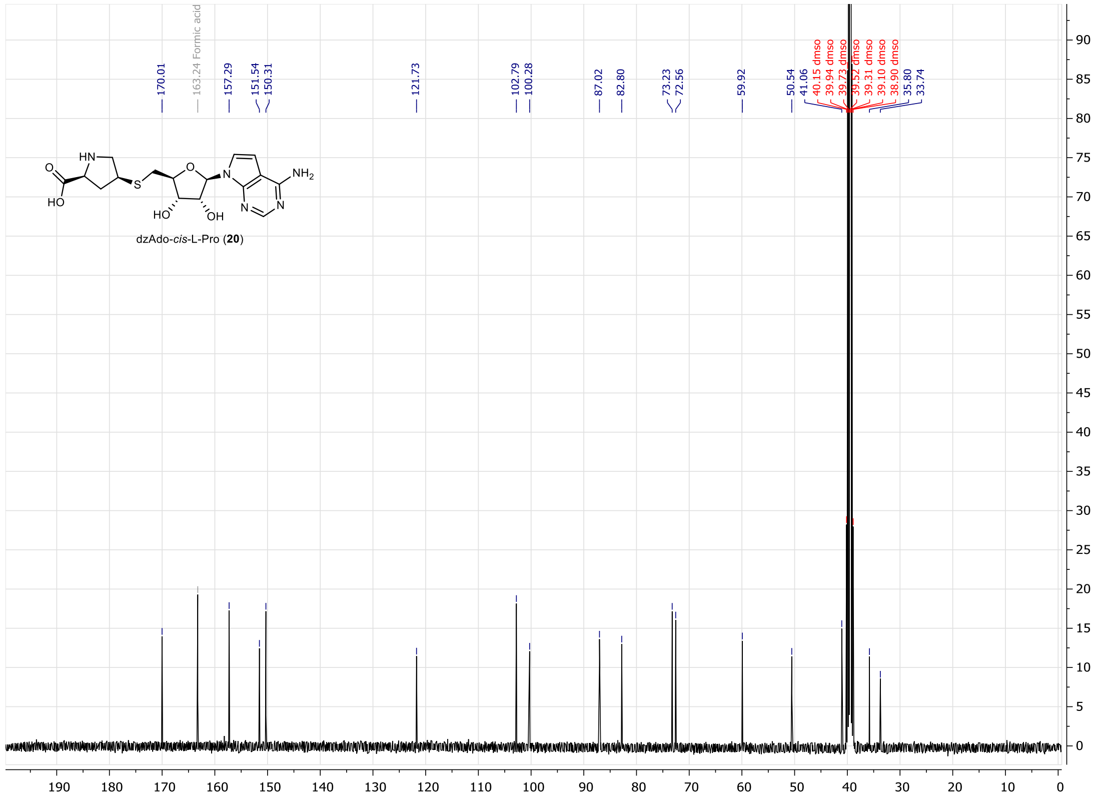

**Figure S86.** $^{13}$C NMR of compound **20** (100MHz, $d_6$-DMSO).

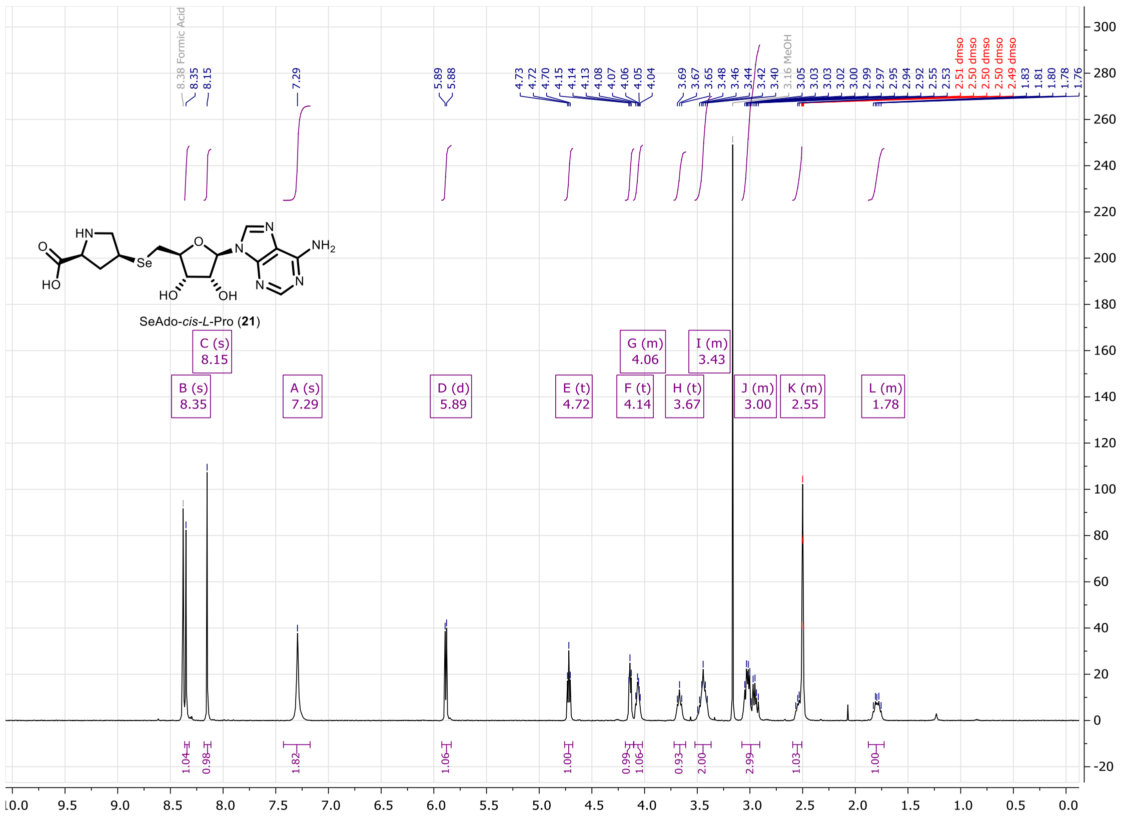

**Figure S87.** $^{1}$H NMR of compound **21** (400MHz, $d_6$-DMSO).

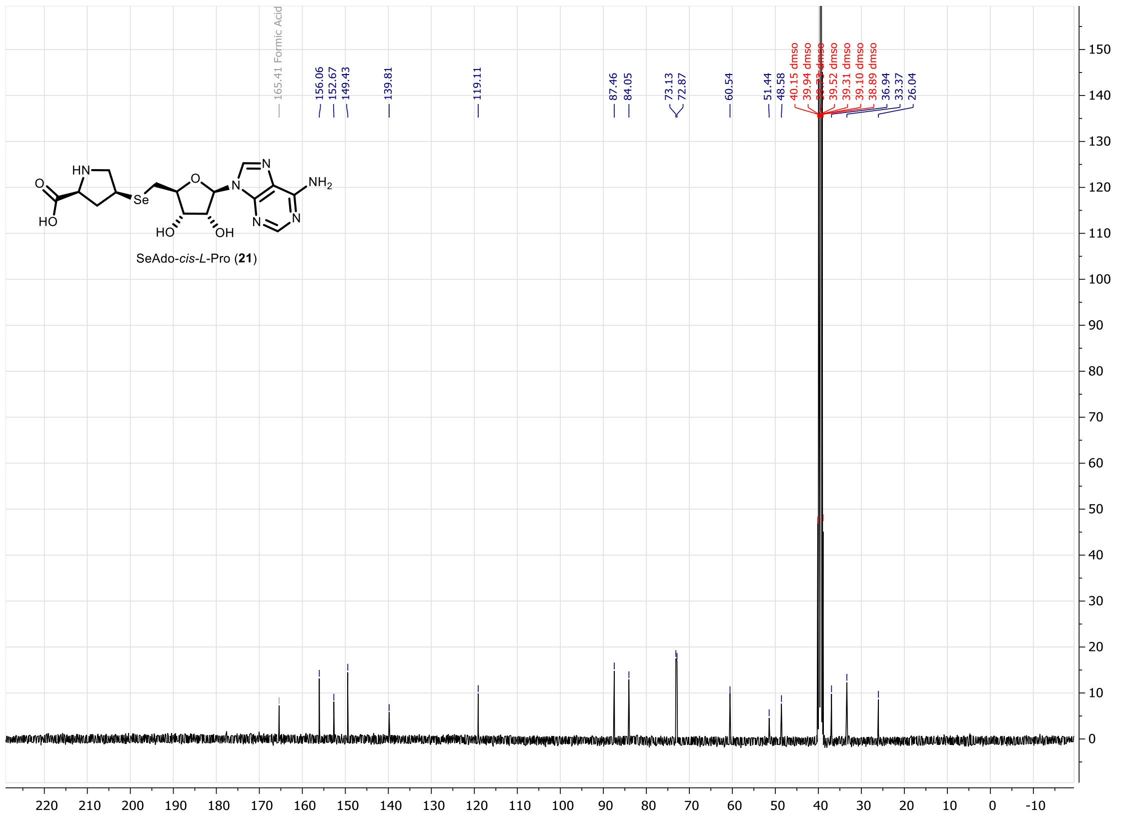

**Figure S88.** $^{13}$C NMR of compound **21** (100MHz, $d_6$-DMSO).

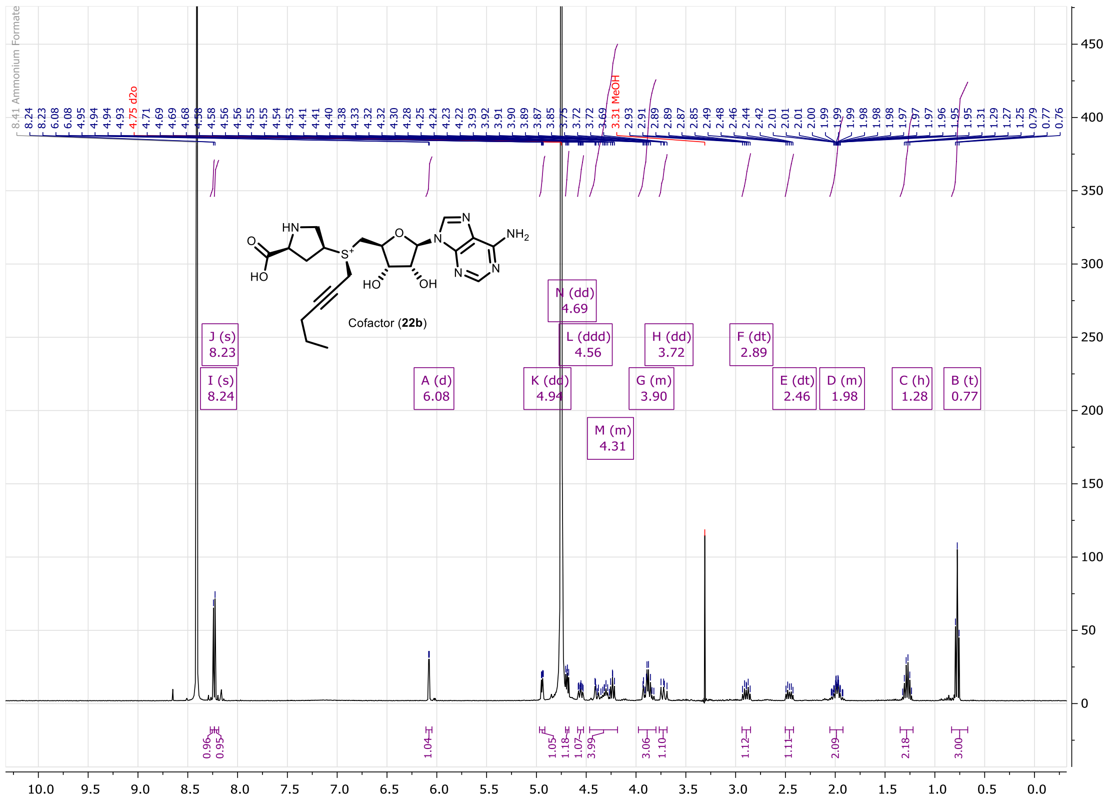

**Figure S89.** $^{1}$H NMR of compound **22b** (400MHz, $D_2O$).

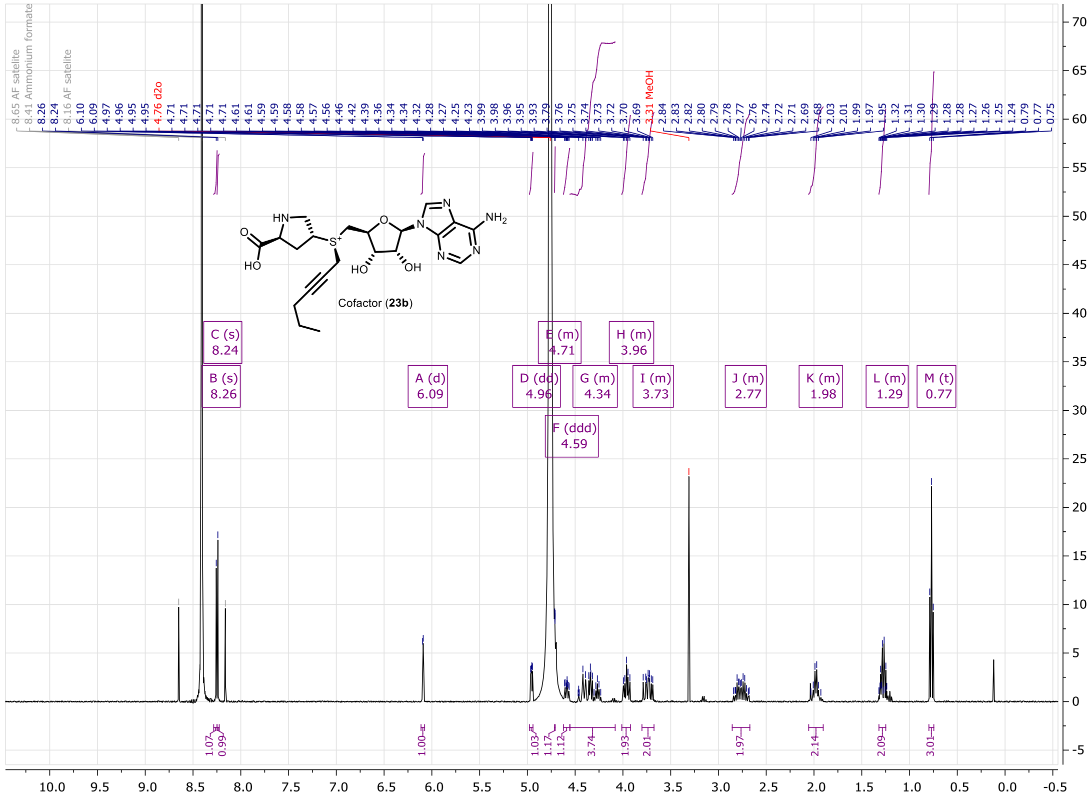

**Figure S90.** $^{1}$H NMR of compound **23b** (400MHz, $D_2O$).

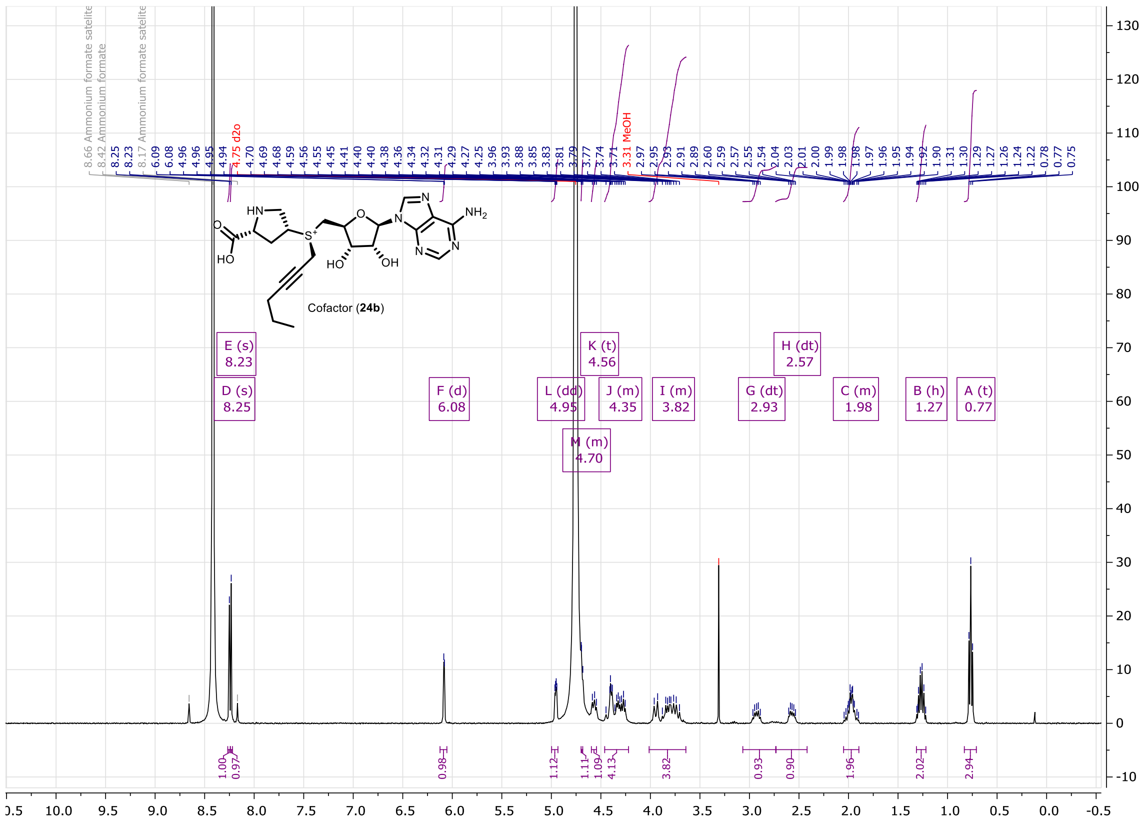

**Figure S91.** $^1$H NMR of compound **24b** (400MHz, $D_2O$).

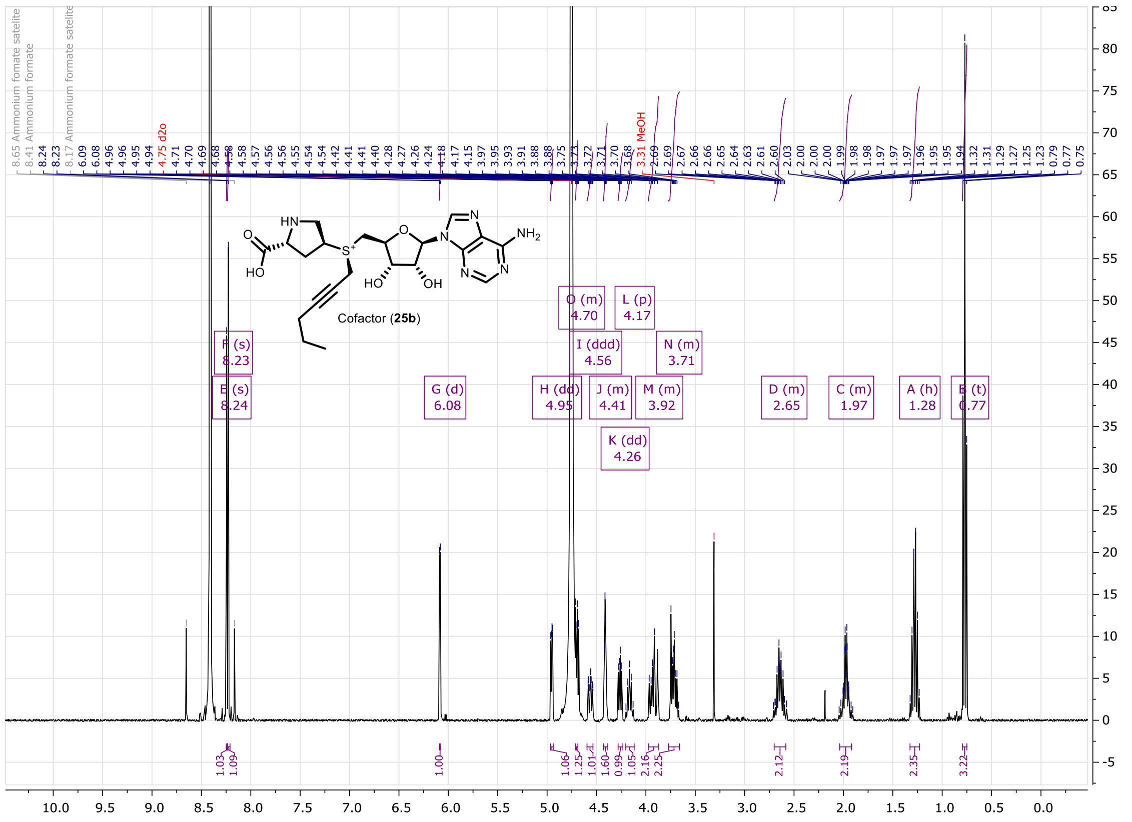

**Figure S92.** $^1$H NMR of compound **25b** (400MHz, $D_2O$).

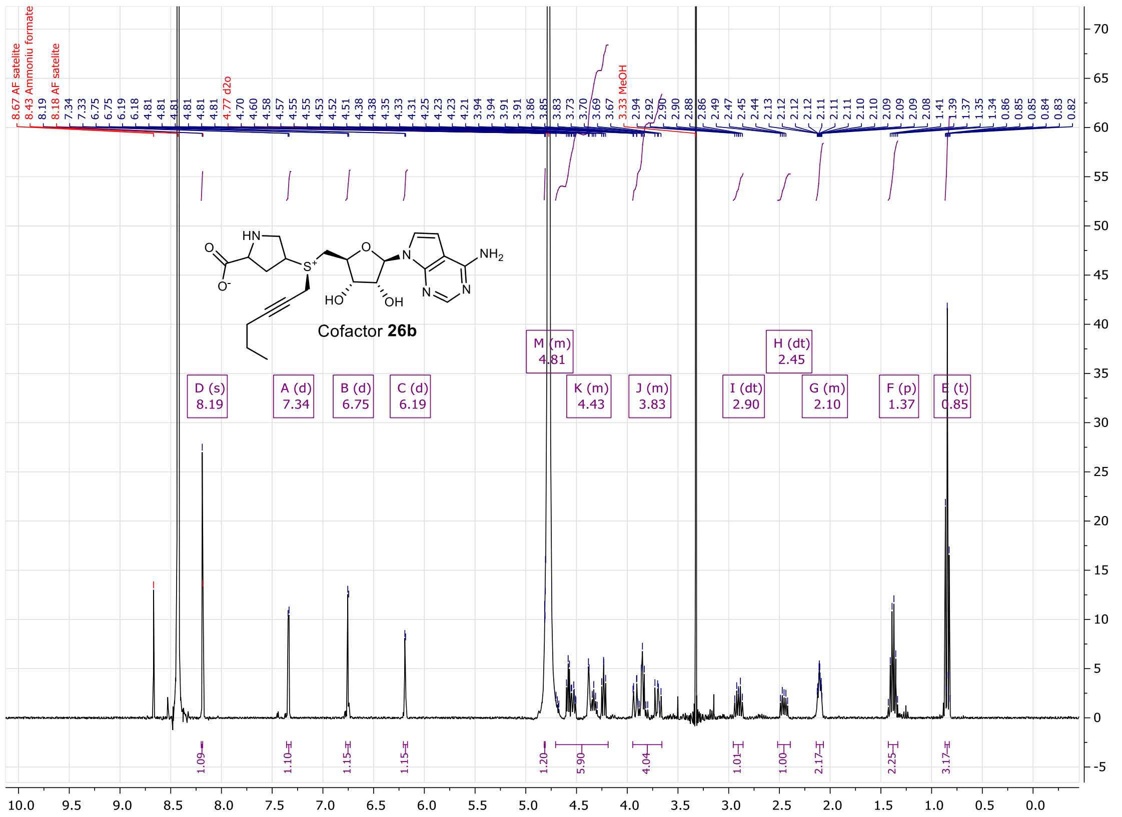

**Figure S93.** $^{1}H$ NMR of compound **26b** (400MHz, $D_2O$).

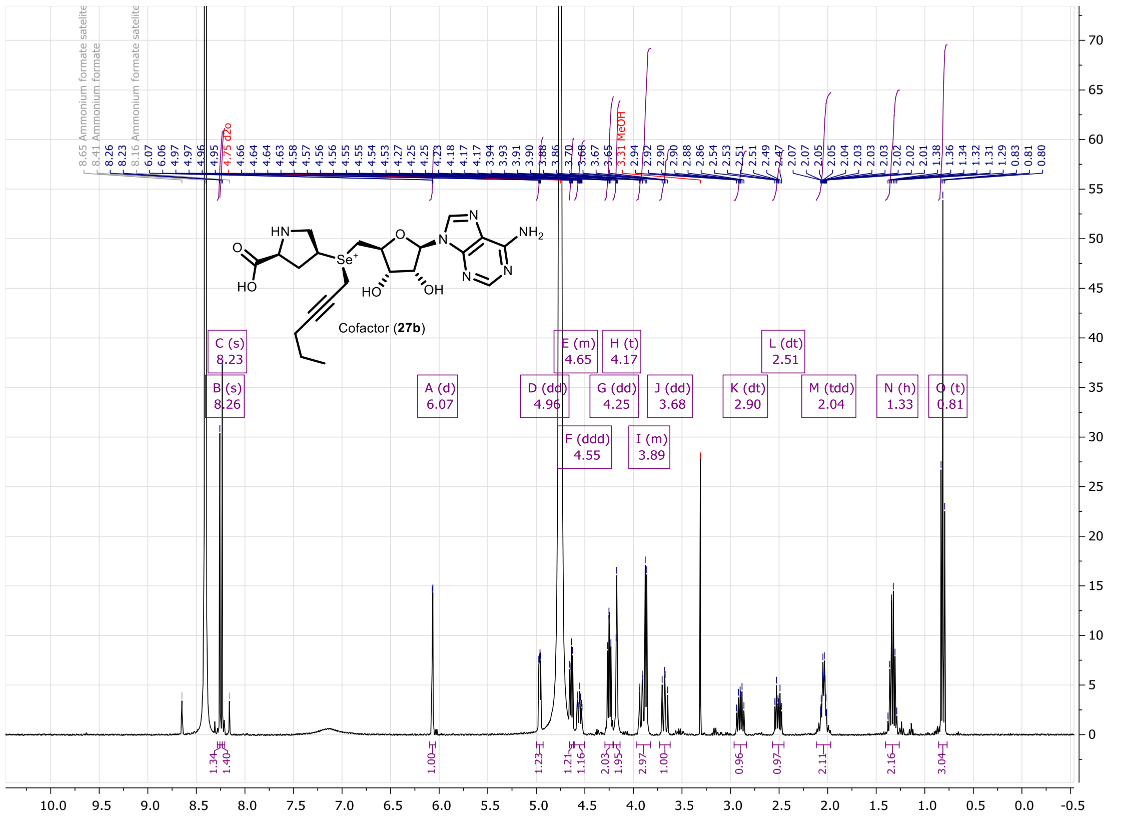

**Figure S94.** $^{1}H$ NMR of compound **27b** (400MHz, $D_2O$).

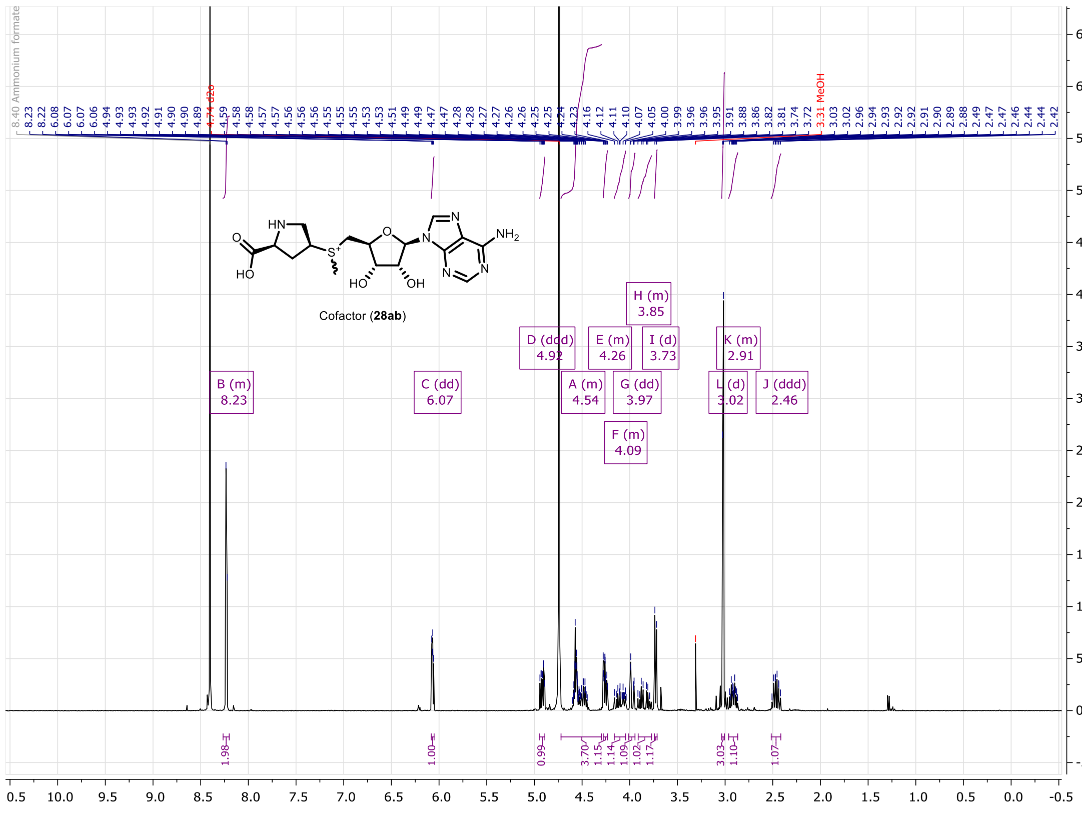

**Figure S95.** $^1H$ NMR of compound **28ab** (400MHz, $D_2O$).

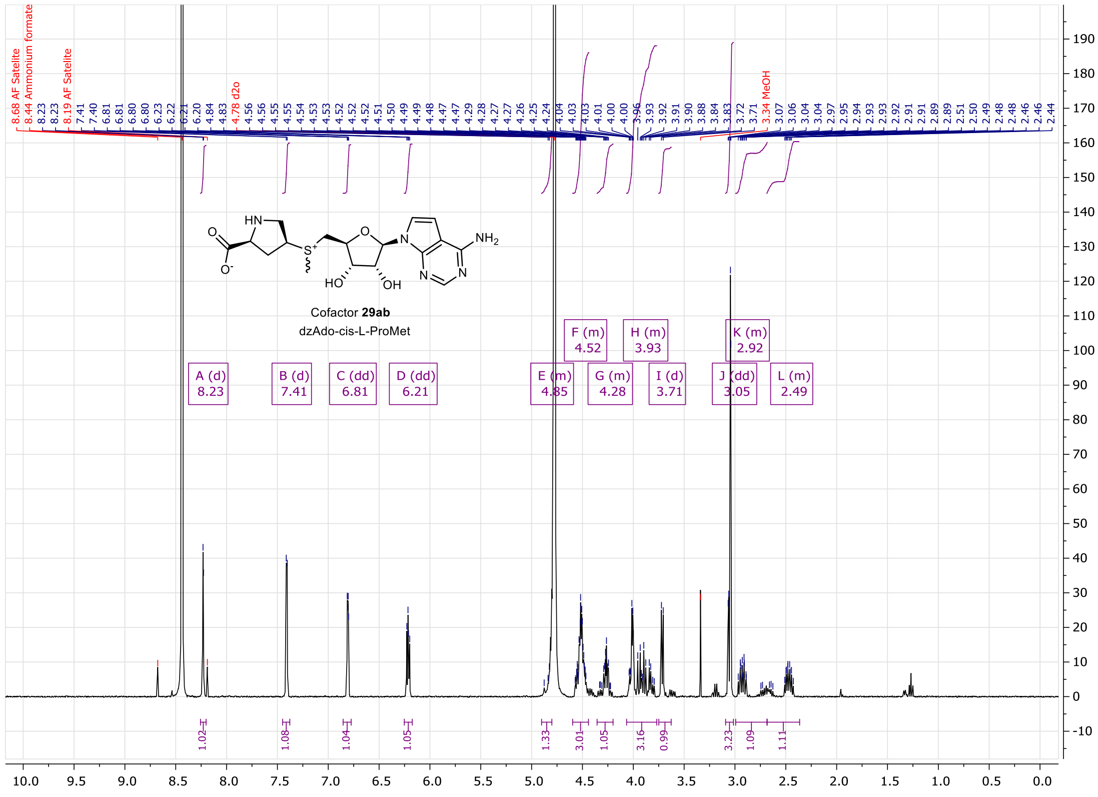

**Figure S96.** $^1H$ NMR of compound **29ab** (400MHz, $D_2O$).

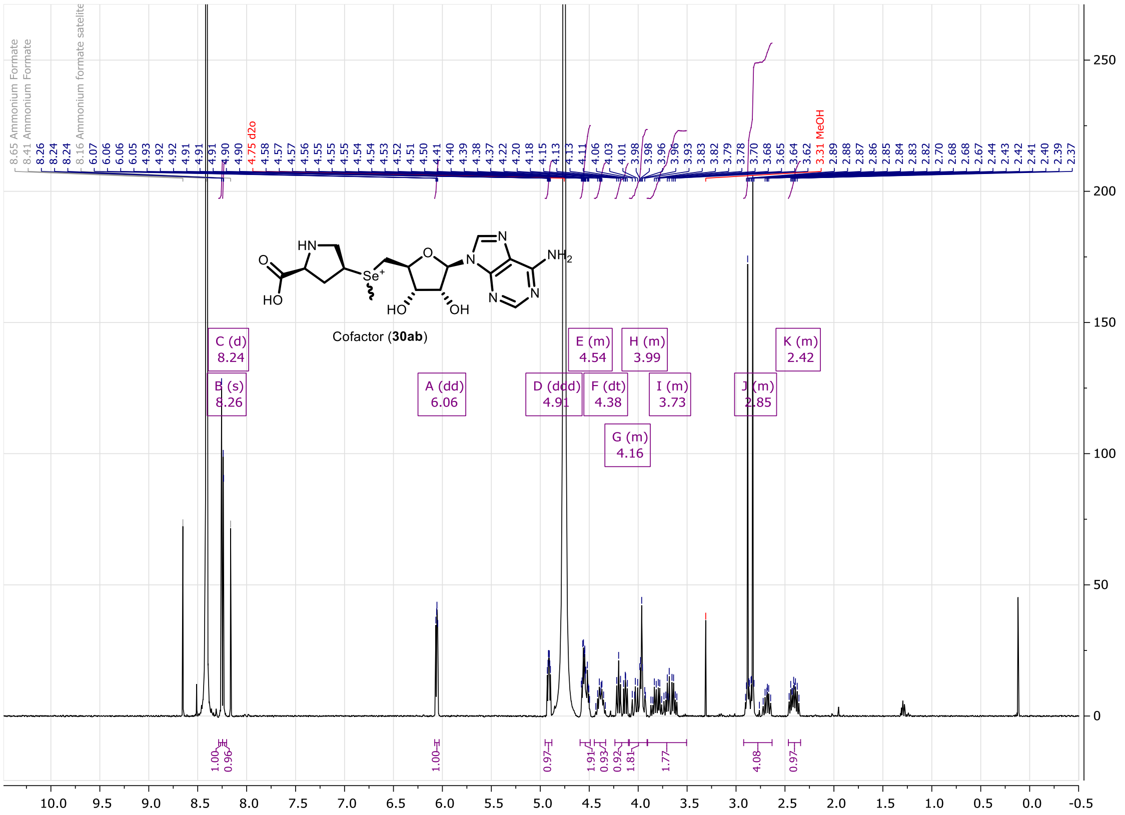


**Figure S97.** $^{1}H$ NMR of compound **30ab** (400MHz, $D_2O$).

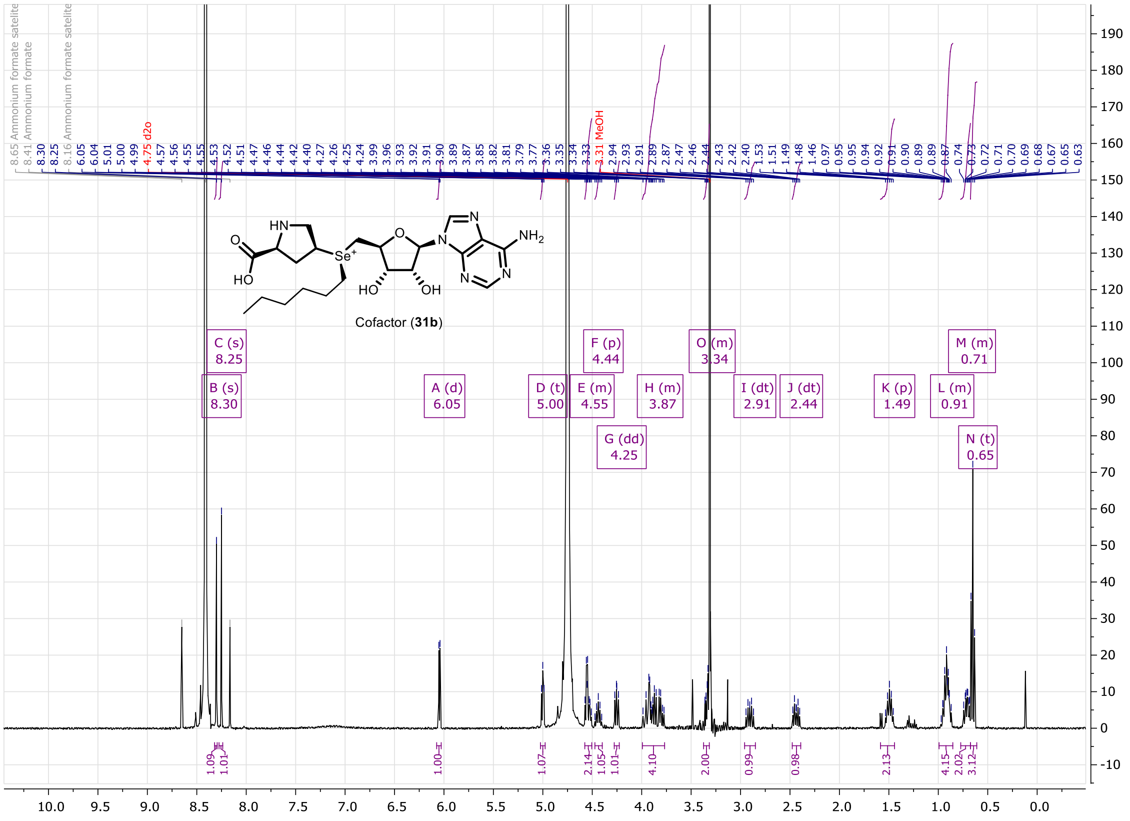


**Figure S98.** $^{1}H$ NMR of compound **31b** (400MHz, $D_2O$).

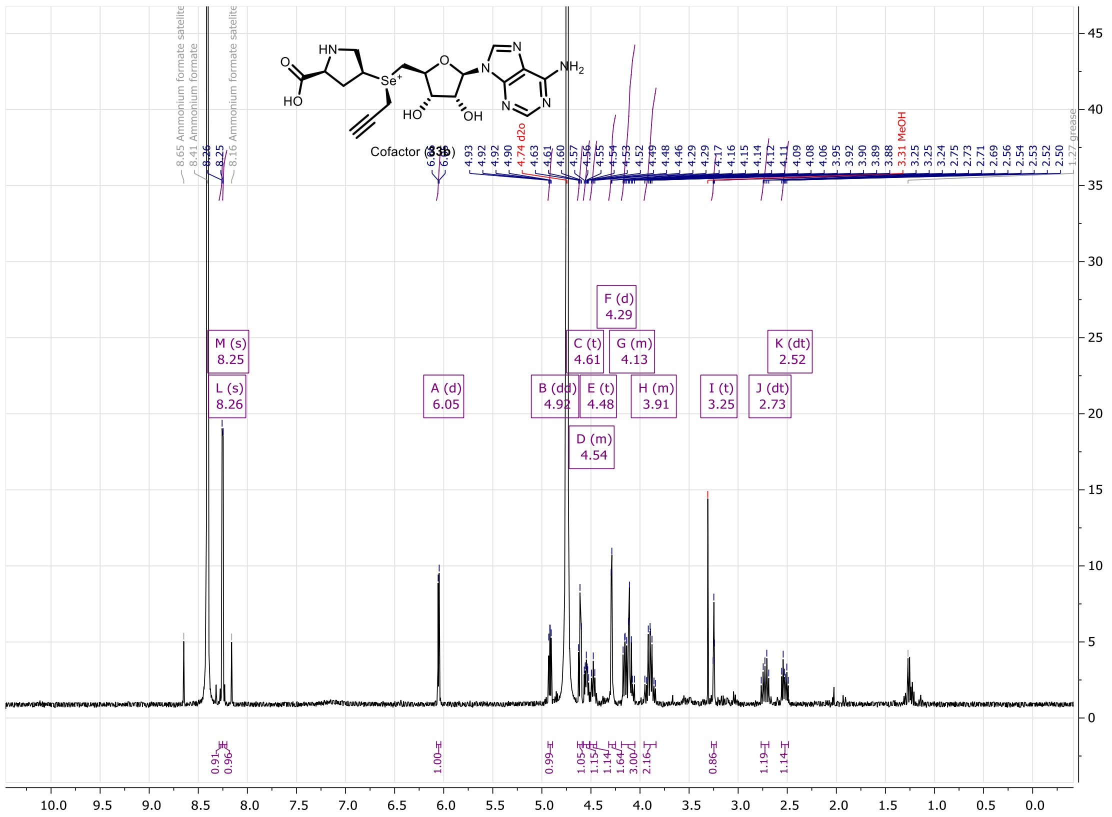


**Figure S99.** $^{1}$H NMR of compound **33b** (400MHz, $D_2O$).

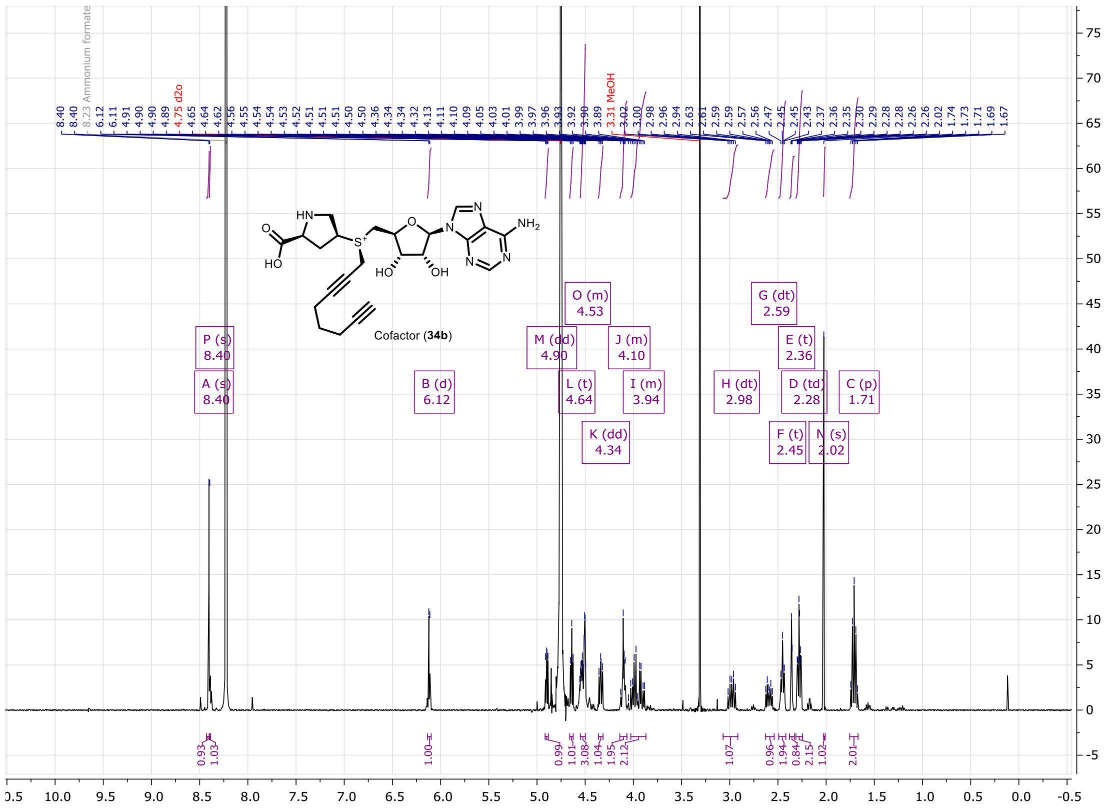


**Figure S100.** $^{1}$H NMR of compound **34b** (400MHz, $D_2O$).

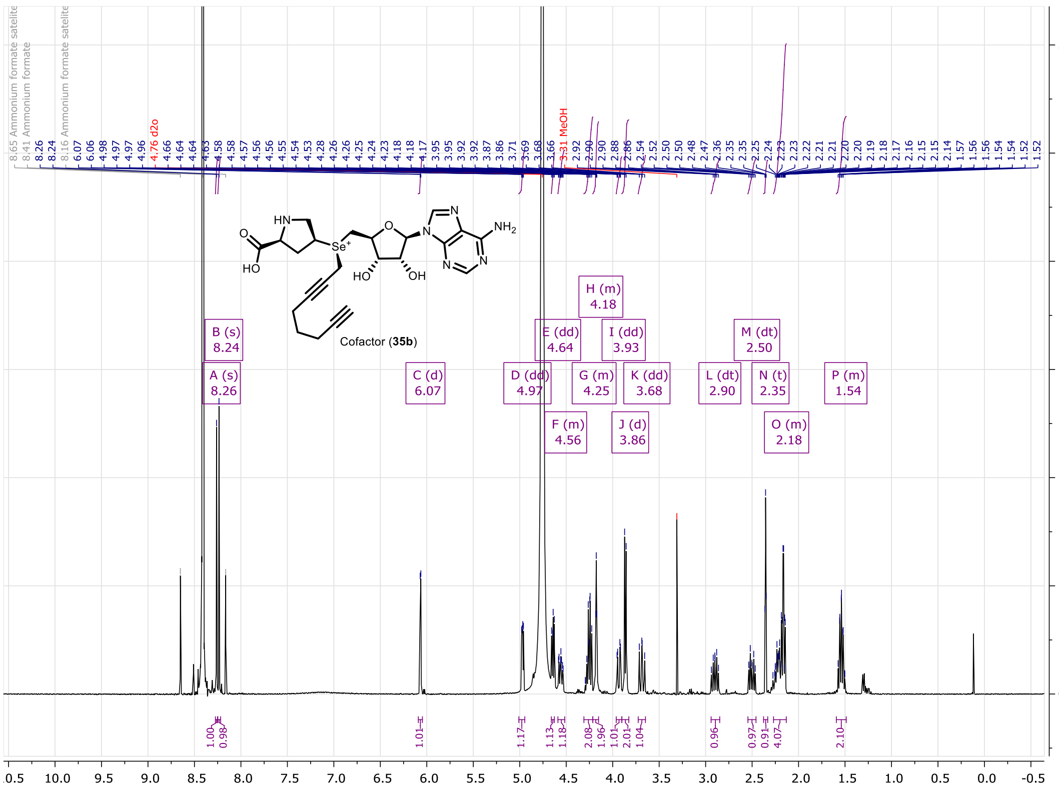

**Figure S101.** $^{1}H$ NMR of compound **35b** (400MHz, $D_2O$).

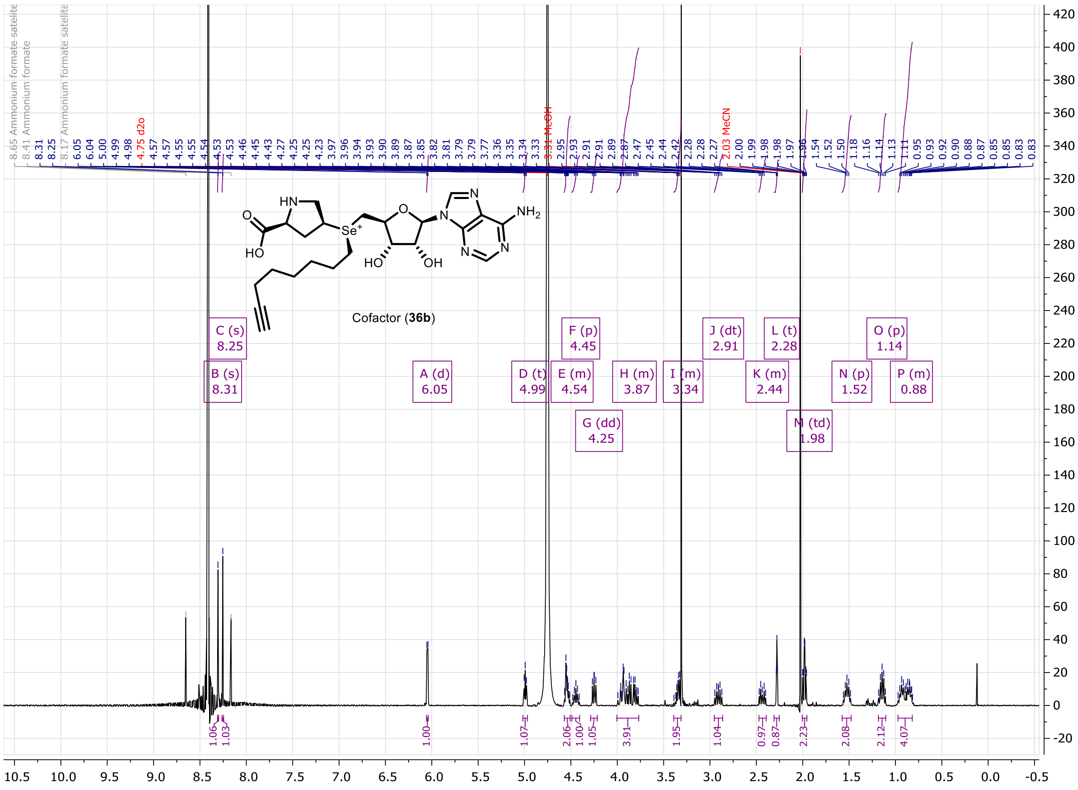

**Figure S102.** $^{1}H$ NMR of compound **36b** (400MHz, $D_2O$).

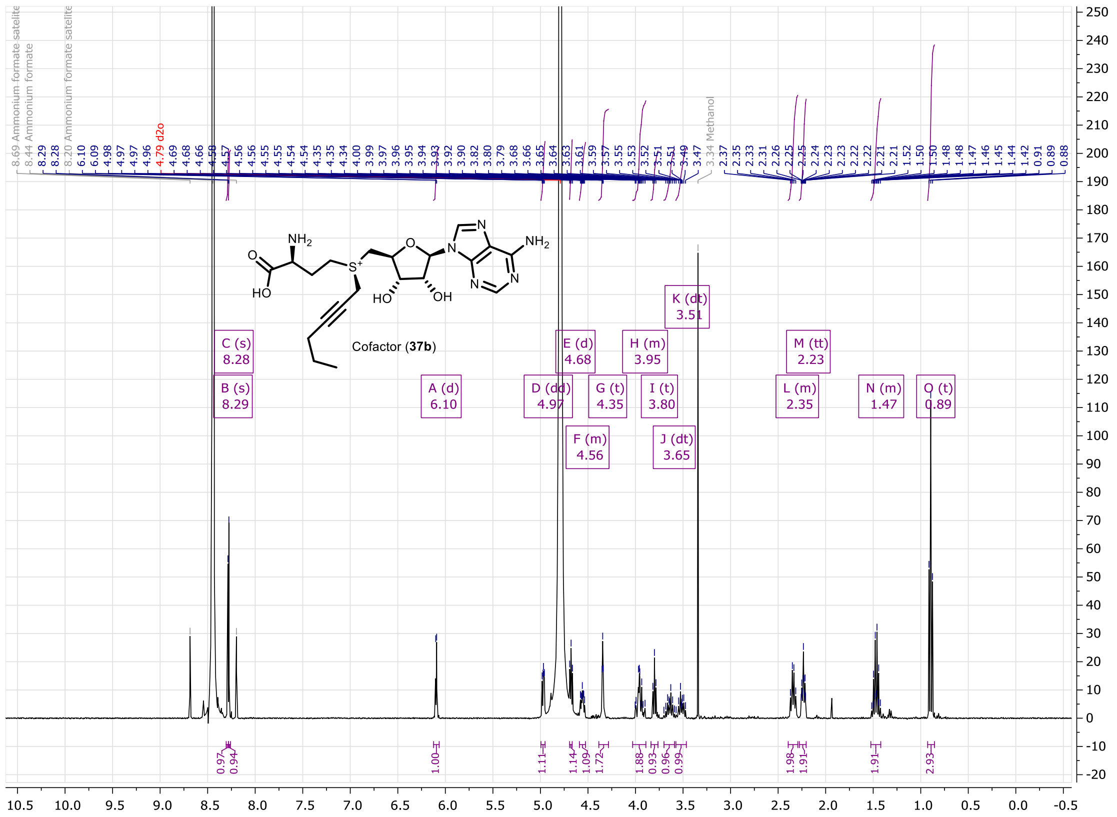

**Figure S103.** $^1$H NMR of compound **37b** (400MHz, $D_2O$).

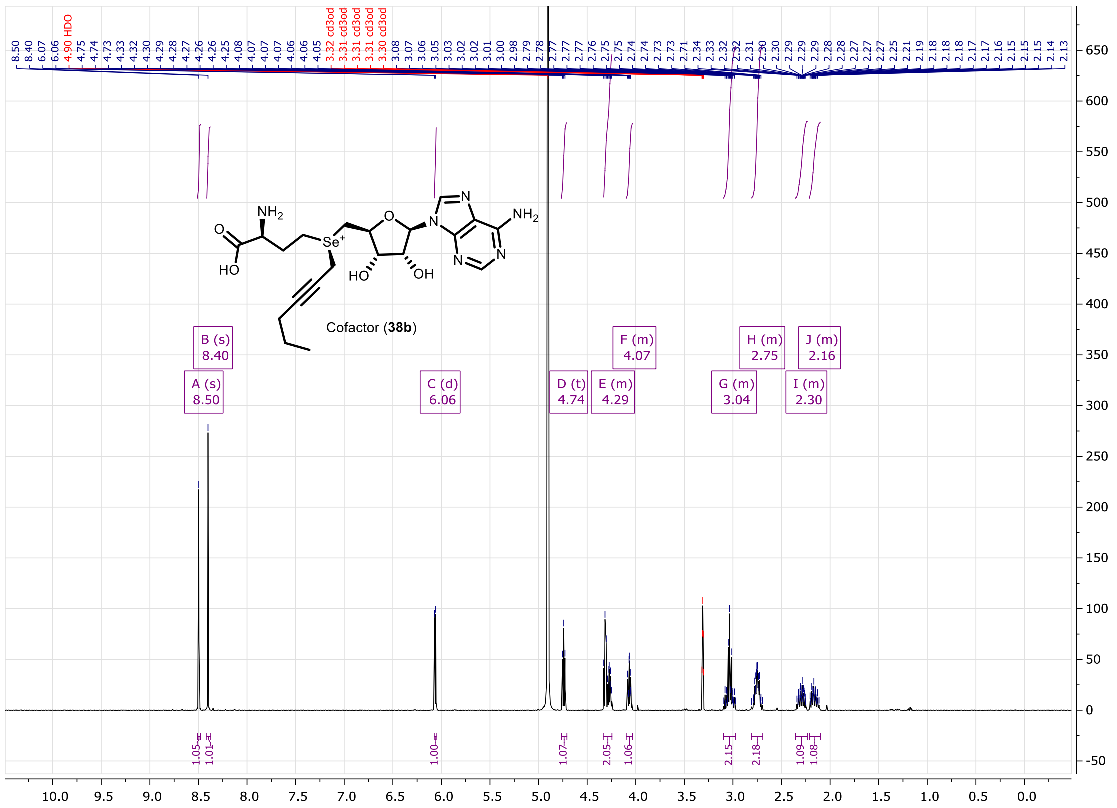

**Figure S104.** $^1$H NMR of compound **38b** (400MHz, $D_2O$).

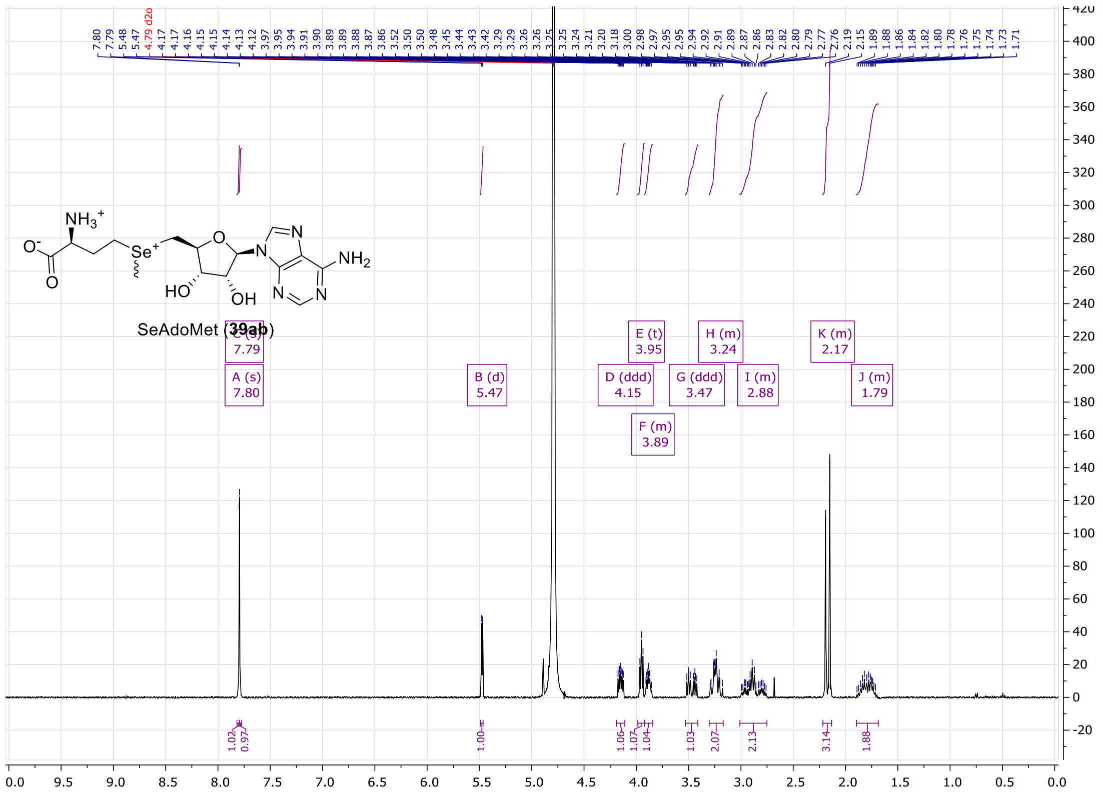

**Figure S105.** $^{1}$H NMR of compound **39ab** (400MHz, $D_2O$).

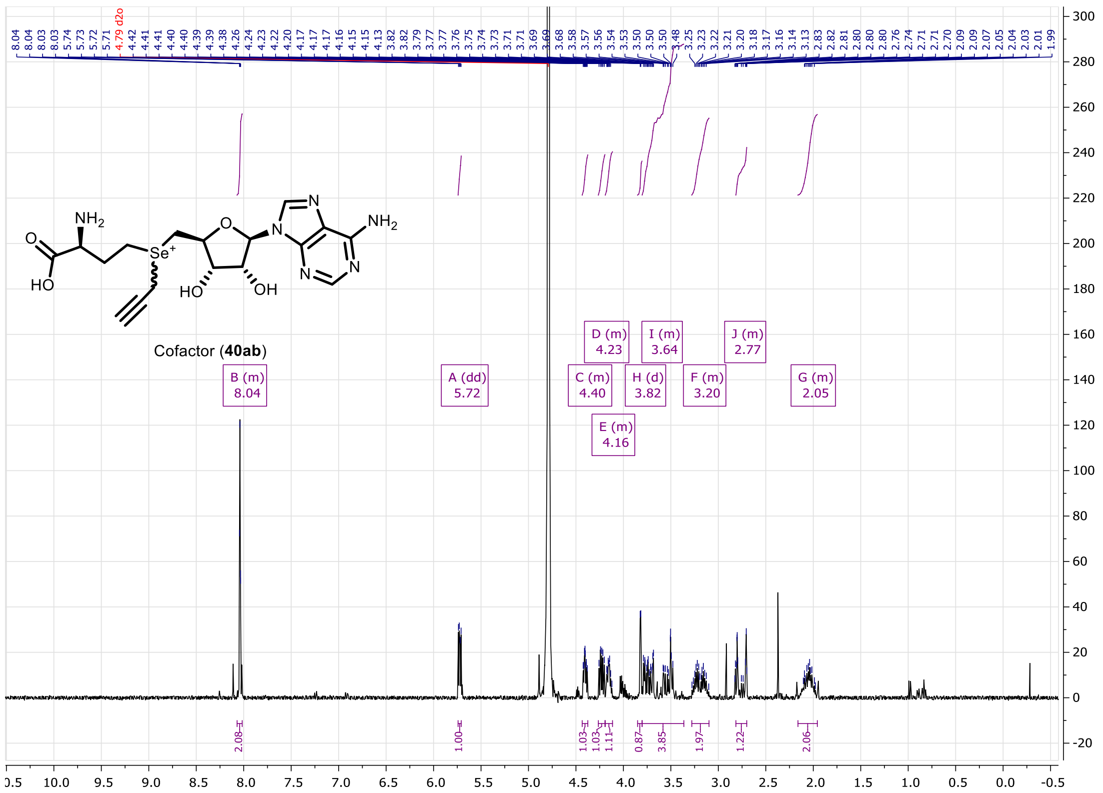

**Figure S106.** $^{1}$H NMR of compound **40ab** (400MHz, $D_2O$).

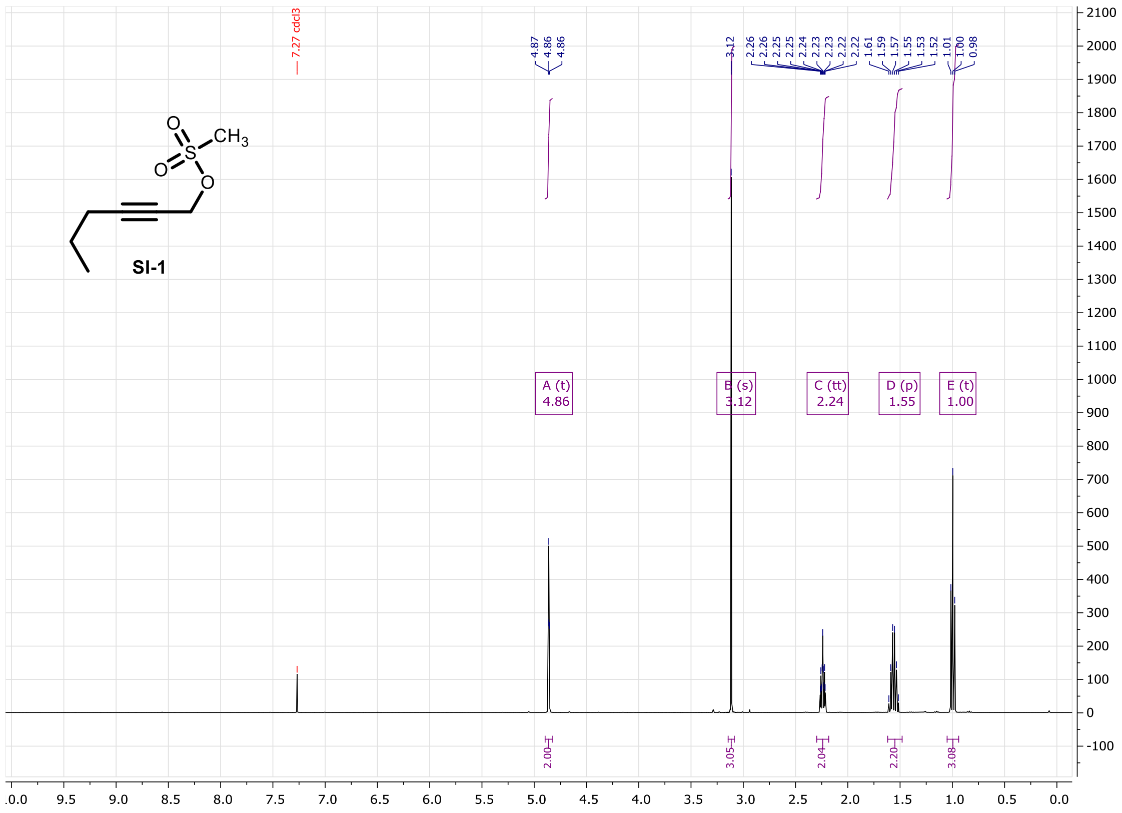


**Figure S107.** $^{1}H$ NMR of compound **SI-1** (400MHz, $CDCl_3$).

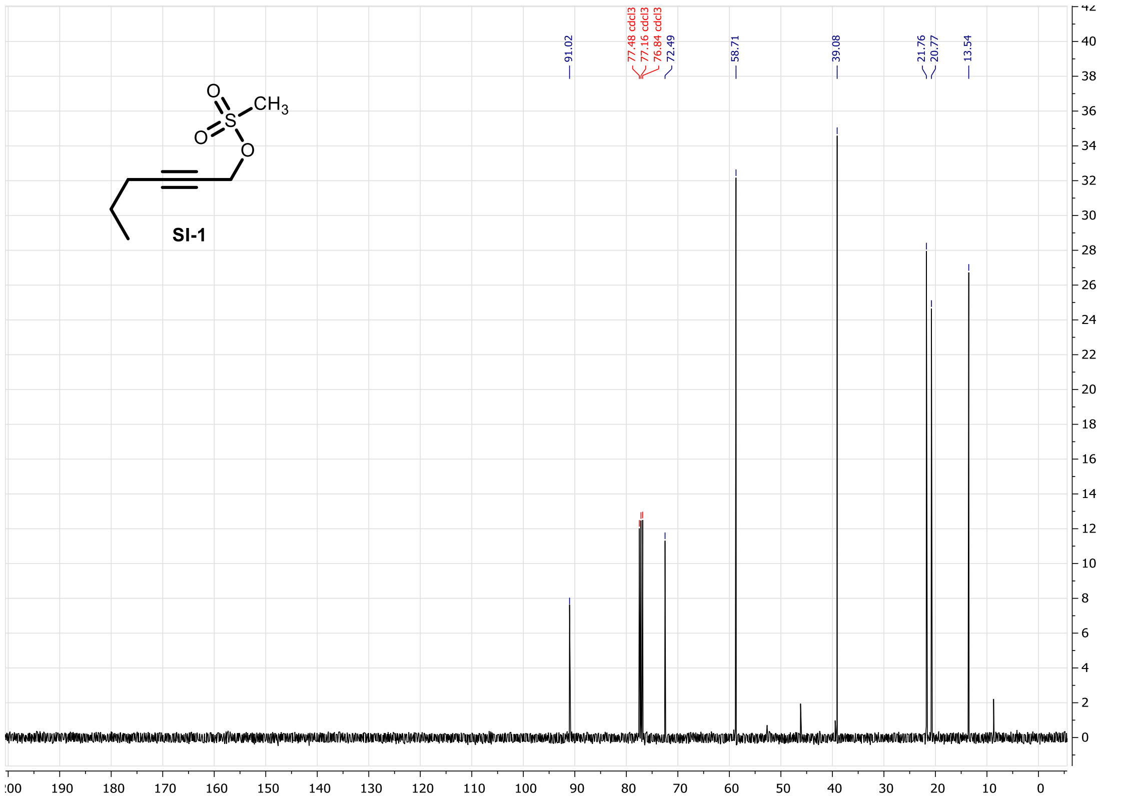


**Figure S108.** $^{13}C$ NMR of compound **SI-1** (100MHz, $CDCl_3$).

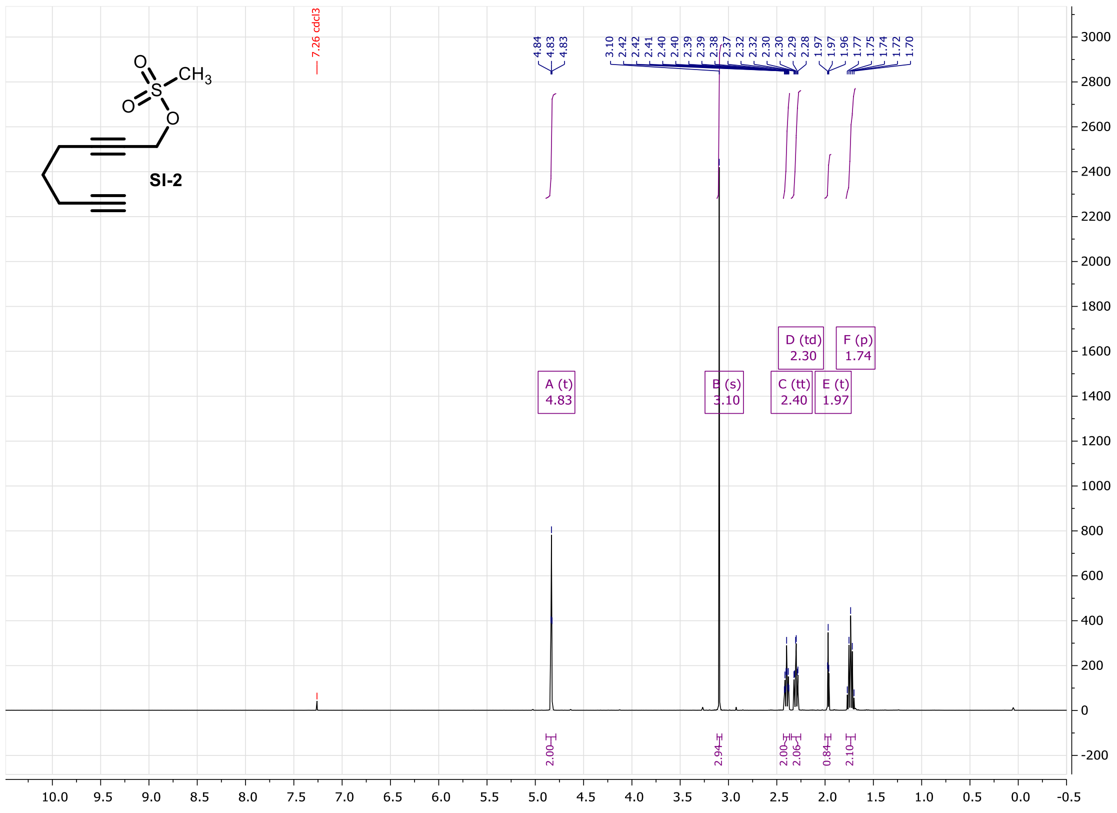

**Figure S109.** $^1$H NMR of compound **SI-2** (400MHz, $CDCl_3$).

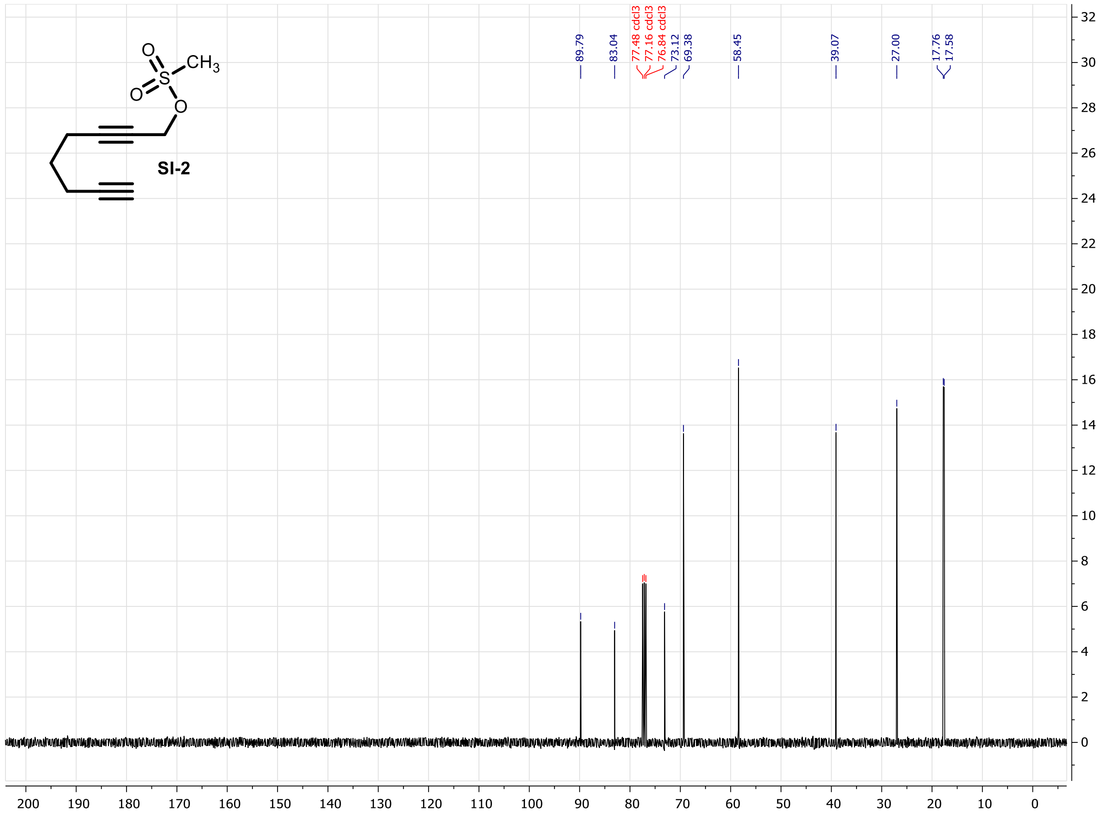

**Figure S110.** $^{13}$C NMR of compound **SI-2** (100MHz, $CDCl_3$).

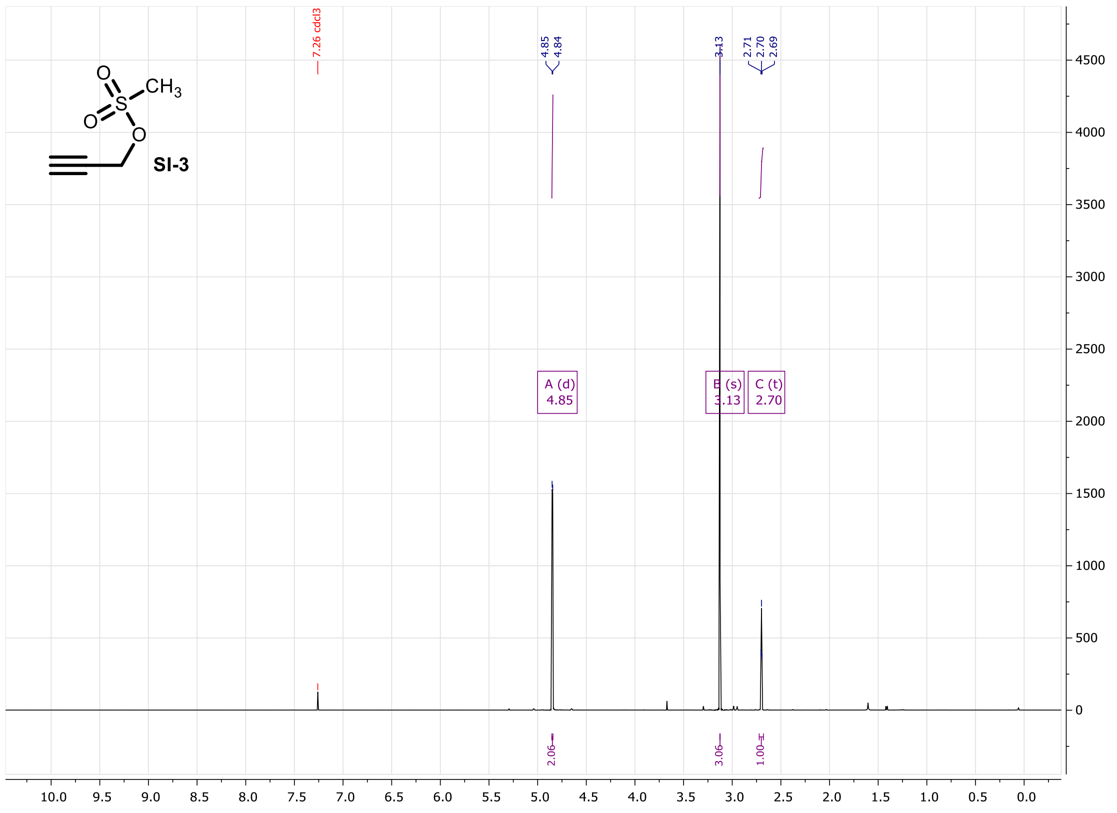


**Figure S111.** $^{1}$H NMR of compound **SI-3** (400MHz, $CDCl_3$).

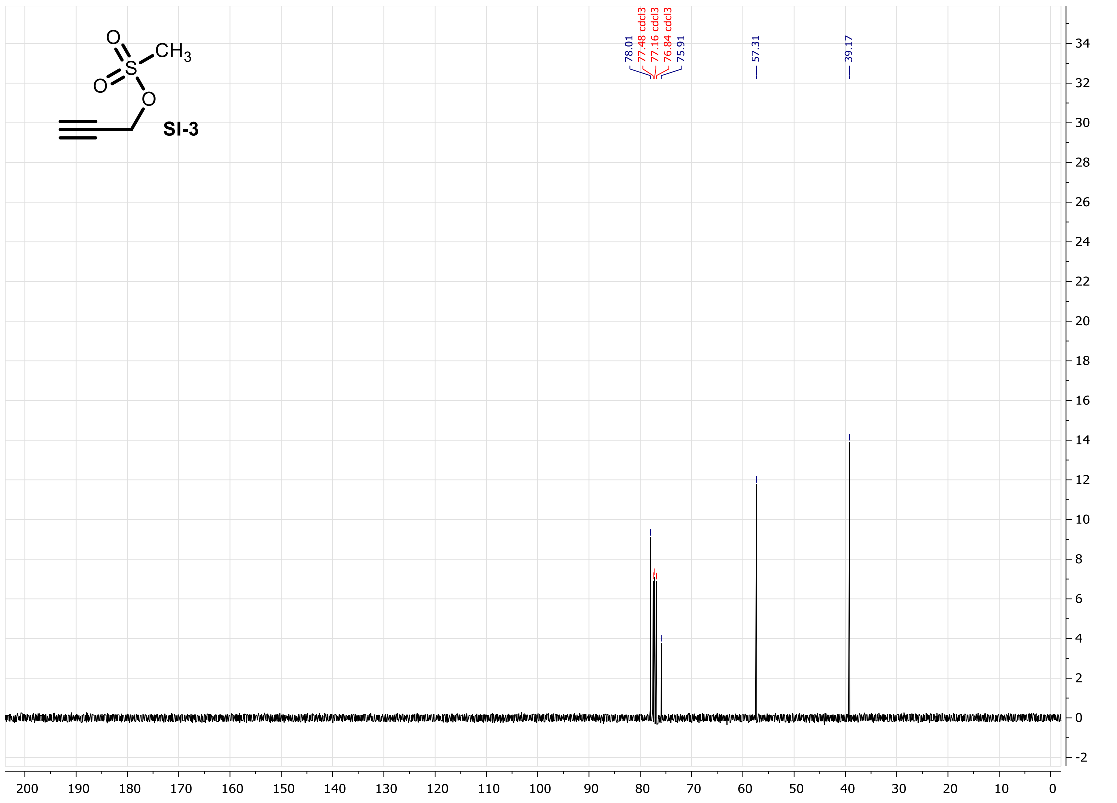


**Figure S112.** $^{13}$C NMR of compound **SI-3** (100MHz, $CDCl_3$).

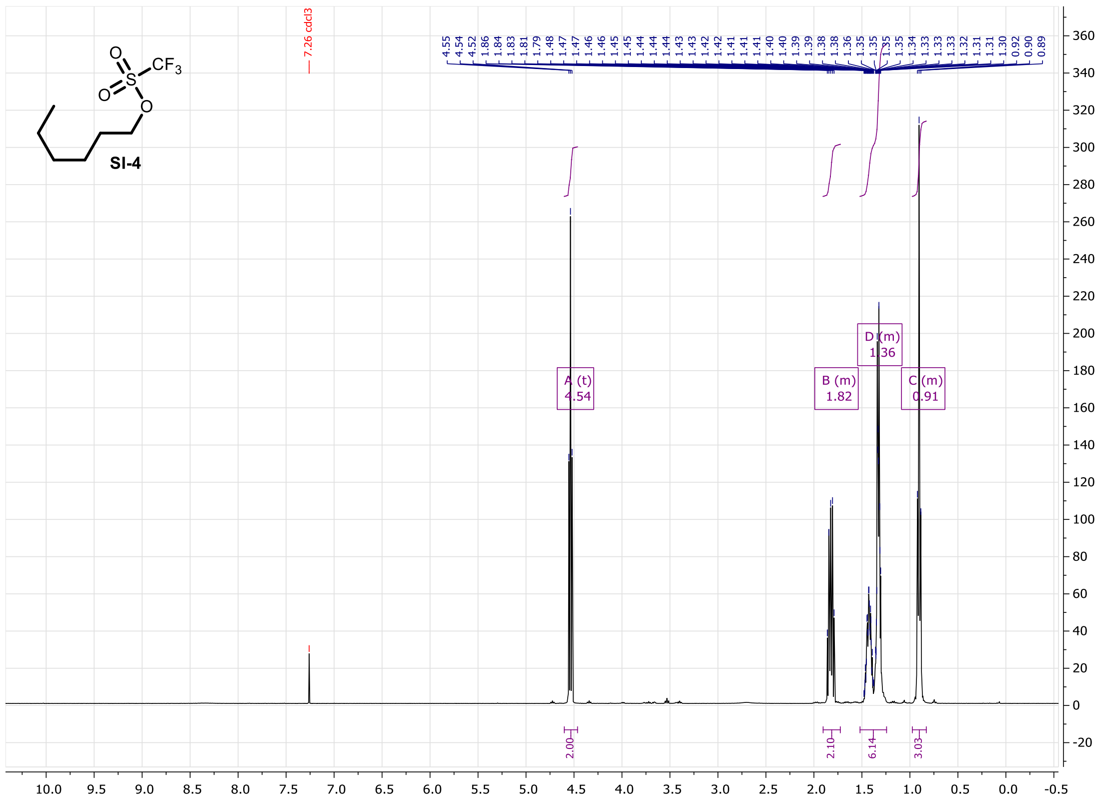

**Figure S113.** $^1$H NMR of compound **SI-4** (400MHz, $CDCl_3$).

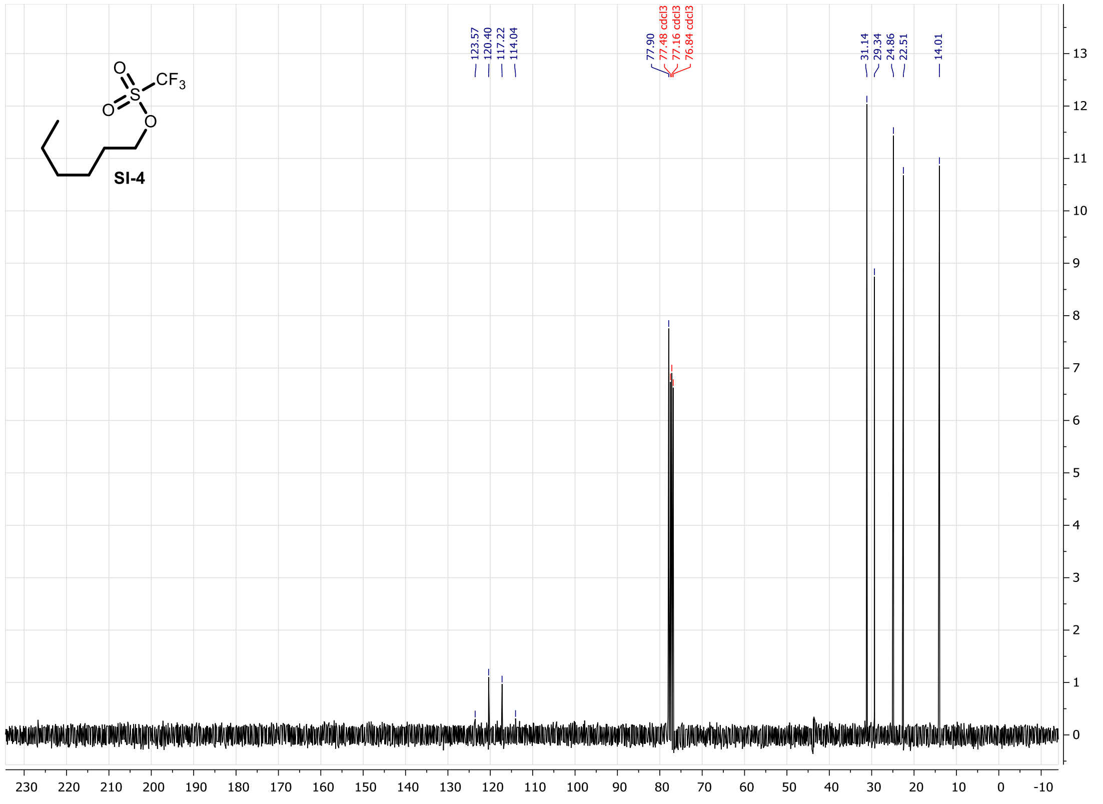

**Figure S114.** $^{13}$C NMR of compound **SI-4** (100MHz, $CDCl_3$).

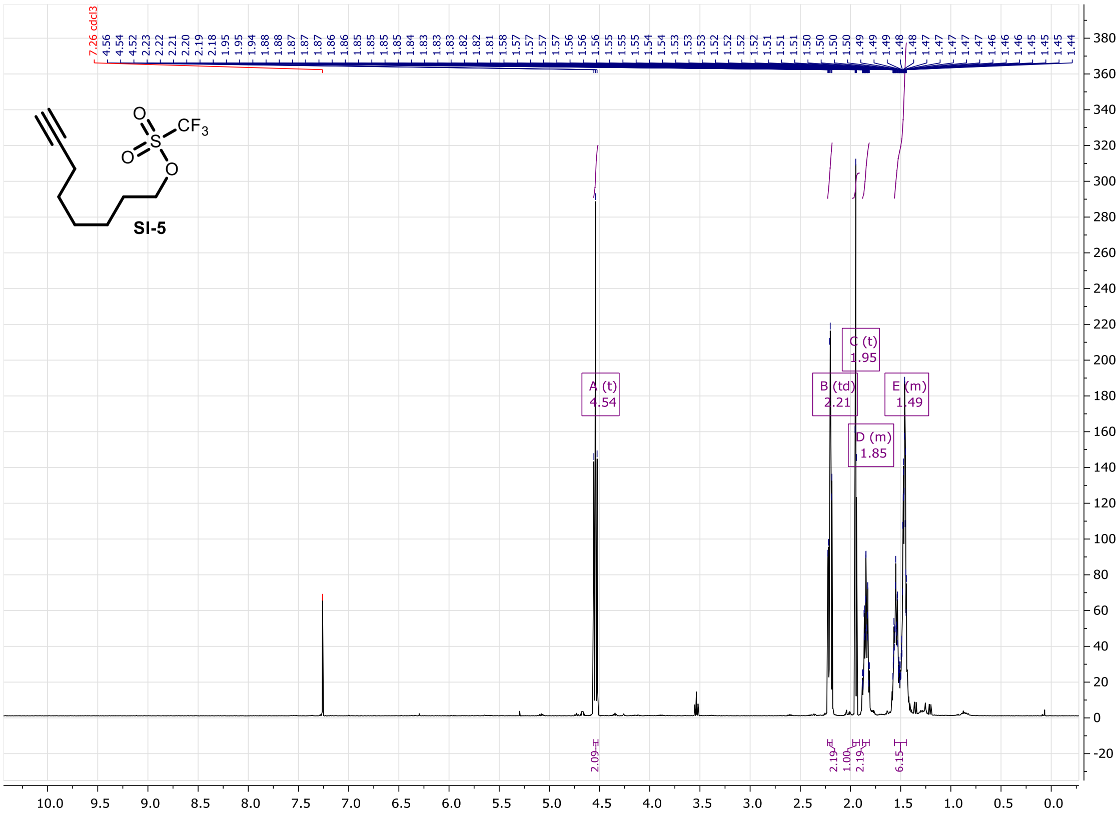


**Figure S115.** $^{1}H$ NMR of compound **SI-5** (400MHz, $CDCl_3$).

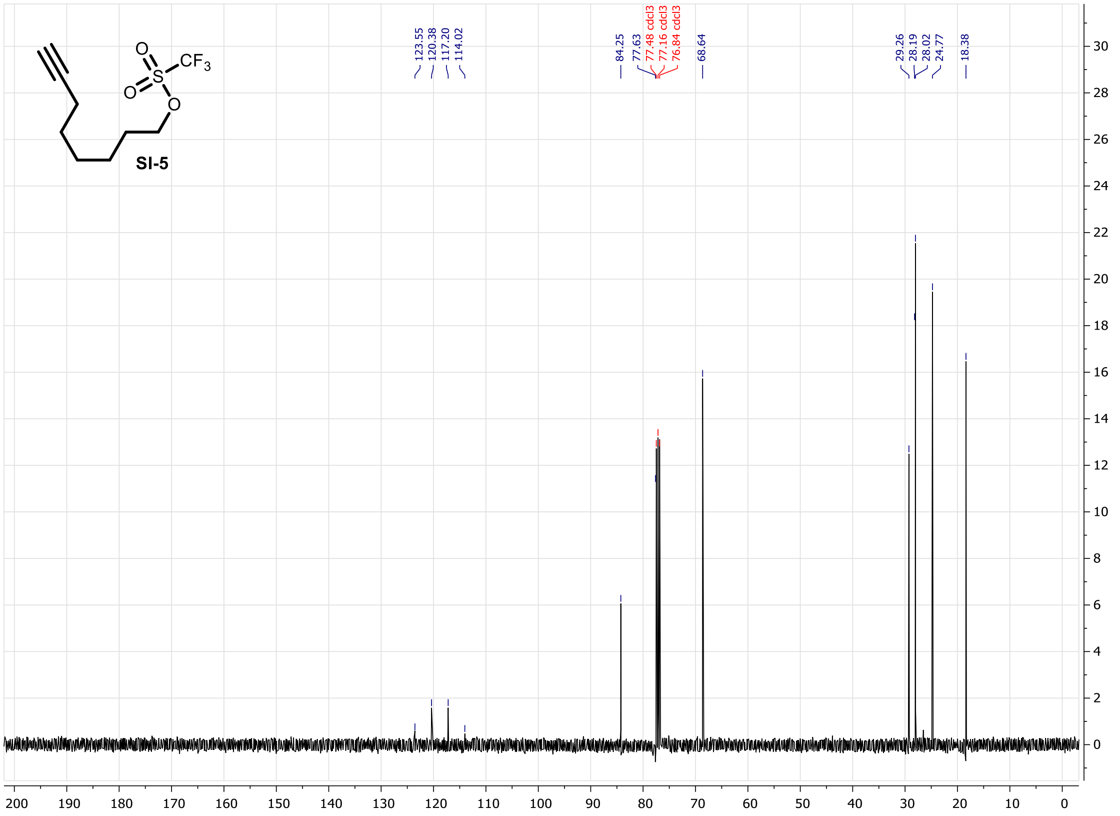


**Figure S116.** $^{13}C$ NMR of compound **SI-5** (100MHz, $CDCl_3$).

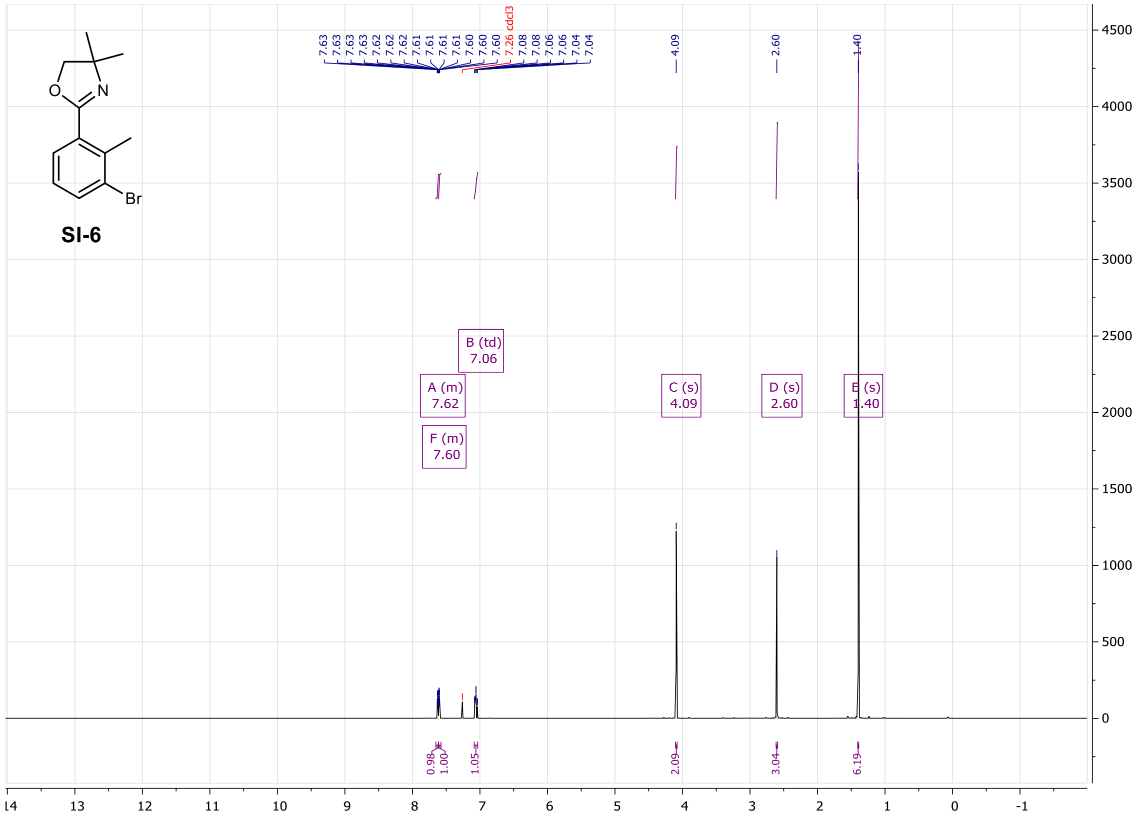


**Figure S117.** $^{1}$H NMR of compound **SI-6** (400MHz, $CDCl_3$).

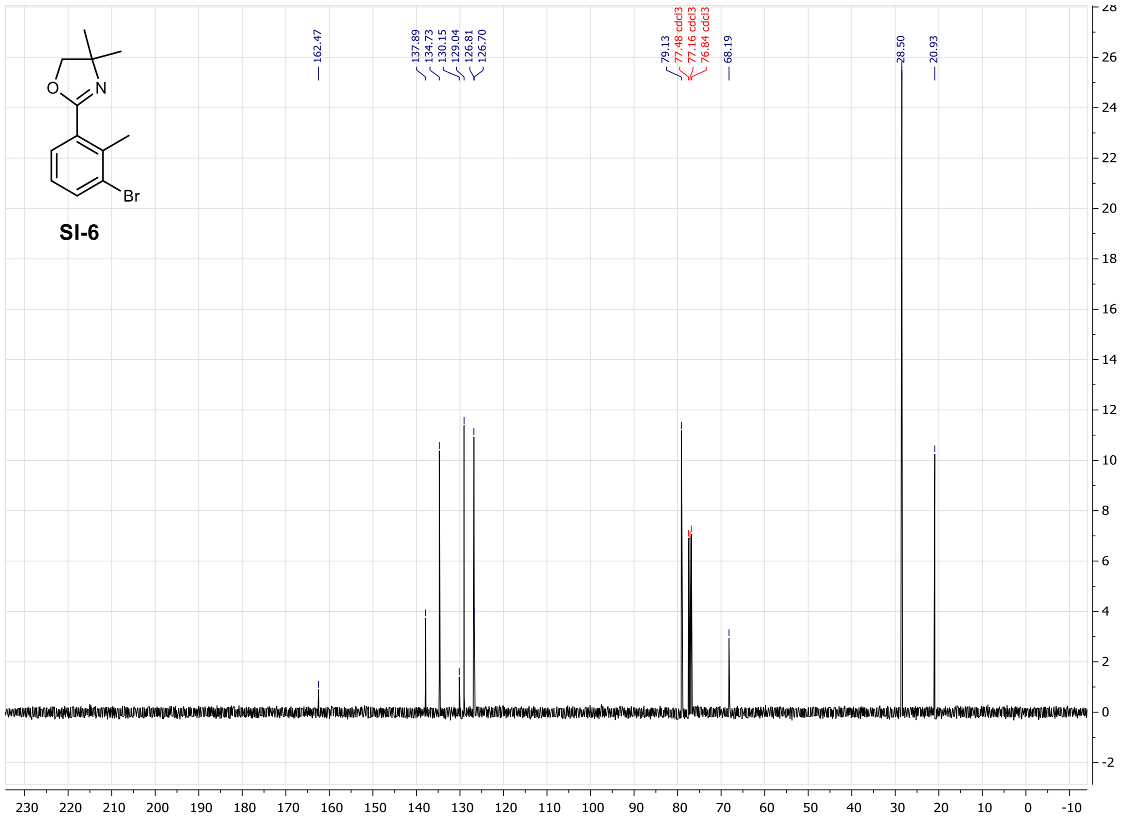


**Figure S118.** $^{13}$C NMR of compound **SI-6** (100MHz, $CDCl_3$).

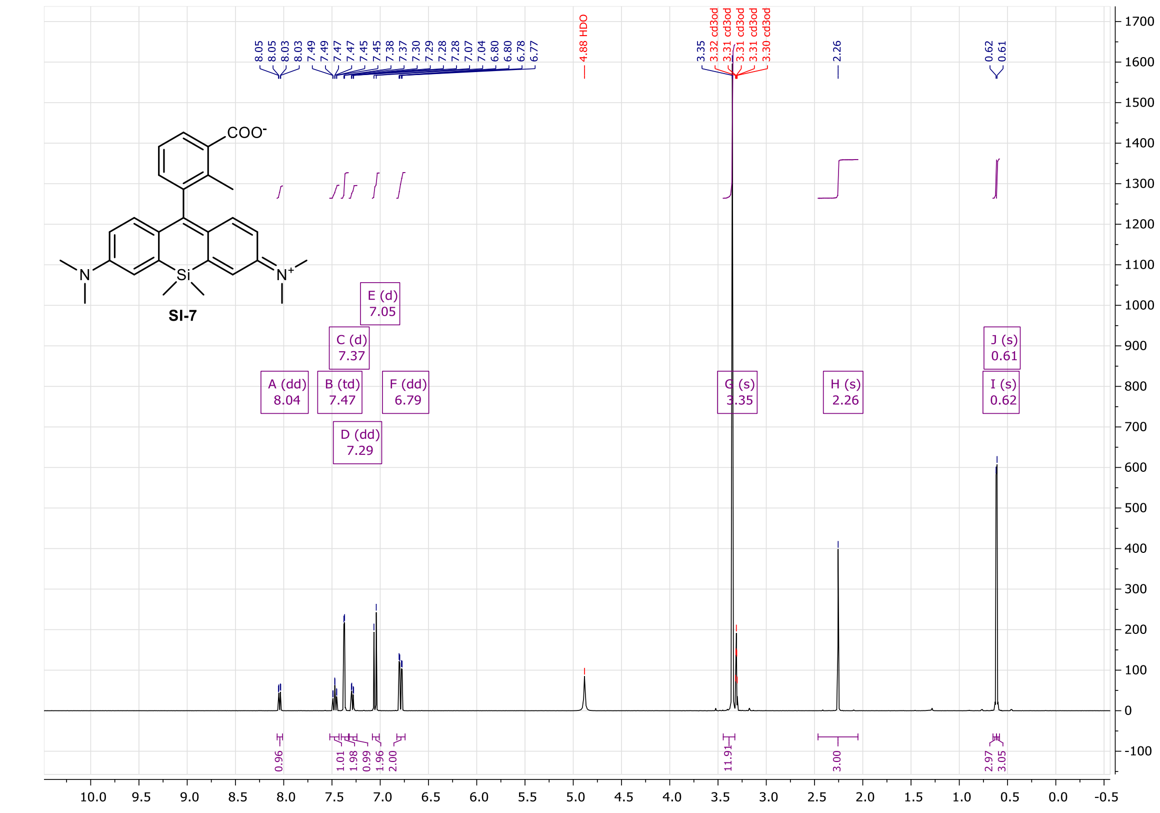


**Figure S119.** $^{1}$H NMR of compound **SI-7** (400MHz, $CD_3OD$).

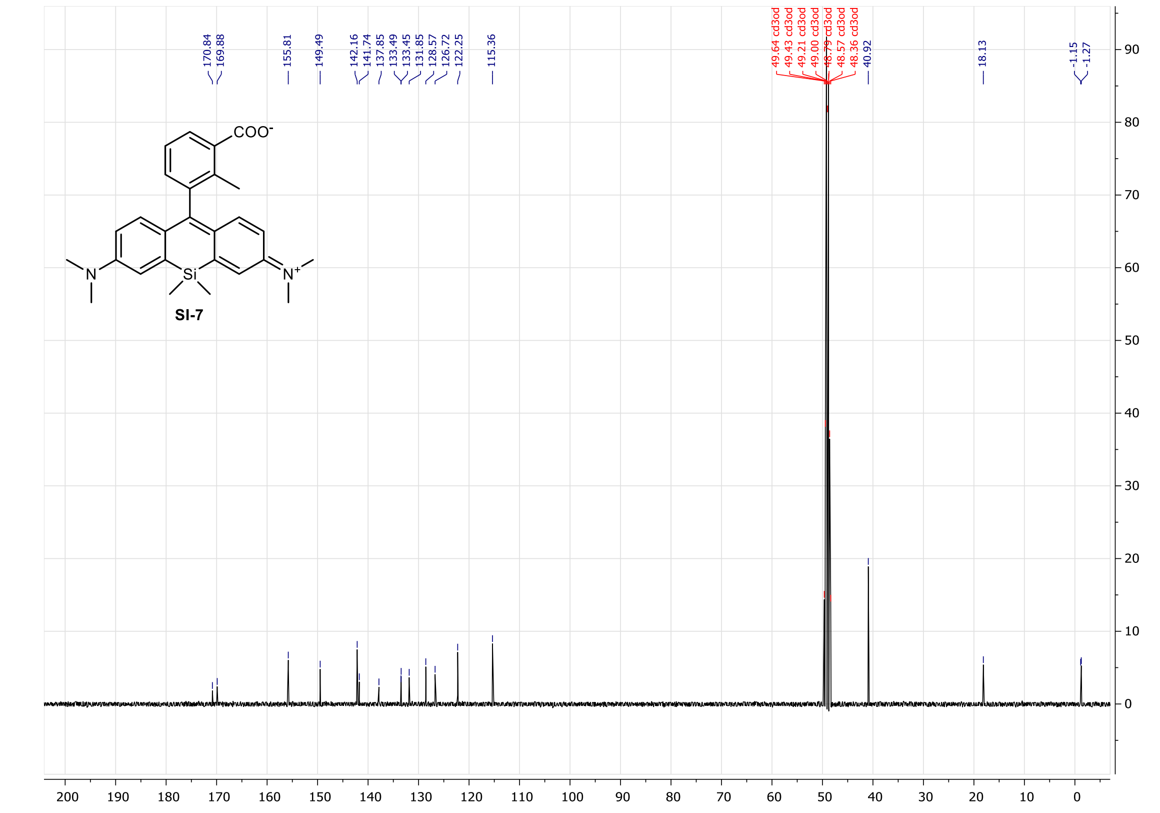


**Figure S120.** $^{13}$C NMR of compound **SI-7** (100MHz, $CD_3OD$).

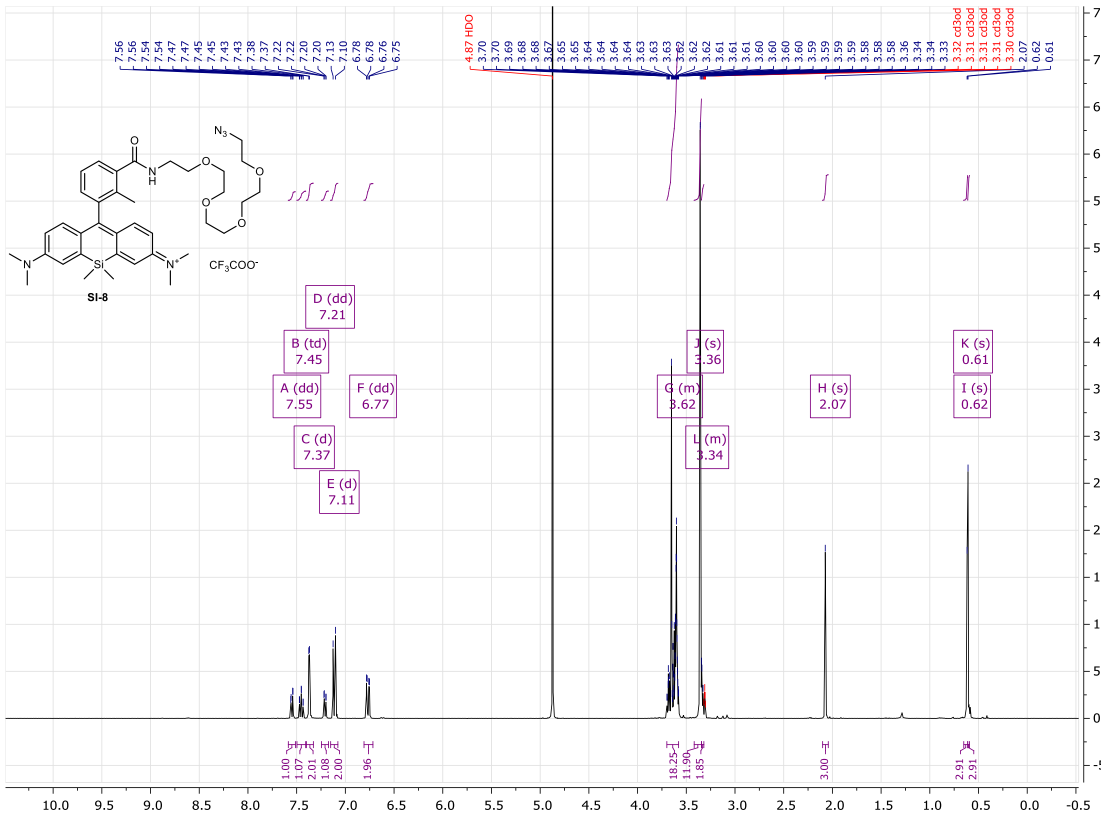

**Figure S121.** $^{1}$H NMR of compound **SI-8** (400MHz, $CD_3OD$).

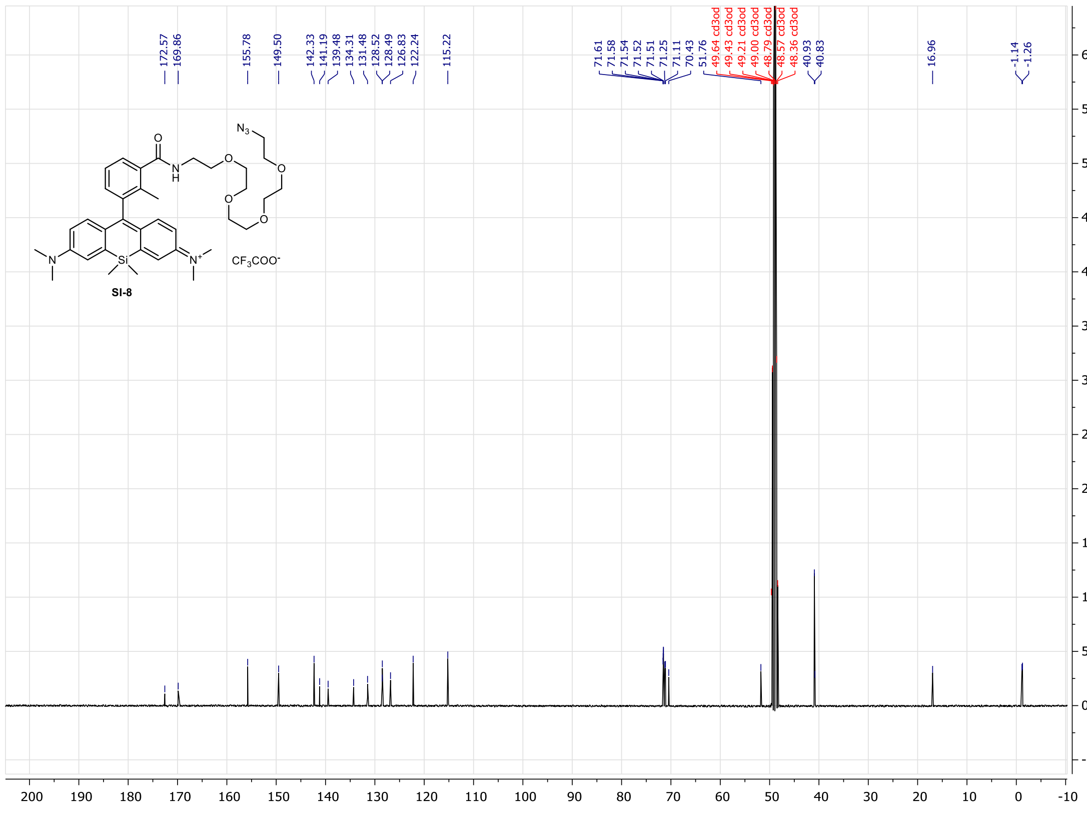

**Figure S122.** $^{13}$C NMR of compound **SI-8** (100MHz, $CD_3OD$).

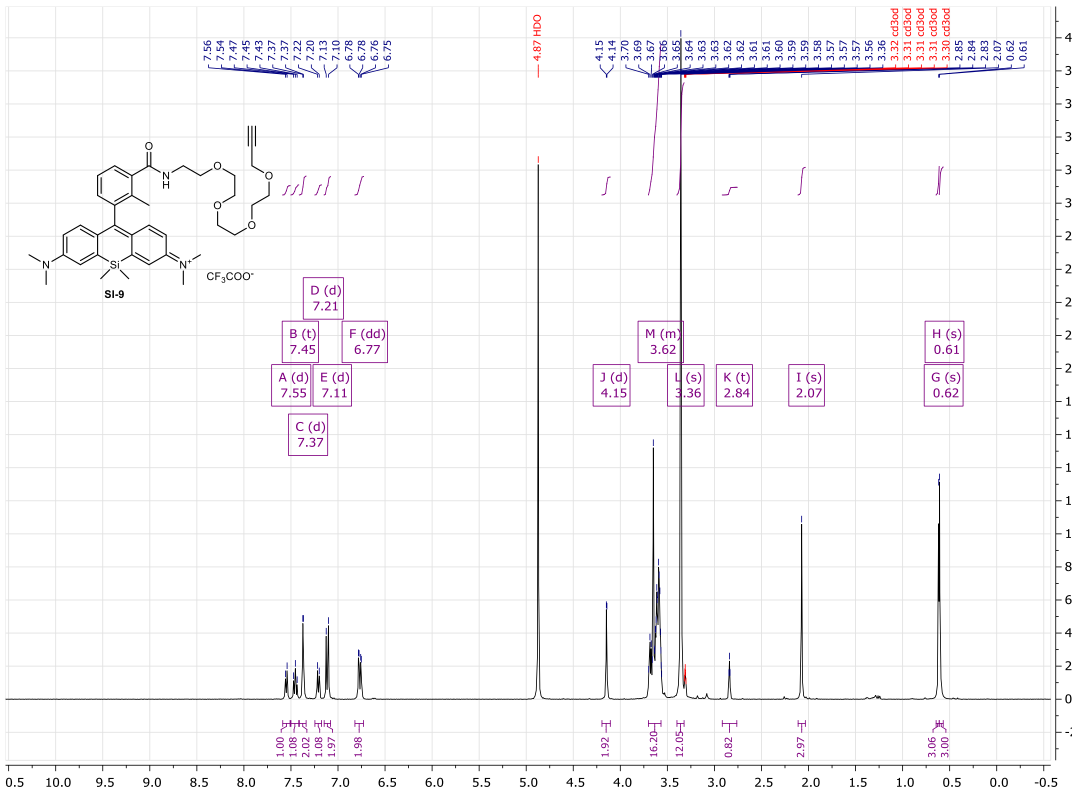

**Figure S123.** $^1$H NMR of compound **SI-9** (400MHz, $CD_3OD$).

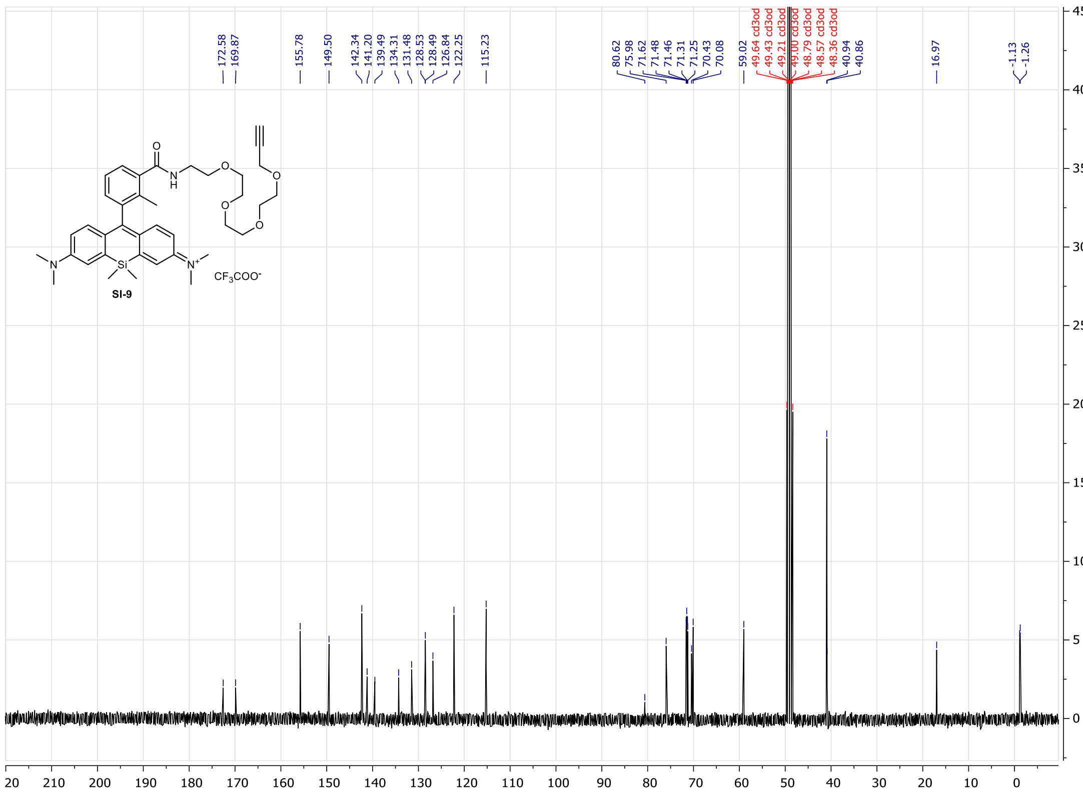

**Figure S124.** $^{13}$C NMR of compound **SI-9** (100MHz, $CD_3OD$).

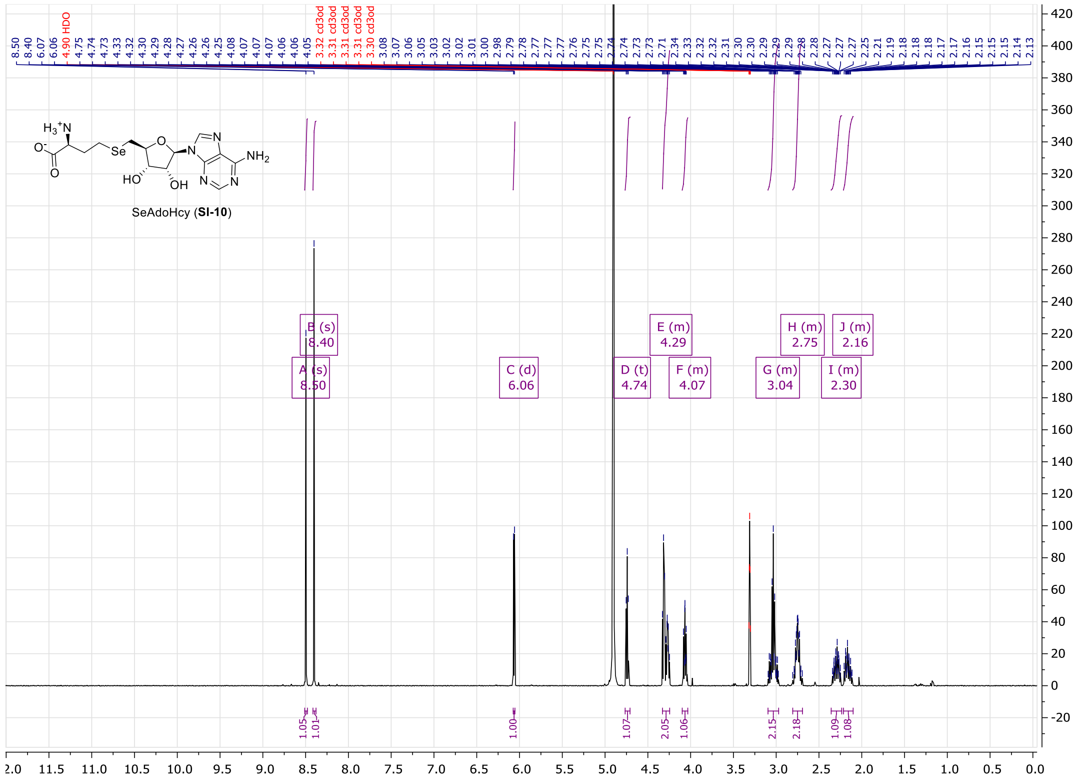

**Figure S125.** $^{1}$H NMR of compound **SI-10** (400MHz, $CD_3OD$).

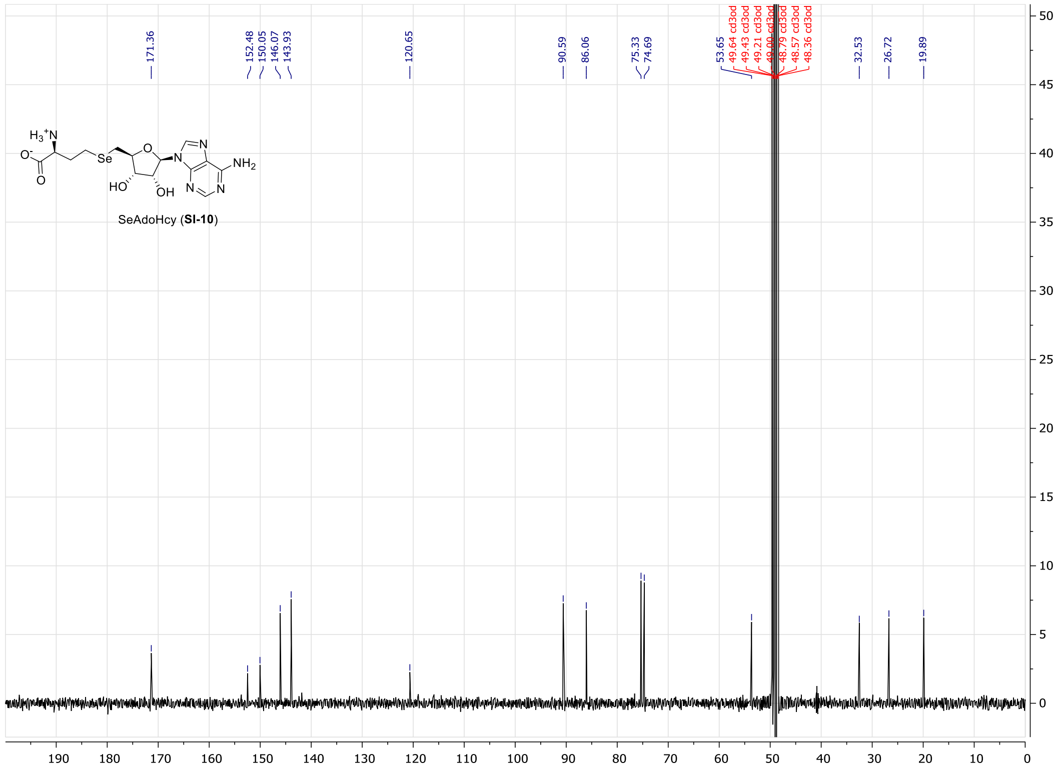

**Figure S126.** $^{13}$C NMR of compound **SI-10** (100MHz, $CD_3OD$).
